# The ecogenomics of dsDNA bacteriophages in feces of stabled and feral horses

**DOI:** 10.1101/2020.07.24.219436

**Authors:** V. V. Babenko, A. Millard, E. E. Kulikov, N.N. Spasskaya, M. A. Letarova, D. N. Konanov, I. Sh. Belalov, A.V. Letarov

**Affiliations:** FSC Physico-chemical medicine FMBA, Russia; Dept Genetics and Genome Biology, University of Leicester, UK; Winogradsky institute of microbiology RC Biotechnology RAS, Moscow, Russia; Zoology museum, Faculty of biology, Lomonosov Moscow state university, Russia; Faculty of biology Lomonosov Moscow state university, Russia

## Abstract

The viromes of the mammalian lower gut were shown to be heavily dominated by bacteriophages; however, only for humans were the composition and intervariability of the bacteriophage communities studied in depth. Here we present an ecogenomics survey of dsDNA bacteriophage diversity in the feces of horses (*Equus caballus*), comparing two groups of stabled horses, and a further group of feral horses that were isolated on an island. Our results indicate that the dsDNA viromes of the horse feces feature higher richness than in human viromes, with more even distribution of genotypes. No over-represented phage genotypes, such as CrAssphage-related viruses found in humans, were identified. Additionally, many bacteriophage genus-level clusters were found to be present in all three geographically isolated populations. The diversity of the horse intestinal bacteriophages is severely undersampled, and so consequently only a minor fraction of the phage contigs could be linked with the bacteriophage genomes. Our study indicates that bacteriophage ecological parameters in the intestinal ecosystems in horses and humans differ significantly, leading them to shape their corresponding viromes in different ways. Therefore, the diversity and structure of the intestinal virome in different animal species needs to be experimentally studied.

**Short abstract (needed in some journals as eLife):** The viromes of the mammalian gut were shown to be heavily dominated by bacteriophages; however, only for humans were the composition and intervariability of the bacteriophage communities studied in depth. Here we present an ecogenomics survey of dsDNA bacteriophage diversity in the feces of horses (*Equus caballus*), comparing stabled horses, and feral horses that were isolated on an island. The viromes equine fecal viromes feature higher richness than in human viromes, with more even distribution of genotypes. No over-represented phage genotypes were identified. Additionally, many bacteriophage genus-level clusters were found to be present in geographically isolated populations. Only a minor fraction of the phage contigs could be linked with the bacteriophage genomes. Our study indicates that bacteriophage ecological parameters in the intestinal ecosystems in horses and humans differ significantly, leading them to shape their corresponding viromes in different ways.

**Importance. (needed for mBio):** The study presents the first in depth analysis of the composition and variability of the gut dsDNA bacteriophage community in the mammalian species, other than humans. The study demonstrates that the bacteriophage ecology in the gut is substantially different in different animal species. The results also indicate that the genetic diversity of the equine intestinal bacteriophages is immense and almost totally unexplored by the moment.

## Introduction

The existence of microbial populations inhabiting different niches of the human and other animal bodies was first observed by Antony van Leeuwenhoek in 17^th^ century (1) and has since become a commonly accepted paradigm that is mentioned in almost any microbiology textbook. Remarkable progress has been made in this field over the last 15 years due to the introduction of the culture-independent tools for the analysis of the composition and function of the microbial component of the human (2) or animal holobiont (3). Much emphasis has been placed on the gut microbiome, representing the largest microbial community associated with humans or other mammalian bodiesies The intestinal microbiome is now considered as a “new organ”, exerting strong and multifaceted influence over the physiology of the macro-host (4). The gut microbiome is involved in the pathology of numerous conditions, including Crohn disease (5, 6), obesity (7, 8), cancer (9) and even behavioral alterations (10–12).

It is well established that in all vertebrate animals the intestinal microbiome is associated with the corresponding virome - the community of the viruses infecting or produced by the microorganisms comprising the bacterial microbiome (13–16). Although the bulk of the intestinal viromes are comprised of bacteriophages (13, 14, 17), these viral communities are also believed to be involved in multiple physiological effects and pathological processes *via* alteration of the composition and activity of the microbial community, and through direct interaction with the macro-host tissues and immune system (14, 18–20).

Bacteriophage diversity, biogeography and dynamics in the human gut has been investigated in depth in numerous studies using metagenomic approaches (16, 17, 21); see also reviews (17, 22, 23). It has to be mentioned, however, that in almost all of the studies the viral community of the feces was used as a proxy of the intestinal viromes. The “normal” composition of the human fecal bacteriophage community has been established and the “core phageome” composition defined as bacteriophage genotypes present in more than 50% individuals worldwide was evaluated (24). The first identified core phage lineage, named CrAssphage, that is highly prevalent in some of the samples (up to 90% of the viral reads) was initially identified using bioinformatic approaches (25) and was later cultured and shown to be a large podovirus infecting *Bacteroides* (26).

Therefore, the main characteristics of healthy human viromes have been established as follows: a diverse community, that is highly stable in time (17, 21), highly individual with larger inter-individual distances compared to different time points (16, 17, 27) even if the dietary interventions were applied (21). Human viromes are suggested to be dominated by temperate bacteriophages (27) although the prevalence of the contigs containing integrases or site-specific recombinases genes is found to vary greatly (0-68%) between individual viral metagenomes (17).

Despite significant progress in the understanding of bacteriophage ecology in the human gut, the data on other animal species are scarce. Although a significant number of metagenomic datasets from various species have been published (28–31), the vast majority of these studies focus on detection and interpretation of the animal viruses sequences, and bacteriophages have not been given significant attention. Only a few studies give emphasis to bacteriophage diversity in these samples, although this is limited to identifying differences between health and disease states in rhesus monkeys (32, 33) or specifically focusing on the diversity of ssDNA viruses (34).

In the present work we focus on the ecogenomics of dsDNA viruses present in horse feces. The equine intestinal microbiome plays an essential role in animal nutrition, allowing the horse to digest cellulose which is the major component of the grass consumed (35). In contrast to ruminants where microbial cellulose digestion takes place in the forestomach (rumen), in horses the cellulolytic microbial community develops in the cecum and large intestine that have a cumulative volume of about 100 L with food retention time about 48-72 hours (36). The large intestine content is not subjected to any subsequent digestion (such as in ruminants) and is pushed by peristalsis into the rectum where it is subjected to partial dehydration to form the feces (37). The average time intervals between food intake or between the defecation acts in horses are much shorter than the indicated retention time (38). Therefore, the horse large intestine functions as a natural chemostat with highly stable physical and chemical conditions and fairly constant flow through. Adult horses do not show any coprophagy, but at the same time they do not avoid contact with feces of other individuals or fecally contaminated objects (38), potentially enhancing the exchange of bacteria and viruses between the individual viromes.

Only a few studies have been dedicated to horse intestinal bacteriophages. A limited Sanger-based metagenomic analysis of a single sample allowed the estimation of richness of the viral community, finding 1200 bacteriophage genotypes (39). A more recent metagenomic study compared fecal microbiomes and viromes of cattle and horses held on the same farm (40). However, the amount of data for each virome in this study was limited and no information concerning the specific characteristics of the bacteriophage communities of the samples was reported. There have also been several studies of the horse gut bacteriophage community based on other approaches. In a limited electron-microscopy study of horse feces almost all VLPs identified were classified as tailed phages, with 69 morphologically distinct types reported out of <200 particles measured, indicating a high level of diversity (see (13) for review of earlier work). A comprehensive study of coliphage diversity and dynamics (41) in the feces of four horses held in the same location suggested high prevalence of virulent coliphages.. The *E. coli* host population was found to be highly divergent, and represented by hundreds of strains simultaneously present in the same sample. The overlap of the sensitivity of these strains to co-occuring bacteriophages was limited (41, 42) with ~1-5 % of the total *E. coli* counts being suitable hosts for any particular phage isolate. The data of Golomidova et al. (41, 43) and the results of the longitudinal study of G7C-related bacteriophages persistence and evolution within the ecosystem of a horse stable (44) indicated the flow of the coliphage genotypes between the animals. However, *E. coli* and its phages make for only a minor fraction of the total quine microbiome and viromes respectively. Currently, to the best of our knowledge, no methods exist that allow the translation of the findings made using this model system to the total community.

Here we present the ecogenomics of horse fecal dsDNA viromes of three separate horse populations including two groups of stabled horses and one herd of feral horses isolated on an island. Our data indicate that equine intestinal viromes are highly diverse communities dominated by the tailed bacteriophages. Although the site of sampling or/and the life conditions of distinct populations have marked influence over the composition of the individual viromes, it was possible to identify the equine intestinal core virome

## Results

### The sampling strategy, workflow and sequencing results

In order to characterize the virome composition and diversity in horse feces we collected samples from three populations of horses in Russia. These included two groups of stabled animals and one feral population. The stable 1(S1) population was kept at the equestrian center in the city of Moscow. The stable 2 (S2) population was located in the country side ~90 km from Moscow. The lifestyle and diet of these two populations differed significantly (see material and methods). In addition, we sampled from feral horsesat Rostovsky national reserve, that have been isolated on the island for several decades (see material and methods for detail) – population F. In population S1 we sampled four animals, two of which were sampled twice. In the S2 population five animals were sampled a single time (October 2018). In the F population, six animals (two harem stallions and two mares belonging to each of the stallions) were sampled both in May and October 2018 (Table S1)

The viromes were extracted from all samples: viral DNA was extracted and sequenced using IonTorrent technology. It is important to note that the procedure applied for virome isolation (Fig.1) did not include any gradient centrifugation or ultrafiltration steps that may selectively remove some types of viral particles. We also did not use DNA amplification to avoid the biased representation of sequences that can occur. Based on previously published research (39, 45), see also (40) the bulk of the horse intestinal virome is composed of tailed bacteriophages, so we decided to focus on dsDNA viruses.

**Figure 1.**
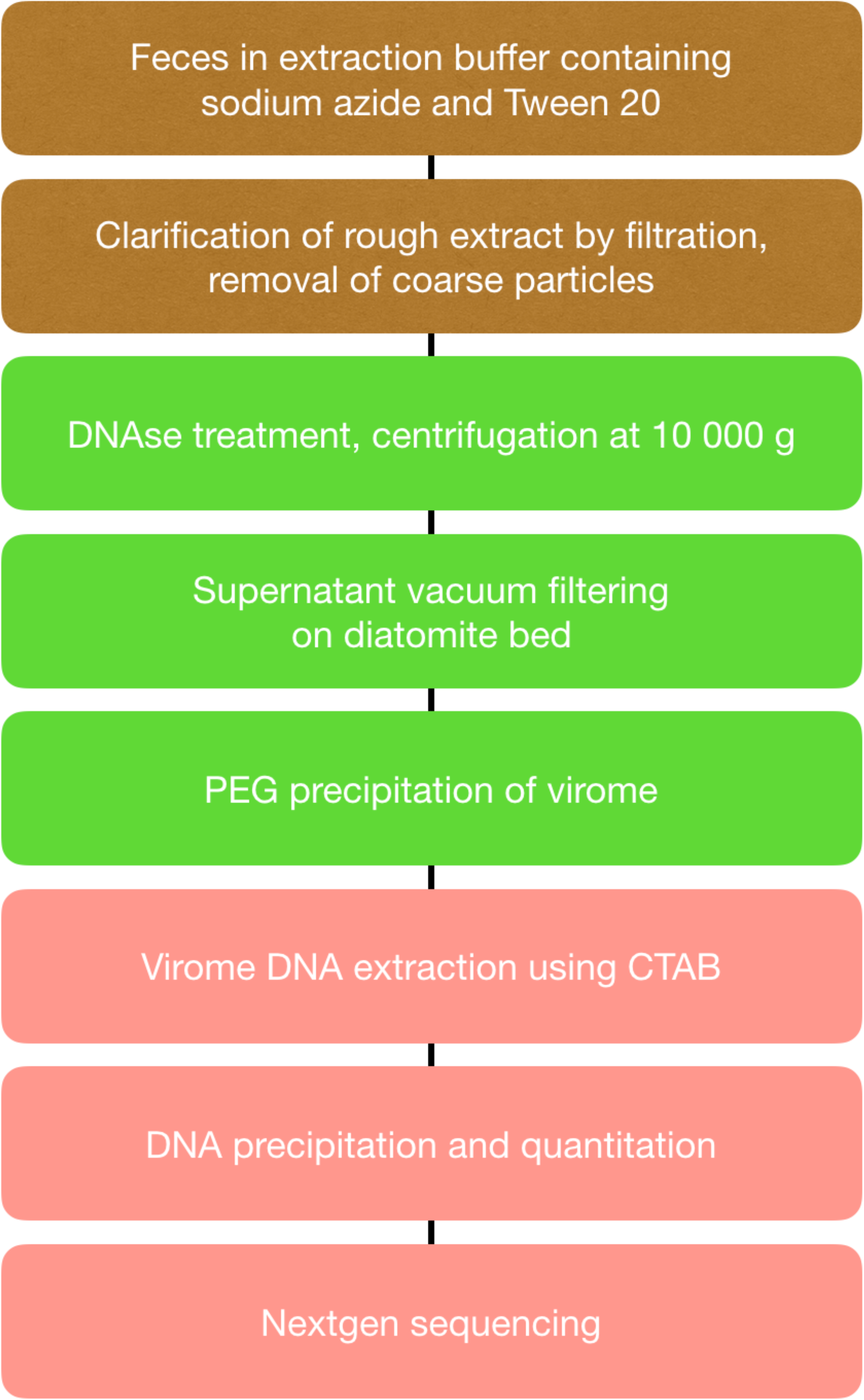
Sample processing workflow

To check for bacterial contamination both virome QC and sortmeRNA were used, and both methods suggest the samples were highly enriched for viral DNA with minimal bacterial contamination. A total of 8097 nonredundant viral contigs >5 kb were identified, and were used for all further analysis. Among these contigs we identified 46 contigs longer than 30 kb. that may represent complete or almost complete phage genomes.

### Complexity of the individual viromes

To estimate the alpha-diversity Shanon and Simpson indexes (Fig. 2) were calculated from relative abundance (how) and revealed that individual virome diversity tends to be higher in feral horses than in stabled (population S1). The population S2 falls in between, being closer to the feral populations. The samples ranking by Shanon and by Simpson indexes (TableS2) were almost identical (the maximum difference in a sample rank was 1). Shannon index is known to give more weight to species richness while Simpson index gives more emphasis on evenness (46). High correlation between these index values in the samples indicate that changes in richness are not associated with significant alterations of the evenness of the horse viromes, so all the samples contain high numbers of viral genotypes, none of which are significantly overrepresented.

**Figure 2.**
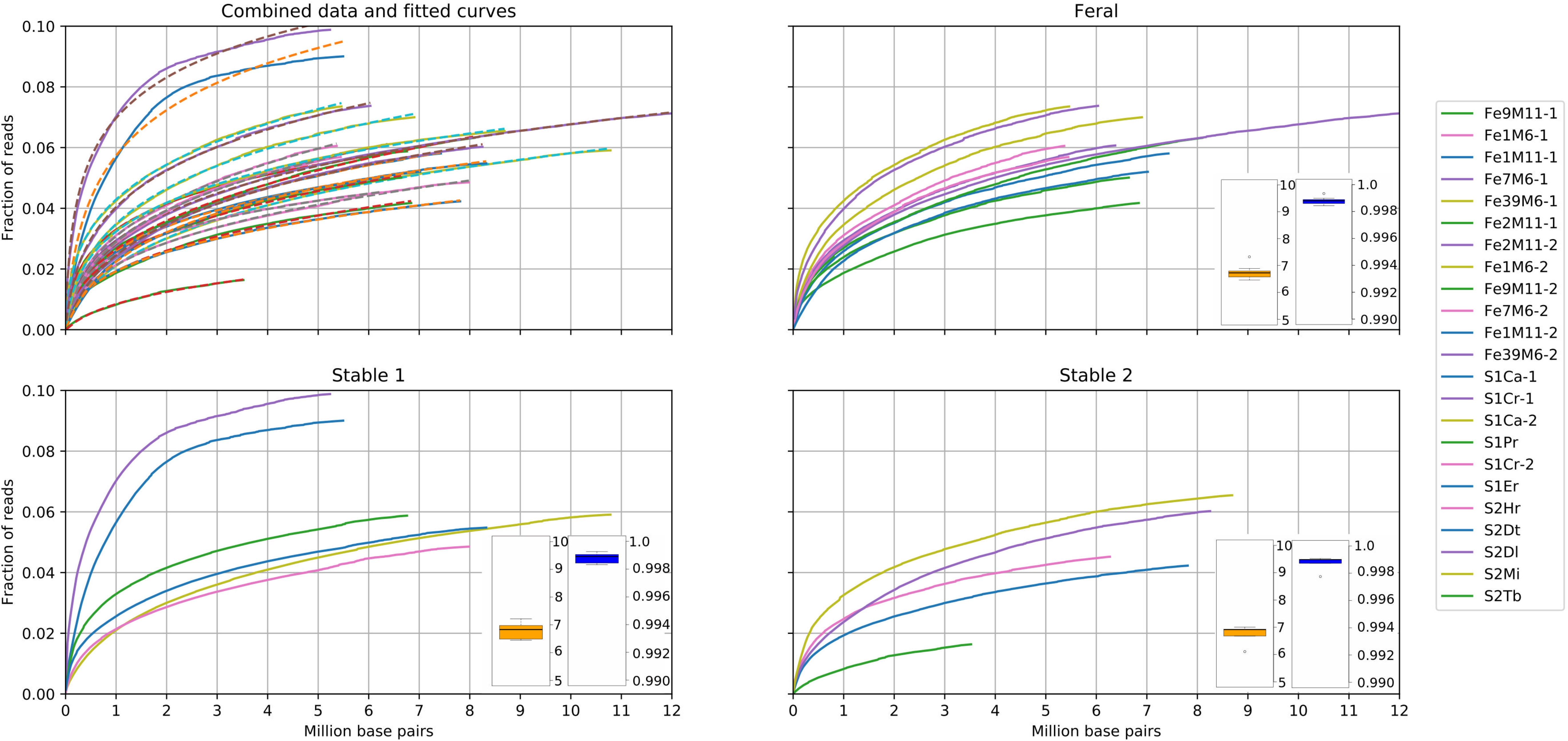
Individual viromes richness. The curves represent the cumulative fraction of the reads recruited by the contigs plotted against the cumulative length of the contigs. Contigs were sorted by CPM, taken as a proxy for their relative abundance. The dotted lines on the panel with the all the samples combined indicate the modelling function fitted to each of the curves. The distribution of the Shannon and Simpson diversity indexes in the samples from each population are plotted as box and whisker plots as inserts of respective panels.

To estimate the richness we used the approach mimicking that of Torsvik et al. (47) to estimate bacterial population richness in a soil sample. These authors estimated the complexity of the bacterial DNA extracted from soil using DNA re-association kinetics measurements. Knowing the average bacterial genome size, these authors calculated the approximate number of unique bacterial genotypes present. Instead of experimental determination of the viral DNA re-association we computed the plots of the cumulative read recruitment against the cumulative length of the contigs ranked by the abundance (TPM) in the given sample (Fig. 2).

The samples appear to differ by abundance of the most prevalent viral genotypes. This is reflected by different slopes of the initial rise of the curves, though after this initial rise the curves are almost parallel, indicating the similar law of the genotypes abundance distribution in the viromes of different animals belonging to different population. Noteworthy, in none of the samples could we detect the presence of a over-represented genotypes. If we estimate an average phage genome as 50-100 kbp, the top represented 20-40 genotypes would account for 2 Mbp of the cumulative non-redundant DNA sequence. This value corresponds to 1.3 – 8.6% of the total amount of the viral DNA (Fig. 2). After this initial rise the curves are almost parallel that indicates the similar law of the genotypes abundance distribution in the viromes of different animals belonging to different population. We were not able to reveal the law of the distribution of the genotypes fractions within the communities study and, therefore, we did not find any reliable function to extrapolate the curves outside of the available data interval. However, to estimate the lower limit of the richness we used the function f(x) = ax^b^ log(cx+1), where x is the cumulative length of the contigs, f(x) is the fraction of reads recruited by the most covered contigs with the cumulative length x, and a, b and c are the parameters fitted to minimize the square deviation from the experimental curves. As shown in the Fig 2X, within the range the modeled curves run always higher than the experimental curves. Therefore, if the distribution law remains the same, the real x values corresponding to any (x) threshold chosen will be higher than the values predicted by the function. The calculated lowest estimates of the non-redundant length of the genomes sequences of the phage particles comprising 50 % of total community for most (20 out of 24) of the curves were in the range 10^9^ – 10^11^ b.p. This translates into 10^4^ – 10^6^ distinct bacteriophage genotypes without taking into consideration of possible overlap of the sequences in many different but still related viral genomes.

### Composition of horse intestinal virome

Having established the high level of diversity of the equine gut dsDNA bacteriophage community, we asked how related are these numerous viral genotypes to known viruses. First, we attempted direct classification of the filtered reads using centrifuge. However, only ~ 0.1% of reads can be classified this way. Out of them, 97% matched viruses and 94% could be classified as dsDNA containing viruses, 81% of which were assigned to the order *Caudovirales* (tailed phages). Of these 43% were *Siphoviridae*, 41% *Myoviridae* and 13% *Podoviridae* (see Supplementary file Fig S1 for the interactive Krona plot this analysis). However, given only a minor fraction of the reads could be classified, all the subsequent analysis was performed on the assembled contigs.

Initially we used pVOGs (48) to annotate all predicted proteins on viral contigs, with a simple scoring matrix. Out of 8097 contigs, 7483 (92%) had at least one pVOG detected. Only seven contigs where pVOGs were detected, were found to have a pVOG not found in the order *Caudovirales*. Further suggesting that the vast majority of contigs originated from tailed phages. We than analyzed the relatedness at the protein level using vCONTACT2, including RefSeq genomes plus other available phage genomes at the time (May 2019). The horse virome contigs were spread across 1156 viral clusters (VCs), but only 31 were found in VCs that contain a known bacteriophage reference sequence, allowing classification at the genus level (Fig. 3, Tables S3). A further 2873 virome contigs remained singletons, once more highlighting the diversity of phages present.

**Figure 3.**
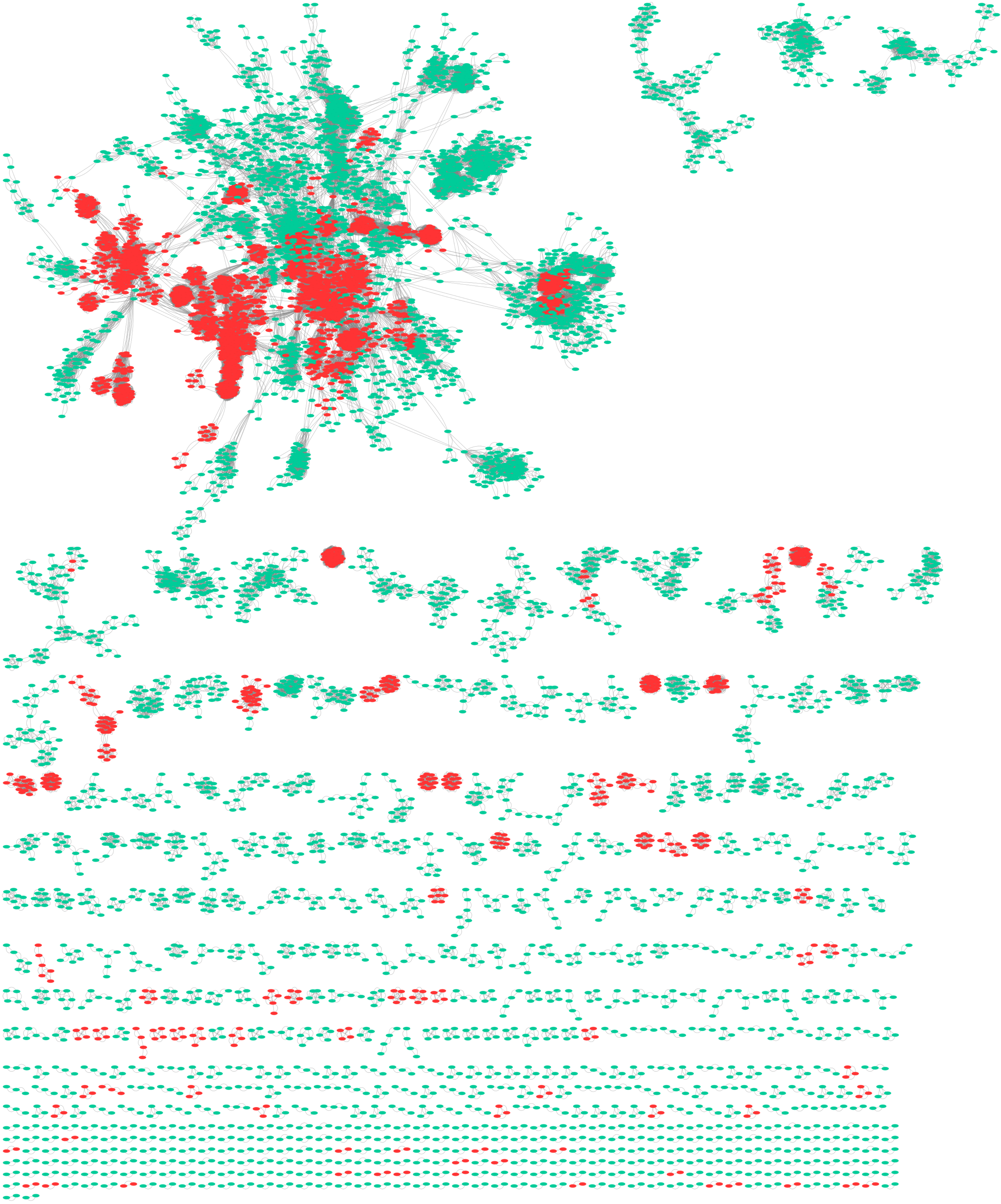
Network graph produced using vCONTACT 2. Red dots – bacteriophage reference sequences, retrieved from GenBank, green dots – the horse viromes contigs.

Due to the inability to link the majority of contigs to any known phage at the subfamily or genus level, we manually inspected the 10 largest contigs that belonged to 10 different VC clusters. Gene products were analysed with both BLASTp and HHpred (49, 50) along with gene order and orientation in the genomes. We confirmed that even for the large (35-65 kbp) contigs the links to the known viral genomes were barely detectable and lie beyond the genus or subfamily level (Table S4, Fig.S3) at which vCONTACT2 is able to operate. Only in one case (the contig 070k255_67966) were distant, but reliable relationships to the known *Gordonia* phage Gravy discovered, which was not in the vCONTACT database at the time of analysis. The results of the manual analysis further confirmed the viral diversity of the horse gut is to date very poorly sampled.

We also detected pVOGs that may be considered markers of the temperate life style (transposase, integrase, recombinase, resolvase and excisionases; see Material and methods section for detail). At least one of such pVOGs was detected in 462 (5.7%) of the 5K contigs (Table S5). Among the contigs detected in the individual samples, the highest prevalence of the temperate lifestyle markers was observed in the population S1 (7%), in the populations F and S2 the prevalence of the “temperate” contigs was about 4%. Taking into consideration that the mean length of viral contigs was 8.3 kbp, the average number of the temperate lifestyle markers per genome is three, and estimating the average length of a temperate phage genome as 40-100 kbp, we can estimate the prevalence of the phage genotypes carrying these markers as 10-25%.

### Variability of the fecal viromes between the individuals and between the populations

To compare samples at the read level, we computed the Jacard’s distances between the datasets using mash (Fig.4). The samples clustered according to the populations they were collected from. Interestingly, the feral horses cluster tighter than the stable animals within the populations S1 and S2. At the same time, we did not observe any cluster formation according to the social (harem) groups of population F. Interestingly, in population F the samples collected at the different time points from the same animal were closer to each other than to any sample from the other animals. This remarkable stability of the individual viromes was observed despite the fact that between two sampling points the animals lived out a very hot summer that was associated with severe water deprivation because the debit of the water hole was decreased about two times for several months because the hole was blocked by the sand (it was cleaned by the rangers in late September). The clustering of the samples collected from the same animal at different time points was not observed in the population S1. However, in this population the period between sequential sampling was longer (Table S1). The population S2 appears to be much closer to the population F. This may reflect the fact that the diet of these two populations is much closer to each other than to the population S1.

**Figure 4.**
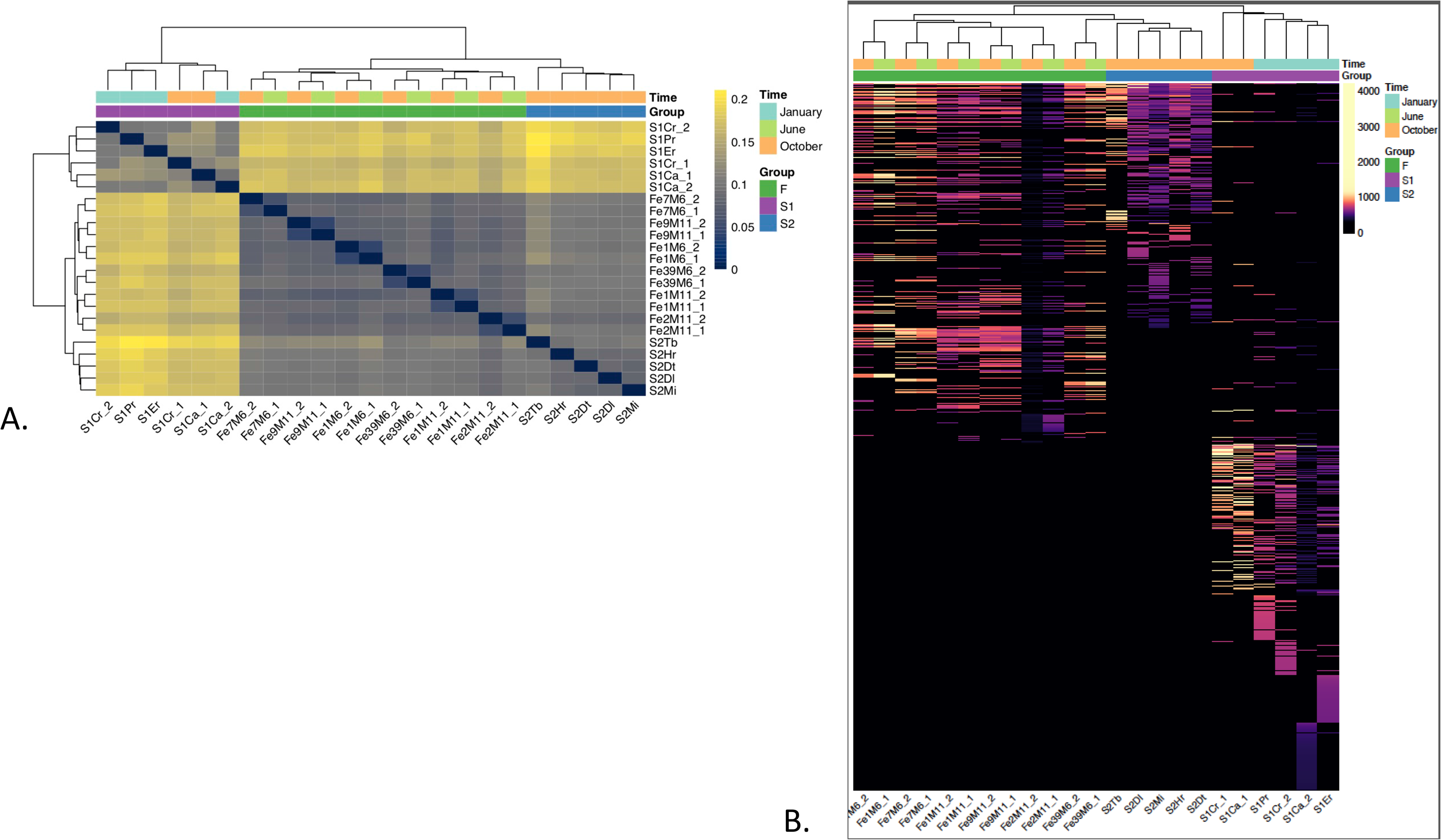
Individuality of horse fecal viromes. A – Jaccard’s distances between the samples. B. Abundance of contigs in different samples.

To compare the individual samples and populations at the contig level we mapped reads from each sample against viral contigs. Contigs were considered present in a sample if the contig had ≥ 1x coverage ≥75%, when mapping at 95% identity. Contig abundance abundance was normalized, for both contig length and depth of sequencing. Thus we used “counts per thousand per million” (CPM) value as a proxy of contig abundance (52). An average of 913 contigs (range 655 – 1105, Table S3) were detected per sample.

The heatmap of the contig abundance in the samples is shown on the Fig 4. One can see that many contigs are shared by the animals belonging to the same group but much fewer are shared between the animals. The existence of the core-virome, defined as the assortment of the viral lineages present in the majority of the sequenced samples, was recently demonstrated for human feces (24). The crAssphage –like viruses that were shown to be highly abundant in some of the samples (24, 25, 51) also belong to the human feces core-virome. In order to reveal a possible horse core-virome we identified the contigs detected in all the samples or in more than half of the animals (the samples collected at different time points from the same animal were thus joined together). These criteria were applied for each population to reveal the local core-viromes, and for all the samples to retrieve the universal core-virome.

Venns diagrams of the local core viromes relatedness are shown on Fig. 5. Only 1 contig was omnipresent in all the samples, however 192 contigs were shared by all three populations, among them 14 contigs were simultaneously present in 50% of the samples in each population (Fig. 5). Given the F population was completely isolated from S1 and S2 due to ca. 1500 km distance and protection by national reserve regimen (indirect exchange of viruses between the animal of the populations S1 and S2 is also highly unlikely though could not be completely excluded), these 192 contigs can be considered as potential candidates for a “equine core viromes”. At the same time no contig exhibited abnormal coverage comparable to the values reported for human CrAssphage (25). The largest fraction of a single in the sum of all CPMs of all contigs of a sample was 0.006 (range 0.001 – 0.006). So, no equine analog of the over-represented and wide spread crAssphage group was detected in our dataset.

**Figure 5.**
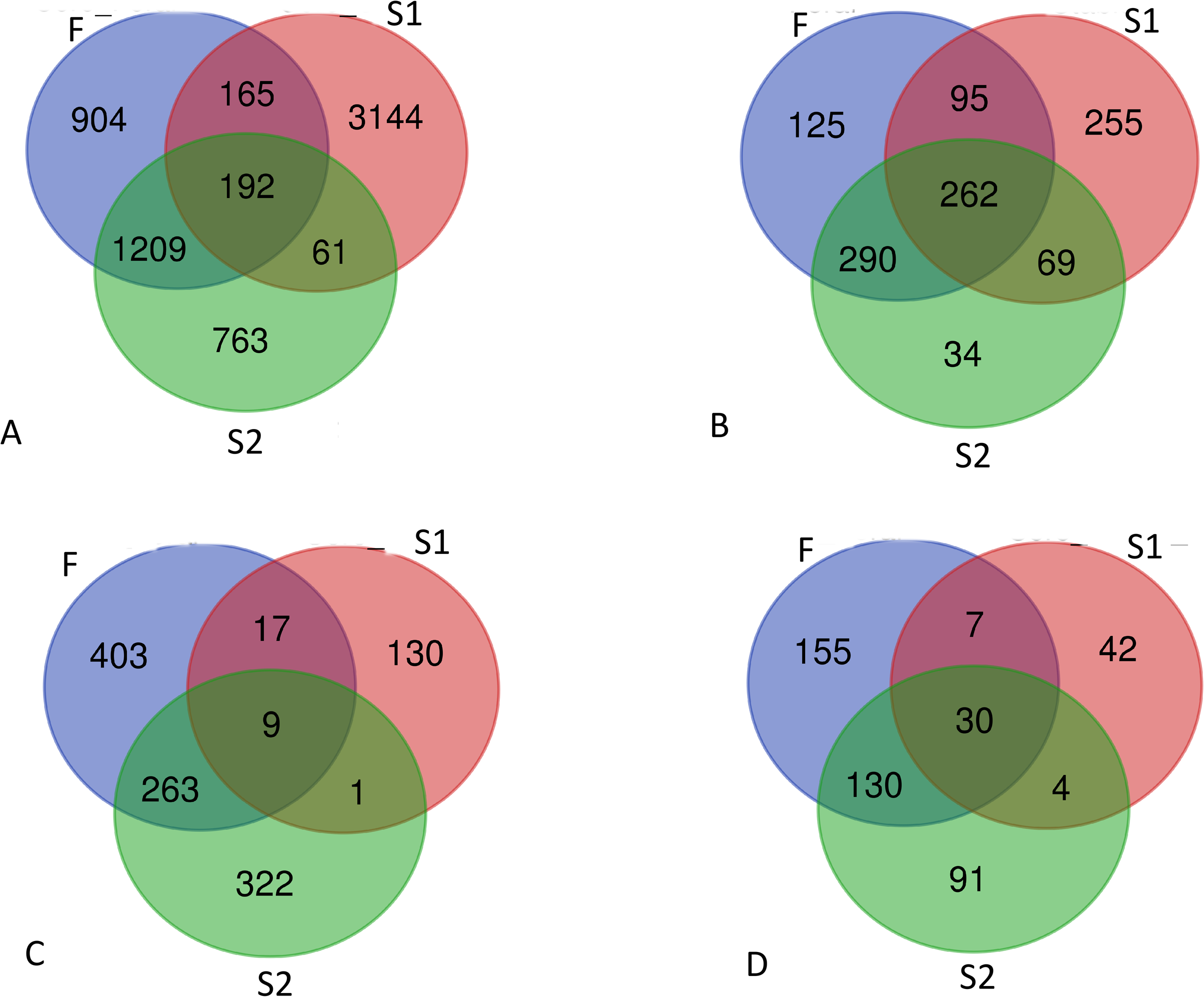
Composition of viromes in the different horse populations. A – distribution of the contigs detected in three populations. B. – distribution of the VC clusters between the populations. C. – distribution of the VCs, detected in 50% or more of the samples at least in one of three populations. D. – distribution of the contigs, detected in 50% or more of the samples at least in one of three populations

The distribution of VCs between the populations (Fig. 5B) revealed more of commonality between locations studied. Out of 1156 VCs, 262 (23%) VCs were detected in all three populations. This equates to 34-40% of VCs present in any of these populations (the VC was considered as present in a population if at least 1 contig belonging to this VC passed the detection criteria for at least 1 of the samples from this population). Interestingly, in each of the populations many VCs were present in 50% or more of the samples (Fig 5.C). The fractions of such prolific VC are larger in the populations F and S2 (89% and 90%) compared to S1 (23%). The fraction of contigs present in 50% or more of the samples in each of the populations are smaller (13%, 11% and 2% for the populations F, S2 and S1 respectively) with only 30 contigs present in 50% in all three populations simultaneously (Fig. 5D.). Thus, viromes appear to be highly individual at the level of the viral genotypes, but they appear to consist similar sets of bacteriophage genera.

### Host-phage relationships

High prevalence of the common VCs in three geographically isolated populations may reflect the presence of similar bacterial groups in the gut of horses belonging to different populations. To test this hypothesis we performed sequencing of bacterial 16S rRNA genes libraries for all samples. We also predicted putative hosts for viral contigs using WiSH (53). The prevalence of the bacterial genus level OTUs and the prevalence of the contigs predicted to belong to bacteriophages infecting these host groups is shown in Fig 6.

**Figure 6.**
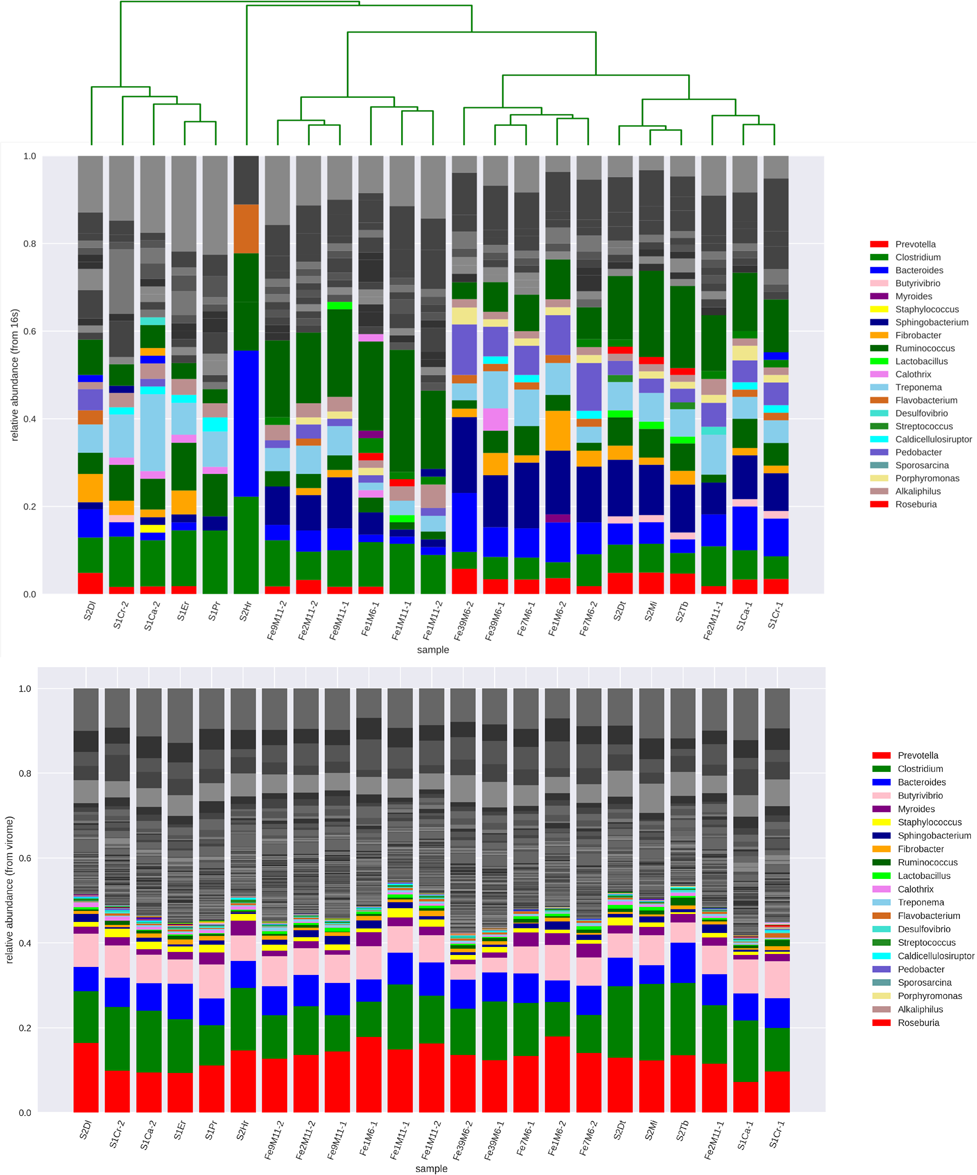
Comparison of the samples by abundance of genus-level 16S bacterial OTUs (top) and by predicted host of phage contigs. Only the OTUs or genera overlapping between the 16S sequencing results and viral host prediction are shown in color. Other OTUs are shown in gray scale.

The samples clustered according to 16S pattern still reflects the location (Fig 6.). At the same time the distribution of the prevalence of the phage genotypes predicted to infect different hosts was close to uniform. It has to be mentioned that the list of the genus-level bacterial OTUs and the list of bacterial genera predicted to be the hosts of the bacteriophage contigs only partially overlap. However, the non-overlapping OTUs have low prevalence as inferred from 16S sequencing data or from the statistics of the prevalence of the contigs allocated to the particular host.

## Discussion

Although the viral component of the intestinal microbiome is now widely believed to be an important factor in both shaping the microbial community of the gut and mediating its interactions with the macro-host (14, 18–20). The main component of the gut viromes – the community of tailed bacteriophages has only been comprehensively studied in humans. The results of our study provide a basic understanding of the composition of the dsDNA viromes of one more mammal species – *Equus caballus*. Given the very long history of domestication (54), until very recently the integral involvement of domestic horses in almost all spheres of business and military activity, and the multifaceted influence of the relationship with this species on human culture (54). It would not be an exaggeration to say that this animal species is the second most important in the development of our civilization after *Homo sapiens.*

The equine intestinal bacterial community has been extensively studied, and it is now considered to influence horse organism homeostasis and health, to the same extent as bacteria in the human gut (reviewed in (35, 55), see also (56, 57). The equine gut bacterial community is involved in pathology of specific diseases such as equine metabolic syndrome and laminitis (58). The horse behavior was also suggested to be influenced by the gut bacterial community (59).

In contrast to the bacterial community, intestinal viromes, in particular, its main component – the dsDNA containing bacteriophages – were not investigated in any detail. Our data confirms that tailed bacteriophages (order *Caudovirales*) comprise the majority of total dsDNA viromes, so for brevity we use here below the term “virome” to describe the community of dsDNA containing bacteriophages.

An individual horse virome appears to be highly diverse including more than 2000 viral genotypes (the extrapolation of the curves Fig. 2 gives estimates of more than 10^4^ - 10^6^ viral genotypes per sample). The viromes richness did not differ much between the samples analyzed. In contrast to human viromes where the contig number per sample has been shown to vary more than three orders of magnitude (17), in our samples the variation was limited to a factor of less than 2.

At the same time the evenness of the viral genotype abundance was much higher in horses as could be seen by comparison of Shannon and Simpson diversity indexes and also inferred from the reads recruitment analysis (Fig. 2). No analog of human crAssphage that is hyper-dominant in some human gut samples, was observed. The most prolific 20-40 viral genotypes, between them only accounted for 1.3-8.6% of the total number of reads in all the samples. This makes a striking contrast to the situation described for human viromes where ~2% of the contigs that are so called persistent personal viromes recruited 92.3 % of VLP reads per sample (17).

Most of the bacteriophage genotypes comprising the bulk of equine intestinal dsDNA viromes are unrelated or very distantly related to known phage genotypes. Only 31 out of 1152 identified VCs contained simultaneously horse virome contigs and known bacteriophage genomes. Moreover, the manual analysis of the 10 largest contigs, did not allow (with a single exception) to assign these sequences to any known tailed phage group. Interestingly, we estimated that only 10-25% of this immense phage diversity are represented by temperate bacteriophages. These values are in marked contrast to the human fecal viromes that are dominated by the temperate bacteriophages (27, 60). Noteworthy, the high prevalence of virulent bacteriophages in horses is in agreement with previous data on the diversity of the coliphages isolated from horse feces (41) (13). The coliphages isolated form the human feces were reported to be mainly temperate (13, 60). At the same time, high richness and high evenness of the viral community as well as the lack of the correlation between the abundance of the host 16S-based OTUs and abundance of the predicted phages to these hosts (Fig. XX) indicate that the community most probably contains multiple viruses for many (if not for all) the bacterial species present in the samples. This pattern may support the elevated diversity at the strain level, might be maintained by a kill-the-winner type mechanism (61). Our metagenomic data does not provide any direct estimates of the bacterial diversity and/or phage-host relationships at the strain level. However high intraspecies diversity of *E. coli* within horse feces associated with high diversity of co-occurring coliphages, having relatively narrow host ranges, was previous demonstrated using the culture-based approaches (41, 42). Additional culture-based evaluation of the strain-level diversity of a more prevalent species than *E. coli* combined with characterization of its co-occurring phages, may help to shed more light over the pattern of the phage-host relationships in the horse gut ecosystem. Despite the observed high virome diversity, our data suggest that a healthy horse (in the feral population the animals without visible abnormalities, wounds and marked anomalies were considered as heathy) intestinal virome includes a certain number of conserved components. The human core-virome was defined by (24) as a set of viral genotypes that are present in more than 50% of the human fecal viromes. However, the limited amount of data (22 samples from 14 animals) makes this criterion less useful for evaluation of our data. At the same time, we may benefit from the known history of strict isolation of the population of the feral horses preserved on an island in Rostovski national reserve (population F) from any contacts with other horses. The factor of geographical isolation makes direct transfer of the viral genotypes even between the ancestors of these animals over last 80-100 years unlikely. Nevertheless, we were able to find significant number of the bacteriophage genotypes present in all three populations. Approximately 3% (192 out of 6438) of contigs detected in the samples were universally present in all three locations. At the higher taxonomy level 262 out of 1130 VCs (approximately corresponding to the genus or subfamily level of relatedness) detected in the individual samples were present in all the populations (Fig 5). Moreover, despite the long history of isolation, populations S2 and F shared 552 out of 875 VCs. Thus, a significant fraction of bacteriophage OTUs of species or genus level are widely present in the horse intestinal viromes, but the fractions of these common taxa may vary significantly. Higher similarity of the fecal viromes compositions of the populations F and S2 compared to their distance to the population S1 (Figs 4 and 5) may be explained by similar diets (grass only or grass and forages compared to grass and grain diets). Given the fact that the distribution of abundances of the viral genotypes in the individual viromes is very even (Fig 2 and Table S2), such variations may obscure the commonality of the viromes composition because many shared components are present below current limits of detection. At the same time, increasing the detection sensitivity using less stringent criteria may lead to a high frequency of false detection of the viral genotypes. The deep sequencing of several viromes from different location using, for example, high output Illumina sequencing and combined with long-reads single-molecule based sequencing (e.g. Oxford nanopore) may allow characterization of the repertory of the core components of equine virome. It is logical to expect that some endemic bacteriophage genotypes may also exist in certain populations, especially in the isolated animal groups, such as the population F. However, high diversity of the viromes does not allow the identification of them at the metagenome sequence coverage levels achieved in our work.

Another remarkable feature of the horse viromes is revealed by clustering of the individual viromes compositions according to the sampling site (Fig 4). The tightest cluster was formed by the samples from the population F. The clustering of these samples did not reflect the social structure of the herd. Only the samples taken from the same animals always clustered together.

Significant fractions of contigs and VCs were found in at least 50% of the samples in each of the populations, however the percentage of the shared contigs and VCs was lower in the group of the horses stabled in the city equestrian centre (S1). These findings may be explained by significant exchange by the viral genotypes between the animals. Horse in stable 1 (S1) are held in the individual boxes, which is much more restrictive of behavior facilitating virus exchange through a fecal-oral route (see (38)). In thepopulation from stable 2 (S2) during the summer season, horses spend most of their time at the pasture, and in the feral population (F) they have no human-imposed restrictions at all. During our field work we regularly observed behavior that may allow viral exchange (for example, during the spring time large groups of horses take the mud baths in the freshwater pools, were some animals may defecate and from which they also may drink), though detailed recording of the behavior falls out the scope of this work. In such conditions, the level of all-to-all exposure may erase the signal from tighter contacts within a harem group.

The phage genotypes transfer between the horses was earlier observed by the detection of the highly related coliphage isolates that could be obtained from multiple animals held in the same stable but could not be discovered in other locations (44, 62, 63). These observations are in good agreement with the metagenomic data indicating that the transfer of the phages between the individual viromes is not limited to the minor viromes fraction(s) such as coliphages. So, the individuality and stability of the intestinal viromes are less pronounced in horses compared to humans (16, 17, 27).

Summarizing all the data, we conclude that horse intestinal viromes appear to be a more open ecological system than has been inferred from the human viromes. The bacteriophages of equine intestinal viromes represent a large pool of novel viral groups including the high level taxa such as families or subfamilies (we mean here new contemporary understanding of phage families, not the old *Siphoviridae* – *Myoviridae* – *Podoviridae* grouping within *Caudovirales*). This work provides an essential starting point from which the full genetic diversity for phages can be explored using long-read sequencing and culture based methods. Additional work is also required to analyze temporal stability of the horse viromes.

## Material and methods

### The horse populations and sampling

The sampling was performed in three geographically separated horse populations. The stable 1 population (S1) was located in a children’s equestrian club in Detski park Fili, Moscow, Russia, and represents typical stabled horses. These animals are kept in the boxes and taken outdoors for a limited time to be exercised (1-4 hours per day) and to have a rest (ca. 2 hours). These animals diet is typical for sportive horses and is comprised of foraging, supplemeted with, oats, offal and carrots. The animals have an *ad libitum* access to water. The population from stable 2 (S2) was a group of horses living in a stable located in the village of Tretyakovo, Klinski district of Moskovskaya oblast, Russia. Horses are stabled in boxes and fed by forages (carrots or apples are occasionally given to them), but they spend 8-16 hours (depending on the season) per day in the field where they are able to graze. Access to water is not limited for this population, with water provided twice a day during dry weather, where the horses can can drink *ad libitum*. In the population III four animals were sampled only once (in October 2018). Thus, the living conditions and diet of these three populations represent almost the whole spectrum of the conditions th

The herd of feral horses (F population) inhabiting Vodny Island in the salty lake Manych-Gudilo in Rostovskaya oblast, Russia. This island belongs to the core part national Reserve “Rostovski” and therefore no business activity is allowed there, and the visits to the island are restricted. The horses on the Vodny island do not receive any feeding from humans and their diet includes is only grass they forage. The access to drinking water is variable. From late autumn to spring the animals can drink from the pools or consume snow, the grass is also frequently wet due to rain and/or to dew-fall. In summer the watering is limited to water piped from the terrestrial beach once per day, and limited (ca. 3 L per min) output to an old water hole that exists in the middle of the island (64). The water from the holes is slightly saline. The access to these water sources differs significantly for different animals depending on their ranks in the social groups (harems) and on the rank of the harem stallion of their group among the harem stallions of the herd (at the moment of sampling in May and October 2017 there were 17-18 harem groups and a group of bachelor stallions). Samples of freshly voided feces were collected immediately after the natural defecation and placed in sterile plastic containers. The containers were placed on ice and transported to the laboratory. The samples from the populations S1 and S2 were processed within 24 h, the samples from the population F – within 72 h.

### Extraction of the viromes, DNA isolation and sequencing

The viromes were extracted as it described in (65) with minor modifications. Briefly, the 10 g of the fecal sample was suspended in 100 ml of the extraction buffer (0.2M NaCl, 0.1 mg/ml NaN_3_, 1 mg/ml Tween 20 (Sigma-Aldrich, USA)) and extracted on the planetary shaker at 200 rpm, room temperature for 4 h. The coarse material was then filtered out using meltblown tissue (Miracle cloth meltblown fabric), then pelleted by centrifugation at 10000 g for 15 min. The supernatant was carefully separated. The samples were filtered through the combined filter composed of Whatman GF-F glass fiber paper and a layer of diatomite (Hyflo Super-Cel). The DNAse was added to the filtrate up to 0.01 mg ml^−1^ and the filtrate was incubated for 1 h at room temp. The virus-like particles were then PEG-precipitated by adding dry NaCl to 0.6M and dry PEG 6000 (Panreac), dissolving both on orbital shaker (45 min) and allowing precipitate to form in refrigerator (+4°C, 5-6 days). Brown-greenish precipitates containing VLPs were directly extracted with CTAB using a protocol described in (66).

Ion proton shotgun sequencing metavirome DNA (approx. 1000 ng) was fragmented to a mean size of 200-300 bp using the Covaris S220 System (Covaris, Woburn, Massachusetts, USA). Then, an Ion XpressPlus Fragment Library Kit (Life Technologies) was employed to prepare a barcoded shotgunlibrary. Emulsion PCR was performed using the OneTouch system (Life Technologies). Beads were prepared using the One Touch 2 and Template Kit v2, and se-quencing was performed using Ion Proton 200 Sequen-cing Kit v2 and the P1 Ion chip. The reads were deposited to Sequence read archive (SRA) database, the accession numbers are given

### Bioinformatic analysis

Prior to assembly reads were quality controlled by trimming with Sickle (67) with default settings. Contaminating of horse DNA was removed by mapping all reads to EquCab3.0 (GCA_002863925.1) as reference genome, using bbmap with the following settings ‵minid=0.95 ‵ with any reads that mapped removed prior to assembly(68). Mapping suggested minimal contamination of horse DNA with the highest percentage of reads that mapped from any library at 0.07%. Metagenomes were assembled with MEGAHITv1.1.2, using the following parameters ‵-k-min 21 --k-max 255 --k-step 10 -t 30‵. Reads were mapped back against resultant contigs using BBmap ‵minid=0.95 covstats rpkm ‵ (68). Resultant bam and sam files were processed using Samtools v1.6 (69). To assess the level of bacterial DNA contamination all samples were processed with SortMeRNA v2.1 to check for contaminating rRNA reads ‵ sortmerna --ref /usr/local/bioinf/sortmerna-2.1/rRNA_databases/silva-bac-16s-id90.fasta,/usr/local/bioinf/sortmerna-2.1/index/silva_b90:/usr/local/bioinf/s$ ‵ (70) and also using viromeQC. For read based assessment of viral diversity centrifuge was used with default settings and database of known phage genomes.

For contig based assessment of viral diversity, contigs were first filtered with DeepVirFinder to remove any contigs that are of likely bacterial origin, only contigs with a p value <0.05 and were greater than 5 kb in length were considered for further analysis (71). Contigs were considered to be present within a sample if the average coverage of mapped reads was ≥ 1X over ≥70 % of the sample as recommended in other studies (4). The relative abundance of contigs within each sample was determined by counting the number of reads mapped to each contig, divided by the length of the contig (Kbp) to give RPK. The sum of all RPK values per sample was divided 1 000 000, with each RPK divided by a 1 000 000. Processing of data was carried out in R, using the PhyloSeq (72) library to calculate diversity statistics. To identify circular contigs lastal ‵-s 1 -x 300 -f 0 –T‵ was used to identify the ends of contigs that overlapped (5).

Contigs were annotated automatically using Prokka v1.12 using the following settings ‵ --meta ‵., using a custom database phage proteins (73). This database was constructed by extracting all the proteins from publically available phage genomes within the European Nucleotide Archive (5) and then further annotated using the scripts associated with prokka to do so []. Further annotation was provided by the use of hmmprofiles using hmmscan with the prokaryotic Viral Orthologous Groups (pVOG) collection of hmm profiles using a cutoff value of 1E^−5^ (7, 8). To identify putative temperate phages a method akin to Sh & Hill was used. We utilised the a set of PFAM hmms (PF07508, PF00589, PF01609, PF03184, PF02914, PF01797, PF04986, PF00665, PF07825, PF00239, PF13009, PF16795, PF01526, PF03400, PF01610, PF03050, PF04693, PF07592, PF12762, PF13359, PF13586, PF13610, PF13612, PF13701, PF13737, PF13751, PF13808, PF13843, and PF13358) that are specific to bacteriophage transposase, integrase, recombinase, resolvase and excisionases.

Putative hosts were predicted using WIsH (53). A database of 9075 complete bacterial genomes was downloaded from Genbank (Jan 2018) and models were constructed for each bacterial genome within WIsH. Null parameters were calculated for each bacterial model using 7000 bacteriophage genomes. Hosts were predicted for each phage contig in the virome, with only predictions that had a pvalue of < 0.05 considered for further analysis.

### Closest relatives

A custom database of all known phages genomes was produced by extraction of ~ 10,000 complete phage genomes from genbank as previously described. A MASH database was produced for using sketch –s 10000. Each contig was queried against this database using mash dist function, with the top hit that had a distance of < 0.05 assigned as it closest known relative. To cluster contigs at the genus level, vContact2 was used with the following settings “--rel-mode ‘Diamond’ --db ‘ProkaryoticViralRefSeq85-Merged’ --pcs-mode MCL --vcs-mode ClusterONE”. The network graphs was visualized in Cytoscape and using Python package graphviz_layout.

Diversity indices were produced by use the R Phyloseq package (72)

## Acknowledgements

We acknowledge T. Redgwell for help with proofreading and V.N. Filatova for help with the samples collection. The work was partially supported by RFBR grant #18-29-13029 (the field work and the pre-sequencing samples processing were performed before the grant acquisition). AM was funded by MRC grants MR/L015080/1 & MR/T030062/1. Bioinformatic analysis was in part carried out on infrastructure provided by MRC-CLIMB.

---

**Figure S1** Krona plots of reads classification using Centrifuge algorithm.

**Figure S2.**
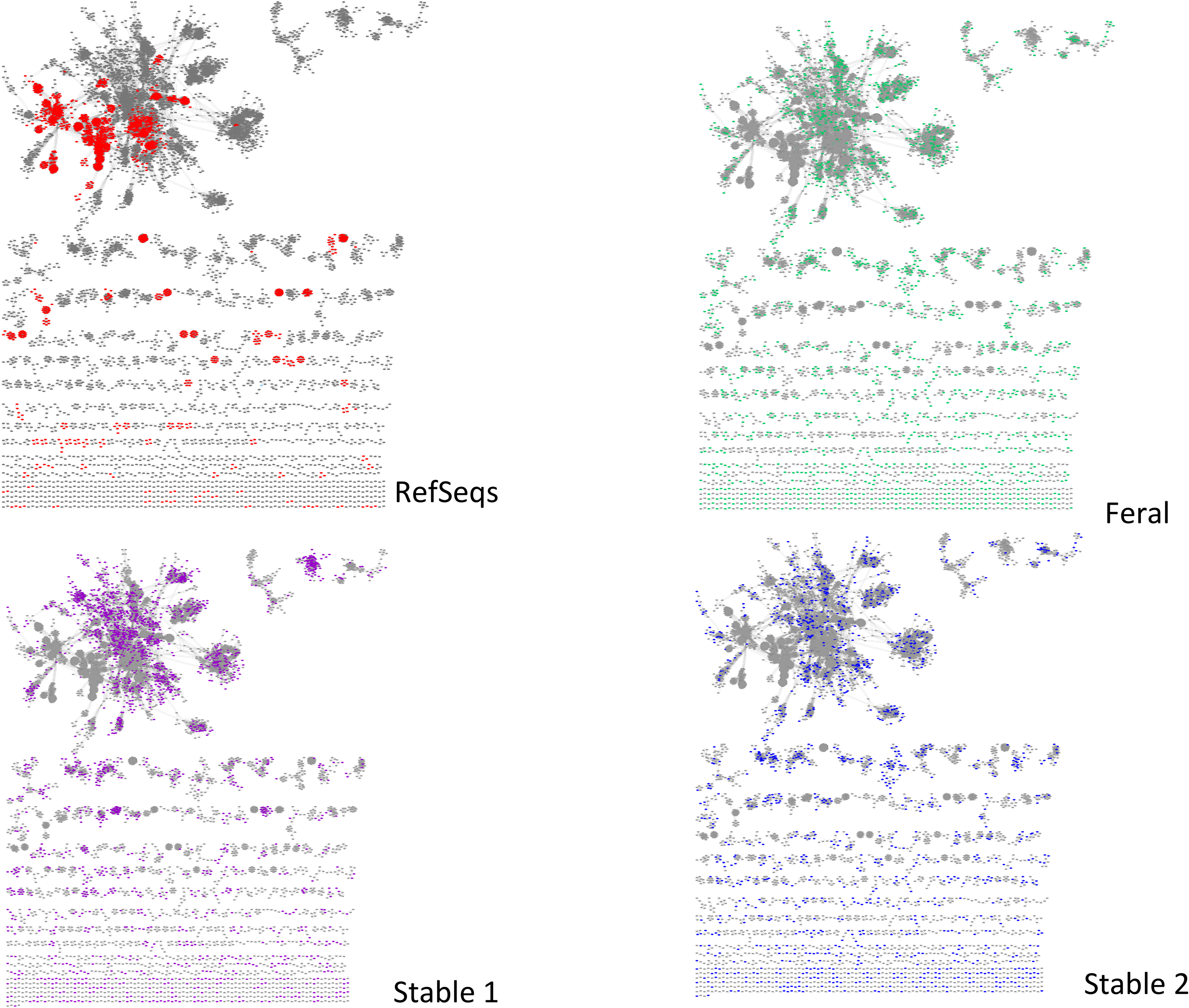
Graphs of contigs clustered using vCONTACT 2 algorithm by populations.

**Figure S3.**
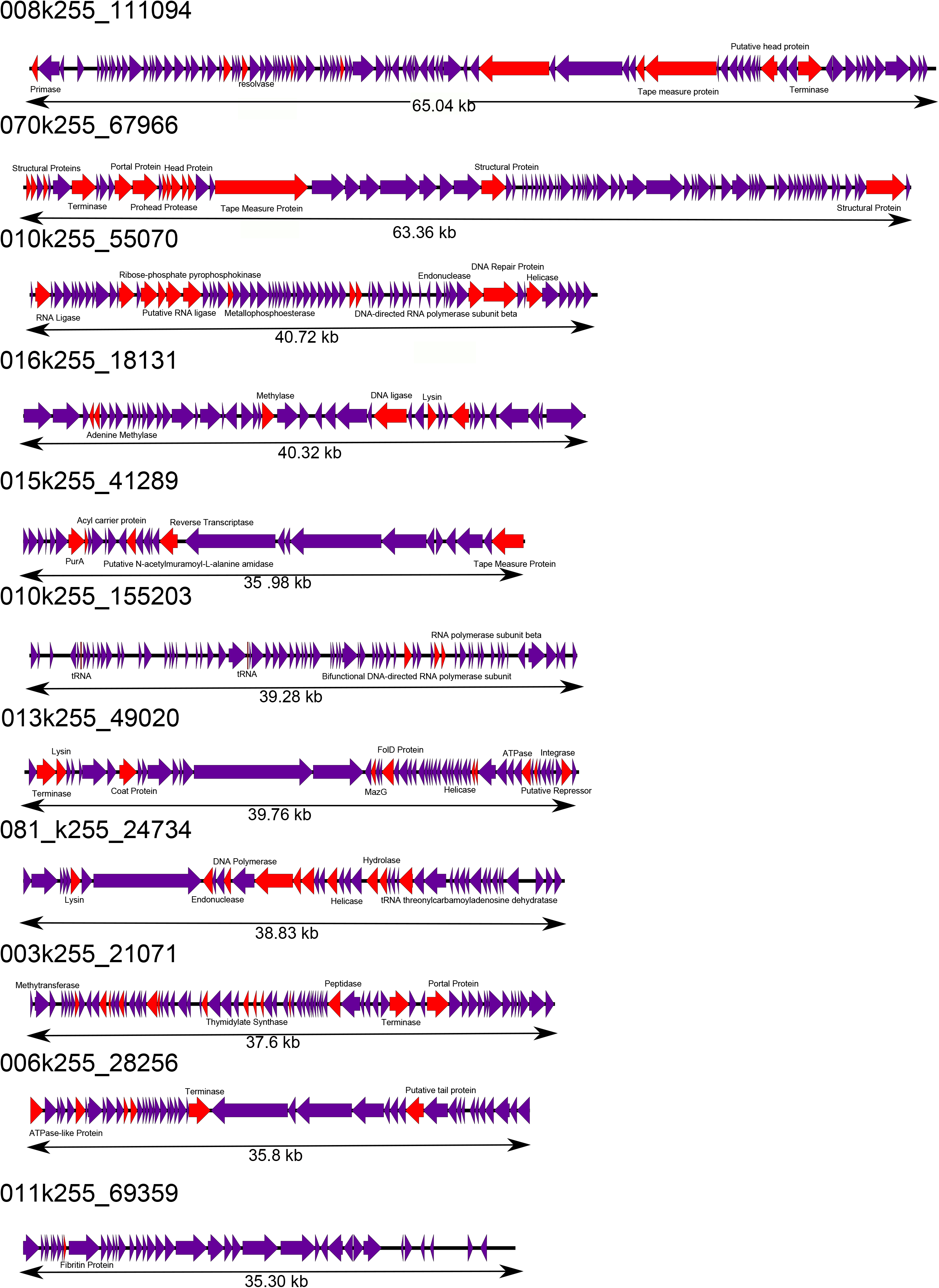
Genome maps of 11 largest contigs subjected to manual analysis. Genes coloured had a putative function assigned, those in purple have no known function.

## Notes

### Competing Interest Statement

The authors have declared no competing interest.

