## Supplementary figures and images for "The ecogenomics of dsDNA bacteriophages in feces of stabled and feral horses"

### Figure S3

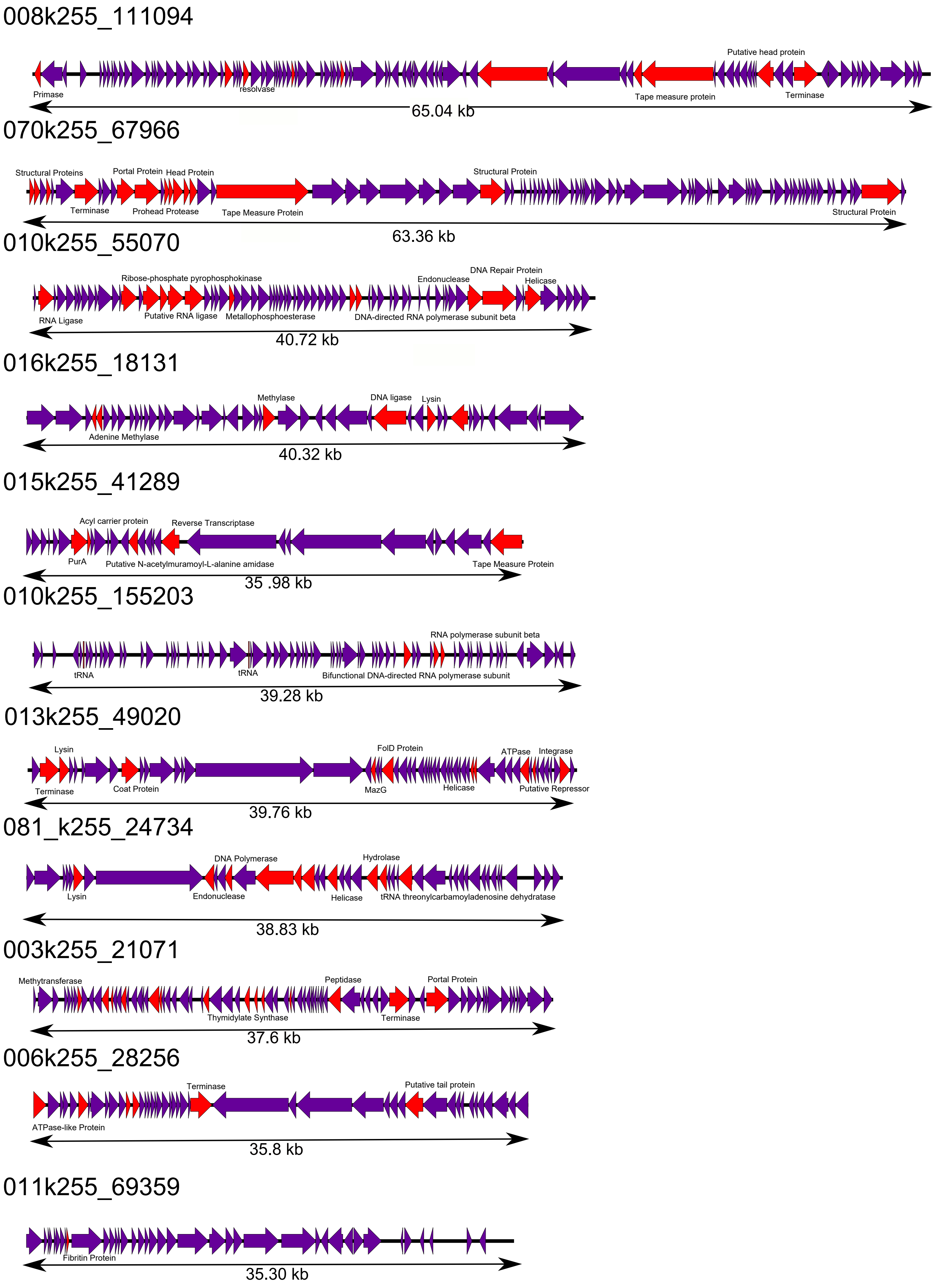
