## Supplementary material for "The ecogenomics of dsDNA bacteriophages in feces of stabled and feral horses": Figure S1: FigS1.html

Javascript must be enabled to view this page.

magnitude
magnitudeUnassigned
count
unassigned
taxon
rank
score

Fe1M11-1
Fe1M11-2
Fe1M6-1
Fe1M6-2
Fe2M11-1
Fe2M11-2
Fe39M6-1
Fe39M6-2
Fe7M6-1
Fe7M6-2
Fe9M11-1
Fe9M11-2
S1Ca-1
S1Ca-2
S1Cr-1
S1Cr-2
S1Er
S1Pr
S2Dl
S2Dt
S2Hr
S2Mi
S2Tb


68.5264.6567.9667.6970.9167.4666.467.0468.8264.7364.6465.9970.5471.3972.6873.8977.8976.3170.1870.9269.1469.9468.26
504.719999999989475.24999999999498.049999999986497.899999999987521.28999999999499.529999999979486.33999999999492.66999999999504.319999999988476.559999999981478.569999999988486.169999999989519.25519.989999999986529.589999999999527.749999999985562.009999999982547.039999999987511.629999999983515.989999999988507.239999999988512.509999999985496.769999999987

432.819999999989407.969999999991427.699999999987427.499999999988446.919999999992429.099999999981415.399999999992421.939999999991432.09999999999408.809999999984411.159999999989417.74999999999446.920000000003445.889999999988453.949999999995451.839999999988482.609999999983467.529999999991438.989999999984442.039999999989434.649999999988438.429999999986425.369999999988
4.379653.524031.956773.540893.027526.744092.885842.496732.237352.131483.186063.321653.4518916.34182.779633.207347.973616.99468.563326.772588.532347.44344.7277
68.3864.4867.8367.3570.7967.2966.2566.8668.5864.4364.565.8470.4571.2872.5273.8577.8576.1969.9570.7968.8569.7268.16
10239
superkingdom

0.530.260.150.450.230.350.10.10.190.20.080.160.050.050.090.090.650.25
0.3773580.3846150.20.5111110.1739130.7142860.20.20.21052600.40.250.250.400.20.22222200.2222221.1538500.68
1714266
0.110.060.030.090.050.080.020.020.050.040.020.040.010.010.020.020.160.05
no rank

0.180.120.090.140.060.120.030.030.41
0.5555560.6666670001.33333000.285714000.3333330.3333330000.33333300.3333331.73171
family
0.060.040.030.050.020.040.010.010.14
1714267

0.83333310002000.444444000.50.50000.500.52.62963
0.120.080.060.090.040.080.020.020.27
0.060.040.030.050.020.040.010.010.14
1714268
genus

0.040.020.010.09
02000000100000000017
species
1154689

0.040.020.010.02
20000000100000001002
species
1282967

species
326574
10000400000110000002
0.020.030.020.040.02

family
1911603
0.020.01
0000.25000000000000.25
0.080.04

0.06
0000.333333
genus
0.02
1911605

species
0.02
1987108
0.04
0000.5

0001
0.02
1156722
no rank

0.01
1911604
genus
0000000000000000.333333
0.03

0.01
1987101
species
0.02
0000000000000000.5

no rank
634168
0000000000000001
0.01

0.4166670.250.250.750.2222220.7222220.250.25000.5000.5000.2500.250.500.85
0.240.080.120.280.180.180.080.080.160.040.040.040.080.2
family
0.060.020.030.070.050.040.020.020.040.010.010.010.020.05
1914302

1932924
0.060.020.030.070.050.040.020.020.040.010.010.010.020.05
genus
0.5555560.3333330.33333310.3076920.9285710.3333330.333333000.666667000.666667000.33333300.3333330.66666701.13333
0.180.060.090.210.130.140.060.060.120.030.030.030.060.15

species
1932927
0.040.020.030.070.020.030.020.040.010.010.010.04
0.080.040.060.140.040.060.040.080.020.020.020.08
10.50.51.50.51.50.50001001000.500.5002

0.040.020.030.070.020.030.020.040.010.010.010.04
2113131000200200101004
no rank
543939

0.040.040.040.040.040.02
0.50000.5100.500000000000100.5
0.020.020.020.020.020.01
1932926
species

0.020.020.020.020.020.01
1000120100000000000201
no rank
1411887

0.130.150.220.210.10.210.050.20.160.060.150.120.160.330.070.240.220.060.130.160.090.090.24
12429
no rank
1.111111.862071.954554.50.7368424.1555613.102560.81250.52.333332.166671.2510.49250.53.673473.10871.751.562.43752.722220.5882351.47826
0.270.290.440.420.190.450.10.390.320.120.30.240.320.670.140.490.460.120.250.320.180.170.46

0000000000000002
0.02
species
1843765

0.010.01
0000000000000000101
1955196
species

1958785
species
00000000000000000000001
0.01

1685779
species
0.010.01
00000100000000000001

00000000201
0.050.02
species
1849539

species
1685735
0000000000000004000001
0.040.01

0.03
00000000000000000000002
species
1654356

1685773
species
0.040.01
00000000000000000003001

species
1911439
0000000000001
0.04

species
1542745
00000000100002
0.020.01

1685740
species
00000000000001001030002
0.010.010.030.03

0000000000000001
0.01
species
1587517

01000321000132915213001
0.020.030.050.020.020.120.210.070.050.020.010.030.01
species
1500510

00100100000000002000001
0.030.010.020.01
1955175
species

0.040.010.010.01
20000000000000000001011
species
1920989

00000000000000011
0.010.01
species
1685739

0.01
00000000000000000000001
159252
species

species
1592828
0.020.01
0000000010000000000001

0.01
000001
1685749
species

0.02
000002
1868621
species

0.01
00000000000000000001
species
1843767

species
1685745
00000000000000000000001
0.01

0.010.030.030.02
0000010001000000400002
species
1955197

000000.500000001
0.020.02
1603065
0.010.01
no rank

000001
0.01
species
1843751

00000000000002
0.01
species
1595998

00000000001000013
0.020.010.03
species
1843770

0.020.02
10001
1888308
species

species
1685786
0.02
000000001

1911438
species
00001000000000001
0.020.01

species
1843766
0.020.010.010.01
010001000000010000001

0.020.030.01
10100000000000000000001
1843763
species

0.01
0000000000000000000001
species
1685785

0000000000000000000001
0.01
1844927
species

species
1678078
0.060.10.160.210.050.120.170.050.030.110.10.080.110.10.050.050.090.080.08
355921407216501101211447606

0.01
00000000000000000000001
1685762
species

no rank
186616
0.02
0.04
00000000000.5

1379788
species
00000000001
0.02

0.450.150.10.130.250.10.070.490.040.230.050.760.240.16
0.17777800.20.200.384615000.16000.200.2857140.142857000.250.3478260.21.131580.7083330.4375
no rank
0.070.030.020.030.050.020.010.070.010.050.010.180.050.04
12877

family
0.040.030.020.030.020.020.010.020.010.170.050.04
1993640
0.160.120.080.10.080.080.030.060.040.530.190.12
0.2500.250.2500.5000.25000.25000000.3333330.6666670.251.603770.8947370.583333

0.33333300.3333330.33333300.714286000.333333000.333333000000.510.3333332.361111.214290.875
0.120.090.060.070.060.060.020.040.030.360.140.08
190729
0.040.030.020.030.020.020.010.020.010.170.050.04
no rank

1428189
species
000000000000000001
0.01

000000000000000000004
0.05
species
219596

00000200000000000020712
0.020.020.090.010.03
species
435488

221036
species
00000000000000000000001
0.01

0.080.060.040.020.040.040.020.050.08
0.500.50.500.5000.5000.500000000.50.42
no rank
0.040.030.020.010.020.020.010.030.04
361689

0.01
000001
species
1175505

706122
species
0.02
000000000001

000000000000000000001
0.01
1246849
species

1234171
species
000000001
0.02

species
713092
100000000000000000001
0.020.01

1506157
species
001
0.03

10010000000000000001
0.020.020.01
species
1507403

species
447603
0.04
0000000000000000000004

0.220.120.060.420.120.05
0.18181800000000.16666700000.3333330.1666670000.33333300.2
0.040.020.010.070.020.01
198601
no rank

361688
0.040.020.010.070.020.01
no rank
0.180.10.050.350.10.04
0.22222200000000.200000.40.20000.400.25

no rank
0.040.020.010.070.020.01
1458186
0.28571400000000.2500000.50.250000.500.333333
0.140.080.040.280.080.03

0.040.020.010.070.020.01
283494
no rank
0.10.060.030.210.060.02
0.400000000.33333300000.6666670.3333330000.66666700.5

0.040.040.020.140.04
0.500000000.5000010.50001
0.020.020.010.070.02
1231296
no rank

0000000000000000002
0.02
species
661505

0.07
000000000000001
945250
species

000000001
0.02
species
1293961

0.02
1
1001263
species

1933092
species
0.01
00000000000002

100000000000000000001
0.020.01
915293
species

no rank
29258
0.290.290.120.280.360.380.170.450.250.470.190.290.040.250.070.160.240.540.330.270.180.30.56
0.3262410.3203120.3478260.6065570.2105260.4766840.8101270.3894740.2393160.4615380.2346940.2720.21.354550.20.6216220.8416671.271930.520270.4601770.2528740.43750.516667
1.411.280.461.221.711.930.791.91.172.210.981.250.21.10.350.741.22.281.481.130.871.442.4

0.250.120.180.080.080.060.220.080.130.060.20.080.190.060.150.040.070.12
0.480.3333330.6666670.250.250.16666700.272727000.250.15384600.33333301.250.50.6842110.6666671.333330.250.2857140.583333
39724
0.070.040.060.020.020.020.070.020.040.010.050.020.050.020.050.010.020.04
family

0000000000000000000000.333333
0.03
227307
0.01
genus

988775
0.01
no rank
0.02
0000000000000000000000.5

species
1548711
0.01
0000000000000000000001

no rank
642248
0.020.020.010.010.050.010.01
0.060.040.020.020.10.020.02
00000.3333330000000.500.50000.50200.50.5

1618238
species
0.01
00000000000001

species
1618245
0.01
0000000000000000000001

0.01
00000000000000000000001
species
1618257

1611039
species
000000000000000001
0.01

0.02
1454552
species
0.04
00000.5

1105383
no rank
0.02
00001

000000000001
0.02
1618243
species

0.05
00000000000000000004
species
1618255

no rank
742826
0.02
0.04
0.5

species
1379718
0.02
1

00.5000000.5000000.33333301.666670.6666671
0.040.010.040.050.030.150.060.12
genus
0.020.010.020.020.010.050.020.04
39725

species
85708
01
0.02

00000001
0.02
species
1194757

00000000000000.502.511.5
0.020.10.040.08
species
2003309
0.010.050.020.04

1732201
no rank
000000000000010523
0.010.050.020.04

00.01
000000000000
133704
species

00.01
345249
no rank
000000000000
00.02

000000000000
00.01
species
720569

0.060.020.060.020.010.050.020.020.010.03
742914
no rank
0.7142860.510.33333300.33333300.363636000.3333330000000100.33333301
0.140.040.120.060.030.110.060.040.030.06

21200001000000000020002
0.040.020.060.020.020.03
1234881
species

0.5000.500.500.5000.50000000000.5
0.040.040.020.040.040.02
no rank
1987730
0.020.020.010.020.020.01

000001
0.01
1506568
species

942034
species
00010001
0.020.02

100000000010000000001
0.020.020.01
1908802
species

0.60.2000.2469140.476190.250.1739130.1666670.20.27777800.25000.2307690.3181820.1666670.4583330.200.2285710.333333
0.30.20.810.630.080.230.120.30.360.160.130.220.180.240.10.350.21
0.060.040.170.120.020.050.020.060.070.040.030.040.040.060.020.080.04
10780
family

40120
0.020.10.080.050.020.040.020.020.020.030.010.030.01
subfamily
0.080.370.320.180.10.160.080.080.090.120.040.140.05
00.25000.3783780.5937500.2222220.200.500000.250.250.2222220.750.2500.1428570.2

1513201
0.020.040.010.030.01
genus
00.3333330001.08333000000000000.3333331000.333333
0.060.120.030.090.03

0.020.040.010.030.01
1513202
species
00.50001.625000000000000.51.5000.5
0.040.080.020.060.02

1513204
no rank
0.010.030.01
0000010000000000003001

010004000000000001
0.020.030.01
797418
no rank

0.150.090.120.03
00000.6666670.55555600000.666667000000000.333333
40122
0.050.030.040.01
genus

0.02
000000.5
1513213
0.01
species

0.01
000001
no rank
1221208

0.10.040.080.02
00001100001000000000.5
species
1513212
0.050.020.040.01

00002200002000000001
0.050.020.040.01
no rank
292208

0.060.030.130.060.030.040.03
00000.3333330.33333300.30769200000000.3333330.3333330.250000.333333
genus
10806
0.020.010.050.020.010.010.01

000000.5
0.02
species
1513187
0.01

159153
no rank
000001
0.01

species
0.020.020.01
1979752
0.040.040.02
00000.5000.500000000000000.5

no rank
210016
0000100100000000000001
0.020.020.01

00000000000000000.5
0.02
1979753
0.01
species

00000000000000001
0.01
1414671
no rank

0000000000000000.500.333333
0.020.03
0.010.01
1513188
species

0.010.01
12524
no rank
0.010.02
000000000000000100.5

no rank
185636
0.01
000000000000000001

no rank
0.02
1169623
00000000.5
0.04

00000001
0.02
species
72003

0.01
1979750
species
0000000000000000.5
0.02

no rank
1642018
0000000000000001
0.01

00000.3333330000.2500000000.333333000000.25
0.060.080.030.050.04
136466
0.020.020.010.010.01
genus

351053
0.02
no rank
0.04
00000.5

0.02
00001
945962
species

0.060.040.03
000000000.33333300000000000000.333333
species
1513193
0.020.010.01

35338
0.020.010.01
no rank
0.040.030.02
000000000.500000000000000.5

no rank
487311
0.020.010.01
00000000100000000000001

0000000000000000000000
0.01
1513196
no rank

species
1513198
0.01
00000000000000000.5
0.02

no rank
1513199
00000000000000001
0.01

subfamily
0.060.020.070.040.020.060.030.040.030.010.010.010.030.03
40119
0.750.25000.2222220.5789470.333333000.250.15384600.3333330000.50.20.250.2500.4545450.5
0.240.080.270.190.060.240.130.120.10.050.040.040.110.12

1507401
0.020.010.060.01
genus
0.080.030.180.03
00000.250.3333330000.3333330000000.333333

00000.3333330.50000.50000000.5
0.060.020.120.02
0.020.010.060.01
1507402
no rank

species
1911555
0.03
0000000001

00000.5
0.04
0.02
1133653
species

1131622
no rank
00001
0.02

0.010.030.01
00000100010000001
1329649
species

00000000000.25
0.08
genus
0.02
10803

species
0.02
1511894
0.06
00000000000.333333

00000000000.5
0.04
341671
0.02
no rank

0.02
00000000001
no rank
281492

0000000.5000000.5000100000.5
0.040.020.080.040.02
no rank
207678
0.020.010.040.020.01

0000000000000000000001
0.010.01
1391667
species

1341019
species
0.02
00000000000000002

0.02
0000001
1208310
species

species
1131485
0.04
0000000000001

0.03
00000000000000000000.333333
genus
0.01
40121

0.01
344889
no rank
0.02
00000000000000000000.5

00000000000000000001
0.01
72197
species

0.020.010.01
1506574
genus
0.060.030.04
00000.3333330.333333000000000000.25

no rank
0.020.01
1506575
00000.50.5
0.040.02

000011
0.020.01
species
740933

species
1511906
0.01
000000000000000000.333333
0.03

000000000000000000.5
0.02
10786
0.01
no rank

0.01
000000000000000001
no rank
10788

0.180.060.060.090.030.060.09
10.333333000.33333310000000000000.333333000.6666670.666667
310911
0.060.020.020.030.010.020.03
genus

no rank
0.060.020.020.030.010.020.03
1233218
0.120.040.040.060.020.040.06
1.50.5000.51.50000000000000.50011

species
1513315
31001300000000000010022
0.060.020.020.030.010.020.03

0.020.020.02
0000000000000000.5000.5000.5
0.010.010.01
535600
no rank

species
1926498
0000000000000000000001
0.01

0000000000000000001
0.01
species
1519097

0.01
0000000000000001
1581151
species

0.2352940.840.251.2075500.7916670.3333330.3333330.2758620.7714290.3333330.43478300.3333330.2500.2857140.250.4166670.5555560.28571410.214286
0.170.250.160.530.240.060.060.290.350.090.230.120.280.010.070.080.240.180.070.120.14
0.050.080.050.160.070.020.020.080.110.030.070.030.070.010.020.020.080.060.030.040.05
10860
family

no rank
0.02
1074043
0000000000000000001
0.04

0.02
0000000000000000002
species
1449897

000000.3333330000000000000
0.030.060.05
genus
0.010.020.02
1977139

species
1977140
0.010.01
0.020.02
000000.5000000

223524
no rank
0.010.01
000001000000

0000000000000000000
0.020.03
0.010.02
70203
species

no rank
181604
0.010.01
0000000000000000000

0.030
000000000000000000.33333300000
1977141
0.010
genus

000000000000000000.500000
0.020
species
0.010
1977299

0.010
00000000000000000100000
no rank
1505528

0.020.040.040.0200.020.020.06
0000000.50.5000000.5000.50000.500.333333
genus
1977144
0.010.020.020.0100.010.010.03

0.01
1977408
species
00000000000000000000000.5
0.02

0.01
00000000000000000000001
185959
no rank

00000011000001001000101
0.010.020.020.0100.010.010.01
455649
species

456491
0.020.070.040.120.050.040.090.030.050.030.030.010.04
no rank
0.51.50.52.501.7000.51.50.510000000.85714310.51.5
0.040.140.080.240.10.080.180.060.10.070.060.020.08

642111
species
0.010.01
0000010000000000000

10864
species
1315040013120000002213
0.020.070.040.120.040.040.090.030.050.030.030.010.04

0000.30769200.5000.30769200000.50.33333300.3333330.33333300.666667000
0.130.020.130.040.2100.030.030.060
no rank
1977138
0.050.010.050.010.0700.010.010.020

127509
species
00000100000001
0.010.01

0.04
0000.5
0.02
1977411
species

0.02
0001
no rank
10868

0.010.010.010.02
2011074
species
00000000000000.5000.50.501
0.020.020.020.04

93224
no rank
0.010.010.010.02
00000000000001001102

0.04
000000000.5
species
0.02
2011073

000000001
0.02
1105327
no rank

00000000000000000000000
0
species
2011075
0

no rank
384360
0
00000000000000000000000

0.140
000000000000000.50
species
2011076
0.070

127507
no rank
0000000000000010
0.070

2011082
0.02
species
0.04
000000000.5

no rank
10872
0.02
000000001

2011079
0.02
species
0.04
0000.5

0001
0.02
no rank
1343494

00000000000000.333333000000000.333333
0.030.03
genus
10875
0.010.01

1977403
0.010.01
species
00000000000000.5000000000.5
0.020.02

no rank
10876
0.010.01
00000000000001000000001

0.020.0600
00000000000000000000000
genus
10861
0.010.0200

00000000000000000000000
0.010.0400
0.010.0200
356623
no rank

0000000000000000
0
species
127508

1150992
species
00
00000000000000000000000

species
1352227
00.02
0000000000

0.333333000000000000000000
0.060.030.030
genus
1977142
0.020.010.010

0.5000000000000000000
0.040.020.020
species
1977304
0.020.010.010

1000000000000000000
0.020.010.010
no rank
10867

00.250000.81.2510000.2500.77777800001.500.401.07692
0.160.20.480.40.080.090.280.20.39
family
0.040.050.120.10.020.030.070.050.11
687329

no rank
363628
0.03
0.09
000000000000000000000.666667

0.06
000000000000000000001
species
0.03
393049

289366
no rank
000000000000000000002
0.03

0.090.360.30.040.210.030.02
000001.333331.666671.33333000001.50000200.33333300.5
0.030.120.10.020.070.010.01
687332
genus

0.030.120.10.070.01
1535289
no rank
0000022.520000000000300.5
0.060.240.20.140.02

000000000000000000001
0.01
species
1859149

0.030.120.10.07
0000045400000000006
1535290
species

species
687372
0.02
00000000000003

0.01
00000000000000000000001
species
687375

00.3333330000.666667000000.33333300.50000000.33333301.57692
0.120.060.060.020.030.26
0.040.020.020.010.010.09
687331
genus

1619215
0.040.020.020.010.07
no rank
0.080.040.040.020.14
00.50001000000.5000000000.502.5

0.020.020.020.010.07
01000200000100000000105
1629657
species

1629656
species
01
0.02

species
687351
0.01
00000000000001

687346
species
0.03
00000000000000000000002

0.080.140.080.240.170.480.060.260.270.060.330.450.070.040.180.040.1
0.250.2857140000.500.250.23529410.3333330.4615380400.6666672.3030340.2857140.250.3333330.250.2
0.020.040.020.070.050.120.020.080.090.020.090.150.020.010.030.010.03
10841
family

subfamily
1910950
0.03
000000000000000000000.4
0.15

genus
0.03
1910952
0.12
000000000000000000000.5

1986034
0.03
species
0.09
000000000000000000000.666667

no rank
0.03
374842
0.06
000000000000000000001

no rank
374844
000000000000000000002
0.03

00.3333330000.66666700.3333330.3333331.3333300.33333300000.500.3333330.33333300.3333330.285714
0.060.060.060.060.360.060.080.030.030.030.07
subfamily
686565
0.020.020.020.020.120.020.020.010.010.010.03

00000000000000000.666667
0.06
genus
40274
0.02

00000000000000001
0.04
species
0.02
2003328

0.02
00000000000000002
105154
no rank

0.020.020.020.010.010.01
1385657
no rank
0.040.040.040.020.020.02
00.5000000.50.50000000000.50.5000.5

1655646
species
0.01
00000000000000000001

1655656
species
01000001100000000010001
0.020.020.020.010.01

0.040.240.040.020.02
000001000200.50000000000.50.5
genus
10852
0.020.120.020.010.01

species
10855
0.020.120.020.010.01
00000200040100000000011

0.3333330.5000000.3636360.33333300.50.83333306014.560.5
0.060.040.110.060.040.120.180.040.160.30.02
0.020.020.050.020.020.060.090.020.080.150.01
117574
no rank

0.020.020.02
1619678
genus
0.040.040.04
0.50000000.50.5

100000011
0.020.020.02
1609634
species

species
1676183
0.020.04
000000010002

000000000001
0.02
1676182
species

1655658
species
0000000000100000001
0.020.01

913970
species
01000000000001202912
0.020.090.020.080.15

0.020.020.020.030.010.010.010.04
251095
family
0.3333330000000.3333330.3333330.3333330000.400.33333300.33333300000.583333
0.060.060.060.090.050.030.030.12

0.010.01
251096
genus
0.020.02
00000000000000.50000.5

12585
species
0.010.01
000000000000010001

genus
0.020.020.020.030.010.010.04
104766
0.50000000.50.50.50000.500.50000000.875
0.040.040.040.060.020.020.08

00000000000000000000002
0.03
67585
species

0.03
0000000001
species
753670

0.010.01
00000000000001000000001
species
1436892

species
283824
1000000110000001
0.020.020.020.01

35342
0.070.030.050.010.010.020.010.030.020.05
no rank
0.130.070.10.020.020.040.020.050.050.1
00000.4615380.85714301000000.5000.5010.50.40.40.8

642260
species
000000000000000000001
0.01

species
1327968
00001202000000001000112
0.020.020.050.010.010.010.03

000011
0.020.01
1454024
species

00000000000000000000001
0.01
species
642253

1592095
species
0.01
00000000000001

0000000000000000000000.5
0.02
species
1249501
0.01

no rank
1114942
0000000000000000000001
0.01

species
1592101
00001
0.02

1454022
species
00000000000000000000001
0.01

0.01
00000000000000000001
642256
species

0.01
000001
1454030
species

species
1592098
0000000000000000002
0.02

0.3076920.33333300.2424240.2424240.59259300.2857140.4285710.3461540.30.30769201.0416700.8666670.4583331.123460.1818180.4864860.20.7450980.727273
0.260.120.330.330.270.140.280.520.20.260.240.150.240.810.220.370.150.510.66
family
10811
0.070.040.090.10.070.050.070.150.060.080.070.050.070.270.070.110.040.130.2

0.02
2022857
genus
000000000000.5
0.04

species
1306546
0.02
000000000001

0.020.02
0000000000000000000.50000.5
10812
0.010.01
genus

0000000000000000001
0.01
species
1302849

889510
species
0.01
00000000000000000000001

0000000000000000000.5
0.02
no rank
70692
0.01

0.01
0000000000000000001
species
1188812

0.4210530.500.3333330.3478260.800.4444440.5714290.4864860.4285710.42857101.4705901.30.6470591.685190.1818180.6923080.27272711.06818
0.190.080.240.230.20.090.210.370.140.140.170.10.170.540.110.260.110.380.44
genus
10814
0.070.040.090.10.070.050.070.150.060.060.070.050.070.270.040.110.040.130.19

0
00000000000000000
species
1195641
0

no rank
290090
0
00000000000000000

species
47838
0
0000000000000000000

species
407837
0.02
00000001

53260
species
0.01
000000000000000000

0.01
00000000000000000001
361731
species

207240
species
01
0.02

000000000000000001
0.01
species
270146

0.01
00000000000001
species
932071

species
123735
0000000000000000000
0.01

00000000000001
0.01
species
10815

681518
species
0000000000
0.02

00000000000000000100002
0.010.03
220340
species

0.010.01
00000000000000000001001
species
180526

00000000000000000
0
species
219744

0.02
0001
species
1266994

00000000000000002
0.02
619599
species

0.02
0001
species
223316

0000000000000000040
0.050
species
71186

0000000000010002
0.020.02
77647
species

species
759390
0000000000000003
0.03

10000101001
0.020.010.020.02
species
1266977

223264
species
0
00000000000000000

333361
0.01
species
0000000000000000000000.5
0.02

293284
no rank
0.01
0000000000000000000001

species
0.02
77811
0.04
0.5

0.02
1
no rank
429564

0.01
00000000000001
507476
species

species
325111
00000000000001
0.01

333968
species
0.03
00000000000000000000002

223345
species
00000000000000000000004
0.05

0.01
000000000000000000
species
435646

00001
0.02
species
202568

1246650
species
01
0.02

species
377462
00000000000000000
0

0.01
00000000000000000001
species
1195378

species
1513531
0.02
000001
0.04

395349
no rank
0.02
000002

211866
species
000000000000000001
0.01

85752
species
00001
0.02

713949
species
1
0.02

00000000000000000001
0.01
652718
species

0.020.050.050.030.070.060.020.020.010.030.010.030.040.030.120.03
291027
no rank
0.5000.3636360.4444441.42857000.85714310.50.500.5000.8333330.50.3333331.50.3333331.60870.4
0.040.110.090.070.140.120.040.040.020.060.020.060.080.060.230.05

species
302100
0000000000000000000001
0.01

498803
0.02
species
0000.5
0.04

0001
0.02
498805
no rank

559878
species
000001
0.01

0000000000000000000001
0.01
223308
species

0000000000000000001
0.01
1874886
species

935473
species
0.020.030.02
000013000001

0.02
00000000001
452758
species

species
1200970
0.01
0000000000000000000001

0.02
1
932073
species

1391746
species
0.01
00000000000000001

0000100020000000000303
0.020.050.010.040.03
species
604276

species
1225069
0.01
00000000000000000000001

00000000000000000010051
0.010.050.01
species
667119

0.020.06
0000000012
species
1825931

0.02
0001
222441
species

0.01
00000000000001
species
574890

0.02
00000000000000002
species
1311277

1859135
species
000000000000000001
0.01

0.02
000000000000000000000.5
species
0.01
417287

417288
no rank
000000000000000000001
0.01

species
1266458
000000000000000000001
0.01

00000000000005
0.04
676044
species

0
0000000000000000000
species
172394

100755
species
0000000000
0.02

00000000000000000001
0.01
species
86502

000001
0.01
649996
species

000000000000000002
0.02
species
310103

species
10831
0000000001
0.03

000000000000000006
0.07
species
999288

0
00000000000000000
species
128941

266799
species
0.020.020.01
00000000001100001

species
0.010.04
928315
0.020.08
00000000000000000000.5001.5

00000000000000000001003
0.010.04
no rank
913600

000000000100000001
0.030.01
390436
species

species
297249
0.01
000000000000000000001

species
223313
000000000000000014
0.010.05

0.9259261.127520.3658540.4260870.4587630.7463771.217710.8070180.3407410.3333330.3529410.4583330.6835442.171950.50.4141411.786160.9933770.4390241.033820.8947370.7150840.541958
1.891.491.231.151.941.382.711.711.350.810.340.961.582.210.840.991.591.511.642.072.281.792.85999999999999
no rank
0.620.50.420.40.670.440.930.590.470.270.110.320.530.720.280.320.510.530.570.720.770.610.99
1922347

1.270.990.810.751.270.941.781.120.880.540.230.641.051.480.560.671.080.971.071.351.511.181.87
1.377951.696970.5555560.6533330.7007871.095741.853931.232140.5227270.50.5217390.68751.028573.236490.750.611942.629631.536080.6728971.585191.350991.084750.828877
0.620.50.420.40.670.440.930.590.470.270.110.320.530.710.280.320.510.510.570.720.770.610.99
1922348
no rank

1923625
species
000020000000000000002
0.050.03

00000000001
0.02
species
1922540

0.020.050.020.020.020.020.01
10002210001000000020001
1923095
species

0.02
0000001
1923717
species

00000000000001
0.01
1922560
species

1923406
species
0.010.01
0000000000000200001

0.010.01
00000000000002000000001
1923634
species

0.03
00000000000000000000002
1923631
species

0.020.030.010.01
1000000001000000001001
1923601
species

0.050.020.010.010.010.010.01
00000021000002000001111
species
1923754

1922712
species
01
0.02

species
1923268
0.01
0000000000000001

0.030.070.050.010.050.010.01
00132120000000000011
species
1922352

0.020.04
0000000010001
1923448
species

1922930
species
000001
0.01

species
1922710
0.03
0000000001

1922632
species
0.020.01
0100000000000000001

1922624
species
0.02
00001

0000000000000000000001
0.01
1923119
species

00000000000002
0.01
species
1922659

species
1923118
00001
0.02

1922789
species
0.01
00000000000000000000001

0.010.020.03
000001010000000000002
species
1922551

1923322
species
0.05
000000000000000000004

0000000000000100001
0.010.01
1923138
species

1923205
species
0000000000000001
0.01

1922706
species
0.01
00000000000000001

species
1923128
00000000000000000000003
0.04

1922677
species
00000001
0.02

0.01
00000000000000000001
species
1923610

000000000000400002
0.160.02
species
1922886

0.01
0000000000000000001
species
1923609

1923486
species
0.020.03
000000000001000000002

1923272
species
0000000000000001
0.01

0.010.010.01
00000000000001000010001
1923132
species

1
0.02
1923137
species

1923485
species
0.04
2

000000000000000001
0.01
species
1922598

0.02
0000000000000002
1922497
species

species
1922879
0.01
000001

0.01
0000000000000001
species
1922760

species
1923731
0.010.02
00000000000002002

0.02
00000000000000002
1923213
species

species
1923104
0.020.020.01
00001010000000001

0.03
00000000000000000000002
species
1922792

1923557
species
0000000000000000070001
0.090.01

0000100000000100000101
0.020.010.010.01
1923755
species

1923340
species
0.020.020.01
10000010000001

species
1922796
0.04
000000000000000003

species
1922814
000000000000000000001
0.01

species
1922861
0.01
0000000000000000000001

species
1923195
0.060.02
31

0.050.010.05
000000200000000010004
1923487
species

1922673
species
0.01
0000000000000001

species
1922431
0.010.01
00000000000000011

1923614
species
100001
0.020.01

00001000000000001
0.020.01
species
1923460

000000101
0.020.02
1922658
species

species
1922768
0.01
0000000000000000001

000000000000000001
0.01
species
1923612

0.020.010.01
1000000000000000010001
1922934
species

0.01
000000000000000000001
1922387
species

0.02
0000000000000000002
species
1923127

1923264
species
0.01
0000000000000000001

1923647
species
000000001
0.02

species
1922587
3000010000000000001011
0.060.010.010.010.01

0.01
0000000000000001
1923554
species

0.01
0000000000000000001
1922896
species

1922683
species
000000001
0.02

0.01
000001
species
1923605

1923248
species
0.020.030.01
00000001010000000001

0.04
00000000000000000000003
1922638
species

1922636
species
0.04
0000000000001

species
1923604
0.020.010.020.01
0000110000000000002001

20000000000001
0.040.01
1922931
species

0.02
000000001
species
1923507

species
1922791
0.010.03
00000000000000000000102

0.03
00000000000000000000002
1923451
species

1922771
species
000000001
0.02

1923468
species
0.020.050.020.010.040.02
0100000210000000100302

0.04
000000000000000000003
1923696
species

species
1923686
0.04
0000000000001

1923597
species
00000001000001
0.020.01

1922622
species
0000000000000001011
0.010.010.01

0000001
0.02
1923191
species

1923117
species
0.01
00000000000000000000001

1922684
species
0.02
01

0.020.020.01
010101
species
1922561

0100000000000001
0.020.01
species
1923377

01000002000000000000001
0.020.050.01
species
1922385

0.02
0000001
1922386
species

000001
0.01
species
1922644

1922432
species
000000000000000000001
0.01

1923493
species
0.020.01
0000000100000001

0000000000000000022
0.020.02
species
1923157

0.020.03
000000000001000000002
1923742
species

0.130.170.030.010.040.01
7810010000001000000001
1923100
species

species
1923263
01
0.02

0.020.020.070.010.010.01
01013100000000000011
1923327
species

species
1922557
000000000000000006
0.07

species
1923415
0.01
0000000000000001

1923740
species
00000000000000000000001
0.01

0.02
0001
species
1923170

species
1923622
0.020.010.020.040.01
00000000100001000020301

species
1923644
0.020.01
00001000000000000000001

species
1922544
0.020.01
000101

species
1923276
0.010.01
00000000000000000000101

1923615
species
0.02
0000000000000000000002

0.01
00000000000001
1923648
species

species
1922905
00000000000101001
0.020.010.01

species
1923164
00000000000000000000041
0.040.01

species
1923401
0.030.01
00000000010001

species
1922893
0.020.020.030.010.020.030.04
00000011000000003012033

00000000000100000000001
0.020.01
species
1923344

1923108
species
0.020.01
0000100000000000000001

species
1923188
0.01
00000000000000000000001

species
1922651
0.020.020.01
00000200000100000001

0.01
00000000000001
1923109
species

species
1923627
0.020.010.010.01
00001100000001000000001

1
0.02
1923154
species

000000000000000000006
0.08
1922615
species

species
1923002
0000010000000001000001
0.010.010.01

0.01
000000000000000001
species
1923467

0000000000000001
0.01
1923135
species

species
1922629
0.020.030.020.01
00000010010000000020001

species
1923769
0.01
0000000000000001

1922550
species
0.02
0000001

0.01
0000000000000000001
1923593
species

1922608
species
0000000000000110101
0.080.010.01

0000000000002
0.08
species
1922444

0.03
0000000001
species
1923654

1923481
species
0.01
00000000000002

00000200100000000101
0.020.020.010.01
1923449
species

0.01
00000000000002
species
1923079

1923723
species
0.020.010.020.01
00000000100001000021

1923112
species
0.01
00000000000000001

0.01
000000000000000001
species
1923735

1923652
species
0.01
0000000000000001

00000000000000001
0.01
1922626
species

0.010.01
000000000000010001
species
1923166

species
1922621
0.020.020.01
00000010000000000000021

species
1923701
0.020.020.020.010.01
11000010000001000001

species
1922932
0000011
0.010.02

1923156
species
0.01
0000000000000001

0.02
000000001
1923479
species

0.01
0000000000000001
species
1923651

00000011000001
0.020.020.01
1923372
species

1922772
species
0001
0.02

species
1923476
00000000000001
0.01

species
1923619
0.01
00000000000000000000001

0.01
0000000000000001
species
1922992

0.020.020.020.010.01
00000011100000000000011
1922549
species

species
1923003
00000000000000001
0.01

1923246
species
00000000000001
0.01

1923641
species
0.02
00000000000000000002

species
1923355
0000000000000000000001
0.01

10001100100000000000011
0.020.020.010.020.010.01
species
1922558

0.01
00000000000000000001
species
1923488

1923761
species
00000000000000000000001
0.01

1922667
species
0.02
00000000000000000002

species
1923496
0.14
000000000000020

00000000000000000000001
0.01
1922871
species

0.05
000000000000000000004
species
1923115

species
1923780
00000000000000004
0.03

0.04
0000000000001
species
1922889

1922563
species
0000000000000001
0.01

species
1923703
0.020.020.020.060.010.010.010.01
00000201100301000111

species
1923143
0.02
0001

1922936
species
0.020.03
0000000011

1923235
species
0.02
0000001

species
1923162
0.01
000000000000000000001

1922586
species
0.02
00000000000000002

1923559
species
0.05
0000002

0110000000000100001001
0.020.030.010.010.01
species
1923778

00000000000000000011012
0.010.010.010.03
1922730
species

species
1923366
0.01
00000000000000000000001

1922606
species
0.02
1

0000010000000000000001
0.010.01
species
1923744

1922758
species
0.14
000000000000002

1923646
species
0.01
000000000000000000001

0.02
1
1923293
species

species
1923558
0.01
0000000000000001

0.01
00000000000000000000001
species
1923566

1923542
species
00200000100000001
0.060.020.01

1923126
species
0000000000000000000001
0.01

species
1923777
000000000000000000001
0.01

1923136
species
0.030.02
00000000010000000002

000012100000000000101
0.020.020.020.010.01
species
1923599

species
1923009
0.03
00000000000000003

species
1923748
0.02
01

species
1922391
0.020.020.010.050.01
00011100000000000004001

1923187
species
0.01
000000000000000001

1922552
species
000000000000001
0.07

1923416
species
0000060000000000000001
0.050.01

0.02
00000000000000002
species
1922657

0.01
0000000000000001
species
1922822

00000000000000001
0.01
species
1923745

1922818
species
0.02
0000001

species
1922824
0.01
00000000000001

species
1923345
0.01
0000000000000001

species
1923420
0.020.010.010.01
01000000000000000010101

species
1922990
0.02
1

1923113
species
00000010000001000000001
0.020.010.01

0.01
00000000000000001
1923716
species

0.01
00000000000000001
species
1922541

species
1922767
0.020.020.01
1000100000000000001

000000000000000000001
0.01
1923608
species

0.02
1
1922734
species

1922634
species
0.020.01
00000000000000000002001

0.01
0000000000000000000001
species
1923107

00100000000000000000001
0.030.01
species
1923695

1923220
species
00001000000000000000002
0.020.03

00000001
0.02
species
1922798

species
1922531
0.01
00000000000000000001

0.03
0000000000000000003
1922862
species

1923393
species
0.03
00000000000004

00010000100100010111001
0.020.020.020.010.010.010.010.01
1922997
species

1922720
species
00000000000000000001
0.01

0.030.020.01
00100200000000000000001
1922872
species

1
0.02
1923142
species

00000000000001001
0.010.01
1923603
species

1923753
species
0.01
000000000000000001

1922884
species
001
0.03

1922524
species
0.01
0000000000000000001

0.01
0000000000000000000001
species
1922553

1922619
species
000000000000000000001
0.01

0.01
0000000000000001
species
1923219

1923374
species
00000000000000000000003
0.04

0.020.010.070.01
00001000000001000000061
1923564
species

1922434
species
0001100000000300001
0.020.020.020.01

species
1923124
0.01
00000000000000000000001

1923689
species
0.04
0000000000001

0.010.01
0000000000000100000001
1922652
species

0.020.020.020.050.030.020.010.060.010.04
01010010210102000005103
species
1922942

species
1923193
0.03
001

species
1922530
0.01
00000000000000000000001

1923201
species
0.05
00000000000007

1
0.02
species
1923715

0000000000010001
0.020.01
1922904
species

0.01
0000000000000000000001
species
1923532

0.01
000001
1923163
species

00001
0.02
species
1923075

0.010.03
00000000000000013
1923693
species

0.01
00000000000000000000001
1923209
species

0.01
0000000000000000001
1923368
species

0.01
0000000000000000001
1923159
species

1923408
species
0.02
01

1923751
species
0000000000000000000002
0.02

000000001
0.02
species
1923763

species
1922892
001000000000000001
0.030.01

species
1923691
0.020.03
00000000000000000020002

000000000000000001
0.01
1922937
species

0.020.01
00000001000000000000001
1922588
species

1922429
species
0.01
00000000000000000000001

0.020.01
00000000000100000001
species
1923628

species
1922769
0.020.020.01
000010000000000000021

00000000000000007002201
0.060.020.030.01
species
1923706

0000000000000000000021
0.030.01
species
1922784

species
1922793
0.01
00000000000001

0.010.01
0000010000000000001
1923726
species

0001
0.02
species
1923129

1922546
species
0.020.01
00000000001000000001

1922866
species
0.01
000001

0.040.01
000000000000100000001
1923477
species

0.030.020.01
001000000001000001
1923134
species

1923280
species
0.030.010.020.010.02
00100101000001000002

00000000000001
0.01
species
1923076

species
1922596
0.01
00000000000000001

1923450
species
0.010.03
00000100000000000000002

0.01
00000000000000000001
species
1923097

0001
0.02
1923245
species

1923764
species
0.04
000000000000000000003

1923489
species
0.01
00000000000001

species
1923122
0.010.01
0000000000000001000001

1922440
species
000000106
0.250.15

1922786
species
00000000000000000000201
0.030.01

1922998
species
000001
0.01

0.02
0000000000000000000002
1923161
species

0.01
000001
1922811
species

1922895
species
0.01
00000000000001

species
1922835
00000000000000000009
0.11

00000000000002
0.01
species
1923444

0.01
0000000000000000001
1922660
species

1922991
species
000000000000000102
0.010.02

species
1923331
0000100000000002
0.020.02

0.010.010.03
0000000000000000000113
species
1922857

0000000000000112
0.010.070.02
1923534
species

0.010.01
00000000000000000000011
species
1922707

0.12
000000000000000014
species
1922885

000000000001
0.02
1922867
species

0.01
00000000000000001
species
1923106

00000000000000001
0.01
species
1922661

species
1923326
0.010.01
0000000000000000010001

000000000000000001
0.01
1923373
species

1922833
species
0.01
0000000000000000001

species
1923639
0.01
000000000000000001

0.030.01
001001
species
1923537

1923563
species
0.01
00000000000001

0.020.010.03
00000001000000000010002
species
1923441

0.01
000001
1923659
species

species
1923123
0.01
00000000000001

0001000000100000000002
0.020.020.02
species
1923249

00010500001103000110001
0.020.040.020.020.020.010.010.01
species
1923743

1923699
species
0.01
00000000000000000001

1923231
species
0.010.02
000001000001

00000000000000001
0.01
1923494
species

1938656
species
0.02
00000000001

0.01
000000000000000001
species
1926998

0.01
00000000000001
1926997
species

439488
1.190.961.281.121.431.191.280.821.241.110.631.050.651.771.591.841.21.761.251.441.471.321.73
no rank
7.769999999999975.819999999999997.976.329999999999988.689999999999977.909999999999967.469999999999984.439999999999997.469999999999976.924.046.699999999999973.8910.999.5200000000000111.357.6799999999999710.347.709999999999968.559999999999978.759999999999977.9299999999999710.79
0.2947230.2199310.3099120.5513430.4384350.5802780.388220.2229730.3199460.1907510.2227720.1850750.2879184.765240.2794129.672252.308597.889750.3177690.3761680.5022830.4073140.564411

no rank
35278
0.670.541.091.031.140.91.030.620.890.780.440.780.240.770.620.520.60.450.760.860.940.711.08
0.3236070.3054550.3961940.7163870.5972460.8126230.4990060.2873130.4828380.2153470.303030.2291170.20.9708740.2456140.4007350.524390.4221110.4264710.4712640.3377190.40.723708
3.772.755.784.759999999999995.809999999999985.069999999999995.029999999999992.684.374.039999999999992.314.191.24.119999999999982.852.719999999999993.279999999999991.994.079999999999984.349999999999994.559999999999983.449999999999995.60999999999999

000000000000000010000.333333
0.090.03
0.030.01
1513277
genus

1549608
0.030.01
no rank
00000000000000001.50000.5
0.060.02

1100890
species
0.030.01
000000000000000030001

family
11050
0.020.030.10.020.040.010.010.020.020.020.090.010.03
0.080.160.350.10.140.030.040.070.070.080.360.050.09
00000.250.250.6571430.2000.285714000.33333300.250.2857140.2857140.2500.5555560.20.222222

000000.333333
0.06
11102
0.02
genus

no rank
1249508
0.02
0.04
000000.5

species
1325446
000001
0.01

0.01
000001
species
1562038

0.010.010.01
11095
genus
0.030.030.04
000000000000000000.3333330.3333330000.25

0.020.020.03
000000000000000000.50.50000.333333
63426
0.010.010.01
no rank

0000000000000000001
0.01
1562066
species

species
1914447
0.01
0.02
00000000000000000000000.5

no rank
1873455
0.01
00000000000000000000001

188938
species
000000000000000001
0.01

0.040.04
0000000.50000.5
0.020.02
1307799
genus

0.020.02
00000010001
species
1307801

0.010.010.010.01
38144
no rank
00000000000000.5000.50.500000.5
0.020.020.020.02

species
1485186
0.01
00000000000001

00000000000000000000001
0.01
species
1963247

1286143
species
0.010.01
000000000000000011

genus
11051
0.020.020.070.020.020.010.010.010.090.01
00000.3333330.28571410.25000.33333300000.3333330.33333300.33333300.7407410.25
0.060.070.210.080.060.030.030.030.270.04

0.140.02
0000001.500000000000000.5
0.070.01
11071
subgenus

species
11079
000000300000000000001
0.070.01

0.030.060.03
000000.33333300.33333300000000000000.333333
0.010.020.01
29263
subgenus

0.020.040.02
000000.500.500000000000000.5
species
0.010.020.01
33743

172148
no rank
0.010.020.01
0000010100000000000001

no rank
39686
0.010.020.010.01
0.020.040.020.02
000000.500000.500000.5000.5

species
643132
0000000000000001
0.01

64318
species
0.01
0000000000000000001

0.01
000001
1302179
species

0.02
00000000001
species
631267

0.020.010.03
40005
subgenus
0.040.020.06
00000.5000000000000.50001

000010000000000010002
0.020.010.03
species
64295

29262
0.01
subgenus
0.02
000000000000000000000.5

species
64283
0.01
000000000000000000001

000000000000000000001.5
0.08
29261
0.04
subgenus

species
64293
0.04
000000000000000000003

genus
12137
0.020.010.01
00000.3333330000000000000000.3333330.333333
0.060.030.03

no rank
502177
0.020.010.01
00000.50000000000000000.50.5
0.040.020.02

0.02
00001
685899
species

species
1602124
0000000000000000000011
0.010.01

0.240.120.080.10.120.060.050.210.040.160.040.14
0.41666700.250.2500.50000.2500.33333300.40.3333330.2500000.43750.250.214286
186766
0.060.030.020.030.030.020.010.070.010.040.010.04
family

0.55555600.3333330.33333300.7142860000.33333300.500.50.50.33333300000.5833330.3333330.3
0.180.090.060.070.090.040.040.140.030.120.030.1
genus
186768
0.060.030.020.030.030.020.010.070.010.040.010.04

species
88388
000000000000001
0.07

88387
species
00000200000100000000001
0.020.020.01

no rank
0.060.030.020.010.030.010.010.040.010.03
343865
0.120.060.040.020.060.030.020.080.020.05
0.83333300.50.500.50000.50000.66666700.500000.8750.50.4

0.01
00000000000000000000001
279538
species

00000000000001
0.01
1826822
species

100000000000000000001
0.020.01
1837089
species

species
1562379
0.03
0000000001

0.01
000001
species
1816487

001
0.03
1629665
species

species
1162083
00010000000000000000001
0.020.01

0000000000000001
0.01
1848150
species

1562380
species
2000000000000100000021
0.040.010.030.01

0.320.280.060.080.370.50.370.410.180.120.140.590.380.370.330.150.270.250.220.6
0.31250.28571400.3333330.250.5675680.520.4324320.4390240.3333330.3333330.28571402.3220301.105260.9729731.03030.2666670.370370.280.5909091.21667
family
0.090.080.020.020.110.170.120.140.060.040.040.160.110.110.110.050.10.090.080.19
39729

0.060.02
00000000000000000.6666670.5
156208
0.020.01
genus

0.04
00000000000000001
no rank
596956
0.02

46076
species
0.02
00000000000000002

0.01
000000000000000001
species
128818

00.3333330000.333333000000.33333300000000.5
0.060.030.060.02
genus
0.020.010.020.01
137757

0.020.010.02
397543
no rank
00.50000.5000000.5
0.040.020.04

010001000001
0.020.010.02
species
1890365

species
946046
0.01
00000000000000000001

0.020.050.010.010.03
1230390
genus
0.040.10.020.020.06
0000000.51000000000.50000.501

species
53954
0.01
000000000000000000001

00000012000000001000002
0.020.050.010.03
431317
species

000000000000000000000.5
0.02
no rank
157019
0.01

000000000000000000001
0.01
species
1856031

156207
0.020.010.030.010.02
genus
00000100000000.501000.5001
0.040.020.090.020.04

0000000000000000001
0.01
460364
species

species
655837
0000020000000000000002
0.020.02

no rank
0.03
460363
0000000000000001.5
0.06

species
1414644
0.03
0000000000000003

31741
species
0.01
00000000000001

genus
12195
0.070.040.020.020.080.150.050.140.060.040.020.110.080.080.10.030.070.050.050.16
0.4210530.400.50.3333330.8823530.8275860.3636360.6666670.50.50.503.5806501.833331.722221.650.3750.6153850.40.91.91429
0.190.10.040.060.170.290.110.270.120.080.040.310.180.180.20.080.130.10.10.35

00000000201
0.050.02
species
291286

341167
species
0.01
00000000000000000001

12222
species
00000000000000000001
0.01

0.02
000000001
41460
species

000000001
0.02
species
31743

species
299200
0.01
0000000000000000000001

species
187977
0.020.040.01
010000000000000051

000000000000000000001
0.01
12196
species

species
0.01
115232
0.02
0000000000000000000.5

204478
no rank
0.01
0000000000000000001

0.030.040.08
00000300000000040000006
32620
species

no rank
332470
0.040.020.020.020.090.010.010.010.010.010.03
0.080.040.040.040.190.020.020.020.020.020.06
0.50.5000.5000.5000005.7368400.50.50.50.500.501

0.09
000000000000012
species
1406816

0000000000000000001
0.01
796352
species

0.01
00000000000001
species
945960

species
861561
00000001
0.02

0000000000000001
0.01
species
1888953

000010000000000000001
0.020.01
species
310681

515446
species
0.020.020.03
11000000000000000000002

000000000000000001
0.01
species
1963260

species
1072517
10000000000000001
0.020.01

species
91613
000000100000000000001
0.020.01

0.01
000001
species
12208

0.01
000001
12211
species

0.010.01
000001000000000001
373398
species

0000002
0.05
453050
species

species
79917
0000001
0.02

201862
species
0.01
000001

930881
species
0.03
0000000001

species
46917
00000000000000001
0.01

species
12205
0.01
000000000000000000001

0.020.010.040.030.060.01
00000000100002043501
species
12210

515575
species
0.020.04
00010000000000000000003

species
43130
00000000000000000000001
0.01

000000000001
0.02
221262
species

00000000000000000011
0.010.01
species
187978

species
443746
0.020.020.02
00000000101000000002

0.02
0000000000000000000002
species
87097

1000012000000000000002
0.020.010.050.02
species
12844

0.01
000001
species
32619

0000000001
0.03
species
1220025

33763
species
0.020.02
10000001

0.040.020.040.020.02
00.50000.500.5000000000000.50.5
675844
0.020.010.020.010.01
genus

323118
species
0.020.010.020.010.01
010001010000000000011

00000000000000.5
0.02
0.01
39731
genus

0.01
00000000000001
species
12465

0.50000000000003
0.040.08
39730
0.020.04
genus

40666
species
0.020.04
10000000000006

0.02
00000000000000.5
0.01
186783
genus

species
544686
0.01
00000000000001

0000000.50000001
0.040.02
0.020.01
2022856
genus

0.020.01
00000010000002
1094249
species

39733
0.090.060.020.120.020.120.030.060.040.020.020.020.020.040.02
family
0.350.220.060.420.080.430.120.240.160.070.080.060.060.170.06
0.7428570.45454500.3333330.5952380.5000.5813950.250.750.250000.2857140.500.3333330.3333330.4117650.333333

10.500.50.4444440000.50000000.5000.50.500.5
0.080.040.040.090.040.020.040.040.02
no rank
0.040.020.020.050.020.010.020.020.01
352926

0000100000000001
0.020.01
species
1141625

species
645687
00010000000000000001
0.020.01

0.020.01
0000100000000000001
species
1074210

species
1247114
0.02
000000001

1247113
species
0000000000000000000001
0.01

0.040.020.01
21000000000000000001
species
683172

0000000000000000001
0.01
species
1093928

0.060.060.060.090.02
000000000.333333000.33333300000.6666670000.6666670.5
genus
0.020.020.020.030.01
249588

1239574
species
0.01
0000000000000000000001

no rank
526119
0.020.020.020.03
000000000.5000.5000010001
0.040.040.040.06

1435485
species
000000000000000000002
0.03

1027245
species
00000000000000002
0.02

0.02
000000001
species
1662272

000000000001
0.02
species
1105379

10.6666670010.6666670010.33333310.3333330000.33333300000.25
0.180.120.210.060.210.090.180.060.030.04
genus
0.060.040.070.020.070.030.060.020.010.01
249589

no rank
336962
0.060.040.070.020.070.030.060.020.010.01
0.120.080.140.040.140.060.120.040.020.03
1.51001.51001.50.51.50.50000.500000.333333

905828
0.01
species
000000000000000000000.5
0.02

no rank
622438
0.01
000000000000000000001

0.060.040.070.020.070.030.060.020.01
3200320031310001
species
336959

0.060.220.230.050.040.040.040.160.010.010.030.10.01
39740
family
00.750.8815792.768120.2666670.86666700000.250.28571402.56923000.3333330.250.55555601.088240.333333
0.240.760.690.150.150.160.140.650.030.040.090.340.03

0.02
12320
genus
0.04
0000.5

species
12321
0001
0.02

genus
0.060.10.020.030.040.020.160.010.010.040.01
12316
0.180.280.060.090.120.060.470.020.030.120.02
010.53571400.3333331.3333300000.3333330.33333303.53191000.50.333333000.5833330.5

747056
species
0000000000000000000001
0.01

0.010.01
00000000000001001
species
243563

64950
0.060.060.020.030.040.020.150.01
no rank
0.120.120.040.060.080.040.30.02
01.5100.5200000.50.505.50000000.5

0.060.060.030.020.06
03200400001008
269213
species

0000100000110130000001
0.020.020.020.090.01
species
12319

0.030.010.03
64949
no rank
0.060.020.06
000.5000000000000000.5001

001000000000000000002
0.030.03
species
56486

000000000000000001
0.01
37127
species

0.260.420.040.020.040.020.040.12
0024.50.50.5000000.500.50000102.5
genus
0.130.210.020.010.020.010.020.06
12300

species
12303
0.020.020.06
000010000000000000205

12301
species
0.020.01
00000000000101

species
300879
004901
0.130.210.01

0000000000000000000.5
0.02
genus
0.01
12304

species
578305
0.01
0000000000000000001

0.420.06
000000000000000.6666670000000.666667
1513233
0.140.02
family

0.280.04
0000000000000010000001
0.140.02
143901
genus

76343
species
0000000000000000000002
0.02

species
1324959
000000000000002
0.14

1283207
0.010.02
family
0000000000000000.3333330.666667
0.030.06

0000000000000000.51
0.020.04
85651
0.010.02
genus

species
37206
00000000000000012
0.010.02

39738
0.020.020.020.010.010.010.010.02
family
0.060.060.060.030.030.030.030.06
00.333333000.33333300.3333330000000.33333300.33333300.3333330.333333000.333333

genus
0.020.01
12141
0.040.02
00.500000000000000.5

species
12142
0.02
01

species
208086
0.01
0000000000000001

0.02
0000000000000000000000.5
genus
12266
0.01

species
12268
0000000000000000000001
0.01

0.020.01
1299311
genus
0000000.50000000.5
0.040.02

00000010000001
0.020.01
12056
species

genus
0.010.01
39734
0.020.02
0000000000000000000.5000.5

193120
species
0000000000000000001001
0.010.01

genus
1911601
0.01
0.02
000000000000000000.5

species
11987
0.01
000000000000000001

genus
0.02
186844
0.04
00000.5

00001
0.02
1352511
species

464095
0.090.060.380.260.40.280.340.220.210.150.130.120.120.10.280.140.180.060.150.30.140.140.13
order
0.2439020.250.5419350.6526321.310560.7031250.6363640.51.43210.2272730.40.2127660.20.7111110.1680.4285710.50.250.3898310.7164180.3454550.3015870.208333
0.410.241.550.951.611.281.320.780.810.660.550.470.60.451.250.630.880.20.591.340.550.630.48

no rank
0.010.010.010.010.01
464925
00000000000000.500.500.500000.5
0.020.020.020.020.02

species
439015
00000000000001000100
0.010.010.01

0.010.01
00000000000000010000001
species
439016

family
232795
0.020.010.020.020.020.030.050.020.030.010.050.030.030.01
00.3333330000.3333330.3333330.3333330.3333330.3333330001.6666700.3333330.5555560.3333330.46153800.6666670.5555560.333333
0.060.030.060.060.060.090.150.060.090.030.130.090.090.03

no rank
0.020.020.010.020.01
336635
00000000.50.50000000.5000.5000.5
0.040.040.020.040.02

species
932662
0000000000000000001001
0.010.01

0.02
00000001
species
1432856

1926968
species
0.020.01
0000000010000000001

0.01
0000000000000001
1926967
species

genus
144051
0.010.050.020.01
0.020.10.040.02
000000.500000002.50010.5

00000100000006
0.010.04
64279
species

0.010.01
000000000000010001
species
12136

0.02
00000000000000002
species
209529

genus
0.020.010.010.02
1213379
00.500000000000000.50.500001
0.040.020.020.04

0000000000000000000002
0.02
68876
species

species
294365
0.020.010.01
01000000000000011

0.040.060.040.060.02
0000000.5000.50000000010100.5
1922209
0.020.030.020.030.01
genus

00000000000000000020001
0.020.01
species
81583

species
92395
0.03
0000000001

0.020.03
000000100000000000002
64698
species

001.5200.83333311000.50.500.50.500000100.5
0.20.180.060.10.10.040.040.020.140.060.02
675074
0.10.090.030.050.050.020.020.010.070.030.01
no rank

species
1926969
000000000000000000002
0.03

1931371
species
000000000000001
0.07

00000000001
0.02
species
1663127

0.02
000002
1884450
species

00000000000001
0.01
1931369
species

species
1931370
003401220001
0.10.090.010.050.050.02

1105380
species
0.01
00000000000000000000001

0.020.040.030.050.060.070.020.020.040.010.140.040.040.070.010.040.020.05
699189
family
0.3333330.3333330.3333330.66666700.523810.30.3333330.3333330000.250.3333330.1272731.333330.78571400.5714290.3333330.2727270.250.266667
0.060.120.090.150.210.20.060.060.160.030.550.120.140.210.030.110.080.15

0.50.50.5100.50.4444440.50.500000.5021.428570100.400.4
0.040.080.060.10.040.090.040.040.020.080.070.110.050.05
1111709
0.020.040.030.050.020.050.020.020.010.040.030.060.030.03
no rank

species
1345662
0.030.050.010.020.020.010.010.01
00120111000000000010101

0.040.03
00000000000000043
species
1675864

1591444
species
0.020.010.01
0100010000000000001

species
1920698
00000000000000001
0.01

species
1629132
00000000000000000000001
0.01

1965238
species
0.02
1

species
1464786
00000000000001
0.01

1911104
species
000000000000000000301
0.030.01

000000101
0.020.02
1410467
species

1345661
species
01
0.02

genus
232799
0.040.020.040.140.010.010.010.010.020.03
000000.8181820.5000000.33333300.17073200.33333300.3333330.50.50.3333330.4
0.110.040.120.410.030.030.020.020.060.05

species
1460071
0.020.01
00000010000000000000001

1111710
species
00000000000000000001
0.01

0000010000000.50000.500.5000.5
0.040.080.130.020.020.04
no rank
336476
0.020.040.070.010.010.02

0.020.01
00000200000000001
1690666
species

000000000000000
0.03
species
1936269

0000000000000000001001
0.030.010.01
1487185
species

1906245
species
0000000000001000000001
0.040.01

000000000000000000001
0.01
species
1541119

species
89463
000001
0.01

species
1993638
0.07
000000000000000.5
0.14

436447
no rank
000000000000001
0.07

0.020.01
00000200000000000000001
1521188
species

0.140.30.150.280.080.060.240.060.160.090.280.080.180.060.440.090.13
0.285714010.6666670.50.2500.333333000.6666670.3333330.250.3333330.250.250.3888890.33333301.2272700.2222220.230769
675072
0.040.10.050.10.020.020.080.020.040.020.070.020.040.020.110.020.04
family

0.020.040.01
680113
no rank
00000.5000001000000.5
0.040.080.02

1120751
species
00000000000000001
0.01

00001
0.02
species
367301

species
28351
0.04
00000000002

310684
0.10.050.020.040.02
genus
0.20.10.040.080.04
001.510000.50010.5

0.040.02
000000000021
202566
species

101688
species
00320001
0.10.050.02

12057
0.05
genus
0.1
00001

0.05
00002
species
12777

genus
0.010.020.01
675073
00000000000000000.50.50000.333333
0.020.040.03

470166
species
000000000000000011
0.010.01

000000000000000001
0.01
1425364
species

no rank
0.01
1238886
0.02
0000000000000000000000.5

1481465
species
0000000000000000000001
0.01

0.50000.5000000000000000000.5
0.040.040.02
12050
0.020.020.01
genus

species
51354
00001
0.02

0.020.01
10000000000000000000001
35287
species

0.060.060.120.070.210.060.10.330.040.07
0.33333300000.3333330000000.3333330.4285710.3333330.3333330.5001.6363600.250.285714
675075
0.020.020.040.020.070.020.030.110.010.03
subfamily

genus
0.020.020.010.070.010.010.070.010.01
12270
0.500000.500000000.50.50.50.33333300300.3333330.5
0.040.040.020.140.020.030.140.030.02

0.01
0000000000000001
species
12809

000001000000001
0.010.07
12273
species

173976
species
00000000000000000006
0.07

0.01
00000000000000000000001
species
65743

0.02
1
12615
species

352924
0.010.01
no rank
0.020.02
00000000000000000.500000.5

0.010.01
0000000000000000100001
species
1927714

0.01
00000000000001
species
12275

65068
species
0.01
000001

0000000000000.50.66666700.50001.5000.5
0.080.030.020.080.02
12258
0.040.010.010.040.01
genus

0.01
00000000000001
species
12262

species
12264
0.01
0000000000000001

species
128790
0.01
00000000000000000000001

0000000000001
0.04
species
328061

species
12261
00000000000000000003
0.04

0.01
00000000000001
species
12263

0.02
12269
genus
00000000000000001
0.04

00000000000000002
0.02
species
1432135

0.030.03
0000000000000000.3333330.333333
0.010.01
324901
family

0.020.02
0000000000000000.50.5
genus
0.010.01
324902

species
324900
00000000000000011
0.010.01

0.120.580.150.850.580.620.280.480.330.140.250.160.040.150.260.030.10.550.150.23
0.33333300.3620690.2666672.294121.20691.064520.8214292.333330.2727270.2857140.240.250.2500.2666670.7692310.3333330.50.7454550.2666670.434783
12058
0.040.160.050.290.160.20.10.160.090.040.080.040.010.040.070.010.030.170.050.07
family

genus
0.02
138954
0.06
0000000000000000000000.666667

0000000000000000000001
0.04
376147
0.02
no rank

species
172314
0000000000000000000002
0.02

000000.666667000000000000000.333333
0.060.03
1986961
0.020.01
genus

species
0.020.01
1986960
0.040.02
000001000000000000000.5

1661395
no rank
0.020.01
000002000000000000001

0.060.02
0000000000000000.3333330000.5
genus
688449
0.020.01

no rank
0.01
1330992
0000000000000000.5
0.02

0.01
0000000000000001
species
1547495

1330524
0.010.01
species
0.020.01
0000000000000000.50001

0.01
0000000000000001
651733
no rank

genus
0.030.020.01
12109
000.333333000.6666670000000000.333333
0.090.060.03

0.030.020.01
1606764
no rank
000.50010000000000.5
0.060.040.02

0010020000000001
0.030.020.01
1606765
species

genus
0.01
1481451
0.03
00000000000000000.333333

00000000000000000.5
0.02
0.01
1481452
species

0.01
00000000000000001
1481453
no rank

0.210.070.120.070.160.030.020.030.020.050.010.02
586418
genus
0.420.140.240.140.320.060.040.10.040.10.020.04
00004.542.51.53.50.500.5000010120.51

00000000000000001.5
0.06
species
2003649
0.03

00000000000000003
0.03
586420
no rank

0000985371010000102412
0.210.070.120.070.160.030.020.010.020.050.010.02
species
1330491

genus
1431456
0.020.01
0.060.03
0000.33333300.333333

0000.5
0.04
1562063
0.02
no rank

1562064
species
0.02
0001

0.01
1431460
species
0.02
000000.5

no rank
1431464
000001
0.01

194960
0.020.010.01
genus
0.060.030.03
0000000.333333000000000000.33333300.333333

0.020.010.01
655314
no rank
0000000.5000000000000.500.5
0.040.020.02

000000100000000000101
0.020.010.01
1659219
species

genus
0.020.030.01
12103
0.060.090.04
0000000.333333000.333333000000000000.25

1821749
0.020.03
species
0000000.5000.5
0.040.06

no rank
12104
0.020.03
0000001001

0.01
1821750
species
0000000000000000000000.333333
0.03

0.02
0000000000000000000000.5
204711
0.01
no rank

434309
no rank
0.01
0000000000000000000001

0.030.03
000000.33333300000000000000.333333
2025078
0.010.01
genus

0.020.02
000000.500000000000000.5
species
0.010.01
2025079

1578826
no rank
0.010.01
00000100000000000001

0.020.010.010.040.010.04
1330069
genus
0.060.030.030.120.030.12
00000.3333330.3333330000000.3333330.333333000001

no rank
0.020.010.040.010.04
1350000
0.040.020.080.020.08
00000.50.50000000.50.5000001.5

0.010.04
00000100000000000003
species
1477516

1477515
species
00001
0.02

00000000000011
0.040.01
1960206
species

000000000000
00.02
species
00.01
1330070

no rank
980964
000000000000
00.01

0.020.030.020.010.020.030.010.060.010.01
35296
no rank
0.040.060.040.020.040.060.020.110.020.02
0.500.500.50.500000.50.333333000000.500.6363640.50.5

species
1506581
00000000000100000002
0.020.02

species
1534547
0.0200.010.010.010.01
0000100000000000010101

1888309
species
1
0.02

00000000000000000001
0.01
1603963
species

species
1074865
0.02
00000000001

species
1958776
000001
0.01

1609159
species
001000000000000000001
0.030.01

00000000000000000001
0.01
species
1483681

12059
0.020.020.010.03
genus
0.040.040.020.06
00000.50000000.50000.51.5

0000000000000001
0.01
138949
species

species
1330521
0.020.02
000010000001

00000000000000003
0.03
147711
species

genus
686982
0.020.030.020.010.020.02
0.500.3333330.500.500.5000.333333
0.040.090.040.020.040.06

0.030.02
1073966
no rank
0.060.04
000.500000000.5

0.030.02
00100000001
species
1958777

0.020.020.010.02
10010101
686983
species

0.060.010.020.03
1330067
genus
000.666667000.3333330.333333000.333333
0.180.030.060.09

001000.50.5000.5
0.120.020.040.06
species
0.060.010.020.03
1330068

0020011001
0.060.010.020.03
no rank
1150861

no rank
1231622
0.020.020.010.030.01
0000.3333330.250.250000.333333000000.333333
0.060.080.040.090.03

0.01
674970
genus
0000000000000000.5
0.02

0000000000000001
0.01
species
674971

0.020.020.010.03
674962
genus
0.040.060.030.06
0000.50.3333330.3333330000.5

0.040.02
00000.50.5
0.020.01
1522178
no rank

000011
0.020.01
species
1522179

359987
species
0.020.03
0001000001

0.04
000000000000.5
0.02
675070
genus

000000000001
0.02
species
347219

family
0.070.080.140.050.050.020.120.140.090.060.060.060.030.020.060.10.10.090.150.03
69973
0.8695650.83333300.750.3076920.8666670.3333330.2857140.4186050.5555560.454545100.37500.5555560.3333330.56251.058821.517240.5357141.083330.25
0.230.240.440.130.150.060.350.430.270.220.180.240.090.060.160.340.290.280.480.08

00000.50000.500000.500.5000.33333300.333333
0.040.040.040.020.030.03
genus
1513308
0.020.020.010.010.010.01

00000000000000.500000.500.5
0.020.020.02
0.010.010.01
1654602
no rank

0.010.010.01
000000000000010000101
1654603
species

217686
species
0000000000000101
0.010.01

217615
species
000010001
0.020.02

12160
0.060.060.070.030.020.070.090.020.060.010.020.010.070.090.030.070.01
genus
1.51.501.50200.50.8571430.8333330.51.500.3333330100.52.066672.52941130.333333
0.120.120.140.060.040.140.180.040.120.030.040.020.150.170.060.140.03

species
326941
000000001
0.02

0.01
00000000000000000001
12161
species

no rank
0.010.010.01
349945
00000000000000.500000.50000.5
0.020.020.02

00000000000001000000001
0.010.01
1543207
species

0000000000000000001
0.01
species
167634

3303040122130002015626
0.060.060.070.030.020.050.060.020.060.020.010.060.070.030.07
64003
species

399314
species
0000000001
0.03

no rank
217429
0.020.01
0.5000000000000000000000.5
0.040.02

0000000000000000000001
0.01
species
940280

1493829
species
0.02
1

0000.7500.2500.333333000000.500010.666667
0.160.040.060.020.040.06
217160
0.070.020.020.010.020.02
genus

70177
species
0
000000

180903
species
0.05
0002

1381464
species
000000000000010002
0.010.02

no rank
362246
0.020.010.020.02
0000.500.500.500000000001
0.040.020.040.04

0001010100000000002
0.020.010.020.02
203168
species

0
000000
367121
species

genus
0.020.020.020.070.050.040.040.020.020.010.050.080.01
67753
0.040.040.040.130.110.120.090.040.040.020.10.170.02
00.5000.500.50.4615380.36363600.666667000.555556000.5100.50.80.5294120.5

species
72750
0000000000000000000001
0.01

00000000000001000000101
0.010.010.01
species
642478

species
52135
0.020.010.01
0000100000000000000101

00000000000000001
0.01
67754
species

0.020.020.02
010000010000000002
81931
species

0.02
00000001
species
267970

species
103881
0.020.01
0000001000000000000001

000000001000020000002
0.020.010.03
51330
species

000000000.501000.500000001
0.040.080.020.04
no rank
352927
0.020.040.010.02

558690
species
0.020.040.010.02
0000000010200100000002

species
31713
0000000000000000100001
0.010.01

species
227507
0.020.010.010.01
0000000100000100000011

76804
0.220.130.320.120.330.240.270.050.210.210.080.240.150.060.10.060.310.210.180.150.41
order
0.3065690.2133330.3413170.161290.4806631.690140.7462690.1666670.2201830.1551720.1818180.20895500.34117600.2647060.3606560.1666670.5523260.2807020.270.4285711.20909
1.370.751.670.621.811.421.340.241.091.160.441.340.850.340.610.31.721.1410.772.2

0.210.130.290.120.330.240.250.050.210.150.080.220.140.060.090.050.310.190.160.140.41
11118
family
1.070.621.260.51.481.181.010.190.880.740.361.040.640.280.470.211.410.850.720.591.79
0.3738320.2580650.4285710.20.5878382.03390.9702970.2105260.2727270.1621620.2222220.2500.42187500.3214290.4468090.1904760.6737590.3294120.3472220.5423731.48603

subfamily
694017
0.030.010.010.040.01
000000.55555600000000.3333330000000.3333331.333330.25
0.090.030.030.120.04

0.020.08
000000.50000000000000002
species
0.010.04
1986112

1631554
no rank
0.010.04
0000010000000000000004

00000100000000000000000.333333
0.040.03
genus
0.020.01
11155

237020
species
000002
0.02

0.02
00000000000000000000000.5
species
0.01
74501

46473
no rank
0.01
00000000000000000000001

694018
0.010.01
genus
00000000000000.50000000.5
0.020.02

405554
species
000000000000010000001
0.010.01

subfamily
0.210.130.290.120.330.210.250.050.210.150.080.220.130.060.090.050.310.190.140.10.4
693995
0.4651160.3265310.5567010.2631580.7565222.764711.289470.2857140.3582090.203390.2857140.31707300.55319100.4090910.5526320.250.8636360.4242420.452830.4848481.97761
0.860.490.970.381.150.850.760.140.670.590.280.820.470.220.380.161.10.660.530.331.34

0.1818180.251.031250.4444440.3333330.3333330.35294100.4615380.150.40.54545500.500.50.5454550.50.1538460.7307690.2500.678571
0.220.080.320.090.060.030.170.130.20.10.220.020.020.110.020.130.260.080.050.28
genus
0.060.020.130.050.020.010.070.070.060.040.080.010.010.030.010.030.10.030.010.08
694002

species
290028
0.020.020.020.020.010.010.02
00000010101101000102

694006
species
0001
0.02

000000.5000001
0.020.08
species
0.010.04
694009

227859
no rank
0.010.04
000001000002

0.120.060.090.050.030.030.12
0.3333330.3333330.33333300000000000000000.3333330.33333301
694005
0.040.020.030.010.010.010.04
species

0.020.030.010.01
11138
no rank
0.500.500000000000000000.5
0.040.060.020.02

10100000000000000001
0.020.030.010.01
11142
no rank

no rank
31632
0.020.020.010.010.04
0.040.040.020.020.08
0.50.50000000000000000000.501.5

no rank
502102
0.020.020.010.010.04
11000000000000000000103

0.040.040.040.020.08
00000.500.50000.5000000.5001.5
1965093
0.020.020.020.010.04
species

0.020.020.020.010.04
00001010001000001003
no rank
1385427

694007
species
0.10.020.020.01
00310000100000000001

0.020.040.040.040.040.020.02
0000000.500000.500001000000.5
0.010.020.020.020.020.010.01
694003
species

11128
no rank
00000010000100002000001
0.010.020.020.020.020.010.01

0000000000.30000000.500.40.50.501
0.020.10.020.050.020.020.020.06
no rank
696098
0.010.050.010.020.010.010.010.03

0.010.020.010.020.010.01
0000000000000000101010
1160968
species

species
1541205
00000000000000000000002
0.03

1590370
species
00000000010000000001
0.030.010.01

0000000010000001
0.020.01
species
694008

0.270.160.090.110.360.040.340.090.160.060.040.160.10.020.090.330.110.020.10.4
1.111110.6250.3333330.3636361.444440.52.705880.4444440.750.50.50.37501.500.51.3333301.606060.5454550.51.34.875
0.090.060.030.050.170.020.170.050.070.030.020.060.050.010.030.150.050.010.050.2
1159901
genus

0000000000000000001.5
0.06
no rank
0.03
1232736

species
1159906
0000000000000000003
0.03

0.010.010.010.01
00000100000001010000001
1159904
species

000041612101040000220212
0.10.010.150.020.050.030.020.030.020.020.020.16
species
1159907

1965089
0.030.020.030.010.01
species
0.060.040.060.020.02
000.5000000000.500002000.5000.5

0.030.020.030.010.01
00100000000100004001001
no rank
1159905

species
1159902
01000000000002000040131
0.020.010.050.010.030.01

0.040.04
0.500000000000.5
species
1297661
0.020.02

572289
no rank
100000000001
0.020.02

species
0.070.040.020.020.020.01
1297662
0.140.080.040.040.040.02
2100.50.50000.500000000000.5

no rank
572290
0.070.040.020.020.020.01
42011000100000000001

0001201100100000004
0.020.050.020.020.020.05
species
1159908

genus
0.040.040.060.020.140.160.050.060.020.060.040.040.020.020.130.040.090.020.04
693996
0.10.120.150.060.40.490.130.180.060.160.140.090.070.060.330.10.270.060.1
0.40.3333330.40.3333330.8254.63265000.3076920.3333330.3333330.37500.42857100.6666670.2857140.3333331.212120.30.7407410.3333330.3

1000000000000000002
0.020.02
species
694001

0.01
00000000000001
species
693999

species
0.010.010.040.01
693997
00000000000000.333333000.3333330001.500.5
0.030.030.080.02

no rank
1766554
00000000000000000000301
0.040.01

no rank
12663
0.010.01
00000000000000.5000.5
0.020.02

11135
no rank
0.010.01
00000000000001001

0.030.01
001001
species
693998

277944
species
0.020.020.010.02
0000100000010102

00.50000000.5000000000.5
0.040.040.02
1913642
0.020.020.01
species

766791
no rank
010000001000000001
0.020.020.01

366617
0.020.020.020.050.020.030.020.040.010.010.010.010.070.020.050.010.01
no rank
0.50.500.510.50000.50.50.500.500.50.50.52.214290.50.80.50.5
0.040.040.040.10.040.060.040.080.020.020.020.020.140.040.10.020.02

species
393767
0.020.06
0000000000010000005

1503292
species
010001000101000000001
0.020.010.030.020.01

species
1920748
0.010.01
00000000000000011

100001000010000000012
0.020.010.020.010.03
1503293
species

species
1508224
0.010.010.010.01
00000000000001000011001

1503289
species
0.020.010.01
000100000000000001001

0000200000000000000001
0.050.01
1264898
species

species
28295
0000000000000101002
0.010.010.02

0.060.140.280.040.060.020.020.020.020.02
000.501.58000.50.50000.500000.50.500.50.5
species
1244203
0.030.070.140.020.030.010.010.010.010.01

0.030.070.140.020.030.010.010.010.010.01
001031600110001000011011
no rank
1241933

233263
0.060.010.010.010.010.08
no rank
0.120.020.020.020.020.16
001000.500000000.5000.500000.53

0.01
00000000000000001
864596
species

0.060.010.010.010.08
00200100000001000000016
1892416
species

0.060.060.040.060.060.030.030.02
0.33333300000.666667000.5000.33333300.500.33333300.333333000.5
694013
0.020.020.020.020.020.010.010.01
genus

species
694014
0.020.020.020.020.010.01
0.040.040.040.040.020.02
0.500001000000.500.7500.500.5

000000000000010101
0.010.010.01
11152
no rank

10000200000102
0.020.020.020.01
no rank
11120

species
694015
0.020.01
000000001000000000001

family
76803
0.030.020.030.020.010.01
000.3333330000.333333000.33333300.333333000000.3333330000.333333
0.090.060.090.060.030.03

genus
2003403
0.030.030.020.010.01
000.50000000.500.5000000.50000.5
0.060.060.040.020.02

0010000001010000000001
0.030.030.020.01
species
38143

000000000000000001
0.01
1885929
species

0000000.5
0.04
no rank
325675
0.02

28344
species
0000001
0.02

0.020.030.010.010.020.03
1312872
family
0.080.120.060.040.080.1
0.25000000000.250000.333333000.25000.50.2

0.060.050.07
0.3333330000000000000.40000000.285714
0.020.010.03
1312873
genus

species
0.01
1945563
0.02
00000000000000.5

no rank
1286140
00000000000001
0.01

species
0.020.010.01
1312874
0.040.020.02
0.50000000000000.50000000.5

no rank
1041929
0.020.01
100000000000000000001

no rank
325676
00000000000001
0.01

species
0.01
1945562
000000000000000000000.5
0.02

000000000000000000001
0.01
no rank
1482319

2040576
0.030.010.02
no rank
0.090.030.06
0000000000.3333330000000.333333000.666667

species
2040578
0.03
0.06
0000000000.5

1286141
no rank
0000000001
0.03

00000000000000000.5001
0.020.04
0.010.02
2040577
species

00000000000000001002
0.010.02
1286142
no rank

0.2500.3333330.2666670.3076920.3333330000.333333000.3333330.33333300.6666672.047620.66666700.2857140.1111110.3333331
0.080.090.150.130.030.060.090.240.030.060.210.060.070.090.030.12
0.020.030.050.050.010.020.030.080.010.020.070.020.020.030.010.04
675071
family

00000.5
0.04
genus
0.02
12291

species
100887
0.02
00001

0.020.02
186886
genus
0.040.04
000000000000000101

species
46436
0.02
000000000000000002

species
79918
0000000000000002
0.02

0.060.060.060.040.040.060.160.120.050.060.020.08
0.33333300.50.3333330.500000.5000.50003.5000.40.1666670.51.5
12234
0.020.030.020.020.020.030.080.060.020.030.010.04
genus

species
12239
000000000000000000001
0.01

species
327387
0.02
00001

00000000000000007001
0.060.01
12253
species

species
228578
0.030.010.04
00100000000000000000003

0.04
0000000000001
402399
species

species
12241
0.01
0000000000000000000001

185955
species
0.03
0000000001

species
35281
0000000000001
0.04

327386
0.020.020.01
no rank
0.5000.50000000000000000.5
0.040.040.02

species
1501716
10010000000000000001
0.020.020.01

0.01
000000000
species
51680

29272
species
000000000000000000000
0.010.01

00000000000000.5000.5
0.020.02
genus
0.010.01
119492

0.01
00000000000001
28355
species

0.01
00000000000000001
32629
species

0000.500.5
0.040.02
0.020.01
12326
genus

12327
species
000101
0.020.01

0.02
00000000000000000000000.5
0.01
464865
no rank

species
439014
00000000000000000000001
0.01

000000.2500.33333300.500.416667000000000.3333330.250.25
0.040.060.240.240.030.040.04
family
0.010.020.060.060.010.010.01
11018

genus
0.010.020.060.060.010.010.01
11019
0.030.040.180.180.020.030.03
000000.33333300.500.66666700.555556000000000.50.3333330.333333

species
84589
0.01
0.02
00000000000000000000000.5

0.01
00000000000000000000001
78540
no rank

no rank
0.01
177873
0000000000000000000000.5
0.02

0.01
0000000000000000000001
11021
species

000000010000000000001
0.020.01
species
11020

0.020.120.12
000000.5000100.833333
177874
0.010.060.06
no rank

11039
species
0.04
000000000002

48543
species
0.010.02
000001000001

11024
species
0000000002
0.06

0000000000000000.666667
0.06
family
0.02
11989

0.04
0000000000000001
genus
12008
0.02

0000000000000002
0.02
39804
species

0.20.20.30.4893620.20.5277780.3636360.20.40.2500.20833300.500.2307690.20.70.20.6190480.20.2857140.407407
0.10.10.50.470.10.360.330.10.250.240.480.340.260.20.30.20.210.10.210.54
675063
0.020.020.10.090.020.080.070.020.050.060.10.060.050.030.060.050.050.030.040.12
order

family
0.030.020.030.010.040.030.010.04
675064
000000.60.333333000.3333330000.3333330000010.2857140.3333330.583333
0.10.060.090.030.120.070.030.12

genus
140568
0.020.010.03
0.040.020.06
0000000.500000000000000.501

12433
species
0.01
000000000000000000001

species
12432
0.020.03
00000010000000000000002

genus
0.01
675066
00000000000000.5
0.02

species
213034
0.01
00000000000001

0.070.060.080.020.02
000000.8571430000.50000000001.500.50.5
genus
12176
0.030.030.040.010.01

392174
species
0.01
000001

species
12181
000002
0.02

12194
species
0.03
0000000001

445435
species
00000100000000000003001
0.010.040.01

species
85454
0000000000000000000001
0.01

genus
675076
0.01
0.02
000000000000000000000.5

0.01
000000000000000000001
365150
species

0.02
0000000000000000.5
375548
0.01
no rank

0000000000000001
0.01
species
1502101

000000.6666670000.33333300.3333330100.250.250.250.3333330000.333333
0.060.090.060.120.040.040.040.030.03
family
0.020.030.020.030.010.010.010.010.01
249184

genus
12051
0.010.010.01
00000000000000000.33333300.50000.5
0.030.020.02

327106
0.01
no rank
00000000000000000.5
0.02

0.01
00000000000000001
1955730
species

00000000000000000010001
0.010.01
species
312008

0.040.060.040.090.030.03
0000010000.500.501.3333300.33333300.333333
12148
0.020.030.020.030.010.01
genus

0000000000000200.500.5
0.060.020.02
species
2035014
0.030.010.01

1010616
no rank
0.030.010.01
000000000000040101

species
70822
0.020.030.02
000002000101

0.080.080.40.380.080.120.20.080.20.320.130.150.130.20.120.040.140.27
0.250.250.3750.6052630.250.750.50.250.5000.2500.30769200.2666670.23076910.250.2500.3571430.518519
0.020.020.10.090.020.030.050.020.050.080.030.040.030.050.030.010.030.07
675068
family

0.020.02
00000000000000.500.5
1003890
0.010.01
no rank

species
1978413
0000000000000101
0.010.01

0.33333300.250.250.33333300.6666670.3333330.666667000.33333300.37500.2857140.2500.3333330.33333300.50.7
0.060.120.080.060.150.060.150.120.080.070.080.030.030.080.2
subfamily
0.020.030.020.020.050.020.050.040.020.020.020.010.010.020.07
1914297

0.500.33333300.5010.51000.500.66666700.40.33333300.5000.6666671.07692
0.040.090.040.10.040.10.040.030.050.030.020.060.13
12163
0.020.030.020.050.020.050.020.010.020.010.010.020.07
genus

0.060.020.020.04
000.50000000000000.50.500001
327375
0.030.010.010.02
no rank

001
0.03
species
1569052

1569056
species
00000000000000001
0.01

species
1569053
0000000000000000000002
0.02

0.01
0000000000000001
165250
species

0.01
00000000000001
species
12169

species
67761
0.05
000000002

00000000000000000010003
0.010.04
species
797075

0.01
0000000000000001
species
142843

000000000001
0.02
species
367682

species
12166
0.01
00000000000000000000001

species
31722
00000000000001
0.01

1
0.02
species
12167

138982
species
0.01
00000000000000000000001

00000001
0.02
species
404404

0000102
0.020.05
329164
species

0.060.040.030.030.02
0000.33333300000000.500.333333000.333333000.5
0.020.020.010.010.01
129725
genus

0.02
000000000001
75387
species

00000000000000000001
0.01
35350
species

0.040.020.02
0000.50000000000.5000.5
351425
0.020.010.01
no rank

0.01
00000000000001
351428
species

00010000000000001
0.020.01
1211388
species

0.060.180.210.090.120.150.040.03
00.3333330.666667101000000.333333000001.333330.25000.333333
1914298
0.020.060.070.030.040.050.010.01
subfamily

0.03
1914296
genus
0.06
000001.5

1425362
species
0.03
000003

0000000000000000000.333333
0.03
12174
0.01
genus

0000000000000000000.5
0.02
no rank
0.01
1770617

1868472
species
0000000000000000001
0.01

genus
0.060.070.05
184751
0011.500000000000002
0.120.140.1

0.05
000000000000000004
516956
species

0.060.07
0023
species
1112770

0.04
000000000000.5
genus
0.02
1914293

0.02
000000000001
species
1247116

genus
0.020.020.01
40276
0.040.040.02
00.50000000000.50000000000.5

0.020.01
0000000000010000000001
species
1051792

species
131226
01
0.02

0000000000000000.50.500.5
0.020.020.02
0.010.010.01
675069
no rank

0.010.01
0000000000000000101
species
148879

0.01
0000000000000001
species
129143

119163
0.070.010.020.020.040.010.070.010.020.010.010.04
family
0.210.030.060.080.120.050.210.040.060.030.030.16
0000.57142900.33333300.3333330000.250.3333330.20.3333330.25000.3333330.3333330.33333300.75

0.020.010.020.020.040.010.070.010.010.010.010.04
119164
genus
0000.500.500.50000.3333330.50.250.50.333333000.50.50.501
0.040.020.040.060.080.040.140.030.020.020.020.12

328430
species
0.01
00000000000000000001

0.020.010.010.04
328429
no rank
000000000000.500.33333300.50000001.5
0.040.030.020.08

0.01
0000000000000001
1913024
species

1564903
species
00000000000000000000003
0.04

00000000000000.5
0.02
0.01
1856642
species

0.01
00000000000001
no rank
1833824

000000000001
0.02
1913124
species

312295
species
000001
0.01

0.040.07
000000000000101
species
91753

646010
species
0.01
000000000000000000001

0.020.020.01
0001000100000000001
species
156690

0001
0.1
94699
0.05
no rank

0002
0.05
species
1659298

0000000000000000000.5
0.02
genus
12036
0.01

0000000000000000001
0.01
species
12049

no rank
0.020.030.070.010.010.010.020.020.010.03
38173
0.040.060.140.020.020.020.040.050.030.06
00000.500000.500000.50.50.50.5010.20.3333330.333333

genus
1307798
0.010.010.01
0.020.020.02
000000000000000000000.50.50.5

0.010.01
0000000000000000000011
1457165
species

species
1170422
0.01
00000000000000000000001

363716
species
0.01
00000000000000000000001

111470
species
0.03
0000000001

0000000000000001
0.01
439491
species

340187
species
00001000000000001102
0.020.010.010.02

000000000000001
0.07
340921
species

0.01
000000000000000000000
1560351
species

family
11974
0.020.030.020.030.010.010.04
00.2500000000.2500.250000.36363600.33333300.250.466667
0.080.120.080.110.030.040.15

95339
0.01
genus
000000000000000000.5
0.02

11976
species
000000000000000001
0.01

0000000000000000.5
0.02
0.01
142786
genus

0.01
0000000000000001
11983
species

0.06
0000000000000000.5
genus
0.02
95341

0.04
0000000000000000.75
0.02
95342
species

0000000000000000.5
0.02
no rank
515178
0.01

106333
no rank
0.01
0000000000000001

no rank
179239
0.01
0.02
000000000000000000000.5

0.01
000000000000000000001
1172196
species

95337
0.020.030.020.010.03
genus
0.060.090.060.030.09
00.33333300000000.33333300.33333300000000.3333330.666667

00.500000000.500.500000000.5
0.040.060.040.02
no rank
182093
0.020.030.020.01

0.01
00000000000000000001
species
154122

species
436911
0.02
01

1615583
species
0.030.02
000000000101

35612
0.03
species
000000000000000000001
0.06

146073
no rank
0.03
000000000000000000002

0.520.420.160.090.290.280.250.20.350.330.190.280.40.990.971.30.61.310.480.580.530.610.63
35301
no rank
2.812.110.820.451.451.651.160.941.861.771.11.462.045.069999999999995.086.759999999999993.199999999999996.589999999999992.362.772.729999999999993.163.35999999999999
0.3807830.2085310.1829270.1777780.2344830.2848480.3362070.2340430.1505380.2542370.1818180.1917810.4313739.536490.38582716.07695.0031312.25190.296610.4223831.047620.5854430.58631

0.50.50000.50.500000.500.500.5000.83333310.50.50.4
0.040.040.020.040.040.020.020.060.040.020.020.05
988644
0.020.020.010.020.020.010.010.030.020.010.010.03
no rank

0.02
00000000000000000002
1608133
species

1849544
species
00000000000001
0.01

species
1608097
1100011000010000001001
0.020.020.010.020.020.010.01

0.01
0000000000000001
1608123
species

1608107
species
0.01
00000000000000000000001

species
1608126
0.01
000000000000000000001

species
1629671
0.01
00000000000000000000001

0000000000000000002
0.02
1608139
species

0000000000.2500.3333330.25000.20.20.25000.20.5
0.120.060.160.050.050.080.10.16
0.030.020.040.010.010.020.030.04
11308
family

0000000000000000.250.250.25000.250.5
0.040.040.040.040.08
genus
0.010.010.010.010.02
197911

0.010.010.010.010.02
11320
species
0000000000000000.3333330.3333330.333333000.3333330.666667
0.030.030.030.030.06

no rank
114729
0.010.02
0.020.04
0000000000000000.5000001

0.010.02
0000000000000001000002
488241
no rank

00000000000000000.50000.5
0.020.02
0.010.01
102796
no rank

130760
no rank
000000000000000010001
0.010.01

0.01
333278
no rank
0.02
000000000000000000.5

1332244
no rank
0.01
000000000000000001

0.040.020.04
000000000000.5000000.50001
324913
0.020.010.02
genus

0000000000010000010002
0.020.010.02
55987
species

genus
1299308
0.030.04
0.090.12
0000000000.333333000.333333

0000000000.5000.5
0.060.08
no rank
1566308
0.030.04

0.030.04
0000000001001
1566309
species

197913
0.01
genus
000000000000000000000.333333
0.03

0.02
000000000000000000000.5
0.01
11552
species

no rank
11553
000000000000000000001
0.01

0.930.570.30.630.350.180.10.570.540.180.360.322.451.264.541.884.770.520.971.141.011.46
0.5913980.2807020.200.3809520.2857140.2222220.20.1754390.3333330.2222220.2222220.2516.08160.33333323.80628.3670216.88470.3269230.3195882.078951.198020.69863
order
1980410
0.190.130.060.140.070.050.020.120.120.040.080.080.60.281.10.441.180.130.240.270.230.29

000.200.250.25000.22222200000.33333300.2380950.250.9565220.250.2857144.333332.666670.538462
0.150.080.080.180.060.210.080.230.080.070.510.270.13
1980418
0.030.020.020.050.010.050.020.070.020.020.170.090.04
family

000.2500.3333330.333333000.30769200000.400.31250.3333331.3750.3333330.333333000.333333
0.120.060.060.130.050.160.060.160.060.030.03
11584
0.030.020.020.050.010.050.020.070.020.010.01
genus

000000000000000000.5
0.02
0.01
1933186
species

no rank
11587
0.01
000000000000000001

0.02
00000000000000000000.5
species
1933188
0.01

0.01
00000000000000000001
no rank
904699

00000000000000000000000.5
0.02
1933190
0.01
species

0.01
00000000000000000000001
no rank
992212

0.02
00000000000000000.5
0.01
1933187
species

0.01
00000000000000001
11588
no rank

0.090.040.040.040.040.110.020.020.02
000.33333300.50.5000.500000.500.4545450.50.50.5
no rank
0.030.020.020.020.010.050.010.010.01
327794

1926502
species
0.010.01
00000000000000011

904698
species
0.01
0000000000000001

0.020.01
000011
species
1926501

0.01
0000000000000000001
species
1926500

1608048
species
000001
0.01

0.020.01
0000000010000001
1055750
species

000.500000000000.500.5
0.060.020.02
1405806
0.030.010.01
no rank

species
1405807
0.030.010.01
0010000000000101

0.010.010.01
000000000000010101
species
904722

0.040.02
000000000.50000000000.5
0.020.01
1933191
species

0000000010000000001
0.020.01
no rank
11591

0.05
000000000000000004
species
1933183

00000000000000000000.56.541
0.020.340.180.06
0.010.170.090.03
12329
genus

000000000000000000011382
0.010.170.090.03
species
66266

0.060.060.210.030.140.03
0.3333330.3333330000000000000.3333330000.3333330.571429000.333333
0.020.020.070.010.050.01
1980412
family

genus
0.020.020.070.010.050.01
675845
0.50.50000000000000.50000.50.888889000.5
0.040.040.140.020.090.02

species
1980431
10000000000000100002001
0.020.070.020.01

0.02
00000000000000000002
1980426
species

0100000000000000001
0.020.01
species
1980427

0.060.060.060.060.120.030.060.05
0.3333330.333333000.333333000000.3333330.333333000000.3333330.333333000.2
0.020.020.020.020.040.010.020.01
1980413
family

1980442
0.020.020.020.020.040.010.020.01
genus
0.50.5000.5000000.50.5000000.50.5000.25
0.040.040.040.040.080.020.040.04

0.020.02
000010000001
species
1980493

1980471
species
0.020.020.020.01
1100000000010000001

species
1980491
0.020.01
0000000000100000001

1980490
species
000000000000000001
0.01

0000000000000000000000.333333
0.03
no rank
0.01
1986491

no rank
339351
0.01
0000000000000000000000.5
0.02

0.01
0000000000000000000001
species
93830

0000000000000000.3333330010.4166670.6666670.25
0.030.090.120.090.04
1980415
0.010.030.040.030.01
family

genus
1980517
0.010.030.040.030.01
0.020.060.080.060.03
0000000000000000.5001.50.62510.333333

0.020.02
00000000000000000000.500.5
1340802
0.010.01
no rank

1615758
species
00000000000000000001
0.01

species
1381104
0.01
0000000000000000000001

0.01
0000000000000001
1980521
species

000000000000000000322
0.030.020.03
1980529
species

1.021740.384615000.50.46153800.250.21052600.250.2500.3750.500.250.3043480.2222220.3043480.3750.2352941.03409
0.460.260.20.130.080.190.080.080.080.560.080.230.090.230.240.170.88
1980419
0.110.060.050.030.020.050.020.020.020.140.020.060.020.060.060.040.2
family

0.350.20.150.10.060.140.060.060.060.420.060.170.070.170.180.130.68
1.342860.5000.6666670.600.3333330.28571400.3333330.33333300.50.66666700.3333330.4117650.2857140.4117650.50.3076921.33824
11611
0.110.060.050.030.020.050.020.020.020.140.020.060.020.060.060.040.2
genus

species
1933301
0.02
00000000.5
0.04

00000001
0.02
83872
no rank

00000000000000000.5
0.02
species
0.01
1933294

no rank
11612
00000000000000001
0.01

0.01
430606
species
0.02
0000000000000000000.5

0000000000000000001
0.01
no rank
1678786

0.020.02
000000000000000000000.50.5
0.010.01
1933295
species

0.010.01
0000000000000000000011
60456
no rank

no rank
326176
0.110.060.050.030.050.020.020.020.140.010.060.010.060.050.030.2
0.240.140.10.070.090.040.040.040.280.020.110.030.110.10.070.48
1.958330.7142860010.857143000.44444400.50.500.75100.50.6363640.3333330.6363640.80.4285711.89583

0.010.140.010.01
000001000000002001001
89471
species

species
460926
0.01
00000000000000000000001

0000000000000000000111
0.010.010.01
485724
species

00000000000000000100012
0.010.010.03
species
1033976

83871
species
0.050.020.020.020.030.01
00002200000000000202201

species
163325
00000000100000000101
0.020.010.01

0.090.040.020.01
52000000001002
species
1414655

0.50.500000000000000000.5000.53.5
0.040.040.020.020.18
0.020.020.010.010.09
1027231
no rank

species
1027232
11000000000000000010017
0.020.020.010.010.09

00000100100101001001004
0.010.020.020.010.010.010.05
species
1857323

1980416
0.040.020.030.050.020.050.020.120.020.080.560.071.050.411.030.010.060.010.080.04
family
0.250.3333330.33333300.6666670.2857140.30769200.250.42857100.250.33333323.01170.33333333.756312.257825.88390.250.5294120.3333331.720.230769
0.160.060.090.150.070.130.080.420.080.241.710.213.21.283.10.040.170.030.250.13

0.040.060.10.020.040.120.080.060.060.020.070.020.14
00.50.5010.50.5001000.5200.83333300.500.7142860.53
no rank
0.020.030.050.010.020.060.040.030.030.010.040.010.07
39718

species
1608060
0.01
00000000000000000001

000020000000100000001
0.050.040.01
species
1590836

0.030.010.06
0010010002
species
1608101

1608068
species
000000100000040101
0.020.030.010.01

0000000000000002
0.02
1608047
species

00000000000000000002
0.02
1457386
species

1266451
species
0100000000000000000006
0.020.07

11572
0.040.010.020.020.060.020.040.530.071.020.411.020.020.010.04
genus
0.120.030.040.060.180.060.081.060.142.090.872.050.040.030.09
0.33333300000.3333330.500.3333330.33333300.3333330.5370.551.660318.034539.13660100.3333330.333333

0.5000000000.5000000.50000000.5
0.040.060.020.02
species
1933304
0.020.030.010.01

no rank
1495866
10000000010000010000001
0.020.030.010.01

0.020.020.01
348018
no rank
00000000000000010100000.5
0.040.040.02

0.020.020.01
00000000000000020200001
1406134
species

species
0.01
1933179
0000000000000000.5
0.02

0.01
0000000000000001
no rank
35304

159150
species
0.530.990.360.990.01
0000000000000740109438100001

273350
species
0.020.040.02
00000010000010000002

000000000000001
0.07
species
35306

0.5
0.04
species
0.02
1933178

no rank
70566
0.02
1

0.020.040.060.040.020.02
000000.5000.50.500.500000.500000.5
species
0.010.020.030.020.010.01
159148

no rank
11582
0000010011010000100001
0.010.020.030.020.010.01

0.04
1933309
species
0.08
00000000000000002.5

00000000000000005
0.04
118655
no rank

0.010.01
1980533
genus
00000000000000.33333300000.333333
0.030.03

species
0.010.01
1980536
0.020.02
00000000000000.500000.5

1608075
no rank
0.010.01
0000000000000100001

0.060.020.030.030.070.020.050.020.030.070.010.010.050.070.08
11617
family
0.180.060.120.090.250.060.140.080.10.210.030.030.150.20.25
0.5555560.3333330.250010.40.3333330.28571400.25000.40.33333300.33333300.3333330.26666700.70.56

1653395
0.030.020.010.01
genus
000.33333300000000.333333000000000.333333000.333333
0.090.060.030.03

0.030.010.01
1653398
species
000.500000000000000000.5000.5
0.060.020.02

1303019
no rank
0.030.010.01
00100000000000000001001

0.04
00000000000.5
1653399
0.02
species

00000000001
0.02
no rank
1223561

no rank
0.01
208898
0000000
0.05

0.01
11618
genus
0.04
0000000

0.01
208896
no rank
0000000
0.03

0.01
483045
no rank
0000000
0.02

300180
species
0.01
0000000

genus
1653394
0.060.020.030.060.020.050.030.070.010.010.040.070.07
0.120.040.060.130.040.090.070.140.020.020.070.130.14
0.8333330.50001.50.7692310.50.44444400000.5714290.500.500.50.42857101.076920.928571

45219
species
000000000000001
0.07

00000000000000001
0.01
species
144752

0.01
11629
species
0.02
0000000

no rank
300175
0.01
0000000

1642458
species
1
0.02

00000000000000000000001
0.01
species
42764

0.030.020.01
00000300100001
45221
species

species
11620
0.03
00000000000000000000002

0000000110000000000102
0.020.020.010.02
45222
species

0.020.010.03
01000000000000000000012
species
45709

0.01
00000000000001
45220
species

00000000000001
0.01
11619
species

0000002000000100001003
0.050.010.010.03
species
11623

species
192848
2
0.04

45223
species
00000000000000000001
0.01

00000000000000000001
0.01
11630
species

0.020.020.010.020.02
568247
family
0.060.060.030.060.06
00.333333000.333333000000000.333333000.666667000.666667

0.040.040.020.040.04
00.5000.5000000000.5001001
genus
0.020.020.010.020.02
88129

01001000000001002002
0.020.020.010.020.02
73561
species

1.140.960.240.360.470.9100000000000010.440.580.80.780.650.721.161.482.640.850.580.431.270.970.941.160.97
0.3508770.2291670.250.2222220.1702130.2967030.5227270.3103450.1750.3076920.2153850.2222220.6551726.006760.5568180.6823530.379310.4186050.3700790.7628870.4893620.3534480.814433
0.260.230.060.090.120.180.10.150.190.180.130.160.280.350.620.180.120.110.310.250.220.260.23
11157
order

000000.50000000000000.5000.25
0.080.080.04
family
0.020.020.01
1884452

genus
1921703
0.020.020.01
0.060.060.03
000000.6666670000000000000.666667000.333333

0.040.040.02
0000010000000000001000.5
1884453
0.020.020.01
species

no rank
1195087
0000020000000000002001
0.020.020.01

family
11270
0.170.150.030.070.050.090.070.020.090.150.10.080.280.320.480.140.080.050.080.070.050.130.11
0.379310.2978720.3333330.2857140.1333330.3529410.750.250.1818180.4117650.2631580.2857140.8636368.676470.8312510.5862070.8666670.5416670.9565220.60.43750.7
0.580.470.090.210.150.340.280.080.330.510.380.280.881.021.60.520.290.150.240.230.20.480.4

genus
1978532
0.020.010.010.010.01
0.060.030.030.030.03
00000000000.33333300000.3333330.333333000.33333300.333333

0.020.01
1978541
species
0.040.02
00000000000.5000000000.5

1608119
no rank
00000000001000000001
0.020.01

species
1978539
0.01
00000000000000000.5
0.02

no rank
1608146
00000000000000001
0.01

species
0.01
1978545
0.02
0000000000000000.5

0000000000000001
0.01
no rank
1408144

species
0.01
1978543
0000000000000000000000.5
0.02

0.01
0000000000000000000001
no rank
1537975

0.02
186778
genus
000000000.333333
0.06

species
11287
0.02
0.04
000000000.5

no rank
1147160
000000001
0.02

0.020.020.070.020.070.03
1299306
genus
0.060.060.210.060.210.09
0.3333330.333333000010000.3333330000.33333300000000.666667

0.50.500001.5
0.040.040.14
0.020.020.07
1972586
species

764599
no rank
1100003
0.020.020.07

0.040.140.06
00000000000.50000.500000001
species
0.020.070.03
1972584

0.020.070.03
00000000001000100000002
318844
no rank

genus
32613
0.020.020.020.020.030.020.020.040.010.070.030.020.010.070.05
0.060.060.060.060.090.060.060.120.030.210.090.060.030.210.15
0.3333330.33333300.33333300.55555600000.33333300.3333330.3333330.3333330.3333330.66666700.333333000.6190481.33333

species
1972592
0.020.020.010.030.05
0.040.040.020.060.1
00.500010000000000000.5001.52

01000200000000000010034
0.020.020.010.030.05
380160
no rank

1972596
0.020.02
species
0.5000.5
0.040.04

no rank
1550518
1001
0.020.02

species
0.010.010.01
1972595
0.020.020.02
0000000000000000000000

0000000000000000000000
0.010.010.01
no rank
318835

1972587
0.02
species
00000000000.5
0.04

no rank
318836
00000000001
0.02

0.040.070.010.02
1972593
species
0000000000000.500.50.5000001
0.080.140.020.04

0000000000001011000002
0.040.070.010.02
no rank
11303

0.010.010.02
1972594
species
0.020.020.04
00000000000000.500.51

00000000000001012
0.010.010.02
318834
no rank

1972589
0.01
species
0.02
0000000000000000.5

0.01
0000000000000001
no rank
31612

0000000000000000000000
0.020.020.02
349172
0.010.010.01
no rank

0000000000000000000000
0.010.010.01
1229186
species

000000.5
0.02
species
1972597
0.01

000001
0.01
1272960
no rank

0.020.020.020.030.02
1972682
genus
0.060.060.060.090.06
0.3333330000000.3333330.3333330.33333300.333333

species
1972686
0.020.020.020.030.02
0.040.040.040.060.04
0.50000000.50.50.500.5

100000011101
0.020.020.020.030.02
no rank
1479613

00000000000000.333333000000000.333333
0.030.03
0.010.01
1308858
genus

species
1094373
0.01
0.02
00000000000000000000000.5

00000000000000000000001
0.01
666363
no rank

0.02
00000000000000.5
948741
0.01
species

no rank
666962
00000000000001
0.01

0.150.160.020.120.040.160.560.040.070.020.060.10.060.04
0.80.50000.5000100.50.515.696400.51.428570.51.5211
0.070.080.010.060.020.080.280.020.030.010.030.050.030.02
35303
no rank

0.02
000000000001
species
1511639

1608112
species
0.020.25
100000000000035

0.060.010.030.02
0000000002000100003002
1608104
species

1620892
species
0.01
00000000000002

1272959
species
0.01
000000000000000001

1459044
species
0.01
0000000000000001

1608138
species
0.02
01

00000000000010000004
0.040.05
species
1608100

1911435
species
00000000000000001
0.01

2
0.04
species
1608114

species
1569258
0.03
00000000000000003

species
1608067
0000000000000101
0.010.01

000001
0.01
species
1288359

species
1911434
0.02
01

0100000000001
0.020.04
species
936149

species
1479612
1
0.02

1608102
species
000000000000000000002
0.03

species
1288358
0.02
01

000000.33333300000000000000.3333330.666667
0.030.030.09
genus
0.010.010.03
11305

000000.5000000000000001
0.020.06
species
1972639
0.010.03

998864
no rank
0.010.03
000001000000000000002

species
1985702
0.01
00000000000000000000.5
0.02

00000000000000000001
0.01
no rank
32612

0.01
1913605
genus
0000000000000000000.5
0.02

0000000000000000001
0.01
152177
species

0000.50.50.500000000.666667
0.040.040.040.03
genus
1972611
0.020.020.020.01

0.010.01
00000100000001
species
1972623

0.020.020.01
000111
1972626
species

00000000000001
0.01
1972612
species

000.5000.5000000.50000000000.50.5
0.060.020.040.020.02
11306
0.030.010.020.010.01
genus

209854
species
0.01
0000000000000000000001

195059
species
001001
0.030.01

00000000000000000000001
0.01
488317
species

species
195060
0.02
000000000001

11286
0.040.020.010.030.160.350.090.010.040.01
genus
0.080.040.020.060.320.70.220.020.080.02
0.50.50000.50000.500201.72.090910.51.50000.5

no rank
0.04
327825
0.08
0000000000000002

642022
species
0000000000000004
0.04

0.030.160.280.050.04
000000000100404603
1072176
species

1
0.02
species
211977

species
57482
1
0.02

01
0.02
species
12538

species
249583
00000000000000101
0.070.01

0.01
000001
species
1167696

species
746543
0.01
0000000000000000000001

1985692
0.010.01
genus
00000000000000000.33333300000.333333
0.030.03

0.020.02
00000000000000000.500000.5
species
1985693
0.010.01

0000000000000000100001
0.010.01
no rank
318843

0.020.010.020.030.040.020.010.02
11271
genus
0000.500.333333000.3333330.3333330.40.33333300.400000.4
0.040.030.060.090.10.060.050.05

species
1972568
00000000001
0.02

0.01
1972573
species
00000000000000.5
0.02

no rank
290008
0.01
00000000000001

species
0.01
1972579
0.02
00000000000000.5

00000000000001
0.01
no rank
198833

1972578
0.01
species
0000000000000000000.5
0.02

0.01
0000000000000000001
no rank
11280

000000.5000.50.50.50.5
0.020.040.060.040.04
species
1972564
0.010.020.030.020.02

1344113
no rank
0.010.020.030.020.02
000001001111

1972569
species
0001000000000000001
0.020.01

1513294
0.020.020.01
family
00000.3333330000.33333300000.333333
0.060.060.03

0.040.04
00000.50000.5
0.020.02
1884454
genus

0.020.02
000010001
1034377
species

genus
0.01
1513295
0.02
00000000000000.5

species
644609
00000000000001
0.01

0.120.060.030.030.060.06
0.6666670000000000.333333000000.3333330.3333330.6666670.666667
0.040.020.010.010.020.02
11266
family

genus
0.010.02
186537
0.020.04
00000000000000000.501

species
11269
0.010.02
0000000000000000102

10000000000.50000000.501
0.080.040.020.04
186536
0.040.020.010.02
genus

186540
species
0.040.02
20000000000000000002

0.02
00000000001
species
186541

000000000000000001
0.01
186538
species

family
1883067
0.02
000000000000.25
0.08

000000000000.333333
0.06
genus
1883068
0.02

0.04
000000000000.5
species
1883069
0.02

000000000001
0.02
1483724
no rank

0.080.040.060.020.030.030.03
00000.250.25000000.3333330000000.500.3333330.3333330.333333
no rank
0.020.010.020.010.010.010.01
644608

species
1435457
0.01
0000000000000000001

1884455
0.02
genus
0.04
000000000000.5

1884460
species
0.02
000000000001

genus
0.010.010.01
1884456
000000000000000000000.50.50.5
0.020.020.02

1884461
species
00000000000000000000111
0.010.010.01

0.060.03
00000.3333330.333333
genus
1884459
0.020.01

species
0.020.01
2040572
0.040.02
00000.50.5

1608063
no rank
000011
0.020.01

family
0.020.080.020.020.070.020.10.050.030.020.040.020.140.040.020.040.150.150.120.110.11
11158
0.060.260.060.060.270.060.290.160.090.080.140.080.420.150.080.10.50.430.370.350.31
0.3333330.30769200.3333330.3333330.370370.3333330.4827590.250.3333330.250.28571400.3750.3333330.40.250.30.421.116280.7027030.5142861.6129

genus
1283308
0.02
0.04
000000000000.5

000000000001
0.02
species
122203

00.50000.50.500000000.5000.50.5000.50.5
0.040.040.040.140.040.020.020.02
genus
0.020.020.020.070.020.010.010.01
186938

11215
species
010000000000000001
0.020.01

0000011000000010000001
0.010.020.070.01
species
12730

species
1357321
00000000000000000110001
0.010.010.01

11216
species
000001
0.01

00.50000.66666700000.3333330.333333000.50.6666670.33333300.7692310.50.3333330.833333
0.040.060.060.060.140.060.030.130.020.060.12
genus
0.020.020.020.020.070.020.010.050.010.030.04
39744

species
0.020.020.020.010.01
1979167
00000000000.50.50001000.5000.5
0.040.040.040.020.02

1452514
no rank
0000000000110002001001
0.020.020.020.010.01

00000000000000000001
0.01
1979163
species

1979162
species
0.02
01

0.02
00000000000000000.5
species
0.01
1979161

0.01
00000000000000001
11224
no rank

species
1979160
000000000000001
0.07

000000000000000000001
0.01
1979165
species

0000010000000000001.500.51.5
0.040.060.020.06
species
0.020.030.010.03
1979158

no rank
1261101
0000020000000000003013
0.020.030.010.03

000000.500000000.50000.50.5000.5
0.020.020.020.020.02
260963
0.010.010.010.010.01
genus

1305716
species
00000000000001
0.01

species
623144
0000000000000000001001
0.010.01

000001000000000001
0.010.01
622416
species

0.060.110.020.030.12
000000.333333000.36363600000.500.333333000.5
260964
0.020.050.010.010.05
genus

0.020.040.04
000000.5000.50000000001
1979179
0.010.020.02
species

no rank
1221391
0000010010000000002
0.010.020.02

0.020.010.01
0000000010000100001
63330
species

species
0.010.010.01
1979176
0.020.020.02
000000.50000000000.5000.5

1474807
no rank
0.010.010.01
0000010000000001001

genus
11229
0.020.040.050.010.010.020.040.01
0.040.10.10.020.030.040.070.02
0.50.400000100000000.50.33333300.50.42857100.5

1170234
species
0.020.050.010.010.01
10000002000000010011

00.5000000000000000.5
0.040.02
0.020.01
11241
species

11242
no rank
0.020.01
01000000000000001

species
11240
0.02
01

species
31604
0.010.01
00000000000000000011

00000000000000000001
0.01
species
11234

0000000000000000000001
0.01
11232
species

0.020.020.010.050.030.010.010.10.090.030.09
336871
no rank
0000.50.50.500.44444400.50000.500000.52.315791.263160.8333332.72222
0.040.040.020.090.060.020.020.190.190.060.18

92129
species
00000100000001000000021
0.010.010.020.01

00011001010000000016306
0.020.020.020.030.010.070.040.08
species
241630

341053
species
0.020.01
00000001000000000001

00000000000000000001
0.01
590647
species

0.03
000000000000000000002
120499
species

0.030.01
0000000000000000000021
322067
species

0.120.090.060.060.060.040.060.12
0.66666700.33333300000.3333330.33333300000000.3333330.250.66666700.583333
family
0.040.030.020.020.020.010.020.04
11244

genus
162387
0.040.020.01
10000000000000000.50000.5
0.080.040.02

00000000000000001
0.01
species
38525

162145
species
0.040.010.01
200000000000000010001

000.500000.50.5000000000.333333101
0.060.040.040.030.040.06
0.030.020.020.010.020.03
1868215
genus

0.02
000000000000000000.5
species
0.01
11263

000000000000000001
0.01
270473
no rank

11250
species
0.020.03
000000000000000000202

species
11246
0.030.020.02
001000011

000.33333300000000000.66666700.333333000.50000.666667
0.090.030.030.020.09
no rank
0.030.010.010.010.03
439490

species
1516128
0000000000000000001
0.01

1661063
0.030.010.010.03
family
0.060.020.020.06
000.50000000000100.50000001

0010000000000201
0.030.010.01
species
1849537

0.03
00000000000000000000002
1532180
species

0.080.160.070.080.08
00000.250000000.2500.28571400.250000000.25
family
0.020.040.010.020.03
1910928

1985368
0.020.020.010.010.01
genus
0.060.060.030.030.03
00000.3333330000000.33333300.33333300.3333330000000.333333

0.040.040.020.020.02
00000.50000000.500.500.50000000.5
1985420
0.020.020.010.010.01
species

no rank
1391026
0.020.020.010.010.01
00001000000101010000001

1542744
0.020.01
genus
000000000000.33333300.333333
0.060.03

000000000000.5
0.04
1985393
0.02
species

000000000001
0.02
no rank
1843739

0.01
1910929
no rank
00000000000000.5
0.02

0.01
00000000000001
1930298
species

0.03
0000000000000000.333333
genus
1941236
0.01

2004963
0.01
species
0.02
0000000000000000.5

no rank
1634486
0.01
0000000000000001

no rank
0.01
1941235
0.02
00000000000000000000000.5

species
1941233
00000000000000000000001
0.01

no rank
35268
0.090.040.050.290.10.070.050.090.060.080.120.240.250.210.430.160.220.170.220.340.140.13
0.450.180.231.260.540.320.210.360.240.380.521.041.11.121.920.750.940.780.941.420.60.6
0.3111110.22222200.1739130.3571430.370370.18750.1904760.2222220.250.2105260.2307690.4615382.709090.31254.453121.506670.8297870.5897440.6702132.225350.3166670.233333

186534
0.020.040.020.190.080.050.020.090.060.060.10.20.060.070.030.020.070.10.190.140.10.11
family
0.3333330.28571400.250.5081970.5862070.2666670.3333330.2962960.3333330.30.31250.7333330.3809520.3333330.4545450.2857140.3809521.096771.016950.6078430.406250.342857
0.060.140.080.610.290.150.060.270.180.20.320.60.210.210.110.070.210.310.590.510.320.35

0.010.030.010.010.01
10639
genus
000000.3333330000.50000.500000.3333330000.5
0.030.060.020.030.02

515319
0.010.01
no rank
0.020.02
000000.50000000000000.5

0000000000000000001
0.01
1221449
species

515444
species
0.01
000001

0000000001
0.03
species
194445

10649
species
0.01
00000000000000000000001

0.01
00000000000001
species
515320

genus
186533
0.020.020.010.050.080.010.04
0.060.040.020.140.20.030.11
00000.33333300.5000000000000.50.9285711.10.3333330.636364

no rank
930167
0.020.040.050.010.03
00000.5000000000000001.520.51
0.040.080.10.020.06

0.020.040.050.010.03
00001000000000000003412
species
930168

species
107324
0.020.010.01
000000100000000000011

0.010.010.01
00000000000000000010101
species
38062

0.020.030.040.020.01
187212
genus
0.040.060.080.040.02
000000000.50.5000.5000010.5

0000000011001000021
0.020.030.040.020.01
species
10654

0.50000.3636361.500000.50.51.50.50.50.500.52.500.40.40.4
0.040.110.060.040.040.240.020.140.020.020.120.050.050.05
genus
0.020.050.030.020.020.120.010.070.010.010.060.030.020.03
187214

0.020.010.010.01
00000000000100010000101
35593
species

172220
species
0.020.020.030.020.120.010.070.010.060.01
10001300001031100150001

0.010.01
0000000000000000000011
10651
species

00000.500000000000000000.5
0.040.02
no rank
418399
0.020.01

species
1848040
0.020.01
0000100000000000000001

0.020.020.070.02
446548
no rank
000000000.5000.50000000301
0.040.040.140.04

587370
species
0000000010010000000602
0.020.020.070.02

0.10.060.250.120.060.040.060.10.140.080.110.060.050.080.10.120.10.06
00.400.33333310.5833330.3333330.50.33333300.40.4285710.50.54545500.6666670.40.37500.40.5833330.60.333333
10652
0.040.020.120.040.020.020.020.040.060.040.040.020.020.040.050.040.040.03
genus

328670
species
0.020.01
00000001000000000001

0.01
00000000000000000001
1471299
species

no rank
325348
0.020.020.020.030.020.020.020.020.030.020.010.010.010.040.020.01
00.500.50.50.8571430.500.500.50.500.8010.50.500.50.8750.50.5
0.040.040.040.070.040.040.040.040.050.040.020.020.020.080.040.02

549205
species
0.02
00000000001

000001
0.01
species
1634914

00000000000000000100001
0.010.01
species
52996

000000000000000000002
0.03
1459800
species

646442
species
0.01
000001

species
1016856
0.020.01
01000000000000000001

000100000000000000001
0.020.01
1940252
species

647294
species
00000010000002001
0.020.010.01

1561150
species
0.01
0000000000000000000001

00000000000002
0.01
species
1016855

0000020010010000000001
0.020.020.020.01
species
1227557

1016852
species
0.020.02
0000100000000002

0.010.01
000000000000000011
species
362398

species
157270
0.010.01
000000000000010001

0.01
000001
species
328671

species
10653
00000000000001000000001
0.010.01

1016854
species
000010000011
0.020.020.02

0.07
00003
1016853
species

0000000000001
0.04
species
178354

0.020.020.010.02
0100000000010000000102
species
31559

0.020.010.01
1232640
genus
0.040.020.02
000000000.50000000000.50.5

00000000100000000011
0.020.010.01
species
995049

family
10404
0.020.02
0.3333330000.333333
0.060.06

0.020.02
336063
no rank
0.040.04
0.50000.5

species
259931
0.020.02
10001

family
11632
0.060.020.070.030.020.020.020.020.040.190.140.40.140.150.070.040.190.040.03
0.240.10.30.150.10.10.10.080.20.640.71.380.520.510.30.130.570.140.12
0.416667000.20.40.20.20.2000.20.250.24.531250.46.159422.134621.372550.40.23076950.4285710.166667

0.020.030.020.020.020.020.040.010.090.020.010.020.010.010.03
327045
subfamily
0.080.120.080.080.080.060.160.040.320.080.040.090.040.040.09
0000.2500.250.250.25000.250.3333330.250.5010.250.250.2222220.2500.250.222222

genus
0.020.010.020.040.030.010.01
11646
0.060.030.060.120.070.030.03
0000.33333300.3333330.333333000000.333333000.285714000.333333000.333333

11653
0.010.01
no rank
0.020.02
000000.50000000000000.5

0.010.01
0000010000000000001
species
11741

0
0000000000000000
0
11655
no rank

11673
species
0
0000000000000000

0.08
0000000000000.5
subgenus
11654
0.04

0.04
0000000000001
11665
species

0000000000000000000000.5
0.02
no rank
0.01
348386

0.01
0000000000000000000001
11663
species

0000.5000.5000000000.5
0.040.040.04
no rank
0.020.020.02
11652

0.020.020.01
0001001000000001
11676
species

11709
species
0
0000000000000000

0.01
0000000000000001
11723
species

0.030.060.060.040.030.030.06
000000.33333300.333333000.3333330.500000.333333000.333333000.333333
0.010.020.020.020.010.010.03
140052
genus

000000.500.5000.5000000.5000.5000.5
0.020.040.040.020.020.02
0.010.020.020.010.010.01
347957
no rank

00000000001
0.02
species
1904439

1822365
species
000001
0.01

0.01
00000000000000001
species
723802

239365
species
0.010.01
00000000000000000001001

00000001
0.02
69576
species

0.02
000000000001
11855
species

species
11757
00000000000000000000001
0.01

00000000000000.666667020.3333330.3333330.25
0.030.150.030.030.04
153135
0.010.050.010.010.01
genus

no rank
353769
0.010.050.010.010.01
0.020.10.020.020.03
00000000000001030.50.50.333333

000000000000020611
0.010.050.010.01
11788
species

0000000000000000000.5
0.02
196403
0.01
species

0000000000000000001
0.01
11764
no rank

0000000000000000
0.01
genus
153057
0.01

11884
species
0000000000000000
0

species
11886
0
0000000000000000

0.03
000000.333333
genus
0.01
153136

194440
0.01
species
000000.5
0.02

0.01
000001
11908
no rank

0.020.170.140.290.10.10.010.190.03
35276
no rank
00000.25000000007.552630.513.56674.76193.0476200.57.50.833333
0.080.380.560.60.210.210.020.380.06

00000000000000000001152
0.010.190.02
1435008
species

0.020.010.14
206037
no rank
00000.333333000000000.3333330.666667
0.060.030.42

0.020.010.14
45617
species
0.040.020.28
00000.5000000000.51

166122
no rank
0.020.010.14
000010000000012

348296
species
0.140.270.090.090.01
0000000000000200301170001

206039
0.020.020.010.01
no rank
00000000000001.5010.50.5
0.040.040.020.02

000000000000030211
0.020.020.010.01
204873
species

subfamily
0.060.050.010.020.030.040.050.01
327046
0.180.150.030.060.090.110.140.03
0.5555560000.666667000000000.33333300.66666710.4545450.7142860.333333

0.060.050.010.020.030.040.050.01
11640
genus
0.8333330001000000000.5011.50.7142861.111110.5
0.120.10.020.040.060.070.090.02

species
207343
1
0.02

20000000000001023201
0.040.010.020.030.020.01
53182
species

109270
species
000000000000000001
0.01

0.03
0000000000000000003
species
338079

species
11642
0.050.01
0000200000000000001

no rank
552364
0.030.070.030.050.040.020.040.030.030.050.020.040.010.15
0.060.140.060.090.080.040.080.060.060.10.040.080.020.3
000.500.8571430.83333300.444444000.50.500.62500.8333330.8333331.300.50.8750.55.5

001022010000020221
0.030.050.020.020.010.020.020.01
1557034
species

1335638
species
000000000010010000002
0.020.010.03

000001010011010100011111
0.010.020.020.020.010.010.010.010.010.15
species
1557035

00001000000001001301
0.020.010.010.040.01
1557033
species

0.2385320.2588240.3648650.1818180.650.2857140.3761470.2240.2535210.1463410.259740.1754390.3752.489540.252.765771.137932.022220.2673270.3157890.3885350.3035710.333333
1.090.850.740.551.60.771.091.250.710.410.770.570.962.390.283.331.161.81.010.571.570.560.51
0.210.190.160.120.310.150.250.270.140.070.150.110.20.460.070.610.210.340.20.110.310.110.12
35325
no rank

00000.3333330.2500000.333333000000000.333333
0.060.040.060.03
39748
0.020.010.020.01
family

00000.5000000.5000000000.5
0.040.040.02
no rank
0.020.020.01
1923830

0.01
00000000000000000001
1926644
species

0.020.02
00001000001
1922393
species

0.01
39749
genus
000000.333333
0.03

358104
0.01
no rank
000000.5
0.02

1129873
species
000001
0.01

249310
0.020.010.02
family
0.060.030.08
00.3333330000.33333300000.25

11014
0.020.010.02
genus
0.040.020.06
00.50000.500000.333333

010001
0.020.01
species
158372

0.02
327108
no rank
00000000000.5
0.04

species
1930960
00000000001
0.02

0.040.020.020.06
0000.500.50000000000.50000001
39780
0.020.010.010.03
no rank

1848042
species
0.03
00000000000000000000002

687917
species
0000000000000001
0.01

species
483537
0.02
0001

000001
0.01
species
1568973

00.3333330.666667000.33333300.5217390.50000.3333331.502.895831.666672.454550.3333330.3333330.58333300.333333
0.120.180.030.230.20.120.140.480.120.330.060.030.240.03
family
564644
0.040.060.010.070.050.040.040.150.040.110.020.010.080.01

1680130
0.040.110.040.1
no rank
00000000000002.5062.54
0.080.220.080.2

0000000000000501258
0.040.110.040.1
1840644
species

2003395
0.020.01
genus
00000000.3333330000000000000.333333
0.060.03

0.040.02
00000000.50000000000000.5
0.020.01
2003398
species

no rank
1478132
000000010000000000001
0.020.01

0.040.080.060.02
00.500000000000.5000.8333330000000.5
no rank
331625
0.020.040.030.01

00000000000000020000001
0.020.01
species
1353016

1908807
species
0.04
0000000000001

01
0.02
1908808
species

0.01
0000000000000001
species
1914154

00.51000.5010.66666700000.500.400.50.50.51
0.040.120.020.10.150.020.050.020.040.020.13
genus
0.020.060.010.050.050.010.020.010.020.010.06
2003394

0.020.010.010.01
01000100000000010001
362692
species

species
310749
0000000000000000001
0.01

000000000000010000002
0.010.03
species
1514896

species
2003680
0.050.01
0000000010000000.5
0.10.02

no rank
1468454
0000000020000001
0.050.01

0.050.03
000000020000000000002
species
1131702

species
1408133
0.060.010.010.01
002000000000000001101

family
11012
0.020.020.070.020.040.010.01
0.080.060.210.060.130.030.03
0.2500000.66666710.333333000001.6153800000000.3333330.333333

0.060.140.09
0.333333000001.50000002.33333
1511809
0.020.070.04
genus

species
0.02
674984
0.5
0.04

235994
no rank
0.02
1

0.04
00000000000005
1323529
species

674982
species
0.01
00000000000001

0.07
0000003
species
37961

37960
0.020.020.010.01
no rank
0.040.040.020.02
00000100.500000000000000.50.5

00000001000000000000001
0.020.01
species
1849335

species
1685502
000002
0.02

species
1408895
0000000000000000000001
0.01

00.33333300000.33333300.2500.3333330.222222010000.3333330.18181800.583333
0.120.0400.060.080.040.060.180.1300.030.110.12
0.040.0100.020.020.010.020.050.0400.010.030.04
11006
family

genus
11007
0.030.01
0.060.02
0000000000000200000

species
11008
0.01
0000000000000000000

1167690
species
00000000000004
0.03

00.500000.50000.50000000.50.2500.875
0.080.040.040.020.040.08
39756
0.040.020.020.010.020.04
no rank

1608533
species
010000000000000000001
0.020.01

1856030
species
0.03
000000000000000000002

species
282786
0.020.020.01
010000000010000001

0.020.01
0000001000000000001
species
663282

0.01
0000000000000000000
1921825
species

0.03
00000000000000.333333
genus
39755
0.01

0.02
00000000000000.5
363800
0.01
no rank

species
1497019
0.01
00000000000001

genus
674981
0.0100.020.010.050
0.0300.060.030.130
000000000.333333000.3076920000

939922
0.0100.020.010.030
no rank
000000000.5000.3333330000
0.0200.040.020.060

0000000010000000
0.0100.020.010.010
1460374
species

species
1547580
0.02
000000000001

species
939923
0.02
000000000001

0000000000000000000.5
0.02
0.01
674952
genus

species
674953
0000000000000000001
0.01

0.06
0000000000000000000.666667
family
0.02
1511860

1685110
0.02
no rank
0000000000000000001
0.04

114871
species
0000000000000000002
0.02

0.80.360.40.351.230.440.570.690.290.30.420.280.641.410.212.220.831.10.580.40.90.420.27
0.30.3333330.3750.2285710.8292680.3181820.3157890.2028990.2068970.20.3333330.2142860.53.659570.3333333.518021.34942.563640.3275860.40.4444440.3809520.333333
0.190.080.10.090.290.090.150.170.070.060.10.060.160.30.070.440.170.220.130.090.190.10.07
10880
family

0.340.120.180.060.50.20.060.140.180.180.220.140.050.120.180.040.140.190.07
0.4705880.6666670.6666670.3333331.720.50.3333330.2857140.22222200.5555560.27272700.57142900.20.33333300.3888890.250.2142860.7368420.285714
689832
0.110.040.060.020.170.050.020.050.050.060.060.040.010.030.060.010.040.070.03
subfamily

0.020.020.030.020.050.040.020.010.030.010.010.020.01
10912
genus
0.5000.500.5555560.500.307692000.501000.33333300.6250.3333330.33333310.5
0.040.040.090.040.130.080.060.030.080.030.030.040.02

000000000000000000100.5
0.040.02
28876
0.020.01
species

10942
no rank
000000000000000000201
0.020.01

00000000000001.5000.5
0.040.02
0.020.01
1348384
species

0.020.01
00000000000003001
335103
no rank

0.040.02
000000000.500000000000.5
0.020.01
335100
species

0.020.01
00000000100000000001
no rank
884200

000001000.5
0.040.04
species
0.020.02
183405

994994
no rank
0.020.02
000002001

36427
species
0.01
0000000000000000001

0.020.020.02
0001000000010000000002
183407
species

10000010000100000000001
0.020.020.020.01
28875
species

000000.5
0.02
0.01
101358
no rank

1637496
species
0.01
000001

0.010.010.01
10985
genus
0.020.020.02
000000.500000000.500000.5

0.010.010.01
0000010000000100001
10986
species

00000000000000000.5
0.04
genus
0.02
208294

species
104580
00000000000000001
0.01

246280
species
0.01
00000000000000001

0.040.04
0.50000000000.5
genus
699182
0.020.02

species
338781
10000000001
0.020.02

genus
0.070.040.060.170.020.050.040.020.010.010.010.010.030.040.01
10892
0.81102.60606100.4444440010.2500.500.250.500.500.2857141.250.5
0.150.080.120.330.040.090.080.080.020.040.020.020.070.080.02

0.060.030.03
000000000000.3333330000.33333300000.333333
species
40054
0.020.010.01

000000000000.50000.500000.5
0.040.020.02
0.020.010.01
33720
no rank

no rank
449133
0.020.010.01
000000000001000100001

22206201002000000010001
0.040.040.060.140.020.020.040.010.01
306276
species

species
1679172
0000000000000000000001
0.01

40060
species
0.020.01
10000000000000001

40059
species
0000000000000100000003
0.010.03

100010000000000000001
0.020.020.01
species
40065

40052
species
00000001
0.02

0.040.040.160.040.14
0000.500000.500011.50.5
no rank
0.020.020.080.020.07
36446

species
1427476
000000001
0.02

000100000000001
0.020.07
1670669
species

species
352248
0.080.02
00000000000023

subfamily
689831
0.070.040.030.050.120.040.120.120.060.040.080.240.430.130.220.070.070.160.030.04
0.2962960.250.250.250.3636360.2666670.4444440.26315800.250.28571400.55.3978504.5086723.204550.4444440.5555560.6491230.1538460.538462
0.270.160.120.160.440.150.360.380.240.140.320.931.730.540.880.270.270.570.130.13

00.33333300000.50.5000000.600.40.66666700.3333330.51.235290
0.060.040.040.050.050.060.030.020.170.04
genus
0.020.020.020.020.020.020.010.010.060.01
10882

0.01
0000000000000001
38170
species

75888
species
0.020.020.010.010.01
000000110000020000011

00.5000000000000.500000.502
0.040.020.020.1
no rank
0.020.010.010.05
381638

00000000000001
0.01
species
1821756

species
1462685
0.020.05
010000000000000000004

667093
species
0.01
0000000000000000001

1459003
0.020.01
species
0000000000000000100000
0.040.03

1388882
no rank
0.01
0000000000000000000000

0.020.01
0000000000000000200000
no rank
419697

0.02
0000000000000000.5
species
0.01
351073

538123
no rank
0.01
0000000000000001

0.020.03
000000.50000000000.333333
genus
311227
0.010.01

no rank
0.01
547124
0.02
0000000000000000.5

0.01
0000000000000001
1840528
species

species
311228
0.01
000001

0.570.630.270.54
00000000000008.6666707.666673.666675
genus
10979
0.190.210.090.18

0.190.210.090.18
56094
species
0.380.420.180.36
000000000000013011.55.57.5

173082
no rank
0.190.210.090.18
0000000000000260231115

0.50000.50.3333330.8571430.444444000.5000.500.6666670.5000.333333001
0.040.040.030.140.090.040.020.060.020.030.06
0.020.020.010.070.050.020.010.020.010.010.03
10981
genus

species
205899
0.020.020.020.02
10001011

0.01
00000000000001
species
205895

00000021001000001000002
0.050.020.020.010.03
species
205908

species
0.010.020.01
205910
000000.500000000010000.5
0.020.040.02

134606
no rank
00000100000000020001
0.010.020.01

0.120.060.090.110.280.040.060.130.180.060.240.030.050.060.170.120.240.060.03
0.3333330.3333330.3333330.3636360.50.250.3333330.30769200.3333330.33333300.6666670.66666700.40.66666700.64705910.6666670.3333330.333333
10988
0.040.020.030.050.10.020.020.050.060.020.080.010.020.020.060.040.090.020.01
genus

0.040.040.040.1400.040.040.060.090.080.10.020.02
0.50.500.50.85714300.50.500.5000000001.111111.51.30.50.5
73145
0.020.020.020.0700.020.020.030.050.040.050.010.01
no rank

10990
species
0.020.020.0500.020.020.030.030.040.040.010.01
01012011010000000033311

0.020.020.010.01
100010000000000000101
185954
species

0.040.0600.040.060.040.040.020.02
0.500.500000.500.50.500000100.500.5
no rank
0.020.0300.020.030.020.020.010.01
519496

species
519497
100000010110000020101
0.020.020.030.020.020.010.01

001000
0.030
984915
species

no rank
0.020.010.080.010.010.010.01
73144
0.040.020.160.020.020.020.02
00000.50.50000001100.500000.50.5

species
77698
0.020.010.080.010.010.010.01
0000110000002201000011

0.020.010.01
000100000000000100001
species
33724

0.480.120.03
00000000000000060100.333333
genus
0.160.040.01
699180

no rank
0.160.040.01
1567403
000000000000000901.500.5
0.320.080.02

1567404
species
000000000000000180301
0.160.040.01

0.500000.500000001
0.040.020.02
genus
0.020.010.01
10911

species
62352
10000100000002
0.020.010.01

00000000000002
0.12
family
0.04
10877

0.04
39781
no rank
0.08
00000000000003

00000000000006
0.04
593131
species

0.020.120.020.220.020.02
000000.500010000.50600.500.5
0.010.060.010.110.010.01
451344
no rank

0.110.01
0000000000000001201
762906
species

species
1115696
00000100000001000001
0.010.010.01

0000000002
0.06
1032473
species

0.640.820.871.070.71.050.580.580.861.030.721.040.680.750.650.570.450.621.231.150.780.741.02
12333
no rank
1.281.631.742.141.392.11.141.141.712.071.452.081.351.511.291.120.91.232.462.291.571.462.02
3.382813.509233.883182.0143917.95713.043863.456142.567253.396144.634482.769231.49635.67551.19384.669643.52.6422812.1268.209615.343957.712339.71782

species
564886
000000000000050001
000.010.020.020.040.01

373126
species
0034000040000000006021
0.10.090.090.070.030.01

species
1448276
0.040.010.020.050.020.0400.030.010.030.010.010.01
00010110200010013010101

447909
species
10001120110601010022101
0.020.020.010.050.020.030.120.010.010.020.020.010.01

798951725931262181151042187710
0.130.20.260.220.120.150.050.130.210.090.230.120.080.130.070.140.080.050.240.10.090.080.13
1883368
species

species
280702
0.190.040.010.020.020.020.040.040.010.010.010.010.010.010.01
102000101100116011111011

00100000021000000011002
0.030.060.020.010.010.03
1483485
species

445517
species
00.020.0200.01
000000000000000000000

00000010000000003000001
0.0100.020.0100.030.01
species
1448273

000001000001210112210311
0.010.010.020.020.080.010.030.10.020.020.010.040.010.01
1448274
species

000000000000020000000
00.010.010.0200.010.010.010
430796
species

1302150100020182602411142
0.020.060.050.020.040.020.040.130.140.050.020.050.140.010.040.03
1874539
species

0.060.030.050.020.030.050.040.060.10.040.010.020.020.030.10.0400.03
02121300213505012227302
1883365
species

00000000000000000000142
0.010.040.03
species
1874540

0.020
0000000000000000
species
1838138

1838137
species
0000000000000000
0.020

species
445515
0.040.01
00000000000010001

species
139871
0.010.070.10.050.010.010.050.020.020.10.010.080.020.020.080.130.02
000345002015011022600901

02005321131406045654535
00.040.010.120.030.050.040.030.110.020.080.010.050.040.040.070.060.050.060.030.07
1883369
species

247623008545130041560244
0.040.090.230.140.060.030.010.190.170.080.110.040.020.030.020.060.080.010.260.06
1883379
species

812614663111131313106151121494529211936
0.160.250.190.330.150.540.280.290.070.410.260.220.250.110.070.110.120.110.510.360.280.210.48
1883366
species

1392231
species
0.020.010.030.010.030.010.010.010.010.240.010.010.04
000103000000030111119013

0.020.040.030.020.030.020.030.040.040.010.030.030.01
12110300110210001030201
species
1220717

00001310000000000250001
0.020.030.030.010.010.030.070.010.01
species
1639815

1267169
species
0101001100010504051001
0.020.0200.020.020.030.040.040.060.020.01

1109714
species
00002100001000000001
0.050.010.020.01

977801
species
0.020.030.020.030.020.010.280.010.090.010.010.010.01
10000300110101401701111

species
1080237
0.040.010.01
2000010000000000001

no rank
686617
0.210.060.030.070.050.090.150.050.190.090.120.080.030.210.070.130.050.010.110.120.150.13
0.8543690.3333330.250.18750.41.354171.20.1818181.28090.600.6666670.40.4666670.2142861.61.166670.80.250.6666671.421052.472971.11111
1.030.30.120.320.250.480.750.220.890.450.540.40.150.980.350.660.250.040.510.570.740.63

0.75000.25
0.240.08
family
0.060.02
1285587

1000.333333
0.180.06
genus
1285588
0.060.02

1.5000.5
0.120.04
1285589
0.060.02
species

328668
no rank
0.060.02
3001

family
0.01
1285596
0.04
00000000000000000000.25

0.03
00000000000000000000.333333
0.01
1285597
genus

0.01
1662464
no rank
00000000000000000000.5
0.02

0.01
00000000000000000001
species
1661257

2022639
0.020.020.060.070.030.010.040.010.03
family
0.33333300000000.333333001000.33333300.55555600.333333100.3333330.666667
0.060.060.180.210.090.030.120.030.09

2022640
0.020.060.010.010.040.010.03
genus
0.040.120.020.020.080.020.06
000000000.5001.500000.500.51.500.51

species
631345
00000000100300001013012
0.020.060.010.010.040.010.03

genus
2022641
0.020.070.02
0.040.140.04
0.500000000000000.501

0.020.070.02
10000000000000102
1871153
species

0.520.240.170.20.390.60.170.640.360.240.320.120.560.280.440.20.240.450.560.41
1.307690.41666700.2352940.51.666671.50.2352941.750.7500.750.50.5833330.2521.63636100.8751.83.251.56098
1285590
0.130.060.050.050.090.150.050.160.090.060.080.030.140.070.110.050.060.120.140.11
family

1285593
0.020.040.020.030.020.080.020.070.060.050.010.010.01
genus
0.3333330.66666700.3333330100.33333300000.66666710.33333302.333331.3333300.3333330.33333300.333333
0.060.120.060.090.060.240.060.210.180.150.030.030.03

1285595
0.020.040.020.030.020.080.020.070.060.050.010.010.01
species
0.5100.501.500.5000011.50.503.5200.50.500.5
0.040.080.040.060.040.160.040.140.120.10.020.020.02

0.020.040.020.030.020.080.020.070.060.050.010.010.01
12010301000023107401101
no rank
523909

1285591
0.110.020.020.050.070.150.020.160.090.060.010.070.070.050.050.10.140.09
genus
0.330.060.060.150.210.450.060.480.270.180.030.210.210.150.150.30.420.27
20.33333300.3333330.6666672.6666720.3333332.3333310100.3333330.3333332.666672001.333332.666674.333332.33333

30.500.51430.53.51.501.500.50.54300246.53.5
0.220.040.040.10.140.30.040.320.180.120.020.140.140.10.10.20.280.18
1285594
0.110.020.020.050.070.150.020.160.090.060.010.070.070.050.050.10.140.09
species

no rank
379529
610128617303011860048137
0.110.020.020.050.070.150.020.160.090.060.010.070.070.050.050.10.140.09

genus
674979
0.03
0.09
000.333333

no rank
1379693
0.03
000.5
0.06

001
0.03
species
1379694

35237
64.8261.4164.6363.9266.7663.5362.7563.7965.0861.2162.3762.5767.8266.6269.3169.4774.6871.9965.9966.464.5766.1463.02
no rank
5.412214.307952.385634.315063.745418.233793.546613.040452.742772.601463.847884.070754.1929820.33683.4286941.610559.25821.065410.53978.3771510.60439.177595.88312
348.269999999995331.389999999996346.809999999991347.239999999993358.649999999998345.68999999999334.679999999996343.629999999995349.099999999995331.029999999993338.079999999994338.669999999996366.570000000002354.199999999994366.419999999996357.289999999994390.20999999999372.709999999994353.34999999999354.099999999995347.739999999995353.509999999993336.399999999994

1.292.963.211.961.892.861.632.922.62.953.593.031.042.942.144.662.774.062.031.591.862.881.65
2.279074.665546.598137.433671.777788.849652.220862.945215.734624.8203414.59614.561061.1923127.35030.69626263.238231.996438.3033.083742.383655.0215111.49313.40606
no rank
0.561.151.450.910.671.040.6611.111.231.611.160.361.260.652.111.261.850.720.560.71.20.67
51368

20000000000200000010231
0.040.040.010.030.030.01
species
754069

species
755272
5512263251010130161912653710
0.090.10.030.050.050.050.070.050.120.0300.020.090.150.160.150.070.060.040.080.13

754064
species
0.070.10.020.040.030.070.030.030.060.010.030.040.030.060.040.030.010.03
40310413113004043053212

00000000110002
0.020.030.01
1580591
species

species
756285
0.02
000000000001

0.020.01
0000000000100000000001
754070
species

00000001
0.02
762905
species

species
1351702
0.030.010.020.060.010.010.01
0010010000130100011

717048
0.020.250.10.020.10.20.120.160.120.150.080.160.080.060.140.190.090.130.260.10.090.150.09
no rank
0.060.760.30.060.290.60.340.490.360.460.240.460.240.170.420.570.270.380.780.30.280.440.27
0.3333330.3157890.20.3333330.2758620.90.2352940.6122450.8611110.260870.6666670.3043480.3333330.4117650.3333332.614040.629632.657891.794870.5666670.6785710.5454550.555556

425386
species
000000000000000001
0.01

0.020.250.10.020.10.20.120.160.120.150.080.160.080.060.140.190.090.120.260.10.090.150.09
128706
no rank
0.50.4705880.30.50.4210531.350.3636360.9090911.291670.38709710.4666670.50.6363640.53.921050.9444444.166672.692310.8510.8275860.833333
0.040.510.20.040.190.40.220.330.240.310.160.30.160.110.280.380.180.240.520.20.190.290.18

0100110000000101
0.020.020.010.010.01
species
1273750

species
1273754
0.010.030.0200.010.01
00000000010100000000011

0000010000010000012001
0.010.020.010.020.01
species
1273746

1273747
species
0.020.020.020.030.010.030.01
0000121001000200000021

species
1273755
00001300000100001030230
0.020.040.030.010.010.020.020.010.040.010.030.040.01

0.030.0200.070.08
0000000001010019
species
1273748

0.020.010.030.010.030.050.020.0400.010.030.040.010.010.020.01
00000301202001003031120
species
1273753

1273751
species
0.020.040.010.040.02
120001000021

01100000000000191000011
0.020.0300.070.080.010.010.01
1262528
species

33771
species
0.040.020.020.020.0200.010.080.030.01
0100021000000000107201

species
1262530
0.020.030.010.01
01100100000001

1262527
species
00000000010110000000011
0.010.030.020.0400.010.01

01010513300112022902213
0.020.020.040.020.090.070.020.040.010.020.020.110.020.030.010.04
1273749
species

0100021000000000107201
0.040.020.020.020.0200.010.080.030.01
species
222645

0.020.020.10.070.010.020.030.01
10014000000000110002031
species
491893

family
2023203
0.070.080.060.050.170.190.050.30.070.090.10.20.080.160.280.030.030.060.030.140.130.10.07
0.210.240.180.150.510.570.150.90.210.270.30.60.240.480.840.090.090.180.090.420.390.30.21
1.333331.333330.6666670.6666672.333337.333330.6666674111.666673.333330.6666677.333331.33333111.6666713.666673.3333331.66667

22113.511161.51.52.5511121.51.52.51.55.554.52.5
0.140.160.120.10.340.380.10.60.140.180.20.40.160.320.560.060.060.120.060.280.260.20.14
1805626
0.070.080.060.050.170.190.050.30.070.090.10.20.080.160.280.030.030.060.030.140.130.10.07
genus

44227222123351022243353111095
0.070.080.060.050.170.190.050.30.070.090.10.20.080.160.280.030.030.060.030.140.130.10.07
species
1450746

species
1349409
0.170.270.720.540.10.380.260.320.520.390.880.40.20.180.090.070.10.130.20.120.310.520.14
91322234441013221346205251812111710234810

00000000000000002
0.02
species
712037

4211297205310172413010011791001117662413
0.070.440.40.210.170.170.120.070.230.510.460.260.710.071.620.841.360.090.070.080.260.18
species
1349410

0.03
00000000000000.333333
family
10659
0.01

0.02
00000000000000.5
genus
10660
0.01

species
10661
00000000000001
0.01

10474
0.020.020.050.010.050.050.050.070.130.030.110.050.010.170.050.04
family
00.33333300.250.2857140.3333330.250.666667000001.294120.3333334.025640.5454551.911760.7692310.251.483330.4117650.583333
0.060.080.140.030.160.150.170.210.390.110.340.130.040.60.170.12

no rank
0.010.030.05
693626
0000000000000000.5001.502
0.020.060.1

species
1298531
000000000000000100304
0.010.030.05

10475
0.020.020.050.010.050.050.050.070.120.030.10.010.010.090.050.03
genus
00.500.3333330.4444440.50.3636361000001.833330.56.50.8333333.047620.50.3333332.333330.5833331
0.040.060.090.020.110.10.120.140.240.060.210.020.030.270.120.06

0.01
00000000000000001
species
459290

species
693628
0.02
00001

species
2032573
0.020.010.02
0000.50000000000000000.501
0.040.020.04

no rank
248496
0001000000000000000102
0.020.010.02

species
244589
0.040.120.020.090.01
00000000000005013270001

01001112000001100010012
0.020.020.010.020.050.010.070.010.010.03
459291
species

species
244590
0000000000000000000001
0.01

0.040.020.020.18
0000000.50000000.50000.5003.5
species
0.020.010.010.09
2032563

256994
no rank
0.020.010.010.09
000000100000010001007

1299307
0.010.010.030.01
genus
00000000000000000.50.500100.5
0.020.020.060.02

species
693629
0.010.030.01
00000000000000000100201

00000000000000001
0.01
species
693627

224.070000000001227.290000000001238.940000000001242.620000000002222.760000000002230.089999999999216.450000000001235.200000000002226.160000000002228.910000000002233.900000000001235.730000000001252.46221.929999999998218.979999999998215.52271.549999999996233.229999999999225.289999999999221.329999999999211.429999999999218.219999999999210.539999999999
6.426925.374592.862275.175624.1007810.21954.232393.924192.896533.243024.763155.048575.3655625.36651.5291859.072282.601627.721614.00199.9415811.050810.81396.27491
48.749.2752.0352.3348.149.5247.4951.5549.0749.6150.6751.1255.3849.0248.7449.1462.1452.149.4348.5246.0647.7245.9
28883
order

1.451351.62551.1111.367382.011834.495441.056711.569040.9993510.8345095.590255.5163515.12283.191790.5516182.001064.141569.203351.678981.766621.270912.14841.31865
14.817.4919.4621.6416.9119.7213.9319.7715.4118.4318.0622.3339.7514.8612.989.4699999999999812.1512.5414.8914.4413.5114.9613.62
family
3.964.775.335.764.675.393.975.364.144.985.076.1110.34.043.462.633.213.594.053.933.674.113.65
10744

0.060.060.060.150.180.060.060.03
00000.33333300.3333330.3333330.6666670.66666700.3333330000000.666667000.333333
1982590
0.020.020.020.050.060.020.020.01
genus

00000.500.50.51100.50000001000.5
0.040.040.040.10.120.040.040.02
0.020.020.020.050.060.020.020.01
1982591
species

0000101122010000002001
0.020.020.020.050.060.020.020.01
1462608
no rank

542960
0.040.10.060.070.10.110.020.030.050.120.040.070.060.050.010.010.050.190.050.110.07
genus
0.1666670.50.3333330.476190.3225811.5000.1538460.6666670.6666670.23809501.4444400.7333330.3333330.3333330.4705881.631580.210.619048
0.120.30.180.210.310.340.060.070.130.360.120.210.180.150.030.030.170.570.150.330.21

0.020.040.020.020.020.040.020.08
000000.5000000.500.500.500.5010.501.5
1982580
0.010.020.010.010.010.020.010.04
species

no rank
215158
0.010.020.010.010.010.020.010.04
00000100000101010102103

0.040.060.10.050.0100.040.080.020.080.060.020.060.020.020.060.02
0.50.5010.40000.50100301.50.501001.50.5
species
0.020.030.050.030.0100.020.040.010.040.030.010.030.010.010.030.01
1982581

11021000102006031020031
0.020.030.050.0200.020.040.010.040.030.010.030.010.010.030.01
1125653
no rank

0.020.10.060.020.080.10.020.020.020.120.0200.020.140.020.080.02
010.500.52.500010000000002.501.50
1231360
0.010.050.030.010.040.050.010.010.010.060.0100.010.070.010.040.01
no rank

species
1124654
0.010.050.030.010.040.050.010.010.010.060.0100.010.070.010.040.01
02101500020000000005030

0.020.040.060.020.080.10.020.020.020.120.060.020.020.040.180.040.080.02
00.50.500.52.5000100.500.500.5000.530.51.50
species
1982579
0.010.020.030.010.040.050.010.010.010.060.030.010.010.020.090.020.040.01

01101500020101010016130
0.010.020.030.010.040.050.010.010.010.060.030.010.010.020.090.020.040.01
no rank
343516

1720323
0.020.170.290.210.050.080.20.120.070.120.190.080.240.050.010.060.020.030.010.01
genus
0.51.333331.844831.095240.2222221.8751.384621.086960.8571430.751.236840.4285712.127663.5000.501.166670.410.50.5
0.040.330.580.420.090.160.390.230.140.240.380.140.470.10.020.120.050.060.020.02

0.010.050.020.020.020.020.020.020.01
0010021100110
species
1195073

242527
species
0.020.080.180.080.020.050.10.070.050.060.070.020.20.050.010.040.020.030.010.01
14531543223157001031211

species
1136535
0202000101110000001
0.050.050.0100.010.020.030.030.020.010.02

species
644524
0.020.060.060.010.010.060.020.030.0700.0100.01
01220120113000000000

1921122
0.020.020.020.010.020.010.010.01
genus
0.060.060.060.030.060.030.030.03
00000.333333000000.3333330.33333300.3333330000.6666670.333333000.3333330.333333

species
1921123
0.020.020.020.010.020.010.010.01
0.040.040.040.020.040.020.020.02
00000.5000000.50.500.500010.5000.50.5

no rank
1113547
00001000001101000210011
0.020.020.020.010.020.010.010.01

0.070.030.060.010.010.050.020.010.040.020.030.050.170.040.010.040.030.010.03
186794
genus
0.43750.250.461538000.50000.50000.90.34146300.6666670.500.2222220.250.3333330.666667
0.160.080.130.020.020.120.040.020.080.050.070.10.410.090.020.090.080.030.09

00000000000000000000
0.010.010.01
173443
species

no rank
329158
0.020.020.030.030.010.010.020.070.010.010.030.010.03
0.040.040.070.060.020.020.030.140.020.020.050.020.06
0.50.50000.83333300000000.6666670.500.50000.40.51

1168548
species
0.020.070.010.010.010.03
00000200000000101000112

00000000000000000000
0.01
species
714583

0.01
00000000000002
221029
species

1
0.02
329864
species

348058
species
0.020.020.010.010.0100
01000100000000000

000000000000000000001
0.0200.01
590739
species

00000000000000000000000
00
876449
species

10000000000000002001
0.020.010.010.010.020.01
species
10761

1000010001000100110100
0.030.010.010.010.040.020.010.030.010.010.010
155148
species

00000000000001100000000
00.010.070
10754
species

0.380.440.70.950.460.450.260.550.630.420.330.430.110.330.240.210.330.320.470.410.610.40.34
542835
subfamily
0.5069440.4855490.6472871.543910.4337350.7964070.5647060.4285710.53750.3293410.43750.350.2272731.394960.1521740.3414631.057381.953270.8502990.4722221.620691.183210.522727
1.441.732.583.531.661.670.851.822.41.671.281.60.441.190.920.821.221.071.671.442.031.311.32

0.090.060.090.06
000.3333330.333333000000.3333330.333333
1985733
0.030.020.030.02
genus

000.50.5000000.50.5
0.060.040.060.04
1985736
0.030.020.030.02
species

no rank
1415146
0.030.020.030.02
00110000011

0.150.040.30.160.220.160.130.230.220.040.090.20.210.060.020.150.320.270.070.460.40.02
1.201.10.8750.7272731.250.3076920.6956521.0909100.4444440.904.3333300.51.066675.656252.259260.4285713.89133.4250
0.070.020.150.080.110.080.070.120.110.020.040.10.10.030.010.070.160.130.040.230.20.01
1132574
no rank

00000000000100003
0.020.03
species
1206559

0000000100000100212
0.020.010.020.15
1455076
species

00000000001200000111116
0.010.020.040.010.010.010.140.07
925983
species

00000301000001100000109
0.030.020.080.010.1
445693
species

0.010.010.010.020.010.01
0000011000000000100
species
444875

445692
species
3031201130000101105112
0.060.10.020.060.020.020.070.010.010.010.060.010.010.02

1410331
species
0.010.04
000000000000000010003

species
1273711
0.010.020.020.060.040.030.020.050.030.020.010.040.010.030.0600.040.010.01
00021302100201000050310

00100100001
0.030.010.02
913183
species

0.375000.40.50.6666671.538460.92000.50.2500.3750.3333330.51.500.2857140.16666740.3333330.5
0.160.090.250.040.060.260.250.060.040.120.080.210.120.140.070.060.20.030.02
542837
0.060.030.090.020.020.120.120.020.020.040.040.070.040.050.030.020.10.010.01
genus

species
344022
00001000000002000001
0.020.010.010.01

000000000
0.02
species
0.01
1985737

0.01
000000000
no rank
418443

0001004300100000001
0.020.040.10.070.0200.01
194966
species

0.020.010.010.10.01
10000000000000001000801
species
187764

979725
0.040.020.060.020.020.020.010.040.020.070.040.040.020.010.01
no rank
0.080.040.120.040.040.040.020.080.030.140.080.080.030.030.02
0.5000.50100.50000.37500.3333330.50.752.500.333333000.5

00010200000100020000
0.020.020.010.0300.020.01
979726
species

1543201
species
0.020.020.040.010.0100.010.01
10010000000000010000

1217810
species
0.020.01
0000000100000001

species
1141139
0000000000000000501001
0.040.010.01

species
1636249
0.020.010.010.0700
1000000000000110000000

0.270.50.180.130.060.150.20.630.270.40.060.030.030.060.030.170.210.220.110.03
1.111110.560.6666670.3076920.3333330.93333300.31.063490.5555560.950.33333300.33333300.3333330.3333330.3333330.6470591.142860.6818180.4545450.333333
genus
0.090.170.060.050.020.050.070.210.090.130.020.010.010.020.010.060.070.080.040.01
477967

0.04
0000.5
0.02
1985731
species

386793
no rank
0001
0.02

1.666670.84848510.50.51.400.4444441.595240.8333331.407410.500.500.50.5011.714291.166670.7142860.5
0.180.330.120.040.040.10.090.420.180.270.040.020.020.040.110.140.120.070.02
0.090.170.060.020.020.050.050.210.090.130.020.010.010.020.060.070.060.040.01
627479
no rank

0000000030000100001
0.070.010.01
1141519
species

0.020.020.030.020.050.030.050.04
110004012000000000343
species
1406779

species
1402875
0.020.03
0000100001

species
551790
00200100003000000000001
0.060.010.060.01

species
386792
01000001
0.020.02

641832
species
0000000000000001
0.01

0.070.040.060.010.01
420000000030000000101
species
1401297

1415145
species
0200000040100000100001
0.040.090.020.010.01

0.020.020.02
00010000000100000002
species
693582

0000000000000000000000
0
species
627480

0.040.010.060.010.010.02
0200010002000000100012
1873949
species

species
1141525
0000000000000000000000
0

0.010.01
1985729
species
0.020.02
000000000000000000.5000.5

0.010.01
000000000000000001001
no rank
881915

0.04
00000000.5
species
0.02
1985730

943335
no rank
0.02
00000001

genus
1985711
0.020.020.030.020.050.020.070.020.060.030.020.020.010.020.03
0.060.060.090.060.130.040.210.080.180.090.060.060.010.040.07
0.3333330.3333330.3333330.33333300.25000.571429000.44444400.33333300.3333330.666667000.25000

000.50000000000000.51
0.060.0200.020.04
species
0.030.0100.010.02
1985717

no rank
1701808
0.030.0100.010.02
00100000000000012

species
0.020.020.020.010.0100
1985713
0.040.040.040.020.0200
0.50.50000000.500000.5000000

1735582
no rank
11000000100001000000
0.020.020.020.010.0100

0.020.020.02000.02
000000.500000000000000000
1985715
0.010.010.01000.01
species

no rank
691318
0.010.010.01000.01
00000100000000000000000

0.0100.010.01000.01
1985714
species
0.0200.020.02000.02
00000000000000000000000

00000000000000000000000
0.0100.010.01000.01
1235824
no rank

00000000000000.5
0.020.02
0.010.01
1985716
species

no rank
1701807
0.010.01
00000000000001

0000000000000000.50000.5000
0.020.020.020
species
0.010.010.010
1985718

no rank
1273713
0.010.010.010
00000000000000010001000

0000.50000100100.5
0.040.10.080.02
0.020.050.040.01
1985712
species

1647373
no rank
0.020.050.040.01
00010000200201

542838
0.020.010.030.10.020.030.070.010.020.020.010.030.080.010.040.050.040.020.03
no rank
0.500.51.20.51.4285701.5000.50.500.500.8333334.50.51.2857121.50.51
0.040.020.060.20.040.070.140.020.040.040.020.060.160.020.070.10.080.040.06

0.020.070.020.030.070.020.020.020.080.010.030.050.040.010.03
10031303001100029134312
151528
species

0.010.030.030.010.010.010.01000.01
0011010000000101000001
268748
species

1985720
0.10.070.040.02
genus
01.13333000000.571429000000.900.666667
0.30.210.10.06

0
1985723
species
0
00000000000000

1647374
no rank
0
00000000000000

0.04
00.5
species
0.02
1985726

0.02
01
no rank
674082

00000001000000
0.10
0.050
1985725
species

00000002000000
0.050
1701804
no rank

0.030.02
1985724
species
0.060.04
00000000000001.501

no rank
532077
0000000000000302
0.030.02

02000000.5
0.160.04
species
1985727
0.080.02

04000001
0.080.02
no rank
1671396

0.090.060.120.090.060.030.090.060.06
000.3333330.33333301.083330000.3333330.333333000.333333000.3333330.66666700.333333
genus
0.030.020.040.030.020.010.030.020.02
1920774

species
0.010.020.010.01
1980957
000000.500000.5000000.5000.5
0.020.040.020.02

no rank
1796998
0.010.020.010.01
00000100001000001001

species
0.02
1980956
0.04
000000000000000001

no rank
1747282
0.02
000000000000000002

0000000000.50000000.5
0.060.02
0.030.01
1920778
species

00000000010000001
0.030.01
no rank
1610834

0.01
1980955
species
0.02
00000000000000000.5

0.01
00000000000000001
no rank
1701805

0.030.020.030.01
1920777
species
0.060.040.060.02
000.50.50200000000.5

0.030.020.030.01
00110400000001
674081
no rank

0.01
1920779
species
0.02
00000000000000000000.5

no rank
1610835
00000000000000000001
0.01

genus
0.120.130.320.550.240.210.070.080.210.250.060.190.110.110.140.090.080.110.20.160.150.120.26
110456
0.380.370.981.660.710.620.20.240.630.760.180.570.330.30.410.260.230.320.610.490.460.330.78
0.3947370.5405411.122452.933730.7042251.145160.200.4126980.4473680.3333330.4035090.303031.90.1707320.3461540.4782610.68750.9672130.3469390.9347830.303030.782051

000000000.500000000000
0.020.0400.02
1985740
0.010.0200.01
species

0.010.0200.01
00000000100000000000
197783
no rank

no rank
329157
0.120.130.320.550.240.210.070.070.190.240.060.190.110.10.140.090.080.110.190.140.150.090.26
0.5769230.8333331.666674.387391.063831.731710.30769200.6315790.6938780.50.6052630.45454530.2592590.5294120.7333331.047621.487180.5517241.38710.3529411.17308
0.260.240.661.110.470.410.130.140.380.490.120.380.220.190.270.170.150.210.390.290.310.170.52

species
979535
00000000200001000000002
0.010.020.010.010.060.010.01000.03

0000000000000000000000
0.0100
496396
species

00311400000000000050001
0.10.020.020.0300.0600.01
1527506
species

0.010.020.0400.020.010.030.020.0100.020.020.020.010.030.010.010.01
00010000001000022020010
928293
species

species
532078
0.010.0100.010.010.040.020.0100.020.020.010.010.030.010.010.010.01
00000000011000011020010

0.01
00000000000000000001
1587520
species

species
526118
0.01
00000000000000000000

329886
species
00000010000000000000003
0.010.020.0100.04

10000100000019000000102
0.020.010.010.010.010.010.040.060.010.010.010.03
species
532076

0000050000000000000001
0.040.01
532075
species

species
369257
0.010.0100.010.010.040.020.0100.020.020.020.010.030.010.010.010.01
00000000011000012020010

species
1562076
0.020.010.01
000000000000000000000

0.020.010.010.010.070.010.010.010.03
10000000000000100000102
344021
species

species
1407608
0000000000000000
0

species
988656
00010000000100000001
0.020.020.01

1500756
species
0.0100.01
00000000000000000000

0.0100
0000000000000000000000
227720
species

species
866889
135144500220301000023514
0.030.060.160.340.10.050.020.050.0700.070.010.010.0100.030.040.080.020.06

10862
species
0.040.020.030.010.05
200013000000000014

942165
species
0.020.010.020.010.020.01
000001001000000000200

669008
species
0.020.030
0001000000000000

species
10759
0.020.010.010.0600.01
01000100020000000001

110457
species
00000000000010000001
00.060.01

0
0000000000000000
species
1536611

0.020.010.01
000000000000000100000
species
1204538

0.010.01
000001000000000001
species
1051673

species
1543199
0.020.010.02
0000111

0.010.020.040.010.010.010.0100.010.010.010.03
00001000000001001100102
644563
species

species
1985738
00.0100.010.010.02
0000000000000000000.50.501
00.0200.020.020.04

no rank
10760
00.0100.010.010.02
0000000000000000001102

1.621.591.862.082.22.341.671.561.361.571.81.951.141.661.141.241.111.81.51.451.141.351.06
196895
no rank
1.7841.686781.517162.864695.091449.616511.929541.665751.140411.346042.057641.766671.352237.137740.6882134.193434.0201628.76712.714732.754661.604173.172661.59641
3.753.484.084.729999999999995.139999999999995.449999999999993.693.622.923.413.994.499999999999992.473.632.632.742.483.653.193.222.42.782.23

species
948071
0.020.030.050.020.010.01
101020001000000100001

species
1636314
0.01
000001

species
1054968
0.020.010.040.070.010.050.030.050.010.060.100.030.030.010.01
1013051200340000003210

species
1090133
0.010.0100.01
00000100000000000001

21022410311202221112101
0.040.020.050.050.030.020.070.030.020.040.010.140.020.010.010.010.020.010.01
1283079
species

1357704
species
010000000000000000001
0.020.01

00.020.010.0100.040.21
000100000000100025
species
1558467

species
1874000
0.010.040.010.010.01
0000000000001000101001

0.020.020.020.08
010119
species
461711

1235661
species
0.020.02
00001000001

species
1654601
00000000001
0.02

species
503388
0.060.030.020.01
30000300100000001

1558469
species
0.020.0100.010.0200.020.0100.020.010.010
00000000000000001000

0.020.060.010.030.02
0000000100030000100022
species
1458716

32121466432105051357443
0.060.040.030.050.020.030.150.150.090.090.040.020.040.050.010.040.060.090.050.040.04
species
1327990

0000000000000000000012
0.010.02
1498172
species

species
1357732
11000100100000000000102
0.020.020.010.020.010.03

00.010.01
0000000000000000000000
1837842
species

species
1283078
014151520226371022143536
0.020.130.020.120.130.050.050.060.110.060.280.010.020.020.010.050.040.060.030.08

10302221611202002121445
0.020.10.050.020.050.020.140.030.020.040.010.020.010.020.010.050.050.07
species
979533

000.010.01
00000000000000010000
1640973
species

species
374810
00.030.040.110
00.030.050.140
00000001000.83.142860000000000

97081
0.010.020
no rank
0000000000000000000000
0.010.030

no rank
194948
00.010
0000000000000000000000

0.020.020.03
0100100001
species
1536597

00000200000001001031011
0.020.010.010.030.010.12
species
1852627

species
1589297
0.050.050.050.120.070.050.040.030.040.060.040.140.040.050.020.090.030.010.030.01
22025820112305246272131

species
1524881
0000020000000110200100
0.020.0100.010.070.020.010

100000000001036000000001
0.020.020.260.01
species
1131315

1821228
species
0000000200000200000001
0.010.010.050.020.01

species
1327983
0.120.040.050.020.080.050.020.020.050.060.060.120.010.140.070.031.110.020.040.030.110.03
621135112230312739123292

4.52.50.540.53.51.5110.51.52.51.54.509.510.51.53.521.50.5
0.340.220.10.380.040.120.140.10.10.060.140.20.240.120.340.040.020.060.180.120.080.02
species
1300003
0.170.110.050.190.020.060.070.050.050.030.070.10.120.060.170.020.010.030.090.060.040.01

951817322135390192137431
0.170.110.050.190.020.060.070.050.050.030.070.10.120.060.170.020.010.030.090.060.040.01
no rank
1300004

1010012200010001011006
0.020.030.010.050.050.010.020.010.010.010.07
1529058
species

1127357
species
00000000000000000000000
0

species
1481785
0.020.030.020.020.04
0110001000101

1636270
species
000000000000000000001
0.01

0.020.02
1000000000000002
1873958
species

0.120.010.020.020.020.040.020.020.010.020.01
000000501011130210011
941970
species

0
00000000000000
434346
species

0.050.02
000200000001
species
1188795

0.040.070.020.020.030.080.040.070.150.02
2000320011041100172
1161906
species

species
1563661
0.020.030.020.020.020.030.050.030.010.060.030.010.03
10101200110006031050212

0.010.02
00000000000000001002
1675027
species

24120324100000000020122
0.040.080.030.050.030.050.10.0200.020.010.020.03
1486423
species

941969
species
0.120.010.010.01
00000050000000000000001

species
1279083
0.01
00000000

species
1316739
100001
0.020.01

1233488
species
0.010.01
00000000000000000000011

0.020.050.020.020.010.020.020.050.01
00001600001101002024001
1309583
species

1610508
species
0000000000001000
0.060

00000001000100000001
0.020.020.01
species
1548902

0.060.040.130.050.050.030.020.020.020.030.010.080.020.030.040.010.010.01
324223110010040110233111
species
1527771

0.060.020.020.060.020.010.020.020.01
0020120002100100221
species
49964

0.040.020.030.040.010.03
210000000000040003102
species
1481187

0.050.010.01
0000000020000001001
species
1235689

0.020.190.220.090.010.120.120.050.270.270.080.040.030.020.050.050.010.080.040.09
197401552914410032441647
species
1864623

species
270673
0.040.010.010.01
20000000000000000010000

0.020.080.030.090.070.070.050.050.020.020.010.140.020.010.030.02
141438221010020152132
1327977
species

670252
species
0.020.030.010.04
00010000010000015

000105010021012052303101
0.020.040.020.040.030.090.050.020.040.040.0100.01
1328030
species

species
1873959
23121610034112002104369
0.040.060.030.050.020.050.020.090.080.020.040.140.020.010.050.040.080.1

0.020.020.010.010.010.010.010.04
10000000100002001011013
species
1461743

species
1622234
00000010001100000000021
0.020.020.020.020.01

1161939
species
00000000010004033020001
0.030.030.030.030.020.01

species
1572753
0.030.010.010.040.010.010.050.04
00100000000202000010403

0.020.05
000100002
species
1560282

56375322020581322734535112
0.10.130.110.170.120.280.050.010.050.010.110.160.040.230.170.060.030.050.060.040.060.120.03
species
1874001

0.110.10.160.350.310.110.120.050.020.090.060.140.040.060.10.10.190.150.20.040.160.04
65515131352133718010121513163143
1540097
species

0.060.040.030.020.030.10.040.040.010.030.030.030.010.030.03
03001301015211030030202
species
994601

0.020.020.010.010.020.0200.010.0200000.010.01
1000020000000000000010
species
1675598

1874002
species
0.010.050.020.020.030.010.090.040.010.010.03
00002000100100001083112

0.0100
0000000000000000000000
1560283
species

1101022010220501102100
0.030.030.020.020.010.020.050.020.040.040.040.010.010.020.020
1327985
species

species
1560284
0.01000
0000000000000000000000

0.050.010.020.050.030.040.030.060.030.020.0100.020.010.020.040.060.02
2001030111110000102241
species
1640969

species
340054
0000000001100000001002
0.010.020.010.010.030.02000.010.010.010.02

species
1100790
0.020.030.020.060.060.01
0110020002300000001

1538364
species
000000010001000001
0.020.020.01

species
754040
0.020.010.010.01
1000000000000001000001

species
754041
0.020.010.01
000000100000000000011

469660
species
0.010.020.040.020.010.01
0000010100001300100001

01000300001001
0.020.030.020.01
species
754059

1221835
species
00010300010004032103105
0.040.030.030.030.030.020.010.040.010.07

1572754
species
0.020.01
00000000001000001

1283077
species
1110011140110011102703
0.020.020.030.010.020.020.090.020.020.070.010.010.020.090.03

species
885867
0.010.010.020.030.020.01
000002000100000000101

species
1639814
0.010.0200.020.010.010.01000
00000000000001000000

1161935
species
12232701111103001076455
0.020.050.060.070.050.060.020.020.030.030.020.020.010.080.080.060.060.07

0.050.010.010.03
00000000200000000011002
species
1417599

1168281
species
000003000000000000101
0.030.010.01

0.010
00000000000000000
1640972
species

3.109091.488372.296884.5454513.006925.47793.96973.241.303030.8478261.388892.6170213.829790.71.8753.714291.25.452383.840911.850.7333331.26923
0.550.430.640.771.451.360.6610.330.460.540.940.160.470.70.160.350.150.420.440.20.150.26
0.270.220.320.390.720.680.330.510.170.230.270.470.080.230.350.080.170.070.210.220.10.080.13
1758253
no rank

0.040.10.110.050.040.050.120.030.060.020.060.060.0200.020.020.05
25320525121308020011004
1690684
species

0.020.050.020.030.030.070.040.030.01
0100060101000410000321
1647284
species

1647474
species
01012002123501103000002
0.020.020.050.050.020.060.060.10.010.0700.030.03

00000000000100000000
0.010.020.01
species
1150869

9261228631010425901225331411522
0.170.040.190.280.670.540.250.250.090.060.10.180.090.140.050.030.040.160.140.060.020.03
1589270
species

0.040.020.010.01
21000000000000001
species
1647475

species
1647286
00000300100000000000110
0.010.010.020.030.020.0200.010.010.010.01

species
1151014
0.020.020.070.01
00010000000100101

1647385
species
0.020.010.010.02
0000000000010100001002

0.020.020.010.080.030.010.10.010.010.010.01
000000110000230111011011
species
35344

0.010.030.05
000000000012
936054
species

379329
species
0.020.010.010.020.010.03
100000000010000002

1647458
species
0.010.020.010.020.010.0100.01
0000020000000100001

1982594
0.050.02
genus
0.170.06
000000000000000000000.7058820.666667

1982595
0.030.02
species
0.060.04
0000000000000000000011

0000000000000000000022
0.030.02
1678082
no rank

0.06
000000000000000000001
species
0.03
1982596

655097
no rank
000000000000000000002
0.03

genus
0.050.050.040.020.040.050.070.010.010.030.0100.0100.010.010.040.02
1921184
0.3076920.2352940000.86666710000000000000.3333330.4285710
0.130.170.10.060.120.150.210.010.050.110.0100.0100.010.030.140.06

1921186
00.0100.0700000.010.030
species
00.0200.1400000.020.060
0000001.50000000000000.510

00.0100.0700000.010.030
0000003000000000000120
1597966
no rank

species
0.020.040.010.010.020.0300.010.020000.010.01
1921185
0.040.080.020.020.040.0600.020.040000.020.02
0.50.50001.50000000000000000

0.020.040.010.010.020.0300.010.020000.010.01
1100030000000000000000
no rank
1406786

1921188
0.020.020.010.010.020.0200.010.0200000.010.01
species
0.040.040.020.020.040.0400.020.0400000.020.02
0.5000010000000000000000

1406789
no rank
1000020000000000000000
0.020.020.010.010.020.0200.010.0200000.010.01

0.7638890.3684210.2857140.85454500.5098040.1538460.1764710.4090910.6250.250.2340431.155563.341770.3333330.9795924.9809510.7258060.8311690.7894740.5454550.493333
0.720.380.210.550.050.510.260.340.440.640.20.470.91.580.210.491.050.540.620.770.570.550.75
subfamily
0.210.110.060.190.020.150.10.120.140.210.060.140.280.470.070.160.320.160.20.240.170.170.23
542836

0.333333000000000000.3333330.666667001
0.060.120.030.09
0.020.040.010.03
1982583
genus

0.040.080.020.06
0.5000000000000.51001.5
1982584
0.020.040.010.03
species

0.020.040.010.03
10000000000012003
no rank
10747

0.2666670.6666670.333333000.4666670.500.36363600.3333330.23529405.8625028.810341.666670.3333330.6842111.275860.9047620.454545
0.150.150.090.010.150.040.110.060.170.10.80.170.580.180.060.190.290.210.22
genus
0.050.050.030.010.050.020.050.020.060.040.310.080.260.060.020.060.120.080.08
680287

0.10.10.060.10.040.040.10.040.380.040.120.120.040.120.10.10.13
0.410.5000.7000.500.50.408.63158012.583332.50.51.083330.710.692308
1234428
0.050.050.030.050.020.020.050.020.190.020.060.060.020.060.050.050.07
no rank

1453367
species
121001000011021026513031
0.020.050.030.010.020.020.150.020.050.060.010.040.030.01

00000100000000000000101
0.010.010.010.010.01
470865
species

204090
species
0.020.010.020.010.010.01
000000001000010000000

0.020.020.010.030.010.01
10000200000000001000211
species
1543206

species
320832
0.010.020.030.010.010.03
00000100000104000011002

0.020.010.020.010.010.01
000000001000010000000
species
204086

species
230871
0.020.10.050.20.060.030.01
0000001000000140624000531

0.040.060.140.040.040.090.040.40.040.020.10.090.060.08
0.500.51.5010.50.4444440000.52.50.7500.5020.888889000.8333330.875
0.020.030.070.020.020.050.020.20.020.010.050.050.030.04
545935
no rank

00130011000051010420012
0.030.070.020.020.20.010.010.050.020.010.03
species
1128071

0.02
000002
species
338473

35238
species
0.020.020.020.010.020.020.01
10000001000102000020021

0.25001.3333300.7000.4444440.520.3750.41666702.363640.510.50.3751.434781.70370.7272730.750.875
0.080.040.180.020.10.040.040.090.250.080.120.220.140.120.060.080.230.270.110.080.16
0.040.020.090.010.050.020.020.050.120.040.060.110.070.060.030.040.110.140.050.040.08
1162303
no rank

0.020.010.080.010.050.030.020.010.070.030.010.010.050.070.010.03
10030100011101131145012
1162306
species

1162305
species
00000200110005011022021
0.010.020.010.010.020.030.010.040.010.010.010.030.020.010.020.02

0000000000000101001121
0.010.010.010.010.010.030.01
1010614
species

species
1162304
0.010.010.010.020.010.010.020.050.010.030.050.010.010.020.030.030.020.03
00000100110106011122102

0.180.080.040.070.040.090.10.180.020.020.020.040.070.030.06
2.50.500.501.1428600.22222211.5000100.500.50.50.714286001
0.090.040.020.030.020.050.050.090.010.010.010.020.040.010.03
186846
genus

species
10778
0.020.020.010.010.050.090.01
0100020023000000000000

0.020.010.010.010.01
00000001000000010111
species
12345

0.090.020.020.020.010.010.010.010.020.010.03
51010200000002000012002
10756
species

542959
0.020.050.040.050.070.090.040.040.040.010.050.030.070.020.020.050.040.01
genus
0.040.10.080.090.140.180.080.080.080.030.10.060.140.040.040.110.080.02
00.50011.2500.4444440.8571430.83333310.500.66666701.3230.50.51.0909120.5

01002101112101020611201
0.020.050.010.020.020.030.040.020.020.010.020.070.010.010.030.01
279280
species

0.030.020.050.060.020.020.010.030.030.010.010.030.04
0000030122010103401124
species
194699

0000.2352940000.7865170.2502.147443.108610.250.454545000.7826090.7500000
0.170.10.890.080.031.562.670.160.1100.230.160.040.040.080.04
subfamily
0.050.030.260.020.010.420.750.040.0200.050.040.010.010.020.01
1981156

genus
1981157
0.030.20.020.360.60.01
0.070.50.0611.620.01
00000001.360.33333303.355.06790000000000

0000000000010000000000
0.020.040.040.10
1981161
0.010.020.020.050
species

0.010.020.020.050
0000000000020000000000
10730
no rank

species
0.010.020.070.10
1981162
0.020.040.140.20
000000000.501.520000000000

0.010.020.070.10
0000000010340000000000
489779
no rank

000.020.060
000000000000.50000000000
species
1981164
000.010.030

194949
no rank
0000000000010000000000
000.010.030

00000002005.58
0.020.240.440.66
species
1981160
0.010.120.220.33

866553
no rank
0.010.120.220.33
00000004001116

0.020.050.010.050.110.040.010.030.01
1981159
genus
0.060.110.020.120.230.120.030.090.03
0000.3333330000.1818180000.3913040.3333330.333333001000

species
0.010.020.03
1981168
0.010.030.03
000000000001

no rank
1660365
00000000000
0.01

0.010.020.04
000000000001
species
0.010.010.04
1981169

1124655
no rank
000000000000
0.010

species
1981167
0.020.020.010.030.040.01
0.040.040.020.050.080.02
0000.50000.50000.40.50.5

1458706
no rank
0.020.020.010.020.040.01
00010001000111

0.010.030.01
1981170
species
0.010.060.02
00000000000000001.5000

1675531
no rank
0.030.01
00000000000000003000

genus
0.020.010.010.030.0200.030.040.010.010.01
1981158
0.060.020.020.070.0600.090.120.030.050.03
0000.3333330000000000.666667001100000

0.010.040.01
1981166
species
000000000000000001.50000
0.020.080.02

0000000000000000030000
0.010.040.01
1498227
no rank

species
1981165
0.020.010.010.020.0200.030.010.010.01
0000.50000000001001.5000000
0.040.010.010.020.0400.060.020.020.02

00010000000002003000000
0.0200.0200.030.010.010.01
1416032
no rank

1.192.111.831.081.021.160.781.471.651.531.491.620.510.390.810.660.770.810.590.510.83
3.361345.478673.174862.333332.176477.551721.666673.367353.696972.843144.201344.5061703.8431402.179495.8888933.714293.70372.372882.50983.27711
genus
0.390.710.610.360.340.390.260.490.550.510.490.540.170.130.270.220.250.270.190.170.27
1918522

0.20.360.320.180.180.240.130.220.250.240.260.260.090.070.090.110.130.150.10.090.16
1918525
species
0.40.720.640.360.360.470.260.440.50.480.520.520.180.140.180.220.260.30.20.180.32
58.553.53.513.21282.54.5546.56.50603.55.54.55.563.545.5

10171077275910813130120711911127811
0.20.360.320.180.180.230.130.220.250.240.260.260.090.070.090.110.130.150.10.090.16
1327972
no rank

0.40.680.580.360.320.30.260.540.60.540.480.560.160.120.360.220.260.240.20.160.24
584.53.538.52.55.564.56705.50310.54.55.553.53.54
species
0.20.340.290.180.160.150.130.270.30.270.240.280.080.060.180.110.130.120.10.080.12
1918524

1328029
no rank
1016976175111291214011062191110778
0.20.340.290.180.160.150.130.270.30.270.240.280.080.060.180.110.130.120.10.080.12

genus
1922329
0.070.070.060.040.030.050.070.090.050.080.050.10.130.140.050.450.060.060.020.010.03
0.190.210.180.10.110.150.210.270.150.240.130.30.390.420.131.350.180.180.040.030.09
0.94736810.1666670.200.8181820.66666710.7777780.20.8333330.3076920.5333335.230770.6666671.2307716.25930.7222220.7222220.250.33333300.666667

species
00.0100.010.060000.010.02
1922332
00.0200.020.120000.020.04
00000000001.50000000.51

00.0100.010.060000.010.02
0000000000300000012
no rank
260827

0.120.140.060.040.060.10.140.140.060.040.040.160.240.280.080.860.080.060.020.020.06
1.51.50.50.501.511.51.50.50.50.518.51225.51.51.50.50.501
species
0.060.070.030.020.030.050.070.070.030.020.020.080.120.140.040.430.040.030.010.010.03
1922330

no rank
1076759
3311032331112172451331102
0.060.070.030.020.030.050.070.070.030.020.020.080.120.140.040.430.040.030.010.010.03

000000000000.500000.5000
00.040.020.020.020.040.040.040.0200.040.020.020
species
00.020.010.010.010.020.020.020.0100.020.010.010
1922331

260372
no rank
00.020.010.010.010.020.020.020.0100.020.010.010
00000000000100001000

0.060.080.030.050.110.150.010.070.050.140.040.170.010.310.040.020.040.150.040.170.020.1
1982576
genus
0.180.240.090.150.330.450.030.210.150.420.10.510.030.930.120.060.120.450.120.510.060.3
110.3333330.6666671.333335.66667010.6666671.333330.32.6666700.3333331.333331.333330.66666714.3333314.333330.6666672.33333

1982577
0.060.080.030.050.110.150.010.070.050.140.030.170.010.310.040.020.040.150.040.170.020.1
species
0.120.160.060.10.220.30.020.140.10.280.060.340.020.620.080.040.080.30.080.340.040.2
1.51.50.5128.501.5120.5400.52211.56.51.56.513.5

no rank
1429767
0.060.080.030.050.110.150.010.070.050.140.030.170.010.310.040.020.040.150.040.170.020.1
33124170324180144231331327

00000.3076920.333333000000.333333000000.3333330.333333000.666667
0.130.060.060.060.030.06
genus
1984798
0.050.020.020.020.010.02

00000.50.5000000000000.5
0.040.020.02
1984799
0.020.010.01
species

no rank
1133293
0.020.010.01
000011000000000001

0.040.020.040.020.020.04
00000.50.5000000.5000000.50.5001
species
1984800
0.020.010.020.010.010.02

0000110000010000011002
0.020.010.020.010.010.02
1133292
no rank

1720324
0.050.030.020.060.010.010.03
genus
0000000100.500.50400.50.5000001
0.10.060.040.120.020.020.06

00000002010108011000002
0.050.030.020.060.010.010.03
species
280701

genus
0.040.060.10.090.050.060.20.150.020.030.10.11.50.170.210.020.020.070.070.070.110.190.3
680277
0.3333330.3750.2666670.5185190.3636360.7222220.7272730.9555560.3333330.3333330.51.230772.689414.6250.20.250.250.750.4285710.5263160.5666671.368423.40909
0.120.160.30.270.110.180.550.450.060.090.30.264.250.480.60.040.040.20.210.190.30.570.88

1987940
0.010.050.0100.440.12000.010.0100.01
species
000000100000500.5000.50000
0.020.10.0200.880.24000.020.0200.02

no rank
1237159
00000020000010010010000
0.010.050.0100.440.12000.010.0100.01

0.50.50.51011200.512100001.500.50.53.57
0.040.040.10.120.040.10.20.060.10.160.20.04000.080.020.020.160.38
species
1987941
0.020.020.050.060.020.050.10.030.050.080.10.02000.040.010.010.080.19

1436889
no rank
0.020.020.050.060.020.050.10.030.050.080.10.02000.040.010.010.080.19
111202240124200003011714

1987938
0.050.0100.390.02000.010.010.040.01
species
0000001000004.500000.5001.50
0.10.0200.780.04000.020.020.080.02

1598146
no rank
0000002000009000010030
0.050.0100.390.02000.010.010.040.01

species
000.310.02000.020.010.010.020.05
1987942
000.620.04000.040.020.020.040.1
0000000000003.51.5000010.500.52

00000000000073000021014
000.310.02000.020.010.010.020.05
no rank
1519788

0.040.040.020.060.060.0400.040.040.040.040.04
00000.50.500.5000.50000000110.50.5
0.020.020.010.030.030.0200.020.020.020.020.02
1987939
species

0000110100100000002211
0.020.020.010.030.030.0200.020.020.020.020.02
no rank
1604355

490103
species
0.020.020.020.0500.230.030.03000.010.010.020.02
01001220000054000100111

0.50.50.50.501000.500.5008.500.50.500.510.521.5
0.040.040.060.040.040.040.040.240.020.020.020.040.020.080.08
no rank
1054967
0.020.020.030.020.020.020.020.120.010.010.010.020.010.040.04

0.020.020.030.020.020.020.020.120.010.010.010.020.010.040.04
111102001010017011012143
1141137
species

genus
545932
0.240.270.10.20.180.140.120.160.120.180.060.20.030.070.010.020.040.130.070.050.070.15
0.860.870.280.60.520.390.380.460.420.540.20.650.070.210.010.060.090.410.190.170.230.45
0.9534880.5862070.4642861.40.4807690.9743590.05263160.1956520.2857140.2962960.250.35384600.1428570.33333300.1666670.1111110.8292680.5789470.05882350.2173910.577778

species
0.010.050.020.140.070.030.020.020.050.020.050.010.070.010.010.060.010.010.020.04
2040655
01031.520.50.5100.5100.50.500.50.52.50.50.50.51
0.020.10.040.280.140.060.040.040.10.040.10.020.140.020.020.120.020.020.040.08

02063411201201101151112
0.010.050.020.140.070.030.020.020.050.020.050.010.070.010.010.060.010.010.020.04
1589273
no rank

2040656
0.010.010.020.010.020.040.010.030.020000.010.010.02
species
0.020.020.040.020.040.070.020.060.040000.020.020.04
000000.500.4285710000.500000000000.5

0.010.010.020.010.020.030.010.030.020000.010.010.02
00000101000100000000001
1548906
no rank

no rank
1417865
0.130.110.050.030.020.040.040.060.050.060.040.080.0100.010.020.040.040.020.030.06
1.150.60.214286000.33333300.3636360.1428570.3333330.30.3684210000000.40.300.251.16667
0.40.30.140.060.040.090.10.110.140.120.10.190.0200.020.030.10.10.040.080.12

0.060.050.030.020.010.010.010.020.010.030.030.010.010.010.020.020.010.010.04
32100100101100000021013
species
188350

0.210.120.0600.030.0300.060.010.030.06000.030.0400.030
1.333330.6666670000.333333000000.33333300000000.2500.3333330
0.070.040.0200.010.0100.020.010.010.02000.010.0200.010
1158721
species

1229675
0.070.040.020.010.010.020.010.020.010.010.01
no rank
210000.5000000.500000000.500.5
0.140.080.040.020.020.040.020.040.020.020.02

1234701
no rank
4200010000010000000101
0.070.040.020.010.010.020.010.020.010.010.01

species
484895
00000101010100000000001
0.020.010.020.010.020.040.010.040.020000.010.010.010.02

species
2040657
0.010.020.010.020.020.020.020.010.030.020000.010.010.010.01
00.5000100000000000000000
0.020.040.020.040.040.040.040.020.060.040000.020.020.020.02

0.010.020.010.020.020.020.020.010.030.020000.010.010.010.01
01000200000000000000000
1429758
no rank

0.180.140.020.080.040.040.040.020.120.080000.020.020.020.04
21.5000.51000100.50000000000.50.5
0.090.070.010.040.020.020.020.010.060.040000.010.010.010.02
2040658
species

no rank
1406974
0.090.070.010.040.020.020.020.010.060.040000.010.010.010.02
43001200020100000000011

0.060.040.10.190.120.110.110.020.050.050.030.266.130.110.070.060.060.070.220.270.060.390.11
1920753
genus
0.180.120.280.570.340.330.310.060.130.150.070.7618.430.350.210.20.180.230.660.830.161.170.33
0.4444440.6666670.5357140.8070180.4705880.8484850.2580650.3333330.3076920.20.2857140.77631631.62070.9714290.3333330.550.4444440.3913041.954553.566270.2581.9697

species
0.040.010.070.240.010.030.030.01
1920759
0.080.020.140.480.020.060.060.02
100000000001.52.50.501.5010.5

no rank
1239384
0.040.010.070.240.010.030.030.01
2000000000035103021

000011.500000020.500.500.51100.5
0.020.10.060.360.020.0200.020.040.060.02
1920761
0.010.050.030.180.010.0100.010.020.030.01
species

1414738
no rank
0.010.050.030.180.010.0100.010.020.030.01
0000230000004101012201

species
0.040.060.10.030.020.020.030.020.044.820.040.070.010.110.190.010.320.09
1920754
0.080.120.20.060.040.040.060.040.089.640.080.140.020.220.380.020.640.18
0112020.500.50.50.5159.52.50.500.5057.50.514.53.5

1054461
no rank
0.040.060.10.030.020.020.030.020.044.820.040.070.010.110.190.010.320.09
0224041011121195101010151297

species
0.010.020.020.010.030.0100.010.310.020.010.020.010.030.020.010.010
1920756
00000.50.50.5000003.51.5001010.50.500
0.020.040.040.020.060.0200.020.620.040.020.040.020.060.040.020.020

1395983
no rank
00001110000073002021100
0.010.020.020.010.030.0100.010.310.020.010.020.010.030.020.010.010

000.50.50.50.50.5000015.51000.50.510.50.500.5
0.020.060.080.040.020.060.0200.10.90.040.020.040.040.060.040.040.020.04
species
0.010.030.040.020.010.030.0100.050.450.020.010.020.020.030.020.020.010.02
1920755

no rank
1391223
0.010.030.040.020.010.030.0100.050.450.020.010.020.020.030.020.020.010.02
001111100002112001121101

species
1920758
0.020.020.020.020.020.080.010.010.010.010.010.01
00000.5100.50.5000.50.50.5000.500.5000.5
0.040.040.040.040.040.160.020.020.020.020.020.02

no rank
1604356
0000120110011100101001
0.020.020.020.020.020.080.010.010.010.010.010.01

species
0.020.010.010.060.070.010.010.010.010.0100.03
1920757
0.040.020.020.120.140.020.020.020.020.0200.06
0000.500.5000001.50.50.500.5000.50.501

1458848
no rank
0.020.010.010.060.070.010.010.010.010.0100.03
0001010000031101001102

686602
0.10.110.110.130.130.170.260.230.260.240.160.20.050.140.140.050.060.050.180.190.180.280.24
no rank
0.20.220.220.260.260.340.510.450.520.480.330.40.10.270.280.10.120.10.350.380.360.570.47
1.50.5454550.6818180.6923080.8461542.735291.960781.444441.923081.3751.090911.20.42.740740.51.60.8333330.52.0285722.083335.035093.65957

0.020.020.010.020.030.02
010011010100000002
136084
species

0.020.010.030.020.010.020.150.070.140.120.010.030.010.010.010.010.070.050.050.16
10110263640102001064414
1327971
species

species
440250
0.010.040.040.050.010.040.020.050.040.030.010.04000.010.050.030.030.050.12
02012111202105000142249

388452
species
0.070.020.020.050.070.020.020.040.020.060.070.010.020.060.080.010.09
41012810102108101025617

0000001
0.02
species
400567

0.010.020.010.010.03000.010.01
000000000100000000000
species
67571

species
347326
0.010.060.050.050.020.10.020.060.060.080.040.010.070.040.020.010.050.010.070.01
00220614123412142014160

194802
species
00000100101202002020001
0.010.020.020.040.010.020.020.01

0.09
0000000000.333333
1982587
0.03
genus

0.03
1982588
species
0.06
0000000000.5

1244856
no rank
0.03
0000000001

1920762
0.0200.0100000.03
genus
0.0400.0100000.07
00000000000000000000000.285714

1920765
00000.01
species
00000000000000000000000.5
00000.02

1837850
no rank
00000000000000000000001
0.01

1920763
0.0100000.01
species
0.0200000.02
00000000000000000000000.5

655098
no rank
0.01000.01
00000000000000000000001

genus
371730
0.580.891.141.090.721.040.551.330.581.060.830.710.640.540.90.550.40.560.630.580.70.80.62
2.43593.565791.983821.712421.822927.285712.159242.903391.643751.06252.066392.103630.6576093.73751.140743.289475.785712.840493.695651.958822.308082.79312.24
1.562.283.093.061.922.81.573.831.63.042.411.931.841.62.71.521.121.631.841.71.982.321.75

species
10752
101810710372565510250169542977
0.190.380.330.180.240.320.060.140.150.150.10.210.080.040.140.080.070.050.020.120.080.09

no rank
0.390.510.820.90.480.720.491.190.430.910.730.50.560.50.90.410.320.50.580.550.580.720.52
481851
0.791.011.621.790.961.440.962.360.871.831.481.011.121.021.80.830.6411.161.11.161.441.04
2.405061.277231.746912.223461.145835.944443.406254.415251.988511.355193.027031.940590.93755.666671.711113.325394.285.689662.990913.008624.111113.16346

species
1477404
00213201121012162502122
0.060.020.070.020.020.020.060.020.040.020.070.060.020.0600.030.010.020.04

10200201001001000200611
0.020.010.060.010.020.020.020.030.010.020.010.080.010.01
species
490913

02121523010106020022661
0.050.050.060.040.040.050.090.010.050.010.020.020.040.030.0200.030.030.080.070.02
species
1173761

species
1406780
00200100021011310035311
0.060.010.060.020.040.010.210.010.030.060.040.010.01

species
1283071
0.180.090.220.160.050.210.270.420.180.060.250.160.040.170.210.120.220.20.230.140.130.150.13
947722411177213812331325162011101410

species
754052
0.010.020.080.090.070.060.020.20.020.120.040.040.140.070.060.020.010.050.090.010.070.01
01233718142239062147061

species
754073
0.040.020.050.010.080.020.040.010.080.050.010.030.010.01
210209110042010300001

12000300000211000100610
0.020.050.010.030.020.010.040.040.010.010.010.080.11
490912
species

0.020.020.040.010.020.070.180.040.020.040.0100.060.010.070.12
11010101362111000501069
species
942016

species
754072
110415110021030344
0.020.020.090.040.040.020.040.010.040.020.020.030.040.06

20001110211011521001
0.040.020.010.020.050.030.020.040.010.350.020.010.01
1051675
species

1636189
species
0.020.020.020.060.010.01
010010001200010000001

11210101013202000002001
0.020.020.060.020.010.020.030.060.040.010.0200.01
1416009
species

species
1566995
0.010.110.130.20.020.010.020.120.050.080.080.050.130.070.0400.040.090.020.010.040.07
05481115224239040381035

0.010.0100.020.010.010
00000000000000000
1530400
species

0.030.050.060.040.10.050.110.010.050.010.020.020.030.0100.060.050.080.090.02
011211224010003010054681
species
1173762

species
754046
0.020.030.030.070.10.070.070.010.060.040.060.030.020.010.030.090.070.040.080.08
11134803022304021276375

1987146
0.020.010.020.040.010.010.01
genus
0000.33333300.3333330.3333330000.333333000.40000000.33333300.333333
0.060.030.060.120.050.030.03

0.040.040.02
0000.50000000.5000.5
species
1987149
0.020.020.01

no rank
1208236
00010000001001
0.020.020.01

1987148
0.010.020.020.010.01
species
000000.50.50000.5000.5000000000.5
0.020.040.040.020.02

no rank
1154995
00000110001001000000001
0.010.020.020.010.01

species
0.01
1987150
0.02
000000000000000000000.5

no rank
1204178
000000000000000000001
0.01

102294
0.760.590.610.430.40.410.690.450.540.430.610.450.130.170.140.120.20.210.340.280.280.370.4
no rank
5.20533.119663.247931.770111.430383.938274.246382.133333.364492.651165.463412.422220.51.441180.51.562.12.404762.283582.581822.684214.630145.51899
1.511.171.210.870.790.811.380.91.070.861.230.90.260.340.280.250.40.420.670.550.570.730.79

00.010.010.010.03
00000000000000014
396034
species

1000100000010100014001
0.020.010.010.020.010.020.010.010.0500.010.01
444876
species

1472912
species
41120113611000000021031
0.070.030.030.050.010.320.160.020.030.020.010.040.01

1636200
species
0.020.050.01
00000220000000000000001

species
1352230
01012400001100000122021
0.020.020.050.030.020.020.010.020.020.020.01

species
179237
000102
0.020.02

000000000000
00.010.01
365048
species

0.330.190.290.140.070.090.150.080.080.190.260.170.020.020.040.090.10.110.090.170.19
1799621063361380202578861514
444861
species

species
1503929
0.020.020.020.02
00001001001003

616674
species
5202110010110412300122
0.090.040.050.020.010.020.020.020.030.070.020.030.020.030.02

0.010.01
00000100000000000001
322855
species

1337877
species
0000000000000000000000
00.010.010.010

species
444878
41003520411213023122041
0.070.020.010.070.050.050.010.090.030.030.040.050.020.020.030.020.020.020.010.040.02

0.020.010.02
0100000000000100002
1543205
species

0.150.190.190.050.120.120.120.120.260.180.250.10.040.010.070.010.020.060.060.070.10.010.15
896251455116135111125568111
175950
species

1120100000001400001007
0.020.020.060.020.040.030.010.08
species
1235559

01140402200303000031101
0.020.030.090.030.050.050.060.020.030.010.010.01
species
981335

species
1197951
01000100000000056
0.020.010.050.05

10662
30.1528.7628.4829.2226.6426.8427.8529.6625.4126.128.9127.5422.7726.1825.5926.8529.0827.7631.2129.1929.0528.6928.69
family
11.393510.04484.864668.635027.1793216.397.169256.905174.575915.356177.932257.49445.3783650.17562.1595125.70698.701850.248822.863417.219719.401617.58199.20409
113.37109.47108.1111.02103.39104.210000000001105.82111.8897.620000000000199.8400000000003110.99106.2186.770000000000297.260000000000494.669999999999794.2500000000002103.600000000001101.79115.39108.62109.15107.09109.07

1.140.690.60.391.590.841.281.141.081.320.570.871.20.780.30.540.450.570.621.350.840.91.08
6.666673.3333321.666677.3333310.66675.57812554.6666734.666673.33333120.3333336.666675.6666755.4838712798.66667
genus
1980928
0.380.230.20.130.530.280.430.380.360.440.190.290.40.260.10.180.150.190.210.450.280.30.36

0.760.460.40.261.060.560.850.760.720.880.380.580.80.520.20.360.30.380.410.90.560.60.72
10532.511168.47.57.574.575180.5108.57.58.292681810.513.513
species
1980931
0.380.230.20.130.530.280.430.380.360.440.190.290.40.260.10.180.150.190.210.450.280.30.36

1572704
no rank
201065223217151514914103612017151736212726
0.380.230.20.130.530.280.420.380.360.440.190.290.40.260.10.180.150.190.20.450.280.30.36

0.090.10.110.190.120.090.120.050.030.120.070.040.060.040.030.040.080.120.080.01
186789
genus
1.090911.545450.6333331.751.615382.142860.80.36363600.51.444440.6470590.3333331.3846200.80.88888900.7272731.777784.52.315790
0.220.220.30.440.260.210.30.110.060.270.170.120.130.10.090.110.180.240.190.04

0.50.5110.51.510000.50.50.51.501100.50.500.50
0.060.040.160.120.040.060.120.020.060.060.080.040.040.040.040.040.040.02
no rank
329160
0.030.020.080.060.020.030.060.010.030.030.040.020.020.020.020.020.020.01

0.030.020.080.060.020.030.060.010.030.030.040.020.020.020.020.020.020.01
11221320001113022011010
1815968
species

species
10678
0.070.080.030.130.10.060.060.040.030.090.040.030.020.020.030.060.120.070.01
34154621014204022025960

0000.500.500000.50.500000.500000.50.5
0.040.040.020.040.080.080.020.020.02
genus
1198139
0.020.020.010.020.040.040.010.010.01

0.010.020.010.020.020.010.01
00010000001100001000001
264729
species

431894
species
0.010.010.010.020.020.01
0000010000110000000001

0.50.370370.2794120.3684210.251.2933300.4285710.2727270.4576270.4516130.35483901.0588200.50.3846150.3333330.560.3563220.1538460.2222220.6
0.20.810.680.760.080.750.060.70.220.590.930.620.170.10.130.030.250.870.130.180.35
857479
0.060.240.210.230.020.220.020.220.070.170.270.180.060.030.040.010.080.260.040.050.11
subfamily

00.4285710.50.52631602.3529400.4444440.500.250.501.8000010.66666700.21.05556
0.140.20.190.170.040.090.040.080.120.10.010.120.120.050.18
1196844
0.070.10.090.090.020.050.020.040.060.050.010.060.060.020.09
genus

0.020.050.040.040.010.020.010.020.0200.030.020.010.03
01110501000103000021002
157239
species

00010000101100000002001
0.020.020.020.020.020.01
species
393598

00000000000000000000000
0.010.01
species
70734

0100000000000000000001
0.020.01
342564
species

10690
species
0.020.050.040.040.010.020.010.020.0300.030.020.010.04
01110501000104000021003

0.060.170.110.140.020.140.170.050.170.230.120.010.030.030.010.020.20.040.030.03
140410
genus
0.140.430.270.340.060.360.390.110.420.580.320.010.070.080.020.050.490.090.080.06
0.7142860.558140.3333330.5294120.3333331.5833300.6666670.3636360.6428570.6896550.50000.7142860.6250.50.40.4693880.2222220.3750.333333

0.03000.01
001000000000000000000
431893
species

species
35343
0.050.030.020.020.030.040.020.020.010.01
00020301112100020101

12407
species
0.01
00000000000000000000

332032
species
000000000010000010000
0.0100.020.010.01

species
406340
0.010.060.060.010.010.01
0000010202000000000111

00000000000000000002
00.03
227940
species

0.50.70.30.4615380.52.6111100.8333330.50.751.043480.750000.5100.50.6190480.50.50.5
0.040.20.10.130.040.180.120.040.160.230.1600.020.040.020.210.020.040.02
329161
0.020.10.050.070.020.090.060.020.080.110.0800.010.020.010.110.010.020.01
no rank

00000000000000000000001
0.010.01
species
1572744

01000000000000000002
0.020.02
species
910473

1647455
species
00000000000100000000
0.020.01

390978
species
0.01
00000000000000000000

species
1357825
0000000000000000000001
0.020.01

0.0400.010.020.020.020.01
0200000000100000000201
species
1530085

10680
species
0000000000000000201
0.020.01

species
1168563
000101000001
0.020.010.02

species
910474
01111700001200000001
0.020.030.020.020.060.020.040.01

0.01
0000000000000001
1647465
species

species
910475
0.020.020.020.020.020.060.06
11010200123

species
1173769
0.02
00000000000000000002

species
1649240
000000020010000000001
0.050.02000.01

29252
species
0.020.010.01
00000000001000000011

species
431892
22110001001100000001001
0.040.050.030.020.020.020.020.010.01

species
10679
00000000000000
0

00000000001000000000
0.0200
species
103216

genus
0.060.10.040.170.020.010.030.030.030.010.020.010.01
1980921
0.180.30.120.510.060.030.090.090.090.030.060.030.03
101001.666672.333330.33333300.3333330001.33333001.333330.3333330.6666670.333333000.333333

1.501.5002.53.50.500.500020020.510.5000.5
0.120.20.080.340.040.020.060.060.060.020.040.020.02
1980923
0.060.10.040.170.020.010.030.030.030.010.020.010.01
species

30300571010004004121001
0.060.10.040.170.020.010.030.030.030.010.020.010.01
no rank
1980924

00000000000000.5000000001.33333
0.060.060.120.080.010.15
1922210
0.020.020.040.020.010.05
genus

0.010.010.010.0100.05
1922213
species
0.020.020.020.0200.1
00000000000000.5000000002

00000000000001000000004
0.010.010.010.0100.05
no rank
631719

species
0.010.010.010.010
1922212
00000000000000.500000000
0.020.020.020.020

0000000000000100000000
0.010.010.010.010
no rank
207597

0.040.020
0000000000000100000000
0.020.010
1922211
species

1229754
no rank
0000000000000200000000
0.020.010

3.729172.089431.101560.6065571.243.892472.90710.5789471.1251.347222.435372.344630.161292.189660.7017540.6666672.040821.311112.294121.320751.814812.853661.77381
1.921.231.280.6110.931.830.570.961.441.471.770.310.581.140.180.490.450.850.530.540.820.84
0.640.410.430.210.340.310.610.190.320.480.490.590.110.20.380.060.170.150.290.170.180.280.28
1921527
genus

species
1921529
0.10.070.060.050.120.040.070.050.090.110.040.110.030.040.010.0200.040.040.020.030.06
2.51.50.45454512.5210.521.512.502.500.5101.510.512
0.20.140.110.10.240.080.140.10.180.220.080.220.060.080.020.0400.080.080.040.060.12

1211280
no rank
0.10.070.050.050.120.040.070.050.090.110.040.110.030.040.010.0200.040.040.020.030.06
53125421432505012032124

8.532.51182.51.51.5333.503.50.51.54.52.54.51.542.53.5
0.640.280.340.120.10.280.260.160.160.380.240.30.020.10.20.060.160.140.220.080.220.120.2
species
1921528
0.320.140.170.060.050.140.130.080.080.190.120.150.010.050.10.030.080.070.110.040.110.060.1

1765221653366707139593857
0.320.140.170.060.050.140.130.080.080.190.120.150.010.050.10.030.080.070.110.040.110.060.1
no rank
879628

0.120.190.110.030.050.10.330.040.110.110.130.190.010.040.070.010.040.040.090.080.030.140.07
1921531
species
0.240.380.220.060.10.20.660.080.220.220.260.380.020.080.140.020.080.080.180.160.060.280.14
341.50.515.56.50.521.534.5030.50.52.51.54316.52.5

no rank
1198013
0.120.190.110.030.050.10.330.040.110.110.130.190.010.040.070.010.040.040.090.080.030.140.07
68312111314369061153862135

0.20.020.180.120.220.060.160.040.080.140.40.280.10.120.420.020.040.080.080.040.040.080.1
2.5011221.500.51530.541.50.511.51.50.50.522
species
0.10.010.090.060.110.030.080.020.040.070.20.140.050.060.210.010.020.040.040.020.020.040.05
1921530

502244301210618312331144
0.10.010.090.060.110.030.080.020.040.070.20.140.050.060.210.010.020.040.040.020.020.040.05
no rank
1198014

genus
0.01
1984782
0.03
0000000000000000.333333

0000000000000000.5
0.02
species
1984783
0.01

1566993
no rank
0000000000000001
0.01

subfamily
1985285
0.270.210.370.380.240.270.240.260.30.20.230.180.150.240.160.161.070.170.280.220.370.230.28
1.080.831.461.520.931.040.961.071.210.810.860.720.620.830.640.634.30.621.110.831.490.921.13
0.7777780.5060240.5821921.532890.5483870.9230771.010420.4018690.7851240.2345680.3488370.5833330.06451611.867470.50.39682524.57670.661291.567570.6265061.187920.760870.902655

0.250.190.260.290.210.20.090.220.190.140.140.10.110.170.160.151.060.160.230.170.280.160.21
1198140
genus
0.750.550.760.870.630.560.270.660.550.430.380.30.350.450.480.443.20.450.680.480.860.480.63
1.093330.7636360.8815792.40230.7777781.1964310.50.6363640.3023260.2368420.33333302.733330.6666670.54545533.02190.9111112.264710.93751.744191.083331.22222

0.060.040.020.020.070.10.020.090.040.040.050.050.010.010.190.020.05
1965384
species
0.50.500.501.7857100.800.5263160.37500000.88888900.888889003.50.50.8
0.120.080.040.040.140.20.040.190.080.080.090.090.030.030.380.040.1

0.030.020.010.040.060.010.050.030.020.020.020.010.010.090.010.01
11000402011000020200710
no rank
925986

no rank
557393
0.030.020.020.010.030.040.010.050.010.020.020.020.010.010.10.010.04
11010301010000020200712

000.5000000000.500010000000
0.060.020.0200.0400.0400.020.02
species
1965385
0.030.010.0100.0200.0200.010.01

576791
no rank
0.030.010.0100.0200.0200.010.01
00100000000100020000000

0.060.0200.020.010.010.040.020.01
1985310
species
0.120.0400.040.020.020.080.040.02
001000000.5000.50100.52.500100.5

0.060.0200.020.010.010.040.020.01
0020000010010201500201
77775
no rank

0.010.0100.0200.0200.010.01
1965376
species
0.020.0200.0400.0400.020.02
0000000000000000100.500

no rank
1229757
0.010.0100.0200.0200.010.01
000000000000000020100

00000100.500000000.50.500.51.500.5
0.020.020.040.10.0400.020.0400.020.080.020.02
1965377
0.010.010.020.050.0200.010.0200.010.040.010.01
species

no rank
1567025
0000020100000001101301
0.010.010.020.050.0200.010.0200.010.040.010.01

0.040.040.020.040.10.060.020.040.060.140.1
0.50000100000003001.50.511031.5
species
0.020.020.010.020.050.030.010.020.030.070.05
1965379

0.020.020.010.020.050.030.010.020.030.070.05
10000200000006003122063
no rank
1340750

no rank
0.040.090.130.230.110.050.090.060.070.030.030.040.020.020.940.050.120.030.050.020.09
1256262
0.080.180.260.460.220.10.180.120.140.060.060.080.040.041.880.10.240.060.10.040.18
1224.5231.511.50.50.50.50101562511.50.53

24494632311102021124102316
0.040.090.130.230.110.050.090.060.070.030.030.040.020.020.940.050.120.030.050.020.09
1815957
species

species
1965383
0.110.020.010.030.020.050.030.020.020.050.160.020.020.040.020.010.01
30.5000.5100100.50.503.51111.50.50000
0.220.040.020.060.040.10.060.040.040.10.320.040.040.080.040.020.02

0.110.020.010.030.020.050.030.020.020.050.160.020.020.040.020.010.01
61001200201107222310000
no rank
1197935

species
0.010.010.0200
1965378
0.020.020.0400
00000.5000000000000

1636202
no rank
0.010.010.0200
00001000000000000

1985308
0.010.010.0100.020.020.010.010.020.010.020.02
species
000000000.500000.500000.500.51
0.020.020.0200.040.040.020.020.040.020.040.04

0.010.010.0100.020.020.010.010.020.010.020.02
0000000010000100001012
no rank
1527531

0.020.020.020.0200.040.0200.02
0000000000000000.5000.500
1965382
0.010.010.010.0100.020.0100.01
species

1555238
no rank
0.010.010.010.0100.020.0100.01
000000000000000100100

0.060.070.330.270.060.210.450.150.360.180.250.240.120.140.030.0300.150.130.260.210.22
0.33333300.5454550.8888890.3333331.380951.555560.6666671.666670.3333330.841.333330.3333332.2857100.3333330.33333301.333330.5384621.038460.8571431.13636
0.020.030.110.090.020.070.150.050.120.060.090.080.040.060.010.0100.050.050.080.070.08
1985292
genus

0.5011.50.52.52.512.50.51.520.5100.50021212
0.040.020.180.160.040.080.280.10.240.060.140.160.080.020.020.10.060.120.060.12
species
1985290
0.020.010.090.080.020.040.140.050.120.030.070.080.040.010.010.050.030.060.030.06

1718606
no rank
10231552513412010042424
0.020.010.090.080.020.040.140.050.120.030.070.080.040.010.010.050.030.060.030.06

1985291
0.010.020.010.030.010.030.010.0300.010.010.030.040.01
species
0.020.040.020.060.020.060.020.0600.020.020.060.080.02
000001.50000.50002000.5000.50.51.50.5

0.010.020.010.030.010.030.010.030.010.010.030.040.01
00000300010004001001131
1718669
no rank

2.333331.666671.333331.333331.333335.666672.333331.666671.66667221.666670.333333100.6666670.3333332.6666713.333331.6666714.33333
0.390.30.390.330.30.450.540.360.390.540.390.360.120.060.060.030.30.120.390.180.120.54
1980937
0.130.10.130.110.10.150.180.120.130.180.130.120.040.020.020.010.10.040.130.060.040.18
genus

species
0.130.10.130.110.10.150.180.120.130.180.130.120.040.020.020.010.10.040.130.060.040.18
1980947
3.52.52228.53.52.52.5332.50.51.5010.541.552.51.56.5
0.260.20.260.220.20.30.360.240.260.360.260.240.080.040.040.020.20.080.260.120.080.36

75444177556651302183105313
0.130.10.130.110.10.150.180.120.130.180.130.120.040.020.020.010.10.040.130.060.040.18
no rank
1647405

1984786
0.020.040.030.070.040.020.020.030.020.020.080.010.010.010.020.05
genus
0.3333330.6666670.333333011.6666700.3333330.3333330.3333330.3333330.3333330.6666670.333333000.333333000.33333300.6666671.33333
0.060.120.090.210.120.060.060.090.060.060.240.030.030.030.060.15

species
1984787
0.020.040.030.070.040.020.020.030.020.020.080.010.010.010.020.05
0.510.501.52.500.50.50.50.50.510.5000.5000.5012
0.040.080.060.140.080.040.040.060.040.040.160.020.020.020.040.1

273133
no rank
0.020.040.030.070.040.020.020.030.020.020.080.010.010.010.020.05
12103501111121001001024

186777
0.040.080.050.050.030.020.040.020.010.010.01
genus
0.6666671.3333300000.28571400.2857140.3333330.33333300.3333330.5000000.3333330.33333300.333333
0.120.240.140.140.090.060.120.060.030.030.03

0.080.160.090.090.060.040.080.040.020.020.02
1200000.44444400.4444440.50.500.50.75000000.50.500.5
no rank
483265
0.040.080.050.050.030.020.040.020.010.010.01

24000010111011000001001
0.040.080.020.020.030.020.040.010.010.01
species
1572750

0.020.01
00000010000002
483266
species

000000001000000000001
0.020.01
665033
species

0.060.020.060.120.160.040.060.06
000.5000.50000.51.51.250000101.501
genus
0.030.010.030.060.080.020.030.03
1198976

0.020.020.03
0000000000010000203
species
261726

0.030.010.030.060.060.03
001001000133000000002
499235
species

genus
0.160.10.040.060.060.090.070.090.040.090.270.090.070.080.050.080.10.090.130.150.08
1912144
2.1251.5238100.3750.4545451.454550.8333330.4285710.7058820.37500.8888891.962261.764710.52.751.090912.2521.444441.769234.066670.875
0.320.210.080.110.110.180.140.170.080.180.530.170.140.160.110.160.210.180.260.30.16

320122
species
300012201002141121344102
0.060.020.010.030.020.060.010.020.040.040.030.070.010.020.010.040.050.050.110.03

00.010.0300.030.020.010.010.010.020.020.010.020.070.010.010.020.02
00010011000003012500101
species
262071

species
559189
0.10.080.020.030.040.050.030.040.20.030.060.020.060.030.060.040.03
54001401210254072052432

0.270.240.240.430.110.150.060.120.120.030.490.30.20.150.160.250.190.20.180.120.660.09
1984776
genus
0.810.720.721.290.330.450.180.360.360.091.470.90.60.450.480.750.570.60.540.361.980.27
4.666673.666672.3333361.3333360.6666671.666671.666670.3333338.3333351.66667705.666679.66667564.66667320.33332

0.540.480.480.860.220.30.120.240.240.060.980.60.40.30.320.50.380.40.360.241.320.18
75.53.592912.52.50.512.57.52.510.508.514.57.5974.530.53
0.270.240.240.430.110.150.060.120.120.030.490.30.20.150.160.250.190.20.180.120.660.09
1984778
species

141171841825512515521017291518149616
0.270.240.240.430.110.150.060.120.120.030.490.30.20.150.160.250.190.20.180.120.660.09
no rank
1429768

genus
1918721
0.130.020.060.050.050.090.020.070.020.040.10.070.070.050.040.070.050.060.080.070.05
0.951220.3333330.3333330.2666670.3076921.7600.5714290.33333300.250.4687501.5714300.60.92857120.6153850.51.083330.5714290.533333
0.410.060.180.150.130.250.060.210.060.120.320.210.190.150.140.210.130.20.240.210.15

species
0.070.020.020.060.010.020.020.030.060.050.020.020.070.020.020.050.030.03
1918722
1.50.5000.53.500.50.500.5103011310.5211
0.140.040.040.120.020.040.040.060.120.10.040.040.140.040.040.10.060.06

no rank
1391188
0.070.020.020.060.010.020.020.030.060.050.020.020.070.020.020.050.030.03
31001701101206022621422

000.500.50.5000000.500.500.50000.501
0.020.060.020.040.020.060.020.060.020.040.04
species
0.010.030.010.020.010.030.010.030.010.020.02
1918724

0.010.030.010.020.010.030.010.030.010.020.02
0010110000010101000102
1296656
no rank

0.120.060.080.020.020.10.020.040.020.060.040.060.040.060.060.040.04
1.500.50.500.501000001011.501110.50.5
1918723
0.060.030.040.010.010.050.010.020.010.030.020.030.020.030.030.020.02
species

30110102000002023022211
0.060.030.040.010.010.050.010.020.010.030.020.030.020.030.030.020.02
1296655
no rank

0.060.360.210.060.360.120.310.320.190.030.140.190.120.430.270.120.510.160.060.150.10.11
01.083330.66666700.7777781.333330.5161290.93750.63157900.3571430.52631603.2558101.666670.752.176470.68750.3333330.5333330.50.272727
genus
1918010
0.020.120.070.020.120.040.110.10.070.010.040.070.040.150.090.040.170.060.020.050.040.03

00.5001.500.50000.50.50503031.50.50.510
00.060.1600.080.020.040.040.060.140.120.020.160.060.040.040.040.02
00.030.0800.040.010.020.020.030.070.060.010.080.030.020.020.020.01
1918011
species

010030100011010060631120
00.030.0800.040.010.020.020.030.070.060.010.080.030.020.020.020.01
57477
no rank

0.040.180.140.040.080.080.120.20.120.020.060.080.020.140.060.060.180.0400.060.020.06
02100.5211.5100.510501.51.53.50.5010.50.5
species
0.020.090.070.020.040.040.060.10.060.010.030.040.010.070.030.030.090.0200.030.010.03
1918012

no rank
1486657
042014232012010033710211
0.020.090.070.020.040.040.060.10.060.010.030.040.010.070.030.030.090.0200.030.010.03

0.450.230.110.150.280.240.230.30.230.360.360.120.210.120.180.270.180.180.360.390.33
1.40.56521700.6666670.8571431.416670.3478260.6666670.3478260.83333300.8611110.3333331.5238100.6666671.333331.185190.6666670.8888891.333331.948721.18182
genus
1921704
0.150.070.030.050.10.080.070.10.070.120.120.040.070.040.060.090.060.060.120.130.11

species
1921705
0.060.030.020.080.030.040.050.040.030.050.040.020.040.040.030.020.060.080.06
1.50.5001.51.50.510.50.50102.5012.51.51123.52
0.120.060.040.160.060.080.10.080.060.10.080.040.080.080.060.040.120.160.12

0.060.030.020.080.030.040.050.040.030.050.040.020.040.040.030.020.060.080.06
31003312110205025322474
no rank
1206558

2.510102.50.510.51.501.50.52011211.5221.5
0.180.10.040.10.020.10.080.10.080.180.140.080.060.040.040.10.060.080.120.10.1
species
1921706
0.090.050.020.050.010.050.040.050.040.090.070.040.030.020.020.050.030.040.060.050.05

0.090.050.020.050.010.050.040.050.040.090.070.040.030.020.020.050.030.040.060.050.05
52020512130314022423443
1589265
no rank

0.090.060.150.030.060.060.0600.060.090.030.03
0.3333330000.33333300.66666700.33333300.333333000000.6666670.66666700.3333330
genus
0.030.020.050.010.020.020.0200.020.030.010.01
1984788

0.060.040.10.020.040.040.0400.040.060.020.02
0.50000.50100.500.5000001100.50
species
1984790
0.030.020.050.010.020.020.0200.020.030.010.01

no rank
1691955
100010201010000022010
0.030.020.050.010.020.020.0200.020.030.010.01

0.3333330.3333330000000000000.33333300000.666667001.33333
0.060.060.050.060.210.010.060.15
0.020.020.010.020.070.010.020.05
1984773
genus

000000000000000000
0.020.020
1984775
0.010.010
species

no rank
1296662
0.010.010
000000000000000000

species
1984774
0.020.020.010.010.0700.020.05
0.040.040.020.020.1400.040.1
0.50.50000000000000.500001002

11000000000000100002004
0.020.020.010.010.0700.020.05
no rank
1296661

25.9927.8627.6726.2426.2928.124.9427.6824.4824.626.4925.3114.3919.7521.2113.3314.8320.2523.221.7625.5224.8524.61
3.88884.678392.076622.656251.873727.20322.144352.580922.411361.90613.032842.981041.132737.499240.9910422.850715.161835.175316.375433.863514.782925.474454.66111
subfamily
6.627.127.016.646.587.16.316.996.186.186.676.393.685.015.383.373.745.135.95.496.436.266.2
1198136

genus
0.020.050.020.030.010.040.040.060.060.020.010.010.050.110.040.05
1985328
0.060.130.060.090.030.120.10.20.180.060.030.010.150.330.140.15
00.23076900.3333330.2222220.33333300.250.20.250.555556000.333333001.6666701.5151500.57142900.866667

0.010.030.020.020.010.010.010.020.040.0100.050.060.020.01
1985338
species
0.020.060.040.040.020.020.020.040.080.0200.10.120.040.02
00.500.50.50.500001000.5002.502.500.500.5

no rank
1129147
01011100002001005050101
0.010.030.020.020.010.010.010.020.040.0100.050.060.020.01

00000000.50.50.50.5000000020101.5
0.020.020.020.060.040.10.040.040000.10.060.08
1985337
0.010.010.010.030.020.050.020.020000.050.030.04
species

0.010.010.010.030.020.050.020.020000.050.030.04
00000001111000000040203
707757
no rank

1.101450.50.3142861.690910.8695652.791040.6086960.7611941.037040.7012990.5789471.134331.8461510.02630.7724552.347831.391331.697371.1251.075473.329111.04255
0.690.520.351.10.690.670.460.670.540.770.380.670.781.141.670.460.230.520.760.480.530.790.47
genus
0.230.180.130.380.230.230.160.230.180.250.140.230.260.380.550.160.090.180.260.160.190.270.16
1914165

species
1914192
0.10.020.120.060.030.010.020.020.060.090.040.160.030.010.010.060.010.010.010.01
2.50.502.511.500.5001.5202.511.50.50.52.50.50.50.50
0.20.040.240.120.060.020.040.040.120.180.080.320.060.020.020.120.020.020.020.01

697227
no rank
0.10.020.120.060.030.010.020.020.060.090.040.160.030.010.010.060.010.010.010
51052301003405231151110

01000000000.500000000.50.51.501
0.020.100.020.040.0200.04000.020.020.020.0800.06
0.010.0500.010.020.0100.02000.010.010.010.0400.03
2004415
species

02000000001000000011302
0.010.0500.010.020.0100.02000.010.010.010.0400.03
no rank
1673871

0.50.50.50.51.550.510.510100.500001.51.51.502.5
0.060.060.10.080.160.180.080.10.040.180.020.080.0200.080.080.080.020.14
0.030.030.050.040.080.090.040.050.020.090.010.040.0100.040.040.040.010.07
1914194
species

no rank
576790
111131012120201000033305
0.030.030.050.040.080.090.040.050.020.090.010.040.0100.040.040.040.010.07

0.070.030.030.170.080.030.030.060.050.110.010.080.020.280.170.030.050.140.080.050.020.10.02
1914195
species
0.140.060.060.340.160.060.060.120.10.220.020.160.040.560.340.060.10.280.160.10.040.20.04
1.50.50.53.51.51.50.5111.502019.511.535.53.520.540.5

3117331223040392361174181
0.070.030.030.170.080.030.030.060.050.110.010.080.020.280.170.030.050.140.080.050.020.10.02
no rank
1567026

0.510.50.5041.51.520.50.50331.550.50.52.522.56.51
0.040.080.060.060.020.140.140.180.220.060.040.480.10.420.180.020.020.120.10.120.280.06
species
0.020.040.030.030.010.070.070.090.110.030.020.240.050.210.090.010.010.060.050.060.140.03
1914193

no rank
1052121
0.020.040.030.030.010.070.070.090.110.030.020.240.050.210.090.010.010.060.050.060.140.03
1211083341106631011545132

6.176817.837183.335774.36162.6966710.7243.192343.651693.840913.013234.81954.723131.8860912.08731.653654.433888.010458.800410.97556.045457.422588.750717.46957
13.9715.1715.0412.9714.1113.8412.7913.5813.6413.6113.1313.46.768.709999999999999.736.729999999999998.619.9711.8411.8813.1114.1213.8
10663
4.795.235.114.414.714.684.334.594.634.564.434.532.352.983.342.272.893.44.074.014.434.764.65
genus

species
1914208
0.020.010.0500.010.020.040.0300.210.02000.0200.0200.02
0.040.020.100.020.040.080.0600.420.04000.0400.0400.04
0.500010000.5000001.51000.500.500.5

1206560
no rank
10002000100000320010101
0.020.010.0500.010.020.040.0300.210.02000.0200.0200.02

329380
3.864.314.33.723.823.933.844.023.844.133.933.821.882.462.922.012.522.913.453.573.733.853.96
no rank
7.748.639999999999998.629999999999997.499999999999997.77.927.679999999999998.057.698.389999999999997.899999999999997.709999999999993.774.909999999999995.8345.045.769999999999996.917.137.479999999999997.739999999999997.97
4.356595.223382.98033.0723.474039.441922.890634.130433.954493.120384.725324.551231.3262612.58861.4945.757511.12910.73489.141826.701267.538778.77398.1744

45406
species
00100000001001000000401
0.010.010.0300.020.01000.010.010.060.02

665681053112231326723157613
0.110.130.180.150.190.090.120.070.040.030.040.040.120.090.140.060.060.020.030.190.090.070.18
1049283
species

821711173261511121411221053171619223135
0.150.450.240.270.40.280.160.390.260.370.270.230.080.150.050.260.090.190.240.290.340.48
species
238854

20110132002202001151310
0.040.020.040.040.010.010.070.070.010.030.040.040.010.020.010.010.060.020.040.11
1340810
species

species
555387
21229764399151320256112121955731452833
0.40.460.30.170.150.370.230.220.360.40.390.510.250.080.150.110.160.060.650.390.590.310.44

species
382359
282761845668603024216168416
0.050.170.090.180.160.160.10.140.150.190.160.140.020.020.040.030.190.20.110.450.09

810761040111513111011380291761626142726
0.150.220.250.150.240.340.270.380.30.350.20.220.120.060.260.150.080.180.320.180.30.35
species
1204516

321235112121020300421150
0.060.040.060.050.080.050.040.030.050.030.040.040.010.030.010.050.030.020.170.01
1493514
species

0.090.040.130.190.140.150.120.070.050.030.150.040.040.030.140.050.030.050.140.170.260.10.11
524861753218214253412142098
1636250
species

0.020.050.040.020.030.050.060.040.010.010.040.02000.030.010.020.02
12101621001100000020101
10693
species

155615332143040223431145
0.020.110.180.150.030.050.070.070.050.030.080.070.030.020.020.040.050.040.020.150.07
1493512
species

20000200000101000002120
0.040.010.020.020.020.010.020.030.0100000.030.020.030
species
1863008

279006
species
63242108432750201751165945
0.110.060.080.110.050.090.20.110.090.060.140.10.150.10.060.040.130.070.060.120.040.07

1391224
species
0.020.030.020.020.030.010.020.020.010.030.010.0100.030.01
00001300000001000000020

1493513
species
0.10.10.050.040.050.090.080.090.180.130.10.10.090.10.170.030.040.080.060.080.040.10.11
5411211337454213235656388

species
1863009
01000300000000001101000
0.010.020.010.010.030.010.010.010.010.0100.010.0100.020.010.010

0.080.060.130.140.190.160.250.310.110.120.10.170.020.110.060.050.060.050.110.040.060.05
434671910124458015066549354
species
926067

124000000001109000010141
0.230.090.020.0100.010.020.010.040.040.070.0100.010.010.010.010.050.02
species
1863010

11120300212202310232231
0.020.020.030.050.010.030.010.020.060.040.060.050.030.020.210.0100.030.030.040.030.040.01
species
1340812

0.040.030.010.010.020.030.0100.020.010.02000.010.040
21000110000000000100030
1837930
species

0000150001203306110222
0.010.010.020.040.030.040.010.130.020.020.060.010.020.010.020.030.02
species
1222338

0.170.020.090.120.090.170.020.090.020.060.160.010.070.040.070.060.110.090.210.050.17
910451071031341148510716513
268746
species

0.150.130.260.180.10.130.140.120.150.150.150.10.140.060.250.120.030.040.050.020.07
868741455657402006291023415
species
1647404

0.370.280.330.160.190.260.160.150.40.220.340.280.120.050.210.120.210.320.190.140.060.130.2
1913107830661771814363132426161141114
760939
species

21240212102230011141113
0.040.020.060.090.010.020.040.070.020.020.050.050.130.010.010.010.050.010.010.010.04
1307804
species

1871313
species
00000000000000000101411
0.010.010.010.020.020.020.010.010.010.010.060.010.02

64123620334404021122712
0.110.10.050.050.090.060.060.010.080.120.080.090.030.030.020.010.020.030.030.090.010.03
species
233894

00000000000010005101412
0.010.0600.020.040.010.010.010.060.010.03
species
115991

1815631
species
0.070.020.030.020.10.070.020.050.120.110.010.010.020.010.020.070.010.020.050.020.020.06
31114812530001018114114

1493510
species
0.120.060.120.060.110.120.110.150.150.120.070.060.090.030.030.060.710.110.080.050.080.11
63324134664320130375896478

120451340432702162472793
0.020.040.010.090.120.110.10.090.090.040.140.010.010.070.050.020.050.080.020.10.10.04
species
762660

0.040.050.260.120.080.060.250.120.040.070.040.060.030.020.550.010.050.010.010.130.050.03
228537104121303816111032
1079998
species

1863007
species
0.020.030.0100.010.010.040.0200.020.030.030.030.03
10100000002100002032202

species
329381
00000010001100010100012
0.020.010.020.020.020.020.010.0100.020.03

0.020.060.020.020.0100.020.010.0400.010.020.01
10000020101000021300111
1141138
species

3524281353563191154546335
0.060.10.060.090.070.080.020.090.120.110.10.120.120.140.070.140.030.060.050.070.040.030.07
227470
species

380676
0.010.050.020.040.060.050.040.050.020.090.010.030.01000.030.030.020.030.05
species
0.020.10.040.080.120.090.080.10.040.180.020.060.02000.060.060.030.060.1
0100.512.333330.510.5100.500.50000110.33333311.5

no rank
293178
0.010.050.020.040.060.040.040.050.020.090.010.030.0100.030.030.010.030.05
02012412120101000022123

197310
species
00000001000002000010100
0.010.020.020.0200.0100.0100

species
260149
23335141955590720254171197926
0.040.070.110.080.120.120.020.240.120.150.110.190.520.030.020.460.210.130.110.10.10.36

01101511101100001010001
0.020.040.040.020.020.050.050.040.030.010.040.020.0100.0100.020.010.02
species
1054480

1010120522205711340212232
0.190.210.030.050.010.050.060.050.070.010.10.160.070.010.210.0400.030.010.030.040.040.03
1493508
species

0.020.020.030.020.050.050.020.020.050.0100.030.020.010.010.070.050.04
11100220201112004211553
species
1673887

1837931
species
0.010.010.010000.1600.0100.0200.01
00000000000000200000101

species
1863011
0.030.030.080.030.040.010.010.040.080.030.030.020.030.010.020.160.030.070.020.04
112104001201030301132513

1493511
species
64323911113107121543363
0.120.10.10.060.070.080.020.040.030.050.060.020.010.050.070.020.010.060.050.040.050.070.04

5171212121461766148022216411141911196
0.10.360.40.280.30.120.150.430.150.20.280.170.020.160.140.140.030.140.160.240.140.220.09
879630
species

1481112
species
0.040.030.190.030.020.020.020.020.010.010.010.020.04
02108301101101001123

0.010.020.020.020.010.010.010.020.060.010.04
00000000000000000101412
1527524
species

268747
species
312128420001030116336723
0.060.030.080.040.050.070.110.050.010.020.020.10.050.040.030.070.090.030.05

01030300111103102112017
0.020.020.080.030.020.040.020.030.020.070.020.010.020.030.010.1
species
1079999

1445860
species
10110311102101021442351
0.030.010.030.020.020.030.020.030.040.020.050.030.020.010.020.010.060.050.030.050.060.03

0.180.30.260.270.130.130.180.10.250.30.30.220.040.040.070.050.090.040.080.090.160.190.03
914811515741010151016151036712172
species
760732

0.040.050.020.030.020.020.010.060.020.040.010.10.170.020.030.020.010.050.06
220104110210111192032144
species
1493509

21001620030002012253021
0.040.030.010.020.030.050.070.010.090.010.010.020.010.020.020.060.040.010.020.02
321023
species

1914206
0000000.010.010.0100.01
species
00000000000000000.50000.500
0000000.020.020.0200.02

no rank
1551641
0000000.010.010.0100.01
00000000000000001000100

0.10.020.020.060.020.10.02000.10.020.1
000010.5000.500100.5000001.5001.5
0.050.010.010.030.010.050.01000.050.010.05
1914204
species

00002100100201000003003
0.050.010.010.030.010.050.01000.050.010.05
no rank
1141141

species
1914207
0.140.080.220.170.210.160.110.180.160.140.190.240.160.180.090.130.150.130.110.230.250.17
0.280.160.440.340.420.320.220.360.320.280.380.480.320.360.180.260.30.260.220.460.50.34
3.523.53.54.5923.53.52561.512.5057.565.54.58.511.56

7477918477410123250101512119172312
0.140.080.220.170.210.160.110.180.160.140.190.240.160.180.090.130.150.130.110.230.250.17
1351740
no rank

348604
0.140.080.220.170.290.150.120.170.160.140.190.230.110.150.070.110.130.210.220.190.280.270.19
species
3.523.53.568.52.53.53.5255.51100.567.58.59.57.510.5127
0.280.160.440.340.580.30.240.340.320.280.380.460.220.30.140.220.260.420.440.380.560.540.38

747712175774101122011215171915212414
0.140.080.220.170.290.150.120.170.160.140.190.230.110.150.070.110.130.210.220.190.280.270.19
10665
no rank

300000000000.500.50000000.5110.5
0.220.040.0200.040.020.040.02000.040.480.02
species
0.110.020.0100.020.010.020.01000.020.240.01
1914202

1204522
no rank
0.110.020.0100.020.010.020.01000.020.240.01
600000000001010000001221

0.040.0400.020.040.020.040.0400.020.040.020.0200.0200.1
0.500000.5000.500001.5000000002
species
0.020.0200.010.020.010.020.0200.010.020.010.0100.0100.05
1914199

10000100100003000000004
0.020.0200.010.020.010.020.0200.010.020.010.0100.0100.05
66711
no rank

1914200
0.090.020.070.020.010.010.020.0200.010.010
species
0.180.040.140.040.020.020.040.0400.020.020
2.50.501.50.50.5000.5000.5000000.50

5103110010010000010
0.090.020.070.020.010.010.020.0200.010.010
1054834
no rank

0.500.50.521.51.50300000.50000.51010.51
0.060.060.060.20.060.180.040.30.020.020.020.0200.040.060.020.060.040.06
0.030.030.030.10.030.090.020.150.010.010.010.0100.020.030.010.030.020.03
1914201
species

1495285
no rank
10114330600001000120212
0.030.030.030.10.030.090.020.150.010.010.010.0100.020.030.010.030.020.03

0.340.940.360.080.240.50.020.120.140.060.020.060.020.10.040.020.040.12
4.5112.50.52.514.5011.50.5000200.530100.512
species
0.170.470.180.040.120.250.010.060.070.030.010.030.010.050.020.010.020.06
1914205

no rank
1002725
9225152902310004016020124
0.170.470.180.040.120.250.010.060.070.030.010.030.010.050.020.010.020.06

0.190.180.570.480.30.30.060.420.170.020.380.120.240.090.060.060.20.350.350.210.60.09
0.5263160.55555622.333330.5333333.033330.3333331.047620.23529401.052630.250.6666670.55555600.1666670.1666670.851.342861.114290.8571434.350.222222
genus
0.070.060.190.160.10.10.020.140.050.010.120.040.080.030.020.020.060.120.120.070.20.03
1913653

0.050.050.190.160.060.090.10.020.010.090.030.080.020.010.010.050.070.070.040.170.01
1913658
species
0.10.10.380.320.120.180.20.040.010.180.060.160.040.020.020.10.140.130.080.340.02
1133.515020020.51100.501.52.51.846151.57.50

no rank
1087482
0.050.050.190.160.060.090.10.020.090.030.080.020.010.010.050.070.060.040.170.01
2267210040041220103543150

species
0.010.010.040.010.020.040.040.040.010.010.010.010.020.050.050.030.030.02
1913657
0.020.020.080.020.040.080.080.080.020.020.020.020.040.090.10.060.060.04
00000.50.50.50.50.500.5000.5000.50.51.333331.5110.5

no rank
1204531
0.010.010.040.010.020.040.040.040.010.010.010.010.020.040.050.030.030.02
00001111101001001133221

genus
0.110.20.270.240.390.290.240.380.210.230.230.220.060.570.270.320.080.470.20.220.240.160.34
1913654
0.290.610.810.71.170.840.711.160.650.690.690.660.181.670.750.940.221.430.60.630.70.481.02
0.3103450.6721310.4814810.7428571.341882.916670.7464791.155171.230770.6231880.7681160.7878790.2222229.311380.5466672.755320.6818183.342662.151.650791.41.56252.44118

0.51.510.52.54.51.5210.520.5011.505.50.550120.50.5
0.040.140.160.080.280.160.180.240.10.10.160.060.320.20.020.260.020.040.120.040.04
species
0.020.070.080.040.140.080.090.120.050.050.080.030.160.10.010.130.010.020.060.020.02
1913685

13215934214102301111002411
0.020.070.080.040.140.080.090.120.050.050.080.030.160.10.010.130.010.020.060.020.02
no rank
1567028

0.020.050.030.020.060.080.020.080.010.030.020.040.020.090.030.040.080.040.060.040.09
1913684
species
0.5100.514.50.51.5000.50.50.51.5051.51.53.51.5223
0.040.10.060.040.120.160.040.160.020.060.040.080.040.180.060.080.160.080.120.080.18

120129130011130103373446
0.020.050.030.020.060.080.020.080.010.030.020.040.020.090.030.040.080.040.060.040.09
1567027
no rank

species
0.020.020.050.040.050.070.060.120.050.050.060.070.080.070.070.010.140.010.020.020.010.16
1913686
0.500.50.5141210.511.505.50.53.50.55.5010.505.5
0.040.040.10.080.10.140.120.240.10.10.120.140.160.140.140.020.280.020.040.040.020.32

10112824212301117111021011
0.020.020.050.040.050.070.060.120.050.050.060.070.080.070.070.010.140.010.020.020.010.16
no rank
1567029

0.51122.52112.51.50.51.5019.511164433.51.5
0.060.080.180.20.260.080.120.140.240.220.060.120.040.560.340.060.040.30.180.20.160.160.1
species
0.030.040.090.10.130.040.060.070.120.110.030.060.020.280.170.030.020.150.090.10.080.080.05
1913683

700939
no rank
1224542253130392221288673
0.030.040.090.10.130.040.060.070.120.110.030.060.020.280.170.030.020.150.090.10.080.080.05

00.050.040.060.020.010.0100.020.060.080.020.040.040.020.050.020.020.04
00.400.3333330000000.5100.50100.50.50.800.50.5
species
1913995
00.030.030.040.010.010.0100.010.030.040.010.020.020.010.030.010.010.02

no rank
645664
01010000001201020112011
0.020.010.020.01000.010.030.040.010.020.020.010.020.010.010.02

0.110.160.130.130.040.160.150.180.070.160.130.040.050.050.030.040.050.030.10.110.05
1985325
genus
0.330.480.390.390.140.480.450.540.230.480.370.120.150.150.070.120.150.090.30.310.15
1.090911.083330.8461541.025640.21428630.8666671.185190.34782602.041670.9729730.3333331020.5714290.3333330.80.66666711.354840.533333

0.080.120.20.180.060.160.180.160.080.040.120.040.10.040.040.080.060.10.120.04
111.520.54.51.51.50.5001.501.50310.51.511.52.50.5
species
1985326
0.040.060.10.090.030.080.090.080.040.020.060.020.050.020.020.040.030.050.060.02

1698360
no rank
22341933100303062132351
0.040.060.10.090.030.080.090.080.040.020.060.020.050.020.020.040.030.050.060.02

species
0.070.10.030.040.020.080.060.10.040.140.060.040.0300.020.0100.050.040.03
1985327
220.50.504.5120.503.51.50.51.50000.5001.51.51
0.140.20.060.080.040.160.120.20.080.280.120.080.0600.040.0200.10.080.06

0.070.10.030.040.020.080.060.10.040.140.060.040.0300.020.0100.050.040.03
44110924107313000100332
no rank
1654928

3.015871.4251.052632.424242.2222213.28072.7397321.666670.6666675.03034.813081.1258.666670.5423731.76192.666674.24495.885062.1904832.333333.33333
0.630.40.570.660.91.140.730.480.390.180.991.070.720.570.590.210.210.490.870.420.360.240.39
0.210.140.190.220.30.380.250.160.130.060.330.350.240.190.190.070.070.170.290.140.120.080.13
1913651
genus

0.020.020.020.020.020.100.0400.060.04
000000.50000000.50000011
1985353
0.010.010.010.010.010.0500.0200.030.02
species

448384
no rank
0.010.010.010.010.010.0500.0200.030.02
00000100000010000022

species
0.010.020.040.010.10.010.020.020.0300.020.0100.020.01
1985354
0.020.040.080.020.20.020.040.040.0600.040.0200.040.02
00.50.5020.50.50000.50.50000.5000.50

no rank
50948
0.010.020.040.010.10.010.020.020.0300.020.0100.020.01
01104110001100010010

0.380.220.280.40.40.720.440.320.260.120.620.640.380.380.320.120.140.320.480.220.240.160.26
52.5244214.532.51882131346.510.544.53.55
0.190.110.140.20.20.360.220.160.130.060.310.320.190.190.160.060.070.160.240.110.120.080.13
1980959
species

1054884296521616426268132189710
0.190.110.140.20.20.360.220.160.130.060.310.320.190.190.160.060.070.160.240.110.120.080.13
576789
no rank

1913650
0.250.340.150.190.340.410.370.260.210.350.270.220.120.160.170.140.060.230.210.230.60.210.15
genus
0.751.020.450.571.021.241.110.780.631.050.810.640.360.480.530.420.180.690.630.691.80.630.45
3.706674.078430.8222221.421054.6666710.95973.459462.769231.619052.466672.444442.296870.5555565.1250.3396233.809521.722225.753623.047623.942038.538893.809522.53333

0.460.580.220.240.680.670.620.480.30.540.360.360.20.260.180.240.10.440.240.360.820.30.24
671.52.5719.20964.534.54.54190.56.5395715.574.5
1913655
0.230.290.110.120.340.340.310.240.150.270.180.180.10.130.090.120.050.220.120.180.410.150.12
species

12143514391296998218113618101431149
0.230.290.110.120.340.330.310.240.150.270.180.180.10.130.090.120.050.220.120.180.410.150.12
no rank
1455074

0.020.050.040.070.080.060.020.060.080.090.030.020.030.090.020.010.010.090.050.190.060.03
1913656
species
0.510.51.504.5101120.5020.510.50.54272.51
0.040.10.080.140.160.120.040.120.160.180.060.040.060.180.040.020.020.180.10.380.120.06

0.020.050.040.070.080.060.020.060.080.090.030.020.030.090.020.010.010.090.050.190.060.03
121309202241041211841452
no rank
709484

1.302330.30769200.3939390.3333331.074070.100.3214290.2093020.2777780.8076920.44.5468701.103450.2222220.5652170.40.4761910.50.740741
0.430.130.080.330.060.270.20.070.280.430.180.260.650.640.260.290.090.230.20.210.270.220.27
genus
0.160.050.030.110.020.10.080.030.110.170.060.090.210.220.10.090.040.090.060.070.090.080.09
1913652

00.040.10.060.020.020.060.060.020.060.060.040.100.060.040.060.120.020.08
00.50102000.50.5000202.5010.51201
species
00.020.050.030.010.010.030.030.010.030.030.020.0500.030.020.030.060.010.04
1914160

no rank
1430444
01020400110004050212402
00.020.050.030.010.010.030.030.010.030.030.020.0500.030.020.030.060.010.04

1914161
0.0100.010.020.010.020.020.040.0100.060.010.030.010.010.010.0100.010.030.01
species
0.0200.020.040.010.030.020.060.0200.120.020.060.020.020.010.0200.020.060.02
0000000000000.50.5000.50000.510

00000000000011001000120
0.010.010.0200.010.020.010.060.010.030.010.010.0100.010.030.01
12353
no rank

0.1700.030.020.040.020.050.030.080.060.010.20.280.020.060.030.010.020.040.02
1.882350000.50.50.4010.3750.5019.5010.333333100.500.5
species
0.0900.020.010.020.010.030.030.050.030.010.10.140.020.030.020.010.010.020.01
1914162

no rank
1541825
40001110011021902100101
0.080.010.010.020.010.020.030.030.10.140.030.010.010.020.01

0.080.040.060.080.020.020.060.060.040.140.060.060.040.0200.060.060.040.040.040.08
10.500.502000.50.50.51.50200.50110.50.511.5
species
0.040.020.030.040.010.010.030.030.020.070.030.030.020.0100.030.030.020.020.020.04
1914155

no rank
1237364
21010400111304010221123
0.040.020.030.040.010.010.030.030.020.070.030.030.020.0100.030.030.020.020.020.04

1.680851.971011.905413.01631.803928.758452.7656.201652.226951.531533.039222.291670.53.989131.186962.739134.85953.086334.908454.683.609372.183913.816
0.941.382.221.841.022.0722.431.411.112.041.440.480.922.30.691.211.391.421.251.280.871.25
genus
1198137
0.320.460.740.620.340.690.660.810.470.370.680.480.160.30.760.230.410.470.480.410.420.290.41

0.220.280.480.520.30.660.621.20.480.420.420.380.080.160.280.120.260.240.480.520.380.180.44
333.55.531961253.55.54.50.5513.57.5510.510748
species
1914020
0.110.140.240.260.150.330.310.60.240.210.210.190.040.080.140.060.130.120.240.260.190.090.22

no rank
115992
667116381224107119110271510212014816
0.110.140.240.260.150.330.310.60.240.210.210.190.040.080.140.060.130.120.240.260.190.090.22

0.180.20.340.460.280.360.30.060.20.080.40.060.160.240.420.20.320.260.220.140.220.10.22
2.522.5531030.520.550.518.51.55.59.554.52.5424
species
0.090.10.170.230.140.180.150.030.10.040.20.030.080.120.210.10.160.130.110.070.110.050.11
1914019

545106206141101217311191095848
0.090.10.170.230.140.180.150.030.10.040.20.030.080.120.210.10.160.130.110.070.110.050.11
75320
no rank

1914021
0.090.180.190.030.070.070.080.050.050.130.140.040.070.010.080.10.020.030.010.080.07
species
2.5430.5041.51.510.53.5302.50.50.5541103.52.5
0.180.360.380.060.140.140.160.10.10.260.280.080.140.020.160.20.040.060.020.160.14

no rank
1610855
586108332176051110822075
0.090.180.190.030.070.070.080.050.050.130.140.040.070.010.080.10.020.030.010.080.07

0.040.080.280.180.10.220.280.20.160.140.280.240.080.140.70.120.060.220.20.120.240.140.04
0.50.52216.52.521.513.530.54.52.531.54.54.524.530.5
1537658
0.020.040.140.090.050.110.140.10.080.070.140.120.040.070.350.060.030.110.10.060.120.070.02
no rank

1262539
species
114421354327619563994961
0.020.040.140.090.050.110.140.10.080.070.140.120.040.070.350.060.030.110.10.060.120.070.02

1985329
0.370.240.060.170.070.040.040.150.090.120.190.10.120.120.020.030.050.090.130.110.13
genus
6.100923.666670.6666672.2411.333330.3333331.590910.92307712.892861.2413814.88235000.6666670.51.333332.333333.333333.333332.72973
1.090.720.180.50.210.120.120.440.260.360.560.290.360.340.060.080.150.270.390.330.37

species
0.370.240.060.170.070.040.040.150.090.120.190.10.120.120.020.030.050.090.130.110.13
1985342
0.720.480.120.330.140.080.080.290.170.240.370.190.240.220.040.050.10.180.260.220.24
9.236115.513.393941.520.52.413791.411761.54.378381.894741.57.545450010.823.5554.20833

1911273415339431300224710108
0.350.240.060.160.070.040.040.140.080.120.180.090.120.10.020.020.050.090.130.110.11
no rank
1540095

0.56410300.10.1904760.682540.906250.9803920.4313731.2941210.50.33333314.651160.3333330.6470593.711863.250.6285711.142860.20.751.14706
0.390.080.30.210.630.320.510.510.680.210.20.120.360.860.30.170.590.480.350.70.150.40.68
0.130.040.10.070.210.10.170.170.220.090.080.040.120.280.10.050.190.160.110.240.050.140.22
1917990
genus

1918008
0.0600.010.020.020.010.020.010.120.150.010.030.030.030
species
0.1200.020.040.040.020.040.020.240.30.020.060.060.060
1.5000.50.50.500.500001.510000.501.5101.50

0.0600.010.020.020.010.020.010.120.150.010.030.030.030
300111010000320001032030
no rank
1406791

0.040.040.060.20.10.280.120.040.040.140.20.060.260.020.10.020.020.06
0.5000.501.521310.50.504.50.51.5800.520.50.51
0.020.020.030.10.050.140.060.020.020.070.10.030.130.010.050.010.010.03
1918005
species

no rank
1141133
100103426211091316014112
0.020.020.030.10.050.140.060.020.020.070.10.030.130.010.050.010.010.03

000010.500000000.500.505.512.5010.5
0.0200.040.020.10.020.020.040.0400.020.040.020.280.060.120.020.040.04
species
1918007
0.0100.020.010.050.010.010.020.0200.010.020.010.140.030.060.010.020.02

0.0100.020.010.050.010.010.020.0200.010.020.010.140.030.060.010.020.02
000021000000010101125021
1486662
no rank

0.020.020.060.020.040.020.10.0800.0200.020.040.020.0400.060.1
000.5000.5010.500000.50000.500.5011.5
species
0.010.010.030.010.020.010.050.0400.0100.010.020.010.0200.030.05
1918009

no rank
1382932
0.010.010.030.010.020.010.050.0400.0100.010.020.010.0200.030.05
00100102100001000101023

1918006
0.020.010.020.070.040.050.010.010.040.020.0500.030.010.050.020.030.08
species
0.040.020.040.140.080.10.020.020.080.040.100.060.020.10.040.060.16
0.50001.52100010.503.5001.50020.51.52.5

10003420002107003004135
0.020.010.020.070.040.050.010.010.040.020.0500.030.010.050.020.030.08
no rank
1406781

1984785
0.0100.020.010.050.010.010.020.01000.010.0100.030.020.010.010.05
species
000010.500000000000.5010.500.51.5
0.0200.040.020.10.020.020.040.02000.020.0200.060.040.020.020.1

no rank
1698451
00002100000000001021013
0.0100.020.010.050.010.010.020.01000.010.0100.030.020.010.010.05

genus
1911929
0.580.380.320.330.290.350.150.360.20.270.30.381.050.420.760.60.270.320.460.260.280.330.41
4.308141.750.6458331.01980.8620693.019420.41860510.6393441.29631.152171.68754.342865.634922.083335.761113.481932.239583.333332.486841.890242.092782.47934
1.721.120.961.010.871.030.431.060.610.810.921.123.151.262.281.80.830.961.380.760.820.971.21

2011.51.52.50.51.5000.52884134.53.56.53.5223
0.140.020.160.180.160.080.040.140.040.161.30.241.10.480.160.180.30.180.120.10.16
1985175
0.070.010.080.090.080.040.020.070.020.080.650.120.550.240.080.090.150.090.060.050.08
species

1195072
no rank
0.070.010.080.090.080.040.020.070.020.080.650.120.550.240.080.090.150.090.060.050.08
402335130014161682697137446

0.060.080.040.020.080.090.050.050.060.150.060.160.020.140.060.030.030.10.120.130.10.16
1985176
species
0.120.160.080.040.160.180.10.10.110.30.120.320.040.280.120.060.060.20.240.260.20.32
1.520.50.51.55.5110.9090912.51.540113214.554.54.56

0.060.080.040.020.080.090.050.050.050.150.060.160.020.140.060.030.030.10.120.130.10.16
34113112225380226429109912
584892
no rank

0.070.170.110.090.020.050.010.070.040.090.060.040.360.170.070.010.050.060.010.020.070.06
1985180
species
0.140.340.220.180.040.10.020.140.080.180.120.080.720.340.140.020.10.120.020.040.140.12
241.51.50301.50.51.51.514.511.5040.522.50.50.532

1498212
no rank
0.070.170.110.090.020.050.010.070.040.090.060.040.360.170.070.010.050.060.010.020.070.06
483306031332923081451164

0.040.120.080.220.180.20.040.10.140.060.160.120.040.140.140.060.060.260.040.060.060.08
0.51.50.521.560.511.50.521.501.50.53.51.515.5111.51.5
species
1985179
0.020.060.040.110.090.10.020.050.070.030.080.060.020.070.070.030.030.130.020.030.030.04

1314312123143031732112233
0.020.060.040.110.090.10.020.050.070.030.080.060.020.070.070.030.030.130.020.030.030.04
1792245
no rank

species
0.350.050.050.030.020.060.040.110.040.090.030.040.090.160.130.120.020.010.030.070.06
1985177
0.70.10.10.060.040.120.080.220.080.180.060.080.180.320.260.240.040.020.060.140.12
9.510.50.503.50.520.5020.50.56.508.57.550.50.5132

no rank
1129192
1921107141041113017151011264
0.350.050.050.030.020.060.040.110.040.090.030.040.090.160.130.120.020.010.030.070.06

subfamily
2.582.311.92.742.732.182.752.362.032.12.272.62.032.332.091.91.461.212.913.592.672.232.65
857473
3.078592.689740.9910452.667021.934314.384621.889462.277781.740331.593832.119112.546581.155715.584150.7720283.322063.465481.705615.167999.025874.237744.079654.02148
8.787.799999999999996.79.559.597.679.687.927.247.468.068.9178.27.156.85.069999999999994.2810.0612.379.387.919.30999999999999

857474
0.430.320.120.30.190.250.290.310.220.330.230.250.230.170.220.150.130.080.190.290.190.170.2
no rank
117.51.56414.55.564.55662.5121.587.53811.577.57
0.860.640.240.60.380.50.580.620.440.660.460.50.460.340.440.30.260.160.380.580.380.340.4

species
857312
2215312829111291012125243161561623141514
0.430.320.120.30.190.250.290.310.220.330.230.250.230.170.220.150.130.080.190.290.190.170.2

genus
0.460.470.330.370.510.380.530.490.360.420.550.510.190.220.360.40.230.180.710.530.440.510.67
1857846
1.221.210.881.041.361.041.361.2311.211.451.30.580.61.051.060.640.521.861.391.191.381.76
2.950822.801651.1251.932692.455885.461542.845592.894311.671.603313.434483.238460.6896553.716670.6095245.150943.468751.923088.107534.992813.966395.79716.27273

3.934433.054551.533.449288322.545452.54.2571431.025644.7666716.867925.1428639.60926.568.112687.90588
0.610.550.440.60.690.560.60.480.550.720.70.560.390.30.640.530.350.320.870.680.640.710.85
no rank
0.310.270.220.30.350.280.30.240.270.360.350.280.190.150.320.270.170.160.430.340.320.350.43
1206898

species
1235660
0.150.140.110.150.170.140.150.120.140.180.170.140.10.080.160.130.090.080.220.170.160.180.21
86367166455862102141061913121616

8636716645596292141061913121616
0.150.140.110.150.170.140.150.120.140.180.180.140.10.070.160.130.090.080.220.170.160.180.21
species
1235659

55510
species
893361291032101101001472241481418
0.150.190.110.070.160.10.230.260.090.070.20.230.080.050.130.060.020.280.180.110.160.24

0.630.631.161.470.450.720.750.60.960.721.050.930.840.420.640.630.660.510.630.550.790.78
1.857141.936512.241385.7551151.641.666673.541671.666673.095242.612906.476190.3333333.781254.968253.257582.529413.698412.327273.645573.61538
genus
0.210.210.380.490.150.240.250.20.320.240.350.310.280.140.220.210.220.170.210.190.270.26
1985181

species
1985182
0.090.050.130.050.070.090.090.040.040.060.120.120.120.070.10.050.150.060.040.080.140.08
0.180.10.260.10.140.180.180.080.080.120.240.240.240.140.20.10.30.120.080.160.270.16
2.51211.551.50.50.5133080.55.52.562.51.535.777783

1126950
no rank
0.090.050.130.050.070.090.090.040.040.060.120.120.120.070.10.050.150.060.040.080.130.08
5242310311266016111512536126

species
0.120.160.260.440.080.150.160.160.280.180.230.190.160.070.110.160.070.110.170.10.130.18
1985183
33.549.51.5933635.54.50110.5692.54.56.545.286.5
0.240.320.520.880.160.30.320.320.560.360.460.380.320.140.220.320.140.220.340.20.250.36

0.120.160.260.440.080.150.160.160.280.180.230.190.160.070.110.160.070.110.170.10.120.18
678193186612611902211218591381113
no rank
1690456

genus
0.030.040.080.050.170.130.20.110.060.040.030.040.050.050.040.020.040.210.050.110.10.04
1857844
0.050.10.180.090.460.350.490.290.120.10.050.110.120.110.10.060.10.590.130.310.280.11
000.2777780.4444440.9130431.542860.8775510.5172410000.18181800.54545500.40.1666670.42.220340.3076922.0322610.181818

40523
species
0.010.020.050.040.010.030.050.030.010.020.010.010.020.020.020.010.020.040.020.010.020.01
00110311000002021131020

0.010.040.0500.280.190.240.150.050.040.010.060.050.040.040.030.040.340.060.190.160.06
00001.52.368420.750.80000.33333300.50000.53.50.3333333.315791.50.333333
330395
0.010.020.0300.140.10.130.080.030.020.010.030.030.020.020.010.020.170.030.10.080.03
species

1173766
no rank
00003511000001000170030
0.010.010.070.040.040.020.010.0100.010.010.010.010.010.010.090.0100.040.01

1458855
no rank
00003522000101000171731
0.010.010.070.050.070.050.010.0100.020.010.010.010.010.010.080.020.090.040.02

0.86363600.1935480.751.251.035710.370370.781250.60.2666670.3461541.050850.483.3157901.148151.971431.4242410.8275861.630430.7727270.666667
0.440.140.310.560.840.280.270.320.250.30.260.590.50.570.240.270.350.330.350.290.460.220.24
genus
0.160.050.110.20.310.10.10.120.080.10.090.220.170.220.080.10.140.130.130.110.170.080.09
1857845

0.85714300.4285711.30.851.222220.7142861.50.8823530.3750.3751.250.8888892.58333011.51.214291.11.3333320.51.33333
0.140.060.140.20.20.090.140.140.170.080.080.160.270.120.160.060.060.140.10.120.10.060.09
0.070.030.070.10.10.040.070.070.080.040.040.080.130.060.080.030.030.070.050.060.050.030.05
365007
no rank

1168744
species
10011120100101000113010
0.020.010.010.020.030.020.050.030.010.010.020.020.010.0300.020.020.040.010

0.050.020.060.080.070.030.020.070.060.030.030.060.120.050.050.030.030.050.030.020.050.020.04
20132303211326023332413
species
1007127

species
1986015
0.040.010.020.060.110.030.010.020.050.030.070.020.080.040.050.040.020.070.020.02
100121.500.500.50.51.505.502301.512.50.50.5
0.080.020.040.120.220.060.020.040.10.060.140.040.160.080.10.080.040.140.040.04

200243010113011046032511
0.040.010.020.060.110.030.010.020.050.030.070.020.080.040.050.040.020.070.020.02
no rank
1406783

0.060.010.020.040.110.030.010.020.020.030.070.020.070.030.050.060.040.020.050.040.02
300143010013010036532431
species
10685

1857843
0.470.220.320.380.640.380.560.230.370.280.390.370.730.640.330.470.270.220.380.760.590.330.52
genus
1.280.620.850.981.741.081.60.511.010.81.130.931.961.820.871.330.710.561.082.231.670.971.43
1.531250.4838710.9529412.040822.959773.370372.118751.568631.524750.93750.6460181.79571.607145.329670.160922.556392.873241.821431.972222.865471.916170.9690723.13986

0.040.040.040.020.020.060.0200.040.020.020.020.060.040.040.04
00.5000000000.500000.50.50010.50.50.5
1924734
0.020.020.020.010.010.030.0100.020.010.010.010.030.020.020.02
species

0.020.020.020.010.010.030.0100.020.010.010.010.030.020.020.02
01000000001000011002111
no rank
1041526

1924729
0.070.10.120.10.060.270.020.090.020.060.060.070.060.030.030.060.040.040.03
species
0.140.20.240.20.120.540.040.180.040.120.120.140.120.060.060.120.080.080.06
1.501.52.523.55.50.501.50.51.5040.53.52012.51.521

0.070.10.120.10.060.270.020.090.020.060.060.070.060.030.030.060.040.040.03
303547111031308174025342
no rank
1239383

0.040.040.160.040.040.060.080.100.040.040.040.040.040.160.10.060.1
0.50.510010.50.50.50100001110.53211.5
species
0.020.020.080.020.020.030.040.0500.020.020.020.020.020.080.050.030.05
1924730

11200211102000022216423
0.020.020.080.020.020.030.040.0500.020.020.020.020.020.080.050.030.05
1450142
no rank

no rank
1970745
0.070.030.010.020.080.080.070.020.060.030.070.070.060.060.010.070.020.010.070.130.080.020.09
0.140.060.020.040.160.160.140.040.120.060.140.140.120.120.020.140.040.020.140.260.160.040.18
20.5001.54.510.5101.51.50.53.503.5103530.53

0.070.030.010.020.080.080.070.020.060.030.070.070.060.060.010.070.020.010.070.130.080.020.09
410039212033170720610616
307898
species

species
1980964
0.020.020.010.030.010.010.020.030.030.020.040.010.010.050.030.020.01
0.040.040.020.060.020.020.040.060.060.040.080.020.020.10.060.040.02
0.50.5000.50.500000.50.501.5020.500210.50

0.020.020.010.030.010.010.020.030.030.020.040.010.010.050.030.020.01
11001100001103041004210
1041527
no rank

0.010.010.010.030.020.010.010.010.010.040.080.020.010.010.040.040.060.050.05
1924731
species
00000.50.50000000.55.50100.51.51.5221.5
0.020.020.020.060.040.020.020.020.020.080.160.040.020.020.080.080.120.10.1

000011000000111020133443
0.010.010.010.030.020.010.010.010.010.040.080.020.010.010.040.040.060.050.05
no rank
1526408

0.020.010.010.010.010.010.020.010.020.020.060.030.010.020.010.010.010.010.040.010.01
10000000001113011011310
species
221915

0.500000.50.50000.501600.500.50.52001
0.060.020.020.020.060.020.060.020.160.180.0200.020.040.10.020.020.06
species
1924732
0.030.010.010.010.030.010.030.010.080.090.0100.010.020.050.010.010.03

100001100010212010114002
0.030.010.010.010.030.010.030.010.080.090.0100.010.020.050.010.010.03
760530
no rank

species
0.050.030.030.020.040.040.030.040.050.060.050.320.080.060.060.050.040.040.040.080.040.07
292029
0.090.050.040.030.070.070.040.070.10.10.080.590.130.110.10.090.070.070.060.120.070.12
10000.4285711.28571000.4285710.51.413.288144.7692302.22.333330.8571431.428571.333332.666671.857141.66667

no rank
1262512
10001300112166044222234
0.040.020.010.010.030.030.010.030.050.040.030.270.050.050.040.040.030.030.020.040.030.05

0.060.160.50.160.040.080.20.120.080.040.060.340.140.140.080.080.060.20.140.10.02
0.5001.554.50.50.52110.50120.53.521.5142.520.5
1924733
0.030.080.250.080.020.040.10.060.040.020.030.170.070.070.040.040.030.10.070.050.01
species

0.030.080.250.080.020.040.10.060.040.020.030.170.070.070.040.040.030.10.070.050.01
1003109114221024174328541
no rank
1524910

0.080.070.030.020.010.030.0700.070.030.040.020.080.020.070.030.040.080.160.10.040.1
1980962
species
21.50.50.501.51.501.50.510.511041.51.53.56.53.51.53.5
0.160.140.060.040.020.060.1400.140.060.080.040.160.040.140.060.080.160.320.20.080.2

431103303121220833713737
0.080.070.030.020.010.030.0700.070.030.040.020.080.020.070.030.040.080.160.10.040.1
no rank
1572703

no rank
0.310.530.340.50.290.340.360.510.280.210.230.330.30.430.40.150.250.120.50.410.240.280.29
1232680
2.950825.169811.602944.52.172417.8676535.431372.517861.674422.913043.727271.7666717.81181.556963.548399.861.565229.166.439023.48986.684214.25862
0.611.060.6810.580.680.721.020.560.430.460.660.60.850.790.310.50.2310.820.490.570.58

442493
species
0.080.090.110.150.150.050.110.040.060.060.040.030.020.030.020.080.060.050.080.04
44366541200305023264373

0.150.310.150.290.080.180.210.350.170.080.090.210.160.320.290.090.20.040.290.220.080.020.15
814412321814724104454924325176111
species
632112

species
298338
0.070.130.080.060.060.110.040.120.050.140.140.060.140.060.070.050.020.050.130.130.120.190.1
462221215247339152411109177

7.80185.311931.364.149535.104768.666673.379313.11342.816093.421054.225816.41862.786679.79732.1688.382025.652173.490214.224630.907714.939812.070810.0294
1.111.090.51.071.050.841.160.970.871.140.931.290.750.741.250.890.460.511.382.711.661.131.36
1857889
0.510.480.210.460.470.350.460.390.340.480.390.560.340.320.560.370.20.220.611.240.740.50.58
genus

species
1204543
0.380.330.130.290.340.210.220.220.170.30.240.370.290.20.430.220.140.130.450.990.540.370.38
2015412142488710121872862316103980413328

0.220.280.160.320.240.280.480.360.360.360.30.360.120.220.260.30.120.160.320.480.380.260.4
2.53132.584.53.53.52.53.54.50.57.50.58336.59.575.57.5
no rank
1857891
0.110.140.080.160.120.140.240.180.180.180.150.180.060.110.130.150.060.080.160.240.190.130.2

0.110.140.080.160.120.140.240.180.180.180.150.180.060.110.130.150.060.080.160.240.190.130.2
5626516977579115116661319141115
1200862
species

6.489954.781484.473265.235435.4627110.38455.553526.014944.196213.296187.053115.674154.50655141.1253.57569321.867265.092152.9514.63539.6540412.056816.4598.93814
25.8822.7925.0625.2319.7121.0422.3324.7621.6622.2524.2920.9316.0326.0721.4734.7935.2729.8328.3526.3923.430000000000126.1924.25
no rank
12.7911.3212.4512.489.8410.3711.1112.4110.6910.9512.0210.377.9712.9510.6917.1817.5214.8613.9913.1211.5913.0112.01
196896

species
1582156
0.020.010.030.020.010.040.010.020.020.030.030.0700.020.0100
10000011010002033501000

0.060.020.020.050.020.010.02
03001001200100012
114777
species

00110100000108001201201
0.010.010.030.020.010.010.010.020.020.010.0600.010.030.010.030.02
363555
species

0.290.170.570.410.10.250.120.40.20.180.280.260.060.10.180.020.120.080.240.280.220.840.41
1571717429516851413114221462122167730
species
687803

1482001001010022022034021
0.260.170.060.010.020.020.160.020.020.030.050.020.01
species
1458843

species
665032
02020400042001021002
0.040.050.030.120.040.010.020.010.02

4021130010000001002211
0.070.060.020.020.030.0200.010.020.020.010.01
551895
species

1837830
species
0.050.040.050.050.070.010.020.01000.070.030.040.040.010.030.010.06
2212210000000013531215

0000100000000000000001
0.010.010.030.010.010.020.0100.010.01
species
1582155

species
1341101
0.010.01
00000000000000000000000

1445859
species
00111213000100000301
0.030.020.020.020.020.070.020.040.01

111505532012021312102441
0.020.040.040.140.010.050.140.080.070.030.030.040.010.020.10.030.020.040.120.030.060.050.02
1837867
species

1673065
species
0.020.060.030.020.010.03
012003000010000001002

0.010.040.020.010.020.030.020.010.010.060.010.0100.040.020.010.03
02010111000007011003112
species
1416334

00000000000000001
0.010.010.010.01
species
55884

1537091
species
20000100000000000011021
0.040.010.010.010.020.01

941058
species
21012202212201126004641
0.040.020.020.050.020.050.050.030.040.040.010.070.020.050.050.080.040.01

1852625
species
13562702001104010023173
0.020.080.160.150.050.070.020.070.010.030.030.030.010.030.050.010.080.04

20110300015602101015301
0.040.030.020.030.030.10.120.010.070.010.010.060.040.01
1262513
species

0.110.080.140.050.030.070.120.040.020.140.030.110.120.020.10.050.040.08
64420300342101903131028446
1229753
species

00000000000000000000002
0.010.010.010.03
species
1542090

species
1348912
00110400211101001053100
0.020.010.030.030.040.020.020.060.060.040.030.010.020.010.060.050.010.010.01

species
1161931
0.020.01
000011

0.01
0000000000000001
1647466
species

species
536444
0.040.020.050.040.070.060.050.060.010.030.010.010.060.090.05
2100242201300401010575

species
1187128
000003000000000000003
0.0300.04

1582157
species
00.010.010.010.030.020.010.010.010.020.040.010.030.070.010.020.010.010.01
00000011000005013501100

species
1654891
31120310002101030000013
0.070.030.040.0700.030.030.010.030.040.020.010.010.030.010.010.010.010.010.010.05

0.04
00000000000000004
279530
species

0.030.010.070.010.050.030.01
0010010000000011043001
1204521
species

species
1605379
301244015333070181473674
0.060.030.050.10.040.040.120.090.060.060.050.010.070.170.080.040.080.080.05

699370
species
11133954144314021044143
0.030.030.050.070.080.080.120.10.040.140.090.060.040.030.030.020.010.010.050.050.020.040.05

species
1458862
0.020.020.01
00000000000100020001

0.020.060.010.01
00010000020000010001
species
1864622

1883372
species
209601000012010310110011
0.040.290.140.010.020.040.010.280.010.0100.010.020.01

00000000100000000000100
0.020.0100.010.01
species
1127518

1655305
species
10010100010012131201103
0.030.010.040.020.010.020.020.0500.010.070.020.070.030.020.040.020.010.010.05

00000051101004012001211
00.010.0100.120.020.030.020.030.010.020.010.010.020.030.020.02
species
1582154

species
1868654
141016051011120004001001
0.040.080.340.390.010.050.030.020.050.050.030.040.0400.0200.010.01

1883370
species
5314510211658412515311487818
0.090.070.050.10.120.090.540.410.120.260.090.030.090.040.070.050.030.010.160.110.090.10.25

species
1437363
100111100100071333022003
0.020.010.020.030.010.040.010.030.010.050.070.30.0300.030.030.010.010.04

species
1161927
0.040.040.030.020.10.030.050.020.020.060.030.070.010.030.010.10.020.010.030.15
221143211000141131921311

species
1279082
00010102212003111043001
0.020.010.050.050.030.040.020.070.010.010.050.040.01

100031200001011000043233
0.190.010.080.030.020.0100.030.010.040.0100.050.050.030.030.04
species
754038

1542092
species
00.030.0100
000000000000000200000

species
1542477
250413220411711379834657
0.040.10.090.020.030.050.050.120.020.020.280.080.210.060.080.10.030.050.080.050.09

0.020.020.020.020.030.030.490.040.070.050.130.010.010.050.030.05
110000110101125151611425
1206110
species

species
1560342
13610192131053494226141181588159
0.250.130.340.460.060.110.250.120.080.120.180.080.10.190.120.040.10.10.170.10.110.160.13

0.030.020.010.020.020.020.010.010.01
00100000100003002200110
10689
species

1477406
species
0.020.030.020.020.050.020.020.020.030.010.010.030.010.040.0700.050.02
11011601101001003035051

522111012311004003290723
0.10.040.060.020.020.090.020.050.070.030.020.030.030.030.020.110.090.020.04
445683
species

1150991
species
1264781121401472151445251115615
0.220.140.130.160.190.090.050.020.090.270.140.080.110.070.040.030.060.290.140.190.070.2

1564096
species
0.020.0400.060.050.080.060.020.030.010.010.04
10001000022409023101003

0.050.020.030.070.110.050.070.060.020.010.060.160.020.010.050.010.030.030.050.06
201031323200343010412244
species
445685

species
1007869
0.040.030.020.090.020.120.030.110.060.040.020.280.020.030.090.030.01
02101100151631342300721

0.190.160.210.060.150.140.060.090.010.20.170.160.040.030.010.080.020.160.040.050.20.05
1076261623069814011011434184
445688
species

1311613733598628202810552426593442
0.240.230.220.310.180.290.130.240.190.210.540.420.10.060.100.040.070.280.330.770.380.56
1458860
species

331278
species
11165624391171520181834621714122641223019
0.210.340.180.160.570.340.270.190.350.620.340.370.120.330.140.160.120.150.30.520.290.330.25

species
445700
0.020.020.020.020.030.020.030.070.010.080.010.010.020.03
110113010001001110110102

species
1458859
0.240.270.220.290.210.270.110.260.210.210.520.380.10.050.10.120.030.070.260.340.770.360.45
131361293141096271927113452227593334

9357241110420130210233121
0.170.060.160.170.050.040.020.020.040.080.050.10.020.010.010.020.040.040.130.02
1647411
species

570111074001202648071033
0.090.150.020.020.090.170.10.020.040.010.410.040.070.080.010.030.04
species
1136534

1200000153210101216151243
0.020.040.020.120.090.040.020.010.010.020.010.070.190.160.040.04
1567484
species

1000230010000100017000
0.020.010.010.020.050.030.010.010.020.010.010.010.020.010.0300.010.020.090.010.010.01
1486421
species

species
1141136
0.750.50.220.430.590.410.760.510.560.590.770.720.160.360.210.20.170.380.450.450.380.420.39
4024718254831202419403645132220313936293929

925984
species
0.020.010.020.010.02
01000100000100000012

13034601022222023092435
0.020.060.070.110.060.010.020.060.040.040.080.010.020.030.110.020.050.030.07
1589733
species

0.170.070.030.050.070.020.040.080.120.060.040.040.070.020.10.090.050.020.040.010.02
931232001263151212741311
species
1608310

species
1883371
6457213211623612172729411610916
0.120.10.180.170.050.110.060.290.140.080.070.120.050.011.180.240.240.060.130.090.140.110.22

1555202
species
1848781869321053721615141318142518
0.340.080.270.160.20.150.150.230.070.080.190.110.120.050.170.150.130.180.150.230.180.280.24

910031910123304112431113763
0.170.210.080.020.080.020.010.020.060.060.060.030.070.110.030.040.130.170.10.070.04
1654919
species

0.150.10.130.060.060.120.160.020.010.080.080.070.060.030.120.030.020.090.030.04
854207561044010063922723
1481186
species

00000100000
0.010.010.010.01
1654884
species

0.020.050.020.040.040.010.10.0300.010.01
00001002102012100020001
1204535
species

species
1647463
00000000000000001000101
0.010.010.01

0002150401030000232334
0.050.020.040.10.030.060.020.040.020.040.040.04
species
755164

0.020.020.0200
000102000010000000
1127514
species

0.220.10.250.140.230.250.210.190.40.190.20.240.060.410.050.240.060.50.30.240.130.29
1247692987176101207652844424181121
1498168
species

33002802201002008320411
0.060.060.050.070.050.050.020.010.070.040.020.050.010.01
1527770
species

20120.7777783.711011133.3333300200.66666721.754.68752
0.150.020.110.160.090.10.10.070.060.130.090.10.360.090.020.060.030.030.110.080.160.1
1873999
0.10.010.070.110.070.070.070.050.040.090.060.070.240.060.010.040.020.020.070.060.110.07
species

no rank
1690431
20120311011134002002142
0.050.010.040.050.020.030.030.020.020.040.030.030.120.030.010.020.010.010.040.020.050.03

1429794
species
9205131432231506101186116
0.170.040.120.020.030.020.10.070.060.040.060.040.040.050.010.130.10.080.120.08

0.080.050.080.050.040.070.080.030.020.050.050.030.010.030.010.020.020.050.140.050.050.06
422218310121030022411344
species
1852564

21010300100105022624211
0.040.020.020.030.020.020.040.020.020.070.020.050.030.010.01
889956
species

6144353221781251131131114122141614
0.110.290.150.080.130.280.050.030.160.240.230.10.040.090.070.120.10.180.140.270.050.180.19
species
1597976

0.050.070.120.120.080.120.10.270.120.080.210.320.080.080.070.010.030.030.080.080.040.070.18
23353134105211162111142663613
species
1815522

0.040.020.020.010.020.050.060.020.070.140.010.030.020.010.070.020.03
210101102210010213021522
536466
species

species
1486413
0.020.010.010.040.020.010.040.010.030.070.010.020.020.020.0100.010.050.010.010.010
10011110120102000040010

1498228
species
0011000101203000000001
0.030.020.020.030.040.120.01

species
0.10.130.210.060.080.130.150.060.190.10.070.10.010.130.070.350.070.080.310.150.20.150.18
756278
0.20.260.420.120.160.260.30.120.380.20.130.20.020.260.140.70.140.160.620.290.40.30.33
2.53311.57.53141.51.384622.508.50.5194313.55.310347.56.56.09091

0.10.130.210.060.080.130.150.060.190.10.060.10.010.130.070.350.070.080.310.140.20.150.15
5662315628335017138862711151311
869724
no rank

species
1236000
0.010.01
0000000000000201

23211020527300000006011
0.040.060.060.020.020.050.130.060.130.0600.070.010.020.01
species
1357715

11000201011203000101
0.020.020.020.020.030.020.040.020.010.01
species
929833

0.040.040.060.050.030.020.020.050.020.010.020.020.010.01
2222031120100000122101
species
1399915

species
1229760
0.020.010.040.010.020.020.010.01
0000110000001202000211

0000000001
0.03
1451050
species

8733610333303022092149351013
0.150.150.110.080.150.090.070.090.080.10.070.160.080.180.060.110.040.060.110.17
species
445686

species
694060
03000400103103011091100
0.010.070.010.010.010.040.020.020.060.020.020.020.010.010.010.110.020.020.010.01

76212417011410501010119214725
0.140.140.70.570.040.060.020.030.040.130.20.120.010.010.010.020.230.260.050.790.08
species
1883373

00000000000001000000002
0.020.010.03
species
1567486

610115552142290313971034
0.110.020.020.020.040.120.120.050.030.080.040.080.060.030.010.040.10.090.130.030.05
species
979525

573175
species
00000000000000000003
0.04

species
230158
0.030.020.010.020.020.040.050.050.010.060.010.010.020.050.080.04
00101111002007017112473

1035662
species
12000010000000000000013
0.020.040.020.010.04

536453
species
4013284100010040120165442
0.070.010.030.080.050.070.110.250.030.030.110.010.070.060.050.050.03

species
1204534
0.020.0100.050.080.010.010.030.030.020.01
100000002000011000002220

1636182
species
0100030052210100112002
0.020.030.120.060.040.020.010.010.010.020.02

0.020.040.020.030.020.020.090.020.010.010.01
12001401130100000001011
species
947843

407392
species
0.050.010.050.050.130.090.10.120.040.050.10.060.080.030.030.030.030.030.020.050.010.050.1
201251045115324033224057

3023283310531819141327249699211024135983362383848
0.570.490.910.770.240.450.440.470.330.410.510.480.44.980.199.973.477.330.380.770.50.420.65
species
1335230

0.020.020.020.010.010.030.020.010.050.020.020.010.030.040.01
10010000011002052212031
species
1445811

1852628
species
6471062111345917110542315212820
0.110.090.240.230.140.180.020.020.070.120.10.190.040.050.070.090.040.050.260.20.280.310.27

1527519
species
1064836641071170353628201214159
0.190.130.130.190.070.050.160.110.230.230.220.150.250.210.050.020.10.230.150.190.170.12

8713162742146453603125272112715
0.160.160.430.390.060.070.10.060.330.190.080.110.120.050.030.010.310.320.270.160.080.21
species
754039

species
1458863
0.040.050.10.020.050.030.020.010.020.01
02002041211001000200001

0.060.050.020.020.020.020.020.020.030.040.020.090.01
300202111011040400271
908819
species

species
1129194
0.020.010.010.01
000100000000020010001

species
929832
0.020.050.030.130.070.10.110.040.050.030.10.120.040.070.020.010.030.070.060.090.01
121531241215606120136581

0.260.20.480.340.120.150.070.110.090.180.220.250.120.140.010.090.20.250.10.750.15
1491514518343611120162107182076911
species
444860

species
1327935
55188235033661181991122181197
0.090.10.050.190.190.20.120.070.090.110.120.040.130.070.080.080.130.250.220.140.10.09

species
1195085
0001020003100021111110
0.020.020.090.020.140.010.010.010.010.020.020.01

0.020.050.020.020.030.020.060.040.020.030.030.01
01021001011310023030
1206545
species

1654883
species
0.240.320.210.240.120.180.230.330.220.220.350.220.080.030.140.090.090.030.140.20.120.120.15
131561052091397181124210102111691011

species
103807
0.010.0100.01
0000010000000000000001

1500713
species
0.040.020.010.010.010.040.03
20000000100001000011302

3841483441551903151116348
0.060.180.130.020.10.070.070.10.090.030.10.10.040.060.030.130.010.130.080.050.040.11
754075
species

0.150.150.120.110.070.180.070.120.210.190.380.170.240.150.050.010.020.090.210.340.060.17
77343212596208621051181625512
species
754037

species
1883377
0.030.020.020.010.020.030.010.010.08
00100200000101024101006

0.040.010.010.010.040.010.020.020.01
0010010010010000100001
66284
species

1357714
species
3614410230202331732782511
0.060.130.030.090.10.090.050.070.060.040.120.020.070.060.030.020.080.10.030.050.15

1124849
species
232410276723108100142990228
0.040.060.060.090.240.230.150.180.050.090.190.160.040.010.030.020.110.120.010.240.11

species
948870
5466712923569451803180831571312
0.10.090.20.160.170.110.230.590.120.180.180.080.20.130.030.150.950.190.10.150.16

1147094
species
7912714437203716110151516136189
0.140.210.050.050.170.120.10.070.160.060.010.070.280.120.070.090.130.190.180.160.080.20.13

1283076
species
0.070.020.030.10.060.020.070.030.040.020.030.030.030.060.110.030.070.07
41104701312104034059265

0.140.040.280.20.120.140.20.160.140.080.020.040.080.180.080.060.020.16
20.50306011.51.521.50.50.50011.53.51.510.52.5
0.070.020.140.10.060.070.10.080.070.040.010.020.040.090.040.030.010.08
687800
species

869662
no rank
410601202334311002373215
0.070.020.140.10.060.070.10.080.070.040.010.020.040.090.040.030.010.08

species
683735
0.040.040.0100.030.050.010.010.010.01
20010000010000000001100

species
984186
20000000000100000000101
0.040.020.010.01

0.030.020.020.010.040.01
00111100000200001
1505227
species

species
1407493
0.020.010.050.010.010.020.010.020.020.010.03
1002010010000000100212

1434323
species
0.130.170.030.0400.040.060.110.020.040.050.030.010.040.060.020.020.040.040.030.020.040.06
68010424012105011332134

species
1640885
0.060.030.010.120.080.050.030.020.020.020.030.010.110.070.060.070.04
31005900210103000095463

03210303201105054701023
0.020.080.070.040.010.030.020.090.070.010.030.030.030.040.050.030.090.010.020.010.030.04
1486658
species

0.10.210.070.10.080.170.140.210.070.110.190.160.030.070.080.060.070.020.120.150.180.10.22
592431958231080101781101213916
species
1891703

species
445696
0.070.040.030.020.020.030.070.020.090.010.0600.030.040.020.010.010.05
42111303130009003302014

45163142514753215291251313276
0.070.10.030.150.080.120.050.120.330.210.10.060.080.110.160.080.10.070.150.160.030.080.08
1026955
species

458639
species
0.020.030.020.020.020.020.060.020.040.030.070.020.070.060.050.070.010.040.11
10101201103114128546148

0.340.320.350.120.140.170.20.370.40.20.310.220.130.070.050.030.010.320.170.210.080.25
181511561981517616110181531281416719
species
546805

0102011141200002100304
0.020.050.010.020.020.090.030.040.020.010.040.04
148943
species

1873990
species
31100000010003013122010
0.070.040.040.0200.030.010.020.020.020.010.030.020.030.030.010

1486422
species
3111067454341603107719102
0.060.030.040.040.010.060.190.110.130.130.060.080.050.050.040.080.090.090.010.120.120.03

species
12336
4030142042663443181632252080155196720108467710044
0.750.630.450.471.010.560.841.060.420.480.620.510.810.581.040.468.110.251.230.571.011.090.59

1582153
species
0.010.010.010.010.010.010.010.02000.010.01
0000000000000000000000

0.030.020.030.070.060.040.070.030.010.010.160.050.050.01
00101303020200131114441
species
1173297

522338101000060132794116
0.10.050.080.080.080.070.040.030.010.010.010.050.010.030.020.090.110.060.120.09
445684
species

24353618206233010248518888
0.050.090.10.120.070.050.440.510.140.080.060.070.010.020.030.10.060.220.10.090.11
species
1137745

0.020.020.01
000000011000000000001
species
1282994

species
1277886
0.020.030.030.020.060.040.050.040.010.020.040.050.110.020.040.070.090.040.020.03
111025110022000122367312

0000103000000000001003
0.010.010.0200.090.010.0100.010.010.04
1628720
species

889955
species
00010001000000050020112
0.020.020.050.020.010.010.03

1857100
species
000010010000036010011031
00.010.020.010.0300.050.020.010.260.020.010.020.020.040.02

754048
species
00000020102000001003111
0.050.020.040.010.040.010.010.01

02100101
0.040.030.010.02
species
557289

451705
species
1100010100000000301002
0.020.020.010.020.030.010.02

91027618421041422116561611149216
0.170.230.080.160.140.150.10.050.230.120.030.090.080.150.070.590.050.20.130.180.120.230.09
1439894
species

0.030.080.010.010.010.030.010.020.040.030.020.040.040.0500.0100.010.01
1300030011111005001000
species
1868169

20000000010203011
0.040.030.040.020.010.01
species
1127515

0.020.040.020.090.020.040.060.040.040.030.020.020.040.060.010.020.010.060.020.020.060.04
11030421111104012152153
1815590
species

1486660
species
0.040.10.110.020.040.040.060.10.130.040.040.040.010.060.030.030.10.020.020.030.010.03
14301423512208023811211

929834
species
0000020001
0.020.03

756280
species
0.020.060.020.020.020.030.010.030.01
130012010100000100301

1204528
species
011011000000000113001
0.030.030.010.020.010.030.0100.010.010.0400.01

species
1582152
7302277430520121210223125106
0.130.060.020.060.060.060.170.10.080.020.10.040.020.090.10.020.080.030.260.150.060.110.09

00000000000000001000101
0.0100.0100.010.01
1225789
species

12402
species
00011100201134015421341
0.020.020.010.050.020.020.120.030.010.040.050.020.010.040.040.01

000100000000287029337962
0.021.130.50.140.840.311.180.02
1051676
species

0.120.070.220.110.070.020.110.050.260.210.010.060.011.240.020.10.150.030.10.07
63743241117030118200912295
1883374
species

89144873396201213462111947
0.170.190.060.110.110.070.180.080.080.280.120.050.090.090.030.040.080.240.140.120.050.1
1654926
species

1173764
species
0.030.020.050.040.030.010.070.040.030.020.020.020.040.010.010.010.030.030.020.010.010.01
10111121100001001221010

0.030.020.030.020.010.020.190.060.090.020.130.120.10.130.120.220.050.020.040.030.03
111002724001317114141841332
1411095
species

444859
species
0.020.060.130.180.050.090.090.070.070.040.030.060.080.020.070.010.020.010.110.020.160.070.08
1247210333113221121911265

1273738
species
218329171710241773148111981132192113
0.390.170.110.050.230.150.420.260.050.140.320.150.120.10.590.10.160.10.130.40.250.230.18

0.070.150.030.050.050.020.020.050.040.020.040.040.010.020.060.120.10.070.07
471206110121160110510765
444862
species

species
1278479
686962054859131804722321161229
0.130.170.210.210.150.170.130.10.20.160.170.260.060.060.050.040.070.040.260.260.210.130.39

1296659
species
0.030.020.050.010.020.020.0100.120.030.010.01
1000102010000000142010

1127516
species
0.020.020.050.020.020.180.040.03
000100121010000001500042

0.020.050.010.08
0000001020000000001007
species
929814

1458861
species
0.050.030.020.140.03
00000020011000200000002

genus
1298971
0.180.310.360.450.110.230.220.150.140.290.330.180.320.040.110.140.080.020.120.150.160.160.23
0.70.7840912.553.289260.4827591.982761.161290.2195120.4210530.1797751.118280.5576920.739130.2857140.2333331.447370.40909101.178571.166671.309520.8292680.731343
0.50.8811.210.290.580.620.410.380.890.930.520.920.070.30.380.220.050.280.360.420.410.67

species
0.0100.010.010.010.040.010.010000
1987988
0.0200.020.020.020.080.020.020000
0000000000000.5000.5000000

no rank
1080800
0.0100.010.010.010.040.010.010000
0000000000001001000000

0.060.080.020.020.040.020.060.060.0400.060.020.0200.0200.040.06
0.50.50000.500000.500001000000.51
species
0.030.040.010.010.020.010.030.030.0200.030.010.0100.0100.020.03
1987155

11000100001000020000012
0.030.040.010.010.020.010.030.030.0200.030.010.0100.0100.020.03
no rank
1446492

0.040.020.020.080.040.120.040.060.020.020.0200.04
00.50000.500.50.50.50.500000000.500.500.5
species
1987993
0.020.010.010.040.020.060.020.030.010.010.0100.02

no rank
867696
01000101111000000010101
0.020.010.010.040.020.060.020.030.010.010.0100.02

1987156
0.010.010.010.010.020.030.01000.010.01
species
000000.50.50000000000.5000
0.020.020.020.020.040.060.02000.020.02

1406795
no rank
00000110000000001000
0.010.010.010.010.020.030.01000.010.01

0.020.040.020.020.040.060.040.020.120.060.060.0200.020.040.020.040.040.06
00000.51.50000.50.50.500.5000.500.50.50.50.50.5
species
0.010.020.010.010.020.030.020.010.060.030.030.0100.010.020.010.020.020.03
1987157

454798
no rank
0.010.020.010.010.020.030.020.010.060.030.030.0100.010.020.010.020.020.03
00001300011101001011111

0.080.040.060.140.020.060.020.020.020.040.060.040.04
10.50.51.501.50000000000000010.50.5
species
1987158
0.040.020.030.070.010.030.010.010.010.020.030.020.02

no rank
1458849
0.040.020.030.070.010.030.010.010.010.020.030.020.02
21130300000000000000211

1987989
0.020.110.270.240.050.040.140.030.050.040.10.060.030.010.05000.030.030.060.020.05
species
0.040.220.540.480.10.080.280.060.10.080.20.120.060.020.1000.060.060.120.040.1
0.52.545122.50.510.52.51.5000300112.50.52

0.020.110.270.240.050.040.140.030.050.040.10.060.030.010.05000.030.030.060.020.05
158102451215300060022514
no rank
1108865

0.0400.060.020.3200.0400.08
0.50000000000.5020000.5000001.5
0.0200.030.010.1600.0200.04
1987994
species

10000000001040001000003
0.0200.030.010.1600.0200.04
637730
no rank

0.010.010.010.010.030.0100.020.010.01
1987153
species
000000.500000000010.5000
0.020.020.020.020.060.0200.040.020.02

0.010.010.010.010.030.0100.020.010.01
00000100000000021000
1406794
no rank

0.060.040.020.020.060.080.080000.02
00.50000.50000110000000000
1987987
0.030.020.010.010.030.040.040000.01
species

926589
no rank
0.030.020.010.010.030.040.040000.01
0100010000220000000000

000000000.5000000.50000000.50.5
0.0200.020.040.020.040.140.020.020.04
species
1987152
0.0100.010.020.010.020.070.010.010.02

1204523
no rank
0.0100.010.020.010.020.070.010.010.02
00000000100000100000011

1091052
species
11011601101001002035051
0.020.030.020.020.050.020.020.020.020.010.010.020.010.040.0700.050.02

1.052630.9444440.4871790.2857140.54166711.043480.9473680.7169810.4464290.743590.2272731.104171.4062510.4166671.0344814.61391.432431.333333.611111.116281
0.570.360.390.280.480.340.690.570.530.560.390.220.960.320.630.120.292.020.370.451.080.430.32
genus
1922327
0.190.120.130.10.160.120.230.190.170.180.130.080.320.10.210.040.090.680.130.150.360.150.1

0.050.010.040.040.020.030.040.020.040.060.030.030.140.030.010.020.030.010.030.020.040.02
1987728
species
0.10.020.080.080.040.060.080.040.080.120.060.060.280.060.020.040.060.020.060.040.080.04
100.50.501.50.500.50.50.50.51.5200110.510.51.50.5

no rank
1486661
0.050.010.040.040.020.030.040.020.040.060.030.030.140.030.010.020.030.010.030.020.040.02
20110310111134002212131

0.030.080.060.020.060.030.050.050.030.060.010.160.030.020.040.60.010.020.130.040
1922328
species
0.060.160.120.040.120.060.10.10.060.120.020.320.060.040.081.20.020.040.260.080
0.5210.5120110.51.502201224.50.51520

no rank
764595
0.030.080.060.020.060.030.050.050.030.060.010.160.030.020.040.60.010.020.130.040
1421240221304402449121040

0.020.010.010.010.020.080.010.010.020.010.010.010.020.030.020.030.010.03
1987727
species
0.040.020.020.020.040.160.020.020.040.020.020.020.040.060.040.060.020.06
0.5000011.50000000.5000.50.51.50.5101

1136731
no rank
10000230000001001131202
0.020.010.010.010.020.080.010.010.020.010.010.010.020.030.020.030.010.03

2.50.50.50.511.5221.5110.502.51.50.51.5133722
0.180.040.060.040.140.060.220.220.160.160.080.040.040.080.420.020.060.040.140.160.360.10.12
0.090.020.030.020.070.030.110.110.080.080.040.020.020.040.210.010.030.020.070.080.180.050.06
1987729
species

no rank
1486659
0.090.020.030.020.070.030.110.110.080.080.040.020.020.040.210.010.030.020.070.080.180.050.06
511123443221053132661444

0.140.310.160.060.10.060.040.090.120.020.180.080.030.030.160.090.040.06
00.7142860.9677420.7500.8333330.20.3333330.500.8888891.500.502.444441.250.3333330.3333331.3750.77777810.333333
1198980
0.060.130.070.030.040.040.020.050.060.010.080.030.010.010.070.040.020.03
genus

species
2006179
0.01
0000000
0.02

538398
no rank
0.01
0000000

417280
species
0.040.10.050.010.020.020.040.060.010.060.030.050.030.020.01
02320101102301073004221

species
2006172
0.0200.01
0.0400.02
0000.50000000000000000.5

0.0200.01
00010000000000000001
no rank
763998

2006170
0.020.01
species
00.5000000000000000000000.5
0.040.02

01000000000000000000001
0.020.01
no rank
1640974

0.020.040.020.020.020.02
000000.50.50000000000.500.50.50.5
species
0.010.020.010.010.010.01
2006169

0.010.020.010.010.010.01
000001100000000010111
no rank
1640970

2006175
0.02
species
0.04
000

000
0.02
no rank
347329

0000000000000000.500.5
00.020.02
species
2006178
00.010.01

00.010.01
000000000000000101
no rank
549446

00.01
2006180
species
0000000000000000.5
00.02

no rank
582382
00.01
0000000000000001

2006177
0.02
species
000
0.04

0.02
000
549445
no rank

0
000000
no rank
1737463
0

species
1158680
000000
0

000000
0
1548907
species

subfamily
0.760.450.320.440.950.480.490.590.560.330.760.583.560.460.350.240.370.231.390.570.570.630.89
1636616
2.431.451.041.442.991.531.611.821.751.092.511.8610.741.491.140.751.180.724.371.881.82.072.79
2.213990.8620691.057690.9236112.341142.875820.8385091.879122.302860.5229361.756971.3924723.03723.17450.1754390.962.661020.98611113.27692.292552.72.753624.4086

genus
1636617
0.50.220.190.260.690.290.250.520.520.180.320.323.320.20.070.120.170.121.10.280.360.290.61
4.51.720932.51.843144.627745.79311.583.269233.846151.108113.109372.9076937.15965.128210.51.782614.058821.6825.40644.228075.643845.103458.69672
10.430.380.511.370.580.51.041.040.370.640.656.640.390.140.230.340.252.190.570.730.581.22

11402912485323280442127459
0.210.080.050.230.110.10.210.120.110.050.070.10.060.040.040.030.140.090.060.060.13
722418
species

7253762510178781414114161181424
0.130.050.190.080.170.060.060.140.250.030.140.183.160.10.070.040.10.060.190.140.110.160.32
1190451
species

8405111436617314034367415611
0.160.080.120.280.120.090.170.150.050.130.080.060.030.030.030.040.760.060.20.070.16
species
722417

1.313430.8947370.4411760.7959180.9850752.212770.9032260.1052630.20.4102562.180181.111111.259264.265620.21.107143.744681.208333.012662.567571.483.186052.48529
0.670.570.340.490.670.470.620.190.150.391.110.630.540.640.650.280.470.240.790.740.50.860.68
genus
1636618
0.260.230.130.190.260.190.250.070.050.150.440.260.240.260.280.120.20.110.30.280.210.340.28

species
934027
22001420004206032244362
0.050.050.020.020.040.040.050.020.010.020.090.050.030.050.060.030.020.020.050.050.040.070.03

species
0.090.070.060.090.110.050.080.020.030.080.150.080.020.060.030.020.050.010.140.130.050.10.09
1980966
0.180.140.120.180.220.10.160.040.060.160.30.160.040.120.060.040.10.020.280.260.10.20.18
21.511.5231.50.50.514204012.50.5651.543.5

no rank
1229752
4323463112840802511210387
0.090.070.060.090.110.050.080.020.030.080.150.080.020.060.030.020.050.010.140.130.050.10.09

species
1110702
42122720005341513134545108
0.080.050.030.060.070.060.060.020.010.020.10.060.170.110.130.030.110.050.060.050.070.110.11

0.050.050.020.020.040.040.050.020.010.020.090.050.030.050.060.030.020.020.050.050.040.070.04
1909388
no rank
11000.521000210301.511221.531.5
0.10.10.040.040.080.080.10.040.020.040.180.10.060.10.120.060.040.040.10.10.080.140.08

species
1541686
22001420004206032244363
0.050.050.020.020.040.040.050.020.010.020.090.050.030.050.060.030.020.020.050.050.040.070.04

genus
0.110.010.150.130.020.140.080.170.040.190.080.140.050.150.110.20.110.170.090.050.09
1921407
0.310.010.390.290.020.360.180.450.080.490.160.340.150.450.310.580.230.430.250.130.21
0.32258100.3846150.72413802.0277800.6444440.250.5102040.50.82352900.733333031.677422.310342.304351.860470.680.3076920.857143

0.060.020.04
00000000.500000000.500.5
species
1921408
0.030.010.02

1231048
no rank
000000010000000101
0.030.010.02

1921411
0.020.030.0200.030.020.030.060.010.040.020.110.050.120.030.050.020.010.03
species
0.5000.501.500.500.50101062.54.511.50.50.51
0.040.060.0400.060.040.060.120.020.080.040.220.10.240.060.10.040.020.06

743813
no rank
100103010102020125923112
0.020.030.0200.030.020.030.060.010.040.020.110.050.120.030.050.020.010.03

0.5000.501000000.500.500.50.50.50.50.50.50.50.5
0.060.060.0400.040.020.040.060.020.040.020.020.020.040.020.040.040.020.02
1921413
0.030.030.0200.020.010.020.030.010.020.010.010.010.020.010.020.020.010.01
species

685892
no rank
0.030.030.0200.020.010.020.030.010.020.010.010.010.020.010.020.020.010.01
10010200000101011111111

0.060.060.0400.060.020.060.060.020.040.020.020.040.020.020.060.040.020.02
0.5000.501.500.50000.500.5000.500.510.50.50.5
species
0.030.030.0200.030.010.030.030.010.020.010.010.020.010.010.030.020.010.01
1921412

0.030.030.0200.030.010.030.030.010.020.010.010.020.010.010.030.020.010.01
10010301000101001012111
no rank
1327966

0.040.060.0400.060.020.060.040.060.020.040.020.020.040.040.020.060.040.020.02
0.5000.501.500.50.5000.50000.50.50.50.510.50.50.5
species
1921410
0.020.030.0200.030.010.030.020.030.010.020.010.010.020.020.010.030.020.010.01

0.020.030.0200.030.010.030.020.030.010.020.010.010.020.020.010.030.020.010.01
10010301100100011112111
no rank
757342

0.120.070.180.130.360.060.060.220.090.120.180.030.070.060.020.130.08
0.33333300.3333330.1538460.6666670.33333300.36363600000.333333000.5555560.33333300.33333300.30769200.25
genus
0.040.020.060.050.120.020.020.070.030.040.060.010.030.020.010.050.03
1925779

0.0400.040.020.020
0000.50000000000010.50000
0.0200.020.010.010
1925780
species

no rank
1548901
0.0200.020.010.010
000100000000000210000

0.500.5010.500.53333300000.5000000.500.500.4
0.040.030.120.040.240.040.040.150.040.080.0200.040.010.080.05
species
1925782
0.020.010.060.020.120.020.020.070.020.040.0100.020.010.040.03

0.020.010.030.010.060.010.010.040.0100.0100.020.01
10102101000000000010101
1327964
no rank

1327965
no rank
00102101000010000010101
0.010.030.010.060.010.010.040.010.040.010.0100.020.01

00000000000000000000
00
981431
no rank

0.020.010.010.030.0100
1925781
species
0.500000000000000100000
0.040.020.020.060.0200

582345
no rank
0.020.010.010.030.0100
100000000000000200000

0.030.030.0600.270.030.0300.030.030.12
00000.33333301000000000000.333333001
1915205
0.010.010.0200.090.010.0100.010.010.04
genus

0.020.020.0400.180.020.0200.020.020.08
00000.501.5000000000000.5001.5
species
1915206
0.010.010.0200.090.010.0100.010.010.04

no rank
1505225
0000103000000000001003
0.010.010.0200.090.010.0100.010.010.04

genus
0.370.610.340.30.40.520.50.590.320.410.470.440.090.110.260.180.160.070.340.350.90.320.59
680114
1.316832.478260.3645830.7439021.247714.867651.385711.891030.7362640.6239321.927421.4833301.7692301.31253.055560.252.206522.566675.9241.880433.1227
1.011.610.960.821.091.361.41.560.911.171.241.20.220.260.660.480.360.20.920.92.50.921.63

2006134
0.180.260.160.190.190.190.30.280.170.220.190.220.020.070.060.070.060.030.170.190.580.20.27
species
0.280.340.280.320.290.220.480.350.290.380.230.320.030.090.060.080.080.030.260.271.040.360.42
2.857146.882350.6785711.43752.9655213.954535.714291.586211.263165.956523.87504.8888904.52.37514.461547.222228.942313.972227.45238

no rank
1340712
00010101001201000023155
0.010.010.020.030.010.010.020.020.030.020.050.010000.030.040.020.060.07

1327936
no rank
0000200000000001100001
0.050.010.010.01

000000000000
0.010.01
205877
no rank

no rank
546807
0
00000000000000000

0.020.020.010.010.0100.040.010.040.02000000.140
10000010100000000000110
1074307
no rank

0.02
1
no rank
1079895

0.020.01
0001000000
no rank
1327762

0.020.020.020.020.0100.050.010.040.020000.0100.140.01
10000020100000000000110
373404
no rank

1327938
no rank
00110101001101000023155
0.010.010.030.030.010.010.020.020.030.020.040.010000.030.040.020.060.07

100001201100000000101110
0.020.020.020.020.010.010.050.010.040.04000.01000.020.140.020.01
663554
no rank

0000000000000000000000
0.010.010.010
no rank
546803

species
2006136
0.060.120.060.030.070.090.050.080.030.050.080.060.010.010.040.040.010.020.030.040.030.040.08
1.52.50.50.51.550.51.50.50.521.500.501.50.50.511.511.53
0.120.240.120.060.140.180.10.160.060.10.160.120.020.020.080.080.020.040.060.080.060.080.16

0.060.120.060.030.070.090.050.080.030.050.080.060.010.010.040.040.010.020.030.040.030.040.08
351131013114301031123236
1698710
no rank

000001.500.5000.50.50000.51.50110.501.5
0.020.0200.060.020.080.020.020.040.060.0200.020.040.060.020.060.060.060.020.08
species
0.010.0100.030.010.040.010.010.020.030.0100.010.020.030.010.030.030.030.010.04
2006138

no rank
1088866
00000301001100013022103
0.010.0100.030.010.040.010.010.020.030.0100.010.020.030.010.030.030.030.010.04

0.030.070.050.020.050.080.050.090.040.070.080.060.010.010.060.020.010.010.030.030.030.030.08
2006133
species
0.060.140.10.040.10.160.10.180.080.140.160.120.020.020.120.040.020.020.060.060.060.060.16
0.51.50.50.50.54.50.51.50.512100.5010011113

no rank
1034128
13111913124201020022226
0.030.070.050.020.050.080.050.090.040.070.080.060.010.010.060.020.010.010.030.030.030.030.08

species
0.080.140.050.050.080.110.10.10.070.060.090.070.020.010.060.030.010.010.070.040.190.040.11
2006135
0.160.280.10.10.160.220.20.20.140.120.180.140.040.020.120.060.020.020.140.080.380.080.22
230.511.562210.521.50001.50.502.51.571.54

no rank
1074305
0.080.140.050.050.080.110.10.10.070.060.090.070.020.010.060.030.010.010.070.040.190.040.11
4612312442143000310531438

1912143
0.060.030.050.020.010.030.050.010.020.060.030.030.020.030.040.010.180.040.110.140.03
genus
1.50.50.5001.50.500.510.50.50101.520.57.51.546.51
0.120.060.10.040.020.060.10.020.040.120.060.060.040.060.080.020.360.080.220.280.06

115987
species
3110031012110203411538132
0.060.030.050.020.010.030.050.010.020.060.030.030.020.030.040.010.180.040.110.140.03

0.830.970.960.990.640.710.770.880.620.90.731.020.30.671.380.490.511.0910.7110.550.79
680115
genus
7.90837.778635.264828.649645.4761915.34078.194296.196514.422365.050637.204199.939161.5365917.47657.793111.256413.121418.03415.0951815.88768.5793110.5171
2.292.622.532.741.681.821.752.291.612.371.912.630.821.72.91.171.42.942.631.932.671.452.05

species
169683
0.20.280.350.230.240.310.580.330.240.350.30.430.080.311.240.30.130.330.370.20.330.20.32
101311101036231310111521243183316263216251824

no rank
0.630.690.610.760.40.40.20.540.370.560.440.590.220.360.140.190.380.760.630.510.670.350.47
1231361
1.261.371.221.520.80.80.41.080.751.120.881.180.440.720.280.380.761.521.261.021.340.70.94
12.785712.2197.762314.07898.520.952.59.166676.293337.2510.522714.52.522.7518.6052621.434229.236822.11111225.512.628614.766

species
198110
0.090.10.060.050.020.050.050.050.060.020.020.020.010.020.070.150.040.04
552202222210030212612043

2928173017446201416222954821944604929512832
0.540.580.550.710.40.380.150.490.330.50.420.590.220.340.140.170.370.740.560.360.670.310.43
species
998086

genus
1980938
0.040.040.01
00000000000.6666670.66666700000000
0.120.120.05

species
1980941
0.040.040.01
0.080.080.02
00000000001100000000

0.040.040.01
00000000002200000000
no rank
1264700

0.02
00000000000000000000
species
0.01
1980942

1325372
no rank
0.01
00000000000000000000

20.513.05177.59.51432.51210.637916.31957.58.1578916.5639.3194.514.561.510.2540.853420.115455.309349.271937
1.521.1610.881.341.11.181.161.690.90.761.320.981.161.240.522.080.641.911.042.912.171.98
0.760.580.50.440.670.550.590.580.840.450.380.660.490.580.620.261.040.320.960.521.451.080.99
686598
no rank

41271519286524223415183312809291232383411119974
0.760.560.50.440.670.550.590.560.810.450.340.660.490.570.620.261.040.280.940.511.451.080.99
species
1084719

340435
species
0.020.020.040.040.010.040.010.010.010.01
0100000110200100031100

genus
0.110.220.120.310.310.170.120.110.180.220.160.170.240.080.740.10.240.580.320.250.110.160.23
1921016
0.320.660.360.910.930.510.360.350.560.680.480.530.70.242.220.30.721.760.980.750.330.480.69
0.718751.212120.4722221.450551.376342.666670.5555560.80.9642860.8823530.8750.7547170.6285711.333332.675681.266674.57.06253.704082.093330.9090911.833331.97101

0.040.080.040.140.220.060.040.080.040.060.060.080.220.080.060.040.080.480.080.120.080.160.08
0.5101.52200.5000.50.513012.59.51.521.53.51
species
1921018
0.020.040.020.070.110.030.020.040.020.030.030.040.110.040.030.020.040.240.040.060.040.080.04

207656
no rank
0.020.040.020.070.110.030.020.040.020.030.030.040.110.040.030.020.040.240.040.060.040.080.04
120344010011260251934372

0.010.060.030.090.080.050.050.070.10.060.060.010.020.660.030.160.040.130.10.030.020.11
1921017
species
0.020.120.060.180.160.10.10.140.20.120.120.020.041.320.060.320.080.260.20.060.040.22
010.51.51.52.51011.51.51014.51.591.55.53.510.54

1406785
no rank
0213352023320293183117218
0.010.060.030.090.080.050.050.070.10.060.060.010.020.660.030.160.040.130.10.030.020.11

1.42.5132.55.511.521.51.51.51102.5212.56.53.51.52.52.5
0.150.240.140.280.240.180.10.160.20.20.140.160.220.040.10.10.080.620.320.180.080.120.16
0.080.120.070.140.120.090.050.080.10.10.070.080.110.020.050.050.040.310.160.090.040.060.08
1921019
species

35265112343332205425137355
0.070.120.070.140.120.090.050.080.10.10.070.080.110.020.050.050.040.310.160.090.040.060.08
1497851
no rank

0.480.510.570.180.690.441.231.460.810.720.930.780.180.150.720.270.180.270.390.530.660.270.78
1.583332.098041.122810.6666671.898553.754.178865.575341.851851.333332.989252.461540.2222221.40.5277782.70371.444441.185193.102562.320755.106061.444444.66667
1910931
0.160.170.190.060.230.140.410.480.270.240.310.260.060.050.240.090.060.090.130.170.220.090.26
genus

10.51011.510.53221.50.50.50.50.50.51.50.52.50.51.51.5
0.10.040.120.120.060.10.080.30.240.180.140.080.020.20.020.020.080.020.140.020.060.08
species
0.050.020.060.060.030.050.040.150.120.090.070.040.010.10.010.010.040.010.070.010.030.04
1910936

0.050.020.060.060.030.050.040.150.120.090.070.040.010.10.010.010.040.010.070.010.030.04
21202321644311111315133
no rank
1527469

1910935
0.110.150.130.060.170.120.360.450.120.120.220.190.020.040.140.080.050.050.120.110.210.060.22
species
33.5213.56.5792.525.54.502.514.52.525482.58
0.220.30.260.120.340.240.720.90.240.240.440.380.040.080.280.160.10.10.240.220.420.120.44

no rank
1204533
0.110.150.130.060.170.120.360.450.120.120.220.190.020.040.140.080.050.050.120.110.210.060.22
674271314185411905295410816516

00000000000002.66667000.3333332.6666700.333333001
0.180.030.30.030.12
genus
0.060.010.10.010.04
1918725

1918726
0.04
species
00000000000000000000001.5
0.08

669875
no rank
0.04
00000000000000000000003

0.060.010.10.01
1918727
species
00000000000004000.5400.5
0.120.020.20.02

1541883
no rank
00000000000008001801
0.060.010.10.01

010.3333330.33333300.3333330.5714290.8214290.3076920.3333330.3333330.33333315.256410.33333301.40.33333301.1851900.3333331
0.180.090.060.030.210.280.130.180.060.060.360.390.210.150.030.270.060.12
0.060.030.020.010.070.10.050.060.020.020.120.130.070.050.010.090.020.04
1980922
genus

species
1980926
0.060.020.020.070.020.030.020.020.010.070.010.010.050.01
01.500.5000.51.50.50.50.50.500.50.500.50.50200.5
0.120.040.040.140.040.060.040.040.020.140.020.020.10.02

0301001311110110110401
0.060.020.020.070.020.030.020.020.010.070.010.010.050.01
1689679
no rank

1980925
0.030.010.050.020.020.030.120.120.040.040.010.04
species
0.060.020.10.040.040.060.240.240.080.080.020.08
000.5000.510.50.50.5001.58.5002.5001.500.51.5

0.030.010.050.020.020.030.120.120.040.040.010.04
001001211100317005003013
no rank
1585840

genus
0.060.150.060.050.080.170.320.050.060.170.080.040.031.110.090.030.070.030.10.030.09
1198138
0.120.330.120.090.170.340.640.090.130.340.160.080.062.210.170.050.150.060.190.060.18
0.8333331.030300.44444401.352943.56.50.4444440.3846151.823530.81250.51.166673.542994.235290.82.333330.8333331.368420.3333331.88889

00000000001000100001
0.010.020.0700.010.01
species
244310

species
446807
0.010.010.020.0200.020.070.010.02
0000010010000300250102

0.020.040.020.020.030.050.10.030.010.48000.010.050.020.06
1201030001510170001415
209052
species

0.040.060.020.020.030.170.320.020.020.050.050.040.550.0800.020.020.010.01
23010471310221089002200
species
260373

0.080.02
010000.5
species
1966265
0.040.01

020001
0.040.01
331627
no rank

0.8151821.279410.7837841.26791.084184.09941.21630.8260870.7891890.987221.745171.935020.4359866.745410.2417582.866922.558333.50542.600541.221431.291971.905922.76206
3.034.763.334.333.925.033.193.453.73.135.185.542.894.361.822.633.64.633.732.82.742.873.11
1911928
0.751.190.851.080.981.260.80.860.930.781.31.390.731.090.460.660.91.160.940.710.690.710.77
subfamily

0.320.540.520.560.360.790.340.310.350.460.660.730.260.640.260.390.50.70.560.310.280.40.49
1914850
genus
0.968752.206251.448052.327381.245456.595741.336540.6526321.076191.683452.848483.557080.41025613.19270.43755.410263.946.30664.392861.747251.829273.408334.95238
0.961.61.541.681.12.351.040.951.051.391.982.190.781.920.81.171.52.121.680.910.821.21.47

species
1914899
0.090.220.20.270.150.250.150.080.120.260.270.310.060.110.070.20.130.20.220.10.120.180.3
0.180.440.40.540.30.50.30.160.240.510.540.620.120.220.140.40.260.40.440.20.240.360.6
2535.5314.531.52.53.9215777.50.57.50.5117.589.544.5811

1162290
no rank
0.090.220.20.270.150.250.150.080.120.250.270.310.060.110.070.20.130.20.220.10.120.180.3
41061162963581415115122151619891622

1980960
0.030.030.040.10.040.120.050.040.030.070.060.020.040.020.030.080.070.050.040.020.030.05
species
0.50.50.520.5700.50.50.51.51.503014.52.521.50.51.51.5
0.060.060.080.20.080.240.10.080.060.140.120.040.080.040.060.160.140.10.080.040.060.1

0.030.030.040.10.040.120.050.040.030.070.060.020.040.020.030.080.070.050.040.020.030.05
111411401113306029543133
1684115
no rank

1.530.51.51.58.51.511.51320.5300.54.50.56.52.5122.5
0.140.280.10.180.140.30.180.140.140.120.260.20.120.080.040.020.160.020.30.140.080.10.14
species
1914898
0.070.140.050.090.070.150.090.070.070.060.130.10.060.040.020.010.080.010.150.070.040.050.07

3613317323264160191135245
0.070.140.050.090.070.150.090.070.070.060.130.10.060.040.020.010.080.010.150.070.040.050.07
no rank
1327934

species
0.060.090.040.090.060.110.060.050.040.050.080.030.020.040.020.020.110.340.110.040.040.050.04
1914897
0.120.180.080.180.120.220.120.10.080.10.160.060.040.080.040.040.220.680.220.080.080.10.08
120.51.51610.50.50.520.50301613.54.51.51.521.5

0.060.090.040.090.060.110.060.050.040.050.080.030.020.040.020.020.110.340.110.040.040.050.04
24132122111410602122793343
no rank
1162295

1.512.5019111.512.55.51281763.512241
0.140.10.360.020.10.30.10.140.160.140.220.460.20.820.280.260.20.180.060.10.10.180.06
1980961
0.070.050.180.010.050.150.050.070.080.070.110.230.10.410.140.130.10.090.030.050.050.090.03
species

3250218223251125621412724482
0.070.050.180.010.050.150.050.070.080.070.110.230.10.410.140.130.10.090.030.050.050.090.03
no rank
1116482

0.730.710.370.651.080.290.690.690.820.370.490.70.720.480.190.390.420.570.620.420.660.570.21
1.479450.845070.3783780.8923081.685191.379310.739130.9130431.097560.4864860.7959180.9857140.6111111.7501.512823.095241.526321.741940.976191.848481.789470.619048
1914852
0.250.230.130.230.360.110.230.230.280.130.170.240.240.160.070.130.140.190.220.140.220.190.07
genus

0.160.140.040.10.340.060.060.120.20.060.120.160.20.060.060.040.20.160.120.120.20.140.02
21.5013.51.50.5120.51.521201632.523.530.5
1914853
0.080.070.020.050.170.030.030.060.10.030.060.080.10.030.030.020.10.080.060.060.10.070.01
species

430273124134240212654761
0.080.070.020.050.170.030.030.060.10.030.060.080.10.030.030.020.10.080.060.060.10.070.01
399183
no rank

species
0.020.030.020.050.020.010.060.030.030.060.010.040.040.050.020.020.060.010.020.020.04
1914855
0.50.5010.50.510.50.51010.530100.52.500.50.51.5
0.040.060.040.10.040.020.120.060.060.120.020.080.080.10.040.040.120.020.040.040.08

0.020.030.020.050.020.010.060.030.030.060.010.040.040.050.020.020.060.010.020.020.04
11021121120216020150113
1131317
no rank

species
1914856
0.020.060.010.020.100.060.020.030.030.030.040.030.020.010.020.030.050.080.020.01
0.040.120.020.040.200.120.040.060.060.060.080.060.040.020.040.060.10.160.040.02
0.5100.520100.50.50.510200.50.50.511.530.50

no rank
1446490
12014020111204011123610
0.020.060.010.020.100.060.020.030.030.030.040.030.020.010.020.030.050.080.020.01

species
1914854
0.120.080.070.090.070.050.080.120.110.060.070.10.050.030.070.030.070.050.020.020.080.01
0.240.160.140.180.140.10.160.240.220.120.140.20.10.060.140.060.140.10.040.040.160.02
31.5121.531.52201.51.51303.51.52.520.50.53.50

0.120.080.070.090.070.050.080.120.110.060.070.10.050.030.070.030.070.050.020.020.080.01
63243634403326073541170
1327956
no rank

genus
1914851
0.190.420.190.30.260.370.220.310.30.190.470.420.220.290.130.130.260.260.170.260.190.130.22
0.591.260.570.920.761.130.660.950.90.591.411.260.660.870.370.410.780.780.490.760.570.390.66
0.7796611.555560.4210531.086961.394744.1769931.684210.9888890.9661022.134751.777780.7575763.724140.2432431.51222.56412.551282.530611.868421.43860.9230771.78788

species
0.040.120.060.110.150.130.180.110.050.080.130.040.050.090.050.060.050.020.080.040.020
1914892
0.080.240.120.220.30.260.360.220.10.160.260.080.10.180.10.120.10.040.160.080.040
12.50.523703.520.5230.530.533.52131.50.50

251461407414616167426310
0.040.120.060.110.150.130.180.110.050.080.130.040.050.090.050.060.050.020.080.040.020
889338
no rank

0.10.140.120.20.120.280.10.140.260.320.120.280.10.060.140.220.240.20.10.10.12
11.51218011241.51.530144.553.5222
species
1914893
0.050.070.060.10.060.140.050.070.130.160.060.140.050.030.070.110.120.10.050.050.06

no rank
1204529
2324216022483360289107444
0.050.070.060.10.060.140.050.070.130.160.060.140.050.030.070.110.120.10.050.050.06

1914894
0.060.110.010.040.020.060.220.080.050.10.10.040.110.030.050.050.10.010.020.080.020.06
species
0.120.220.020.080.040.120.440.160.10.20.20.080.220.060.10.10.20.020.040.160.040.12
1.52.500.50.534.51.5102.52.50.5802.5340130.52

350116932055116056802614
0.060.110.010.040.020.060.220.080.050.10.10.040.110.030.050.050.10.010.020.080.020.06
no rank
1391428

12.50.510.53001.503.5305.500.54.50020.51.53.5
0.10.240.120.120.040.10.020.140.040.260.260.160.020.160.020.10.040.080.2
species
0.050.120.060.060.020.050.010.070.020.130.130.080.010.080.010.050.020.040.1
1914895

0.050.120.060.060.020.050.010.070.020.130.130.080.010.080.010.050.020.040.1
251216003076011019004137
1141135
no rank

genus
1980929
0.020.040.060.050.050.030.020.020.090.060.010.010.020.010.020.010.04
0.3333330.6666670.6666670.6666670.6666671.333330.33333300.33333311000.33333300.3333330.66666700.3333330.6666670.33333301
0.060.120.180.150.150.090.060.060.270.180.030.030.060.030.060.030.12

1984796
0.020.040.060.050.050.030.020.020.090.060.010.010.020.010.020.010.04
species
0.5111120.500.51.51.5000.500.5100.510.501.5
0.040.080.120.10.10.060.040.040.180.120.020.020.040.020.040.020.08

1090134
no rank
0.020.040.060.050.050.030.020.020.090.060.010.010.020.010.020.010.04
12222410133001012012103

family
13.8415.1517.6216.9216.3916.8914.9816.0818.9818.116.0817.0222.1818.6419.5519.5429.6620.5413.8315.1213.0614.5413.16
10699
2.605821.802162.278814.684992.540049.879282.861592.299023.006513.156722.379984.871494.0775211.6061.9759313.8455130.32218.593710.90516.159354.748229.250637.16497
45.690000000000149.8958.140000000000356.760000000000253.570000000000255.829999999999947.8351.100000000000162.990000000000360.170000000000352.950000000000155.170000000000270.300000000000260.449999999999862.310000000000162.409999999999993.260000000000666.380000000000444.9149.199999999999842.1447.720000000000141.16

000000000000000000000.28571400
0.010.010.050.130.080.030.070.010.01
genus
0.010.010.010.050.020.010.030.010.01
1921558

1921563
00.0100
species
00.0200
0000000000000000000000

no rank
1589749
0000000000000000000000
00.0100

00.0100
1921562
species
0000000000000000000000
00.0200

1589752
no rank
0000000000000000000000
00.0100

000000000000000000000.500
000.020.040.0200.0200
species
000.010.020.0100.0100
1921561

00000000000000000000100
000.010.020.0100.0100
no rank
1589755

00.010.020.010.010
1921559
species
00.020.040.020.020
000000000000000000000.500

no rank
1589750
00.010.020.010.010
00000000000000000000100

genus
1982243
0.040.040.10.020.240.10.170.020.30.210.060.10.080.190.070.410.050.10.170.120.110.11
0.3333330.250.5333330.3333331.628571.906251.196080.3333333.755561.384620.5555560.80.3333335.19298011.52891.1333301.56252.392161.666672.242421.29032
0.120.120.30.060.70.320.510.060.90.650.180.30.240.570.191.210.150.320.510.360.330.31

0.5000331.5062.50.51.50.59.5019.5202.5430.52
0.040.020.040.30.120.140.560.340.040.140.080.280.060.70.080.120.20.160.040.1
species
0.020.010.020.150.060.070.280.170.020.070.040.140.030.350.040.060.10.080.020.05
1982246

0.020.010.020.150.060.070.280.170.020.070.040.140.030.350.040.060.10.080.020.05
10006630125131190394058614
no rank
1647387

species
1982245
0.020.030.080.020.080.050.10.020.020.050.040.030.040.050.030.050.010.050.070.040.090.05
0.50.510.51.52.520.50.50.510.50.53030.50231.542
0.040.060.160.040.160.10.20.040.040.10.080.060.080.10.060.10.020.10.140.080.180.1

1647386
no rank
0.020.030.080.020.080.050.10.020.020.050.040.030.040.050.030.050.010.050.070.040.090.05
11213541112116061046384

0.160.020.140.220.020.120.140.20.20.30.170.250.4700.10.140.160.20.08
0.500000.250000.150.40.600.47058800.40.723404000.2857140.251.250
1910976
0.040.010.030.050.010.030.030.050.050.070.050.060.1200.020.030.040.050.02
subfamily

genus
0.010.010.06
1910992
00000000000000.33333300.3333331.16667
0.030.030.18

1911010
0.010.03
species
00000000000000.5002
0.020.06

00000000000001004
0.010.03
no rank
698487

1911008
00.010.03
species
00.020.06
0000000000000000.51.5

00000000000000013
00.010.03
no rank
698486

genus
1910991
0.010.030.050.010.020.030.020.010.010.010.040.0400.020.030.040.02
000000.33333300000000.33333300.50.5000.3636360.33333300
0.010.110.170.010.060.110.060.030.050.030.10.0800.080.110.120.06

species
0.010.010.010.010.0100.020.010.020.030.01
2005694
0.020.020.020.020.0200.040.020.040.060.02
000000.500000000001000.50.500

424949
no rank
0.010.010.010.010.0100.020.010.020.030.01
00000100000000002001100

0.020
00000000000000000
2005699
0.010
species

0.010
00000000000000000
no rank
1115478

species
0
2005697
00000000000000000
0

no rank
1325963
00000000000000000
0

0.040.02
0000000000000000.5
species
2005698
0.020.01

0000000000000001
0.020.01
no rank
1540823

00000000000000.5000
0.040.020
2005701
0.020.010
species

0.020.010
00000000000001000
293644
no rank

0.020
00000000000000000
0.010
2005700
species

00000000000000000
0.010
424944
no rank

0.02
2005696
species
0.04
0000000000000001

0000000000000002
0.02
1340534
no rank

species
2005692
0.010.010
0000000000000000
0.020.020

no rank
424947
0.010.010
0000000000000000

0.040.02
0000000000000000000
species
0.020.01
1981032

1611545
no rank
0.020.01
0000000000000000000

000000.500000000000000.50.500
0.020.020.020.020.020.0200.020.040.020.02
species
2005695
0.010.010.010.010.010.0100.010.020.010.01

00000100000000000001100
0.010.010.010.010.010.0100.010.020.010.01
1163482
no rank

genus
0.040.010.030.040.060.020.020.030.05
1910993
0.120.030.090.120.180.060.060.090.15
0.66666700000.3333330000.3333330.66666710100.666667100001.66667

100000.50000.511.501.5011.500002.5
0.080.020.060.080.120.040.040.060.1
species
1911006
0.040.010.030.040.060.020.020.030.05

no rank
1173749
0.040.010.030.040.060.020.020.030.05
2000010001230302300005

genus
1623293
0.020.020.050.020.020.020.010.010.08
00000.4444440.500000000.5000000001.5
0.040.040.090.040.040.040.020.030.16

species
466051
0.010.010.020.010.010.010.010.010.04
00001100000001000000003

00001100000000000000003
0.010.010.020.010.010.010.010.04
466052
species

genus
0.050.030.040.020.020.050.040.030.020.040.040.030.020.010.020.03
1921709
0.150.090.120.060.060.170.120.110.060.120.120.110.060.030.040.09
0.6666670.22222200.250.3333330.333333000.23529400.2727270.33333301.1666701.33333000.36363600.3333330.250.666667

1921710
0.010.030.010.040.020.010.020.010.040.020.0100
species
0.020.060.020.080.040.020.040.020.080.040.0200
0000.500.5000.5000.50102000.50000

no rank
1406790
00010100100102040010000
0.010.030.010.040.020.010.020.010.040.020.0100

0.050.020.010.020.010.020.020.030.030.020.010.010.010.03
1921712
species
10.5000.50.500000.500200000.500.50.51
0.10.040.020.040.020.040.040.060.060.040.020.020.020.06

no rank
1406792
21001100001004000010112
0.050.020.010.020.010.020.020.030.030.020.010.010.010.03

0000000.33333300.57142900.153846000.333333000000000.333333
0.010.10.060.080.210.080.130.090.030.010.010.03
1921544
0.010.040.020.020.070.020.050.030.010.010.010.01
genus

00.010.010.0100.0100
1921546
species
00.020.020.0200.0200
000000000000000000000

000000000000000000000
00.010.010.0100.0100
1141526
no rank

00.010.010.020.010.010.010
1921547
species
000000000.500000000
00.020.020.040.020.020.020

no rank
1476390
00.010.010.020.010.010.010
00000000100000000

1921548
0.050.020.01
species
00000000100.5000.5
0.10.040.02

0.050.020.01
00000000201001
1132026
no rank

0000000.50000000000000000.5
00.020.040.020.020.020.02000.02
1921545
00.010.020.010.010.010.01000.01
species

00.010.020.010.010.010.01000.01
00000010000000000000001
347325
no rank

0.820.80392201.111110.40.7333330.2222220.4242420.57777800.9677420.3201.6666700000.1666670.30.3333330.50.777778
0.50.510.260.360.30.150.090.330.450.170.620.250.120.0100.120.10.060.20.18
1623299
0.160.170.080.120.10.050.030.110.150.050.220.090.040.0100.040.040.020.060.06
genus

species
0.050.050.030.010.020.010.020.050.020.070.03000.010.0100.020.02
1633150
0.10.10.060.020.040.020.040.10.040.140.06000.020.0200.040.04
110000.500101.50.500000000.500.50.5

no rank
673835
22000100203100000001011
0.050.050.030.010.020.010.020.050.020.070.03000.010.0100.020.02

0.140.10.060.20.120.020.040.040.080.040.120.040.0800.040.0200.060
1.510210.50.500.501.50.502.500000.50.5010
0.070.050.030.10.060.010.020.020.040.020.060.020.0400.020.0100.030
1633151
species

673837
no rank
0.070.050.030.10.060.010.020.020.040.020.060.020.0400.020.0100.030
32042110103105000011020

species
0.050.070.030.010.020.030.010.070.060.020.070.03000.010.010.020.020.04
1633149
0.10.140.060.020.040.060.020.140.120.040.140.06000.020.020.040.040.08
11.50001.501101.50.500000000.50.50.51.5

no rank
673832
23000302203100000001113
0.050.070.030.010.020.030.010.070.060.020.070.03000.010.010.020.020.04

1982088
0.210.220.871.550.590.470.320.260.770.570.080.280.160.090.280.720.311.020.460.310.170.70.11
genus
0.630.662.614.651.771.410.960.782.311.710.240.840.480.270.842.160.933.061.380.930.512.10.33
3.666673.333339228.3333318.33334.333333.33333116.333331.333334.666671.333334.333331.3333326.666712.333327.666713.33338.333334.3333321.33332.66667

0.210.220.871.550.590.470.320.260.770.570.080.280.160.090.280.720.311.020.460.310.170.70.11
1982090
species
5.5513.53312.527.56.5516.59.52726.524018.541.52012.56.5324
0.420.441.743.11.180.940.640.521.541.140.160.560.320.180.561.440.622.040.920.620.341.40.22

0.210.220.871.550.590.470.320.260.770.570.080.280.160.090.280.720.311.020.460.310.170.70.11
111027662555131033194144134803783402513648
1796997
no rank

0.020.010.020.050.030.010.010.01
1982573
genus
0.060.030.060.150.090.030.030.03
0000.33333300.3333330.33333300.66666700000010.33333300000.3333330.333333

species
1982575
0.020.010.020.050.030.010.010.01
0000.500.50.5010000001.50.500000.50.5
0.040.020.040.10.060.020.020.02

00010110200000031000011
0.020.010.020.050.030.010.010.01
1662330
no rank

genus
0.030.060.080.020.040.020.050.030.010.020.010.0100.010.030
1623292
00.50.40.333333010.2500.36363600.25000.66666700.50.5000.57142900
0.060.150.180.040.080.050.110.080.020.030.020.0200.030.070

205879
species
01110010100002011001
0.030.030.020.020.040.010.010.0100.02

0.030.020.010.010.020000.010
1993879
species
000.500000000.500000000000
0.060.040.020.020.040000.020

0.030.020.010.010.020000.010
0010000000100000000000
1052673
no rank

1089119
species
00.040.020.010.020.010.01000.02
00010200000000000001

0.010.010.010.010.040.0100.010.03
00000000000.5000000000000.333333
0.010.010.010.010.020.0100.010.02
1623286
genus

species
389469
0.0200.01
00000000001000000000001

10708
species
0000
00000000000000000000000

species
505292
000000000000000000000
0

505291
species
0000
00000000000000000000000

species
937835
0000
00000000000000000000000

000000000000.33333300.33333300000.3333330000
0.060.060.0300.060.03
genus
1982375
0.020.020.0100.020.01

1982376
0.020.020.0100.020.01
species
0.040.040.0200.040.02
000000000000.500.500000.50000

00000000000101000010000
0.020.020.0100.020.01
1109715
no rank

genus
1623305
0.020.020.060.050.110.10.260.150.060.090.030.120.220.170.830.030.10.020.160.160.110.20.11
0.500.50.2222220.4782610.91.038460.5666670.40.501.1666712.911762.331330.41.30.250.9333330.9032260.523812.307690.55
0.040.040.120.090.230.20.520.30.10.180.070.240.440.341.660.050.20.040.30.310.210.390.2

0000002100000100001200
00.050.020.010.010.020
species
665883

213769
species
0.0200.0200.020.010.01
00000200000000000020111

83129
species
0.0300.010.010.030.030.010.020.010.040.01
00001000000004004011140

665884
species
00101100110100000031141
0.010.030.010.040.010.010.020.030.010.0200.030.020.010.040.01

000001000000275100000
0.010.010.110.060.380.010.010.010.010.01
31532
species

0.010.030.010.040.030.050.050.020.030.010.040.010.030.030.050.010.080.03
00101322110200013033172
63118
species

213775
species
0.010.110.060.3800.010.01000.01
00000000000027501000001

0000010001000000200200
0.010.010.010.010.030.0100.020.0100.020
species
494269

10000121000000000001011
0.020.010.050.020.010.010.010.01
species
287412

0.020.010.080.030.060.010.0700.010.020.030.010.01
00010131000302100120201
1165134
species

1623273
0.030.010.010.030.010.020.020.040.0200.040.020.060.0100.020.01
genus
0.060.020.020.060.020.040.040.080.0200.070.040.120.0200.040.02
000.5000.500000.500.51000.7142860.250.3333330010.5

0010000000100000211
0.0300.010.0200.020.010.01
575610
species

0.01
0000000000000000000
320839
species

species
186152
0.0100.01
0000000000000000000

130478
species
0.010.020.0400.010
0000000000001000100

0000000000000000001
0.010.010.01
215167
species

320837
species
00000100000000000010021
0.010.010.020.010.020.01

0.020.030.020.020.020.030.060.020.010.020.020.01
1993858
genus
0.33333300.3333330.3333330.33333300.333333000.33333310.33333300.333333000.6666670.666667000.333333
0.060.090.060.060.060.090.180.060.030.060.060.03

species
1993859
0.020.030.020.020.020.030.060.020.010.020.020.01
0.500.50.50.500.5000.51.50.500.50011000.5
0.040.060.040.040.040.060.120.040.020.040.040.02

373415
no rank
0.020.030.020.020.020.030.060.020.010.020.020.01
101110100131010022001

0.040.16
00000000000000.250000000.75
genus
1982111
0.010.04

0.010.04
1982112
species
00000000000000.3333330000001
0.030.12

no rank
0.010.04
945083
00000000000000.50000001.5
0.020.08

1391659
no rank
0.010.04
000000000000010000003

00000.51.500000.50415.155606.1041715.53.500.50.50.50
00.020.080.060.070.020.0400.680.450.080.480.540.180.030.040.020.020.02
genus
1623274
00.010.040.030.030.010.0200.340.230.040.240.270.090.010.020.010.010.01

0000000000000001000
000.010.01
species
360398

species
186153
0.020.020.1
00000000001000011

00.010.040.030.020.01000.340.220.040.130.270.090.010.020.010.010.01
00001300000083101431701110
259901
species

0.650.880.591.350.080.660.610.821.231.041.30.980.480.140.190.180.090.160.520.120.20.15
0.40.2727270.2542370.229630.250.696970.2786890.0975610.2439020.1538460.3769230.6632650.750.28571400.5263160.3888890.1111110.1250.5576920.08333330.20.266667
subfamily
1982876
0.180.240.140.350.020.170.160.230.340.280.340.260.120.040.050.050.030.040.130.040.060.06

0.180.060.060.060.360.180.120.0100.050.030.05
0.55555600000.666667000.6666670.3333330.333333000000000.33333300
genus
1982900
0.060.020.020.020.120.060.040.0100.010.010.01

0.5000000010.50.5000000000.500
0.040.020.020.120.060.04000.020.020.02
0.020.010.010.060.030.02000.010.010.01
2006145
species

205874
no rank
1000000021100000000100
0.020.010.010.060.030.0200.010.010.01

0.040.010.020.010.060.030.0200.010.01
1982904
species
0.080.020.040.020.120.060.0400.020.02
1000010010.50.500000000000

2000020021100000000000
0.040.010.020.010.060.030.0200.010.01
no rank
373414

0.6956520.37500.3076920.3333330.975610.4117650.2162160.1481480.3157890.6307690.95384610.444444010.5454550.3333330.3333330.6666670.20.666667
0.230.320.130.520.060.410.340.370.270.190.650.650.360.090.090.110.030.030.240.050.06
genus
1982898
0.070.10.030.180.020.130.120.130.090.070.210.210.120.030.030.030.010.010.080.030.02

0.020.020.020.040.080
000000000010000000000
0.010.010.010.020.040
1983108
species

0.010.010.010.020.040
000000000020000000000
no rank
1698252

0.080.040.020.120.020.120.040.020.020.020.0200.04
100100.510000000000000.501
species
1983107
0.040.020.010.060.010.060.020.010.010.010.0100.02

2002012000000000000102
0.040.020.010.060.010.060.020.010.010.010.0100.02
1429905
no rank

11000100.5000.50.51.50000.50.50.500
0.080.10.020.040.040.020.080.040.040.060.240.020.020.020
species
1983109
0.040.050.010.020.020.010.040.020.020.030.120.010.010.010

220002010011300011100
0.040.050.010.020.020.010.040.020.020.030.120.010.010.010
no rank
1354508

species
0.020.010.020.020.010.02
1983117
00.5000100.5000000000000.5
0.040.020.040.040.020.04

01000201000000000001
0.020.010.020.020.010.02
373406
no rank

species
0.010.010.020.020.020.010.030.01
1983106
0.020.020.040.040.040.020.060.02
0000010.50.500000001.50.5

0.010.010.020.020.020.010.030.01
00000211000000031
1354507
no rank

species
0.020.050.010.010.030.070.070.010.010.01
1983119
0.040.10.020.020.060.140.140.020.020.02
0000.502.50000.51.51.501000000.50.5

no rank
1089137
000105000133020000011
0.020.050.010.010.030.070.070.010.010.01

0.020.040.040.020.020.020.040.080.180.020.08
00000.50.5000.501200.5000001.5
1983118
0.010.020.020.010.010.010.020.040.090.010.04
species

no rank
543152
00001100102401000003
0.010.020.020.010.010.010.020.040.090.010.04

1983110
0.010.020.010.020.030.020.020.010.02
species
0.020.040.020.040.060.040.040.020.04
0000.500.5000.50.50.50.500.5001

0.010.020.010.020.030.020.020.010.02
00010100111101002
1647301
no rank

1982899
0.010.040.0600.040.10.020.030.0100.01000.03
genus
0.010.10.1700.110.290.030.030.0200.01000.05
0000.176471000.272727000.1379310100000.5000000.2

0000.5000000.50000000000000.5
0.020.060.0800.02
species
0.010.030.0400.01
1982917

00010000010000000000001
0.010.030.0400.01
1089122
no rank

00.020.020.040000
00000000000000000000000
species
00.010.010.020000
1982905

no rank
1429903
00000000000000000000000
00.010.010.020000

00000010000000001000000
00.020.010.030.030.0100.0100
species
1089110

species
00.010.010.010.0200
1982918
00.020.020.020.0400
00000000000000000000000

no rank
373413
00000000000000000000000
00.010.010.010.0200

genus
1982897
0.0200.0200.010.090.050.010.020.010.010.010.01
0000000000000000000000
0.0400.0400.020.180.10.020.040.020.020.020.02

1982901
species
0.0200.0200.010.090.050.010.020.010.010.010.01
0000000000000000000000

0.010.220.280.250.020.140.080.140.0300.010.030.120.010.010.04
00.5454550.5357140.4801000.2500.28571400000.333333000.3333331000.75
1623295
0.010.080.10.090.020.070.040.060.0100.010.010.040.010.010.03
genus

00000000000000000000000
0000.02000
0000.01000
1982929
species

no rank
1486403
00000000000000000000000
0000.01000

species
00.0300.010.0200
1986540
00.0500.010.0400
0000.40000000.5000000000000

0000000000000000000000
0
1414748
no rank

0.020.02
00010000001
1675554
no rank

1986538
0000.0100
species
00000000000000000000000
0000.0200

00000000000000000000000
0000.010
1458710
no rank

000000000.50000000000000
0.0400
species
1982926
0.0200

0000000010000000000000
0.0200
1673886
no rank

00.511000000000000.5000.51.5
0.040.120.100.020.020.08
1986350
0.020.060.0500.010.010.04
species

1052671
no rank
0.020.060.0500.010.010.04
01220000000000010013

0.0600
000.500000000
1982927
0.0300
species

no rank
1035482
00100000000
0.0300

0.050.0100.020.0200.01
1982928
species
01000000000.5000000000000
0.10.0100.020.0400.01

no rank
1458725
02000000001000000000000
0.050000.020

no rank
1034143
00000000000000000000000
00000

0.060.040.040.020.040.10.080.080.040.061.060.020.020.02
000.50.50.50.50.5100010.5000.51.521.500.50.500.5
genus
1623300
0.030.020.020.010.020.050.040.040.020.030.530.010.010.01

0000000000000001
0.01
1225794
species

001111120002100134301101
0.030.020.020.010.020.050.040.040.010.030.530.010.010.01
species
235201

1623276
0.010.020.020.040.060.020.010.030.010.01
genus
00.3333330000000.3333330.250.3333330000000.66666700.3333330.6666670.3333330.333333
0.030.060.060.120.180.060.030.090.030.03

0.060.040.02
0000000000.50.5000000000.5
species
1982363
0.030.020.01

0.030.020.01
00000000011000000001
205880
no rank

species
1982364
0.010.010.02
00000000000.5
0.020.020.04

no rank
563121
00000000001
0.010.010.02

1982365
0.020.020.020.020.030.010.01
species
00.50000000.500.500000010010.50.5
0.040.040.040.040.060.020.02

no rank
543153
01000000101000000200211
0.020.020.020.020.030.010.01

0.020.020.01
2005059
genus
0.060.060.03
0000.33333300000000.333333000000000.333333

2005060
0.020.020.01
species
0000.500000000.5000000000.5
0.040.040.02

no rank
1662332
000100000001000000001
0.020.020.01

1922243
0.460.490.190.30.610.410.430.210.330.180.370.270.260.690.210.240.210.660.530.420.270.590.34
genus
1.381.470.590.91.811.231.310.650.990.561.090.810.782.090.610.720.641.981.611.260.831.771.02
43.789121.016952.177784.209947.869923.335881.353852.383841.517862.972482.259261.4743620.63640.3278694.777783.9843810.81827.726715.507943.662659.440684.23529

0.460.40.20.320.520.440.260.220.280.040.360.320.060.340.20.160.240.360.520.440.360.720.26
64.51.53.55.512.52.52304.54011.50.54.56.57.5118.56.516.54.5
species
1922247
0.230.20.10.160.260.220.130.110.140.020.180.160.030.170.10.080.120.180.260.220.180.360.13

1434319
no rank
129371125546098023191315221713339
0.230.20.10.160.260.220.130.110.140.020.180.160.030.170.10.080.120.180.260.220.180.360.13

0.230.290.10.140.340.190.310.110.190.170.180.110.230.530.10.160.10.480.280.20.10.230.21
1922246
species
66.51.537116242.54.52.52.5370.58.55.2105319.51283.510.57.5
0.460.580.20.280.680.380.620.220.380.340.360.220.461.060.20.320.190.960.560.40.20.460.42

0.230.290.10.140.340.190.310.110.190.170.180.110.230.530.10.160.090.480.280.20.10.230.21
121336142212485955741171139241672115
1500386
no rank

0.66666700.3333331.33333010.66666700.33333300.6666670.33333300.66666700.666667000.3333331.333330.3333331.66667
0.120.090.270.090.150.060.120.060.030.060.030.150.030.15
genus
1921696
0.040.030.090.030.050.020.040.020.010.020.010.050.010.05

0.040.030.090.030.050.020.040.020.010.020.010.050.010.05
1921698
species
100.5201.5100.5010.50101000.520.52.5
0.080.060.180.060.10.040.080.040.020.040.020.10.020.1

1229755
no rank
2014032010210202001415
0.040.030.090.030.050.020.040.020.010.020.010.050.010.05

0.080.090.010.150.020.050.120.040.160.070.1500.030.040.040.06
0.500000.33333300000.400.3333330.501.1250.1428570.46666700.333333010.5
186532
0.040.030.010.060.010.030.040.010.050.030.0600.010.010.020.03
genus

36343
species
1000020000100100100002
0.020.020.020.010.010.02

species
31537
1000010000000000020
0.020.010.010.020

no rank
0.030.030.040.010.050.010.040.010.010.01
329156
0000000000000.50.501.6363600.42857100.5000
0.060.060.080.020.110.030.070.020.030.03

0.010.010.040.01
000000000000100001
1862961
species

species
1862959
00000000000000030001
00.030.010.01

species
1862958
00000000000000030
00.030.01

00000000000000000100000
0.010.010.010.010.01
species
1862960

1862962
species
00000000000001000100000
0.010.010.010.010.010.01

0.020.080.030.070.070.030.220.070.210.080.020.530.240.040.320.010.060.070.120.10.04
1918017
genus
0.060.240.090.210.210.110.660.210.630.240.061.570.70.120.980.030.180.210.360.30.12
0.3333330.6666670.3333330.5714290.5714290.9090911.545450.5714292.4444400.50.3333331.452233.8857100.83333310.80610.3333331.166671.1428631.11

0.080.020.040.040.320.080.060.020.040.08
010000.50.50000.5023001.500.5101.5
1918192
0.040.010.020.020.160.040.030.010.020.04
species

1327982
no rank
0.040.010.020.020.160.040.030.010.020.04
0200011000104600301203

species
1918194
0.020.030.050.020.150.020.020.020.160.10.010.010.05
0.040.060.10.040.30.040.040.040.320.20.020.020.1
0.500.510.5030.50.500.502700.500.50002

1327980
no rank
10121061101041401010004
0.020.030.050.020.150.020.020.020.160.10.010.010.05

species
1918193
0.040.020.050.030.050.050.190.040.020.20.090.030.30.050.050.120.010.04
0100.511.5114010.52.5601.517.50224.50.51.5
0.080.040.10.060.10.10.380.080.040.40.180.060.60.10.10.240.020.08

0.040.020.050.030.050.050.190.040.020.20.090.030.30.050.050.120.010.04
0201232280215120335044913
1327976
no rank

0.50.80.1764710.3636360000.5003.388894.8095202.1666700.7500.1666673.650.62500.51
0.040.150.170.110.020.060.240.080.070.360.630.10.120.0800.060.20.080.040.06
genus
1623281
0.020.080.090.060.020.030.110.040.030.180.310.050.060.0400.030.10.040.020.03

205868
species
0.030.01
0010000000000000001

00.0200.010.010.010.010.0100.01
000000000000000000
1168594
species

00000000000000000
0
species
0
1993864

no rank
439809
00000000000000000
0

species
1089132
1100000000000501008201
0.020.020.040.010.090.020.01

species
1071505
00.010.0100.010
00000000000

0
000000
373412
species

563122
species
00010000008120000000001
0.020.020.150.250.01

373408
species
00.050.030.020.020.010.01
00000002000103020101

species
1074303
000000000000
0.01

00000000
0.02
1993880
0.01
species

1204515
no rank
0.01
00000000

685506
species
0.010.010.010.030.01
00000000000000000000

546182
species
0.0500.020.010.020.020.010.03
02000001001000010000002

species
205876
000100000000000000
00.020.0200.010.010.010.010.0100.01

00000000
0.02
species
1993860
0.01

0.01
00000000
no rank
938131

genus
1982562
00.010.0200.01000.03
0000000000000000000001
00.030.0600.03000.09

1982563
00.010.0200.01000.03
species
00.020.0400.02000.06
0000000000000000000001.5

no rank
1838079
0000000000000000000003
00.010.0200.01000.03

10.50000.4000017.384617.79221.50.500.50.50011.333330.1666670.5
0.080.040.050.041.431.540.240.020.020.020.040.090.060.02
genus
0.040.020.030.020.720.780.120.010.010.010.020.050.030.01
1623275

1406784
species
0.040.0100000.01
20000000000000000000001

0.020.01
0000000000100000000000
1406782
species

1273741
species
0.010.01
0000000000000001000000

species
1273742
000.020.120.010.010.020.040.01
0000000000013100100231

species
1406788
0.020.010.690.740
01000100003637000000000

0.010000
0000010000000000000000
1273739
species

species
1273740
0.010
0000000000000000000000

0.180.090.060.130.110.130.060.130.030.120.050.030.05
00.3333330.33333300.3076920.7272730.30769200.30769200000.33333300.66666700000.33333300
0.060.030.020.050.030.050.020.050.010.040.010.010.01
1982236
genus

00.50.500.510.500.500000.5010000000
0.060.060.020.040.040.040.020.040.020.040.020.02
0.030.030.010.020.020.020.010.020.010.020.010.01
1982237
species

no rank
1796995
01101210100001020000000
0.030.030.010.020.020.020.010.020.010.020.010.01

00.5000.510.500.5000000100000.500
0.060.020.040.040.040.020.040.040.020.020.02
species
1982238
0.030.010.020.020.020.010.020.020.010.010.01

1796996
no rank
0.030.010.020.020.020.010.020.020.010.010.01
01001210100000020000100

00.030.240.030.060.030.0300
000.5000000.333333000000000.3333330000
genus
1623280
00.010.080.010.020.010.0100

species
2003505
0.060.020.010
0.120.040.020
001000000.5000000000.50

no rank
1079894
0.060.020.010
0020000010000000010

00.020.040.020.020
0000000000000000000000
00.010.020.010.010
2003506
species

1056830
no rank
0.010.020.010.010
0000000000000000000000

0.0200.020.040.020.02
00000000000000000001000.5
genus
1623307
0.0100.010.020.010.01

00000000000000000002001
0.0100.010.020.010.01
species
1236573

00.7857142.024392.066670.7777780.21.136360.880.510.50.7142860.2631580.501.545450.50000.3333330.50.8
0.020.140.410.30.270.050.220.250.040.160.140.140.190.040.110.0200.020.020.060.020.1
genus
0.010.070.190.140.10.020.110.120.020.080.070.060.070.020.060.0100.010.010.030.010.05
1623306

0100000000100001
0.030.0100.030.01
species
861045

0.0100.01
000000000000000000001
species
861044

00000012001000000000001
0.0100.040.060.020000.010.02
species
1056831

1097753
species
0.010.040.160.120.010.070.050.020.080.020.040.020.040.010.010.010.010.010.03
02550132121200041000112

species
1993872
0.030.020.070.010.010.020.050.0100
0.060.040.140.020.020.040.10.0200
000.50.51.50000000.50.5100000000

0011300000011200000000
0.030.020.070.010.010.020.050.0100
no rank
88870

0.060.0300.030.030.060.030.030.030.210.09
0.33333300000.33333300000000.33333300.666667000.3333330.333333020.666667
2034998
0.020.0100.010.010.020.010.010.010.070.03
genus

2034999
0.020.0100.010.010.020.010.010.010.070.03
species
0.500000.500000000.501000.50.5031
0.040.0200.020.020.040.020.020.020.140.06

1636261
no rank
0.020.0100.010.010.020.010.010.010.070.03
10000100000001020011062

genus
680116
0.060.220.120.140.050.020.220.070.020.060.360.120.070.060.070.10.10.020.080.040.05
0.120.440.240.280.10.040.440.140.040.120.720.240.140.120.140.20.20.040.160.080.1
1.503.52.5330.54.51.500.51.54.58.50.53.5444.51322

species
66797
307566193013917178892644
0.060.220.120.140.050.020.220.070.020.060.360.120.070.060.070.10.10.020.080.040.05

0.020.10.020.020.050.050.030.020.020.010.010.02
1982745
genus
00.33333310.33333300.250.30769200.6666670.3333330.3333330.3333330000.33333300.3333330.25
0.060.30.060.040.130.150.090.060.060.030.030.08

species
1982746
0.020.010.020.010.02
00.50000.50.500000000000.50.5
0.040.020.040.020.04

1636184
no rank
0.020.010.020.010.02
0100011000000000011

species
0.10.0200.020.050.030.020.020.010.01
1982747
001.50.5000.5010.50.50.50000.5000
0.20.0400.040.10.060.040.040.020.02

0.10.0200.020.050.030.020.020.010.01
0031001021110001000
1636183
no rank

1000.4000.5000.50.50.500000.8333331.50000.50.5
0.080.050.010.040.060.040.040.010.060.080.010.020.04
0.040.030.010.020.030.020.020.010.030.040.010.010.02
1623287
genus

00010000010000002300011
0.0200.030.020.0400.010.02
species
1129146

species
31535
20000010001100001000000
0.040.020.020.020.0100

0.020.020.050.020.040.050.090.130.20.360.050.020.010.010.010.010.070.01
1982251
genus
0.060.060.130.060.120.130.250.390.61.080.130.060.030.030.030.030.210.03
0.3333330.33333300.3076920.3333330.500.307692000.640.4102561.6666716.666700.7692310.3333330.3333330.3333330.3333330.33333320.333333

00.01
1982304
species
00000000000000000.5
00.02

1655019
no rank
00000000000000001
00.01

species
0
1982257
0
0000000000000000000000

0
0000000000000000000000
1655015
no rank

00000000000
0
species
0
1982308

00000000000
0
1655023
no rank

1982272
0.020.010.040.020.20.360.010.010.010.070.01
species
0000.500.5000010.52.525000.500.500.530.5
0.040.020.080.040.40.720.020.020.020.140.02

504553
no rank
000101000021550001010161
0.020.010.040.020.20.360.010.010.010.070.01

0000000000100000.50000.5
00.080.020.020.02
1982303
00.040.010.010.01
species

1655018
no rank
00.040.010.010.01
00000000002000010001

0.02
1982305
species
0.04
0000.5

0.02
0001
no rank
1655020

0.02
1982287
species
0.04
00000000.5

no rank
1235656
0.02
00000001

species
1982254
0
0
0000000000000000000000

1655012
no rank
0000000000000000000000
0

00000000000
0
0
1982260
species

no rank
1229782
0
00000000000

0.020.01
1982270
species
0.040.02
000000000000.5000000.5

0.020.01
000000000001000001
no rank
1498188

0.040.06
000000000000.50001.5
species
0.020.03
1982285

0.020.03
0000000000010003
1235654
no rank

0.010
1982262
species
0.020
000000.500000

00000100000
0.010
1229784
no rank

1982256
0.02
species
000000000000.5
0.04

0.02
000000000001
no rank
1655014

0
1982280
species
0
0000000000000000

1235650
no rank
0
0000000000000000

species
1982261
0.020.020.020.020.020.04
0.50.5000.5100.50001
0.040.040.040.040.040.08

1229788
no rank
0.020.020.020.020.020.04
110012010002

0.060.180.0300.030.03
0000.3333330000000100.333333000000
1921434
0.020.060.0100.010.01
genus

species
1921435
0.020.060.0100.010.01
0.040.120.0200.020.02
0000.500000001.500.5000000

no rank
1173743
0.020.060.0100.010.01
00010000000301000000

1299429
0.110.150.220.160.060.10.150.050.060.130.020.070.080.010.010.050.040.03
genus
0.270.390.550.380.140.220.350.130.180.330.020.140.190.020.010.10.10.06
0.5925930.4102560.4181820.68421100.50.3636360.4000.4242420000.51.526320.50000.80.30.333333

1981048
0.020.040.050.030.020.020.030.010.020.0300.01000.010.01
species
0.040.080.10.060.040.040.060.020.040.0600.02000.020.02
0.50.50.50.5010.50.5000.500000.50000000.5

11110211001000010000001
0.020.040.050.030.020.020.030.010.020.0300.01000.010.01
no rank
1718273

species
347328
11120111001000010000111
0.020.040.050.050.010.020.040.020.020.0300.0100.020.010.01

0.040.080.10.060.020.040.060.020.040.0600.02000.02
0.50.50.50.500.50.50.5000.500000.5000000.5
species
1981049
0.020.040.050.030.010.020.030.010.020.0300.01000.01

no rank
1204517
0.020.040.050.030.010.020.030.010.020.0300.01000.01
1111011100100001000001

species
279383
0.040.030.020.070.050.010.03
201000000010001510002

462590
species
1112011100100001000001
0.020.040.050.050.010.020.040.020.020.0300.01000.01

genus
1623282
0.030.040.050.020.090.110.0200.070.020.020.0400.04
000.50.500010.50.51.130430.5000.501001.5000.428571
0.060.080.10.040.180.230.0400.140.040.050.0800.07

00010000011000100000000
0.040.050.020.070.010
species
205872

00100002103100000000001
0.030.050.020.070.020.010.01
561996
species

00000000011000002003001
0.040.030.020.010.040.02
species
1089115

00.3333330.333333000.6666670000.3333330.3333330.3333330000.6666670.6666670000.33333300.333333
0.060.090.060.090.060.060.060.060.030.03
1982113
0.020.030.020.030.020.020.020.020.010.01
genus

00.50.50010000.50.50.5000110000.500.5
0.040.060.040.060.040.040.040.040.020.02
0.020.030.020.030.020.020.020.020.010.01
1982148
species

1486472
no rank
01100200011100022000101
0.020.030.020.030.020.020.020.020.010.01

genus
0.070.020.030.020.050.090.020.020.060.010.050.090.02
1982095
1.333330.3333330.3333330.3333330.6666673.1379300.3333330.3333330.3333330000.66666700000.7692312.3333300.666667
0.210.060.090.060.150.290.060.060.180.030.130.270.06

0.020.040.060.02
000000.5000.50.5000000000.5
0.010.020.030.01
1982098
species

no rank
1175657
0000010011000000001
0.010.020.030.01

0.140.040.060.040.10.180.040.060.020.060.180.04
20.50.50.51500.500.5000100001.53.501
species
0.070.020.030.020.050.090.020.030.010.030.090.02
1982096

41112100101000200003702
0.070.020.030.020.050.090.020.030.010.030.090.02
no rank
1548905

0.500.50.50.50.500001.51.5070022.50.500.50.50
0.040.060.040.040.020.020.120.120.20.080.120.0200.040.020
genus
1623308
0.020.030.020.020.010.010.060.060.10.040.060.0100.020.010

359961
species
101111000033014004510110
0.020.030.020.020.010.010.060.060.10.040.060.0100.020.010

genus
0.040.020.060.070.030.020.050.140.030.020.020.040.020.070.030.020.030.090.05
1982222
0.120.060.180.210.090.060.150.420.090.060.060.120.060.210.090.060.090.270.15
0.6666670.3333330.666667101.333330.3333330.66666720.3333330.3333330.3333330.33333310.33333310.6666670000.6666672.666671.33333

0.080.040.120.140.060.040.10.280.060.040.040.080.040.140.060.040.060.180.1
10.511.5020.5130.50.50.50.51.50.51.51000142
species
1982225
0.040.020.060.070.030.020.050.140.030.020.020.040.020.070.030.020.030.090.05

21230412611113132000284
0.040.020.060.070.030.020.050.140.030.020.020.040.020.070.030.020.030.090.05
1691954
no rank

genus
0.060.030.030.020.020.120.010.020.050.040.050.040.010.04
1623290
0.120.070.070.040.040.240.010.040.10.080.090.080.020.09
0.500001.428570000000.833333000.52.10.51.111110.50.51.33333

1000010000001001111100
0.030.020.010.010.010.0400.010.010.020.010.020.01
species
552525

0.030.020.030.010.010.0800.010.040.020.030.020.010.04
1000030000002001513113
701257
species

2.332.251.991.752.1621.642.051.591.893.113.220.621.860.90.810.951.662.352.472.742.882.36
3.78973.564442.733673.034292.6259.912.170732.71222.050312.460326.032154.338511.096779.284950.3888893.802474.621057.15066.225537.793527.259127.7951414.5212
genus
187218
0.830.830.710.630.790.730.640.770.640.721.211.260.230.690.370.280.340.610.930.941.131.161.04

0.080.080.10.240.080.020.120.180.060.10.20.320.160.10.10.140.040.10.280.06
11012.52.50120.512.5011042.522.51261
1982564
0.040.040.050.120.040.010.060.090.030.050.10.160.080.050.050.070.020.050.140.03
species

no rank
1728958
2202550241250220854524122
0.040.040.050.120.040.010.060.090.030.050.10.160.080.050.050.070.020.050.140.03

2.5100.511.50.50.50.50.511030120.513.51.541.5
0.20.10.060.080.10.060.080.060.040.060.10.10.020.10.040.040.080.040.060.180.080.180.1
species
0.10.050.030.040.050.030.040.030.020.030.050.050.010.050.020.020.040.020.030.090.040.090.05
1982565

1813783
no rank
0.10.050.030.040.050.030.040.030.020.030.050.050.010.050.020.020.040.020.030.090.040.090.05
52012311112206024127383

0.520.160.0200.020.040.060.040.0400.020.0400.060.04
6.520000000.500.50.501000000.501.50.5
0.260.080.0100.010.020.030.020.0200.010.0200.030.02
1945575
species

134000000101102000001031
0.260.080.0100.010.020.030.020.0200.010.0200.030.02
1572715
no rank

02002511003000000160221
0.010.060.030.020.060.040.020.030.010.060000.010.080.010.030.030.02
species
1112008

0110040010151013100314157143
0.030.030.010.040.020.020.030.290.210.040.020.070.0100.040.170.190.10.160.04
species
1567006

977927
species
0.010.020.020.020.020.010.010.020.020.030.060.120.010.07000.050.010.020.010.030.03
01011000013602100411022

491003
species
441227458715051041156282549
0.080.10.050.060.050.060.120.130.190.240.030.110.010.040.070.010.030.020.170.070.370.280.66

species
10726
01011032202303000411910
0.030.030.0400.070.050.050.020.040.060.020.010.050.010.020.120.020

0.30.420.550.390.420.490.330.420.20.380.580.560.160.310.140.150.20.370.380.480.40.340.19
352888
no rank
6.28.23814.917436.329115.5176519.13274.257585.578313.102563.7894710.77598.75213.29030.588.81511.565815.88547.88.735294.05405
0.60.841.090.790.850.980.660.830.390.761.161.120.320.620.280.30.40.740.760.960.80.680.37

0.260.380.380.350.330.390.260.340.170.290.480.420.160.230.070.150.170.370.310.430.240.240.13
141812141345101379252143211620302735182110
1150989
species

0.040.040.160.050.10.10.070.070.020.090.10.140.080.070.030.070.050.160.10.05
22524123313570111040641294
1589298
species

1982359
0.030.050.020.01
genus
000001000.666667000.33333300000000.333333
0.090.150.060.03

species
0.030.050.020.01
1982360
000001.5001000.500000000.5
0.060.10.040.02

0.030.050.020.01
00000300200100000001
1074308
no rank

0.120.10.20.080.340.040.070.070.120.050.040.10.050.060.090.08
00.50121.6251.529410.50000.66666700.60010.500.410.7777780.875
genus
1623297
0.060.050.10.040.170.020.030.040.060.030.020.050.020.030.040.04

species
180504
0.020.010.020.020.010.010.010.020.030.01
01000110000001000100021

255131
species
0.020.050.020.010.0100.020.010.01
0100002000000000010000

431891
species
01024441000202002101212
0.020.050.10.030.10.020.010.040.010.020.010.020.030.010.03

genus
0.010.030.020.010.010.0100.020.03
1623285
0.030.060.050.020.010.0200.040.06
000000.500000000010000000.333333

00000000000000000000001
0.010.020.0200.010.02
205875
species

species
1089135
0.010.010.010.01000.020.01
00000100000000020000000

0.50.7846150.5357140.25010.3939390.780.9555560.5357140.6190480.8823536.666673.29630.66666716.54413.175448.897810.250.90.2142860.4666670.685714
0.20.650.280.240.320.330.50.450.560.420.512.430.270.421.360.571.370.040.40.140.150.35
0.060.220.10.080.110.110.170.150.180.140.170.810.090.140.460.190.450.020.140.040.050.11
1982557
genus

0.160.020.040.020.040.060.040.10.1000.020.0200.10.040.040.08
01.50001000.501100000.50020.511
1982561
0.080.010.020.010.020.030.020.050.05000.010.0100.050.020.020.04
species

no rank
1838078
0.080.010.020.010.020.030.020.050.05000.010.0100.050.020.020.04
03000200102200001004122

0.020.140.120.080.040.060.040.20.180.080.160.0200.020.0800.040.020.020.04
01.510.5010.50211200.5000.5100.500.50.5
1982558
0.010.070.060.040.020.030.020.10.090.040.080.0100.010.0400.020.010.010.02
species

no rank
1838061
03210210422401001201011
0.010.070.060.040.020.030.020.10.090.040.080.0100.010.0400.020.010.010.02

0.020.080.020.040.020.040.040.040.020.02000.120.0400.040.020.020.04
00.50001000000000030.500.500.50.5
species
0.010.040.010.020.010.020.020.020.010.01000.060.0200.020.010.010.02
1982559

01000200000000006101011
0.010.040.010.020.010.020.020.020.010.01000.060.0200.020.010.010.02
no rank
1838065

10.40.50.502.222220.8333331.857140110.5105.51256.515.50.51.50.50.51.5
0.10.050.060.040.090.120.210.120.080.061.620.160.280.90.220.780.020.080.020.020.08
1982560
0.050.030.030.020.050.070.120.060.040.030.810.080.140.450.110.390.010.040.010.010.04
species

0.050.020.030.020.040.050.090.060.040.030.810.080.140.450.110.390.010.040.010.010.04
2111052302212011250133113113
no rank
1838074

0.060.060.090.06
00.333333000000.33333300.3333330.333333
0.020.020.030.02
1982101
genus

species
1982102
0.020.020.030.02
00.5000000.500.50.5
0.040.040.060.04

01000001011
0.020.020.030.02
1486424
no rank

0.940.480.320.171.090.440.380.230.740.30.580.952.811.480.961.031.810.940.430.790.550.760.58
1.989360.5208331.1250.3529412.596331.1590910.260871.256760.61.327591.642114.6156612.26351.5520813.03889.171271.904261.255812.063290.7818181.657891
0.450.220.160.10.560.220.210.130.370.150.290.471.470.750.570.580.90.480.20.380.260.380.29
1623298
genus

320849
species
0.010.010.0100.010.010.02
0000000000000000100002

00001000000000010201001
0.010.020.010.0100.030.020.010.01
species
487152

00000000000011012000001
0.0100.0100.010.060.010.010.020.0100.02
species
320844

1000002000000200000040
0.020.050.010.010.050
1197953
species

53369
species
00000000001100000000002
000.030.040.03000.03

59506
species
000.010.030.040.010.040.010.0200.020.03
00000000001100015010102

326037
species
00
00000000000000000

000000000000010022000
000.010.020.0300.01
species
872294

20000100410002000013140
0.040.010.020.010.10.030.010.020.010.040.020.040.01
species
1197952

0.0100.010.010.010.03
000000000000010012
species
405947

55511
species
30001311100072201223102001
0.060.010.040.030.030.040.020.30.160.040.120.20.020.0200.02

0000000000000000000000
00.01
387910
species

40112301205800001369264
0.070.030.020.050.030.020.050.10.160.010.040.070.110.030.070.05
320840
species

00000000001100000000002
000.030.040.03000.0100.03
species
320835

0000000010000000000000
0.02000
species
320842

0000000000000101201000
0.010.010.010.010.010.020.0100.01
320850
species

species
106284
00001300010083101326701010
00.010.040.030.010.010.030.340.220.040.120.220.090.010.010.020.01

0.060.020.030.020.020.020.040.020.070.020.010.010.020.030.01
31101200102100102112031
320841
species

species
320847
000000000000010012
00.010.010000.010.010.010.010.03

00.01
00000000000000001
326036
species

0.060.0600.010.020.020.010.030.020.060.030.040.01
1636203
no rank
0.120.1200.020.040.040.020.060.040.120.070.070.02
1.51.50000.5000000.500001.500.252.50.8571430.7142860.5

0000000000000000000001
00.010.010.01
species
1636204

33000100000100003015221
0.060.0600.010.020.020.010.030.010.060.030.020.01
species
1636205

000100000000002000000
0.020.020.14000.01
12360
species

399185
species
5130221030221300222206
0.10.030.10.010.060.020.020.010.090.040.040.040.020.020.030.030.030.010.07

species
379501
0.010.010.010.01
000000000000000000001

species
387908
00000.01
000000000000000000000

0000000000000300120002
0.010.010.010.020.010.020.010.02
320848
species

00010000020130000002001
0.020.060.020.120.020.01
species
320834

00001300000093101326700000
00.010.040.030.010.010.360.220.040.130.220.0900
species
320846

00009210001111001201213
0.210.020.020.010.020.020.040.010.010.020.010.030.010.04
species
398839

0100000020000000000001
0.02000.0500.010.010.01
575608
species

00000000000000
00.010
320845
species

genus
0.010.080.080.070.20.120.010.050.40.080.080.150.150.480.220.460.380.180.130.060.10.03
1912760
01.333330.50.6315790.9454551.4242400.1818185.03390.50.5909090.68292702.533331.846725.5454514.71018.070182.563640.9736840.2222221.517240.666667
0.030.240.240.190.550.330.030.111.180.240.220.410.451.370.661.381.140.550.380.180.290.09

0.070.010.0100.01
0000000000000010000000
297254
species

000102.5000.50010300.524.50.50200.5
0.020.040.10.020.10.020.020.040.040.080.080.020.820.020.10.020.04
0.010.020.050.010.050.010.010.020.020.040.040.010.410.010.050.010.02
1981024
species

0.010.020.050.010.050.010.010.020.020.040.040.010.410.010.050.010.02
00020500100206014910401
1500734
no rank

0.080.060.070.030.010.370.060.030.050.070.410.180.040.340.110.040.020.070.03
1981023
species
0.160.120.140.060.020.740.120.060.10.140.820.360.080.680.220.080.040.140.06
02101.5200810.51053102.513.54.51.50.531

no rank
1500733
0.080.060.070.030.010.370.060.030.050.070.410.180.040.340.110.040.020.070.03
042034001621201062052793162

0.040.020.140.020.04
000000000000000100300.5
1981022
0.020.010.070.010.02
species

0.020.010.070.010.02
000000000000000200601
no rank
1500735

508419
species
0.020.050.030.020.040.040.010.01
00012301002200000001

0.140.020.040.040.080.020.020.020.040.02
00001.50.500000.50.502.500.50.50.5010
species
0.070.010.020.020.040.010.010.010.020.01
1981020

0.070.010.020.020.040.010.010.010.020.01
000031000011050111020
no rank
1399942

0000000000.615385000.50000.333333000000
0.010.070.260.020.080.010.030.010.05
genus
1623283
0.010.030.120.010.040.010.020.010.02

00000000020010001
0.020.080.040.01
species
1541887

00000000000000000000000
00.0200.01
1173757
species

0.01
00000000000
1173754
species

1005056
species
00.020.020000.01
00000000000000000000000

species
1141134
00.020000.01
00000000000000000000000

1982878
0.020.10.030.040.050.040.040.0700.020.030.010.010.04
subfamily
0.080.40.120.140.230.160.140.300.060.070.070.010.16
0.250.350.250.21428600.434783000.1250.2142860.10000000.1666670.1428570000.4375

00.3333330000.666667000.3333330.333333000000000.3333330000.333333
0.060.060.060.0900.0300.03
genus
0.020.020.020.0300.0100.01
1983735

species
1983742
0.020.020.020.0300.0100.01
0.040.040.040.0600.0200.02
00.50001000.50.5000000000.50000.5

0.020.020.020.030.010.01
01000200110000000010001
1647302
no rank

0.060.180.090.010.090.060.010.200.010.010.030
0.3333330.5555560.333333000.55555600000.150000000000
0.020.060.030.010.030.020.010.0600.010.010.010
1983734
genus

00.040.020.0400
0000010000000000000
1983737
00.020.010.0200
species

no rank
1698358
0000020000000000000
00.020.010.0200

0.040.10.060.020.020
0.510.5000.5000000000000
1983738
0.020.050.030.010.010
species

no rank
1429793
121001000000000000
0.020.050.030.010.010

0.0100.0300
1983739
species
0.0200.0600
00000000000.500000000

1327765
no rank
0.0100.0300
0000000000100000000

00000000000000000000
00.020.02000.02
species
00.010.01000.01
1983741

no rank
1567472
00000000000000000000
00.010.01000.01

00.33333300.33333300.333333000000000000.33333300000.666667
0.060.090.030.0300.0300.0300.09
0.020.030.010.0100.0100.0100.03
1623277
genus

0.040.060.020.0200.0200.0200.06
00.500.500.5000000000000.500001
0.020.030.010.0100.0100.0100.03
1983736
species

01010100000000000100002
0.020.030.010.010.0100.010.03
1074309
no rank

00.33333300.3333330100.33333300.3333330001.333330.3333331.333330.6666670.33333314.66667112.33333
0.060.060.090.060.090.090.210.120.060.030.090.510.120.090.27
0.020.020.030.020.030.030.070.040.020.010.030.170.040.030.09
1918718
genus

0.020.020.030.020.030.030.070.040.020.010.030.170.040.030.09
1918720
species
0.040.040.060.040.060.060.140.080.040.020.060.340.080.060.18
00.500.501.500.500.500020.5210.51.571.51.53.5

756282
no rank
0.020.020.030.020.030.030.070.040.020.010.030.170.040.030.09
010103010100041421314337

1982367
0.540.571.121.1811.410.520.511.051.260.310.370.690.711.380.450.520.740.730.60.90.950.04
genus
5.481484.660827.199412.175110.0834.83453.344163.830077.57467.328952.580652.889913.0289918.28174.1763311.333311.35910.270312.41114.618.917924.84910.583333
1.621.713.363.5434.231.541.533.153.80.931.092.072.134.141.351.562.222.191.82.682.850.12

species
0.380.340.830.990.831.070.280.350.620.80.180.150.470.471.070.360.340.430.520.570.810.880.03
1982368
0.760.681.661.981.662.140.560.71.241.60.360.30.940.942.140.720.680.861.041.141.621.760.06
108132117.562.55.5713134.53.55.5337.52020.517.522.52331401

20162642351251114262697116615404135454662802
0.380.340.830.990.831.070.280.350.620.80.180.150.470.471.070.360.340.430.520.570.810.880.03
no rank
691964

0.320.460.580.380.340.680.460.320.860.940.260.420.440.480.620.180.360.620.420.060.160.140.02
45.54.543.5204.5397.5352.516.52510.512.591330.5
1982369
0.160.230.290.190.170.340.230.160.430.470.130.210.220.240.310.090.180.310.210.030.080.070.01
species

811987409618156105334102125182661
0.160.230.290.190.170.340.230.160.430.470.130.210.220.240.310.090.180.310.210.030.080.070.01
691965
no rank

genus
0.050.090.150.140.060.120.050.030.110.150.040.110.080.070.010.070.090.080.060.060.08
1623278
0.10.180.30.280.110.240.10.060.210.30.090.220.160.140.020.140.170.160.120.120.17
0.80.8888890.8666672.142860.4545454.1250.40.3333330.6666671.60.5555562.2727302.312502.571430.5111.6251.666670.9166671.23529

1034125
species
02201310201003000121013
0.010.040.070.010.030.030.040.010.060.020.030.010.020.0100.020.030.020.010.020.05

222511001141506061334432
0.040.040.060.120.020.090.020.020.120.020.10.050.060.010.040.030.060.050.030.03
861047
species

00000000000001000010000
0.010.020.0100.010.020.010.0100.010.0200.010.01
species
1086751

genus
1623296
0.230.550.610.430.330.510.240.520.480.640.680.740.040.290.140.10.090.280.360.210.280.220.51
1.826093.458721.841671.785711.134338.521.511632.414142.010423.031253.477613.8175705.3793111.351.222223.210535.042252.166673.018183.177787.06
0.461.091.20.840.6710.430.990.961.281.341.480.080.580.280.20.180.570.710.420.550.451

1211643
species
6116522516591112017244516710920
0.120.240.20.130.050.220.020.160.130.30.220.260.130.140.040.030.070.190.090.140.110.28

0.020.110.120.070.070.060.010.090.120.110.090.110.010.050.010.030.030.050.030.040.03
15332603534506013242202
species
1204537

1211642
species
032228322055030301031245
0.020.060.080.050.070.070.090.050.060.020.110.120.010.020.030.010.130.040.010.030.050.07

1211641
species
24342922216507011213222
0.050.090.110.110.070.080.060.070.060.050.130.120.010.050.010.010.030.020.050.030.020.03

01223604455605000242246
0.020.040.080.050.080.060.010.10.110.160.110.130.010.040.010.010.030.050.030.030.050.08
species
1211644

0.120.060.180.060.010.060.060.060.060.10.090.120.080.060.030.03
0.16666700.33333300000.333333000000.5000.33333310.250.33333300.3333330.333333
genus
0.040.020.060.020.010.020.020.020.020.040.030.040.020.020.010.01
1921117

species
1921119
0.010.010.030.010.010.010.010.010.040.010.020.010.01
0.020.020.060.020.020.020.020.020.080.020.040.020.02
000.500000000001000.51.500.500.50.5

0.010.010.030.010.010.010.010.010.040.010.020.010.01
00100000000002001301011
1610836
no rank

species
1921120
0.020.0300.020.010.010.010
0.500.500000.5000000.5000.5000
0.040.0600.040.020.020.020

1610837
no rank
10100001000001001000
0.020.0300.020.010.010.010

00000000000001000.5000
00.020.020.020
1921118
00.010.010.010
species

1468411
no rank
00000000000002001000
00.010.010.010

0000000000000000000.50
0.020.020.020.020.020.040
1921121
0.010.010.010.010.010.020
species

1610838
no rank
0.010.010.010.010.010.020
00000000000000000010

genus
1623309
0.040.030.020.030.050.070.020.090.040.040.030.010.060.010.050.01
0.080.060.040.060.10.140.040.180.080.080.060.020.130.020.10.02
100.50.501.511.50.50.83333300.50.5200.51.461540.520.5

2001032312011401304
0.040.020.030.050.070.020.060.020.040.030.010.030.05
538979
species

species
470314
0000000001
0.03

00100000000100003001
0.030.020.030.01
species
1188794

329254
species
000000000000000011
0.010.01

0.040.050.0600.030.080.040.10.020.0900.020.02
0.50.40000000.2501000000000
genus
0.020.030.0200.010.040.020.050.010.0300.010.01
1623284

373403
species
00000000000000
0

0.020.06
0000000000000
no rank
1776684
0.010.03

species
1679524
0.010.03
0000000000000

0.020.02
000000000
species
0.010.01
1993871

0.010.01
000000000
1035485
no rank

11000000102000000000
0.020.020.020.020.020.050.010.010.01
546183
species

species
205869
0
00000000000000

01.666670.33333300.6666674.333330.3333330.3333331.6666721.333331.6666700.3333330.33333300.3333332.066670.333333000.666667
0.30.090.150.330.060.060.360.540.240.480.030.420.030.450.030.06
genus
1982569
0.10.030.050.110.020.020.120.180.080.160.010.140.010.150.010.02

0.20.10.220.040.040.240.360.160.240.140.020.120.04
02.50016.50.50.52.5323000.500.52.50001
1982570
0.10.050.110.020.020.120.180.080.120.070.010.060.02
species

05002131156460010150002
0.10.050.110.020.020.120.180.080.120.070.010.060.02
no rank
1636545

species
0.030.040.010.070.090.01
1982571
000.500000000100.50.5003.50.5
0.060.080.020.140.180.02

1636546
no rank
0010000000020110071
0.030.040.010.070.090.01

genus
0.010.030.030.020.010.060.070.030.020.010.030.010.03
1623289
0.040.080.090.060.030.180.210.070.060.050.090.030.06
00.3750.33333300.3333330.33333300000.5555560.28571400.571429000.66666700.55555600.33333301

0.03
00000000000000000000002
624134
species

0.060.060.02
000.5000000000.501
species
1954380
0.030.030.01

673375
no rank
00100000000102
0.030.030.01

0.010.030.02
010000000000
1245890
species

1954378
0.010.020.01
species
0.020.040.02
000000000000010010

000000000000020020
0.010.020.01
1147142
no rank

0.080.02
0000000000100000000.5
species
0.040.01
1954379

1109721
no rank
0000000000200000001
0.040.01

species
0.01
1981027
000000000000000000
0.02

no rank
1527514
000000000000000000
0.01

species
0.010.010.020.010.020.030.020.01
1981026
00000.50.500000.50.5000000100.5
0.020.020.040.020.040.060.040.02

1567004
no rank
000011000011000000201
0.010.010.020.010.020.030.020.01

0.210.190.140.260.30.130.220.370.050.10.270.220.20.330.250.060.10.050.060.240.170.090.7
0.4285710.4210530.2857140.6923081.60.6153850.2727271.37838000.9629630.8181820.43.393940.360.50.50.80.8333331.333330.9411760.2222225.51429
genus
0.10.10.070.120.150.070.110.190.030.050.130.120.10.170.120.030.050.030.030.120.090.040.35
1623303

0.0100.010.010.0100.020.010.020.020.0300
00000100001104000000000
78541
species

0.030.030.010.040.020.010.020.0200.010.040.040.020.020.020.010.020.020.060.010.010.16
110111000022020110241012
species
306323

species
682370
0.010.020.040.050.110.010.030.1200.010.0400.040.010.090.010.010.020.010.040.010.01
01124114002012111200300

0.030.010.010.010.020.030.0300.0100.08000.010.020.020.020.02
100002110000011000011111
species
85154

0.030.030.040.020.010.020.010.010.040.040.040.020.020.010.020.030.010.010.16
110111000022130110021012
species
459845

0.3333331000.666667100.33333300.3333331.333330.666667010.6666670.3333330.3333330.66666700.333333000.666667
0.060.180.180.090.060.090.240.120.060.510.030.030.060.060.030.09
genus
0.020.060.060.030.020.030.080.040.020.170.010.010.020.020.010.03
1982150

0.040.120.120.060.040.060.160.080.040.340.020.020.040.040.020.06
0.51.50011.500.500.52101.510.50.5100.5001
species
1982151
0.020.060.060.030.020.030.080.040.020.170.010.010.020.020.010.03

480808
no rank
0.020.060.060.030.020.030.080.040.020.170.010.010.020.020.010.03
13002301014203211201002

1982372
0.020.020.01
genus
0.060.060.03
0.3333330.333333000000000000.666667

0.50.5000000000001
0.040.040.02
1982373
0.020.020.01
species

11000000000002
0.020.020.01
no rank
562742

0001000.3333330.66666710.3333331.333330.666667
0.210.060.150.210.090.240.12
0.070.020.050.070.030.080.04
1982239
genus

0001.5000.511.50.521
0.140.040.10.140.060.160.08
species
0.070.020.050.070.030.080.04
1982240

0.070.020.050.070.030.080.04
000300123142
1844477
no rank

1982218
0.020.010.010.040.01
genus
0000.33333300000000.666667000000000
0.060.030.030.120.03

species
0.020.010.010.040.01
1982219
0000.500000001000000000
0.040.020.020.080.02

no rank
1622192
000100000002000000000
0.020.010.010.040.01

0.040.030.020.060.020.020.070.020.130.320.03
1982106
genus
0.66666700001.3333300.33333300.6666670.3333330.333333000.3333330.6666675.333338.666670000.363636
0.120.090.060.180.060.060.210.060.390.960.11

species
1982107
0.040.070.020.02
100000000000000.51000000.5
0.080.140.040.04

no rank
1838067
0.040.070.020.02
2000000000000012000001

00000200.5010.50.500008130000.5
0.060.040.120.040.040.260.640.04
1982109
0.030.020.060.020.020.130.320.02
species

000004010211000016260001
0.030.020.060.020.020.130.320.02
no rank
1838071

0.180.110.260.120.360.130.230.110.060.120.030.120.030.120.180.050.050.050.31
0.444444000.30769200.6666670.6666670.3076920.34782600.3333330.16666700.33333300.6666670.33333310.4444440001.22581
1982748
0.060.030.080.040.120.050.070.030.020.040.010.040.010.040.060.010.010.010.11
genus

0000.50110.50.50000001000.50002.5
0.020.040.080.040.120.040.080.040.020.040.040.020.020.020.14
species
1982749
0.010.020.040.020.060.020.040.020.010.020.020.010.010.010.07

0.010.020.040.020.060.020.040.020.010.020.020.010.010.010.07
00010221100000020010005
1821554
no rank

0.020.040.080.040.120.040.080.040.040.020.040.080.040.020.020.020.04
0000.50110.50.500.50000101.50.50000.5
1982750
0.010.020.040.020.060.020.040.020.020.010.020.040.020.010.010.010.02
species

no rank
1838073
0.010.020.040.020.060.020.040.020.020.010.020.040.020.010.010.010.02
00010221101000020310001

0.040.010.020.010.010.020.01
1982751
species
100000000000.500.5000.5010000.5
0.080.020.040.020.020.040.02

1838084
no rank
20000000000101001020001
0.040.010.020.010.010.020.01

subfamily
0.020.0200.020.010.010.010.01
1982875
0.080.0800.080.040.040.040.04
00.250000000000000000.2500.25000

genus
1982880
0.020.0200.020.010.010.010.01
00.3333330000000000000000.33333300.333333000
0.060.0600.060.030.030.030.03

species
0.020.0200.020.010.010.010.01
1982890
00.50000000000000000.500.5000
0.040.0400.040.020.020.020.02

1327758
no rank
01000000000000000101000
0.020.0200.020.010.010.010.01

0.611.031.630.660.660.780.90.80.91.320.891.010.480.720.30.690.650.580.540.850.60.570.68
0.950820.5922330.4233131.015150.5454551.128211.111110.50.4888891.181820.6741570.5445540.251.430560.2333333.304351.461541.275860.6481481.070590.9166670.543861.23529
186765
0.220.360.560.220.210.270.310.270.30.440.310.350.160.250.10.230.220.20.190.30.210.20.23
genus

species
10742
0.020.0100.020.010.020.01
0000001000000000100100

0.0300.01000.03
010000000000000000002
37554
species

0000000000000000000100
0.010.020.010.0100.02000.010.010
10710
species

no rank
0.190.320.520.220.210.260.280.270.30.430.30.320.160.240.10.230.210.20.180.260.180.190.23
335795
0.390.631.030.440.440.50.570.520.590.880.580.640.320.470.20.460.420.380.340.520.360.360.45
0.9230770.9206350.6699031.522730.8181821.761.71930.7692310.7457631.772731.034480.8593750.3752.191490.354.956522.23811.947371.029411.115381.361110.8611111.86667

species
1147158
00202200000011002120011
0.060.010.050.020.010.010.040.010.020.020.020.010.01

0.010.040.050000.010.01
02100000000000000100
species
1147149

02202442242303024034325
0.040.090.010.060.040.110.060.060.150.040.060.020.030.020.0300.040.050.050.030.07
species
1572747

0.020.040.090.030.060.040.110.060.060.180.040.070.020.020.020.0300.030.050.050.030.07
12212542252302024024325
1572743
species

species
563769
000.0400
0000000000020000000000

0
0000000000000000000000
1147144
species

000002
0.02
species
906668

00000101110013001012001
0.010.020.020.030.040.020.010.010.020.01
10724
species

0010000000000900810000
0.030.060.070.010
1147151
species

0.0100.010.010.030.020.020
0000000000000103200100
species
1147143

species
1147152
00000000000000000001
0.020.01

10713
species
00000001000000015
0.020.14

species
1147157
0000000000000000010000
000.010

0.0100.010.01
00000000000000010000
species
1147150

22010301104100100012312
0.050.040.010.020.010.030.020.020.040.080.0400.070.010.010.030.040.010.03
species
1572745

000000
0.010
species
1147154

0000000000000500073
0.020.040.090.03
414970
species

0.010.020.010.0100.0200.0100.010.01
0000000000000000100000
species
1147146

1147147
species
0.020.03
000000000000000

32151622001101000211022
0.060.040.050.120.020.050.050.050.020.020.010.020.010.010.020.03
species
906669

species
1161903
0.01
000

species
335797
0.050.020.010.02
0000000020100000001002

0.020.020.010.020.02
00000000100000002
species
196242

species
1147156
0.030.040.0500.010.03
121000000000000000002

0011000001001300010213
0.030.020.030.040.020.010.020.010.03
species
10717

00000000000
0.01
species
1147153

species
10702
020000001021
0.040.020.040.02

1147148
species
0.030.020.020.010.020.010.020.0100.0100.010.01
1100020000000100010000

genus
0.050.20.120.030.010.01
1920750
000001.42.666671.6666700.333333000000000000.333333
0.150.60.360.090.050.03

species
1920752
0.010.030.010.01
000000.50000.5000000000000.5
0.020.060.020.02

0.010.030.010.01
0000010001000000000001
no rank
1340829

0.080.40.240.02
000002.542.500000000000
species
1920751
0.040.20.120.01

0.040.20.120.01
0000058500000000000
1305710
no rank

genus
1623302
0.220.210.080.110.10.130.020.070.090.120.190.10.170.140.010.040.070.230.320.030.090.08
0.650.610.220.290.290.40.060.210.270.360.570.290.480.420.010.120.190.670.940.070.270.24
1.353851.8852500.8275860.7931033.2250.3333330.2857141.333330.8333332.175440.6206903.041670.33333300.8333330.7368422.776123.446810.2857141.185191.5

0.040.020.020.060.020.040.160.1400.060.040.040.0400.040.02
0.50000000.5000.5005.50.501.50.510.500.50.5
species
1982381
0.020.010.010.030.010.020.080.0700.030.020.020.0200.020.01

0.020.010.010.030.010.020.080.0700.030.020.020.0200.020.01
100000010010011103121011
no rank
1208587

0.210.320.060.160.150.210.060.180.20.30.110.1100.020.080.220.320.040.120.14
2.380953.501.51.45.7142900.521.540.90909103.18182000.51.5560.52.52.5
0.110.160.030.080.080.110.030.090.10.150.060.0600.010.040.110.160.020.060.07
1982377
species

no rank
1173135
57033120143820700131012155
0.10.160.030.080.070.10.030.090.10.150.050.0500.010.040.110.160.020.060.07

0.180.060.060.020.040.060.040.020.020.040.080.040.14000.180.2600.02
20.5000.51.50.50000.5101.50.50004500
species
0.090.030.030.010.020.030.020.010.010.020.040.020.07000.090.1300.01
1982380

41001310001203100081000
0.090.030.030.010.020.030.020.010.010.020.040.020.07000.090.1300.01
1458851
no rank

genus
0.020.030.020.010.050.050.060.020.020.020.020.050.060.10.11
1982244
0.060.110.060.010.150.150.20.060.040.060.060.150.20.30.33
0.333333000.3333330000.2666670.6666670.25000000.3333330.250.3333330.6666670.5333330.91.333331.42424

0.5000.50000.510.5000000.50.50.500.5122.5
0.040.040.0400.080.10.10.020.020.020.040.060.10.14
0.020.020.0200.040.050.050.010.010.010.020.030.050.07
1982247
species

0.020.020.0200.040.050.050.010.010.010.020.030.050.07
10010001210000011101245
1168479
no rank

1982248
0.0200.010.020.0100.010.020.030.040.050.04
species
0000000000000000.500.5111.521.5
0.0400.020.040.0200.020.040.060.080.10.08

0.0200.010.020.0100.010.020.030.040.050.04
00000000000000010122343
no rank
1168478

0.140.120.130.190.230.160.170.060.310.120.120.070.10.090.180.230.270.010.070.14
0.2857140.16666700.31578900.250.1176470.3333330.096774200.1666670.16666700.28571400.30.2222220.3888890.3913040.48148100.2857140
0.060.040.030.070.070.040.050.020.090.040.040.030.040.030.060.070.090.010.030.04
1921125
genus

1921130
00.010.030.020.010.020.020.030.020.020.01000.020.02000.01
species
00.020.060.040.020.040.040.060.040.040.02000.040.04000.02
0000.5000.50.50.500.50.500.500000.51000

00010011101101000012000
00.010.030.020.010.020.020.030.020.020.01000.020.02000.01
no rank
1225800

00.010.0100.020.020.010.020.010.010.0100.030.010.0100.020.01
1921128
species
00000100000000.500.5010.50.500.50
00.020.0200.040.040.020.040.020.020.0200.060.020.0200.040.02

1173748
no rank
00.010.0100.020.020.010.020.010.010.0100.030.010.0100.020.01
00000200000001010211010

species
1921129
0.020.0100.020.010.010.020.01000.010.010.010.0200.01
0.50000000000000000000.5000
0.040.0200.040.020.020.040.02000.020.020.020.0400.02

0.020.0100.020.010.010.020.01000.010.010.010.0200.01
10000000000000000001000
no rank
1458857

0000000000000000.50.5011000
00.020.02000.020.020.040.020.0200.020.020.020.060.06000.02
00.010.01000.010.010.020.010.0100.010.010.010.030.03000.01
1921127
species

no rank
1173747
00000000000000011022000
00.010.01000.010.010.020.010.0100.010.010.010.030.03000.01

0.040.040.020.060.040.020.020.0400.020.020.020.020.02000.02
0.50.500.5000000000000.50.50.500000
species
1921126
0.020.020.010.030.020.010.010.0200.010.010.010.010.01000.01

11010000000000011100000
0.020.020.010.030.020.010.010.0200.010.010.010.010.01000.01
1173745
no rank

0.2941180.3456790.2950820.3648650.3333330.7142860.1481480.50.4133330.2205880.6333330.3333330.3333330.9250.129634.121211.866671.036360.4074070.5593220.3617020.5319150.421053
0.680.810.610.740.60.630.270.560.750.680.60.60.480.40.540.660.750.550.540.590.470.470.57
subfamily
0.170.210.180.210.170.160.090.140.210.180.150.160.120.110.140.170.190.150.150.160.130.130.15
1910999

0.410.380.180.260.280.330.140.360.40.20.350.280.120.40.490.380.280.310.180.130.130.3
0.3902440.4210530.1666670.50.51.212120.1428570.7222220.250.150.9714290.501.666670.1755.551022.921051.750.5483870.6666670.2307690.4615380.533333
0.130.120.060.10.10.110.060.120.140.060.110.10.040.140.170.120.10.110.060.050.050.1
1920866
genus

0.020.020.0200.0200.04000.020.0200
000000.5000000000000.5000
species
1920994
0.010.010.0100.0100.02000.010.0100

0.010.010.0100.0100.02000.010.0100
000001000000000001000
1416030
no rank

10.5000.51000.5000.50000000.50001
0.10.060.0200.040.0400.060.040.020.040.0600.020.020.02000.06
species
0.050.030.0100.020.0200.030.020.010.020.0300.010.010.01000.03
1920995

0.050.030.0100.020.0200.030.020.010.020.0300.010.010.01000.03
21001200100100000010002
1416031
no rank

0.020.010.030.050.0100.070.010.020.010.020.010.03
1921000
species
0.040.020.060.10.0200.140.020.040.020.040.020.06
0000.510.501.5000.5000000.500.5000.51

1147155
no rank
00012103001000001010012
0.020.010.030.050.0100.070.010.020.010.020.010.03

0.030.030.0100.020.0200.030.020.010.020.0300.010.010.010.0100.01
1920993
species
0.060.060.0200.040.0400.060.040.020.040.0600.020.020.020.0200.02
0.50.5000.51000.5000.50000000.50000.5

1416028
no rank
11001200100100000010001
0.030.030.0100.020.0200.030.020.010.020.0300.010.010.010.0100.01

0.040.020.020.020.02
0000000.5000000000000.500.50.50.5
species
0.020.010.010.010.01
1920997

00000010000000000010111
0.020.010.010.010.01
species
1048207

0.060.10.060.10.10.040.10.080.060.160.10.080.140.320.20.160.080.080.040.040.04
0.510.510300.50.50.52102.50.58.55.531.51.50.510.5
0.030.050.030.050.050.020.050.040.030.080.050.040.070.160.10.080.040.040.020.020.02
1920996
species

no rank
1495286
0.030.050.030.050.050.020.050.040.030.080.050.040.070.160.10.080.040.040.020.020.02
1212060111420511711633121

0.060.120.150.060.060.090.360.060.060.180.120.030.03
0.3333330.66666700.66666700.33333300.33333300.333333000.44444400000.6666670.6666670.8888890.5833330.3333330.333333
1920860
0.020.040.050.020.020.030.120.020.020.060.040.010.01
genus

0.04
1981042
species
00000000000000000001.5
0.08

0.04
00000000000000000003
no rank
1654927

species
1981043
0.020.050.080.020.02
0.5001000000001000011
0.040.10.160.040.04

1002000000002000022
0.020.050.080.020.02
1675609
no rank

0.020.040.040.060.020.02
000000.500.500000000000010.50.5
species
1981041
0.010.020.020.030.010.01

1647408
no rank
00000101000000000000211
0.010.020.020.030.010.01

0.080.020.060.040.040.02
010000.50000.500000000010.5
species
0.040.010.030.020.020.01
1920861

0.040.010.030.020.020.01
020001000100000000021
no rank
1129191

genus
0.020.050.040.050.020.030.020.020.040.010.040.01
187217
00.3333330.2307690.2857140.3636360.40000.50.3333330.33333301.16667000000.714286000.333333
0.060.130.070.110.050.060.060.060.120.020.070.03

1921008
species
00000000000001
0.01

species
1921004
0.020.03
0.040.06
000000000000.502

1216926
no rank
0.020.03
00000000000104

00011100000000000002
0.020.020.010.02
1002724
species

species
1073767
0.020.0100.030.010.010.01
00000000010001000001

species
0.020.030.020.010.020.01
1921005
0.040.060.040.020.040.02
00.50.500.50.500000.5000000000000.5

no rank
1572702
0.020.030.020.010.020.01
01101100001000000000001

0.040.040.120.050.040.030.040.140.150.10.050.120.040.020.090.180.06
0.50.510.40.50.3333330.501.50.400.400.6001.6666710.500.77777811
1920864
0.020.020.060.030.020.010.020.070.060.040.020.040.020.010.040.070.03
genus

1945565
0.010.030.020.010.040.010.04
species
000000.50000.500.501002.50000.51.5
0.020.060.040.020.080.020.08

0.010.030.020.010.040.010.04
0000010001010200500013
no rank
1141140

355246
species
0.020.020.060.020.020.020.070.030.020.010.020.010.030.030.03
11211010310101000210222

0.0400.06
00000000000.5000001.5
genus
0.0200.03
1920865

species
1540096
00000000001000
0.020

245685
species
00000000000000003
0.03

genus
0.050.010.030.020.0100.020.010.030
1623288
000.4545450000.50.5000001.500.500.5714290
0.110.020.060.040.0300.040.020.070

1074304
species
0.050.010.030.020.0100.020.010.020
0010001100000301010

0.010.010.02
000000000000000001
species
373407

1623304
0.10.020.050.030.050.020.060.080.050.130.030.470.020.610.380.130.030.060.130.050.03
genus
0.210.030.110.060.10.040.10.140.10.260.050.940.041.220.750.240.060.120.260.10.06
0.66666700.2727270.33333310.250.40.71428610.92307700.408.08511011.10666.013332.083330.8333331.333331.423080.60.166667

201021122000014055601101
0.0400.030.050.010.020.050.050.10.050.040.070.010.010.01
species
112023

species
90410
0.030.01000.0600.140.020.20.050.020.020.050.060.020.01
1000000002000190225223420

0001001002000200302020020
0.020.020.010.060.150.280.170.020.030.01
633135
species

10000000000103016010210
0.030.010.010.010.0100.020.020.010.0600.0100.030.020.01
species
64186

species
72638
0.0100.010000.0100.060.070.050
00000000000008076000000

000.6666670.66666700.83333300000.250.333333000000.250.333333000.333333
0.060.180.150.120.120.060.010.080.030.050.06
0.020.060.050.040.040.020.010.020.010.010.02
1982227
genus

001101.5000000.5000000.50.5000.5
0.020.120.10.060.020.0400.040.020.020.02
species
0.010.060.050.030.010.0200.020.010.010.01
1982229

1589746
no rank
0022030000010000011001
0.010.060.050.030.010.0200.020.010.010.01

species
1982228
0.010.010.0300.010.010.01
0.020.020.0600.020.020.02
000000.500000.500000000000.5

1589747
no rank
0.010.010.0300.010.010.01
0000010000100000000001

genus
0.050.01
1982091
000000000.66666700000000000000.333333
0.150.03

1982093
0.050.01
species
00000000100000000000000.5
0.10.02

00000000200000000000001
0.050.01
1471542
no rank

0.460.581.261.060.481.31.280.910.551.430.660.590.41.991.530.151.910.260.420.170.370.190.22
0.3695650.3620690.5476190.9622640.7708335.861542.656251.538460.3272730.5244760.9848480.1864410.57.095482.065360.93333312.48170.6923080.7857140.5882350.9729730.3157890.636364
genus
0.170.230.50.420.20.550.580.410.210.540.290.230.140.720.730.080.670.120.180.070.180.080.1
186764

species
540068
0.020.110.070.020.220.20.150.050.10.020.210.020.010.020.05
0133126860151003210204

0.030.050.090.020.170.010.030.010.41000.040.050.040.01
0114027001000160034031
species
1089111

species
0.020.010.030.010.020.010.050.010.010.020.010.020.01
1993865
0.040.020.060.020.040.020.10.020.020.040.020.040.02
0.500.5000.500.500.50000.500100.510.5

0.020.010.030.010.020.010.050.010.010.020.010.020.01
101001010100010020121
1071504
no rank

species
0.010.010.010.010.010.010.010.020.190.0100.440.01
1089129
0.020.020.020.020.020.020.020.040.380.0200.880.02
000000000000.501300260.5

00000000000102600521
0.010.010.010.010.010.010.010.020.190.0100.440.01
1445715
no rank

species
1993867
0.020.030.110.040.010.06
0.500.5006.50000000.50.5003.5
0.040.060.220.080.020.12

0.020.030.110.040.010.06
101001300000011007
no rank
1071503

species
1109716
0.020.020.030.010.0100.020.01
000000011100010000101

0.02
00000000000000.5
species
1993861
0.01

1086800
no rank
0.01
00000000000001

0.01
00
species
260121

species
1051140
00000000000001000000000
0.010.01

000
0.01
species
1089128

1076136
species
0.01
0000

0.01
2003503
species
00000
0.02

1071506
no rank
0.01
00000

010000.5
0.10.040.02
species
1993862
0.050.020.01

0.050.020.01
020001
260120
no rank

00000000000001
0.01
1089118
species

00000000000000000010010
0.020.010.010.020.020.020.010.010
1051138
species

species
663557
00000100001000000000000
0.010.010.010.010.020.0100.0100.0100.010

00000000000000000
0.020
0.010
1993886
species

00000000000000000
0.010
701456
no rank

species
540065
0.010.010
000000

0.060.030.130.150.010.060.020.040.090.230.070.040.080.290.070.010.050.020.010.020.020.04
359350
no rank
0.120.060.260.290.020.110.040.080.180.460.140.080.160.580.140.020.110.030.020.040.040.08
0.6666670.3333330.7307691.4137902.2727300.50.7777781.021740.7142860.25111.25860.50.51.636360.3333330.510.250.5

1527516
species
0.040.030.0900.030.020.210.070.010.030.010.010.010.01
20140000011002911311010

1354511
species
00.020.070.010.03000.01
0000000012000300000000

1527535
species
0121050123212700300202
0.010.020.080.020.050.010.040.060.10.040.020.080.050.030.020.010.02

1673873
species
0.010.010.020.030.010.010.010.030.010.010
000100000000000000000

28369
species
0.030.020.020.0700.010.0100.020.0200.030.0100.02000.02
11002000000000030101001

0.030.010.01
000000000000000000000
148603
species

species
540066
000000000000
0.01

species
0.010.04000.0100.02
1993882
0000.5000000000000000.5
0.020.08000.0200.04

0001000000000000001
0.010.04000.0100.02
1175504
no rank

10100334111101800610000
0.020.030.030.070.10.020.060.030.030.130.010.050.020.01
205870
species

0.040.060.140.040.20.040.060.020.040.080.020.06
00.50.501.5120.500.500.50300000.50001
1993885
0.020.030.070.020.10.020.030.010.020.040.010.03
species

555603
no rank
0.020.030.070.020.10.020.030.010.020.040.010.03
01103241010106000010002

00000000000000000000102
0.010.010.010.01000.010.03
species
373405

0000010000000000000
0.010.010.010.010.010.010.01
species
1089116

0.010.01000.0100.01
0000000000000000000
species
31757

000000.50000.50000000.50.500000
0.020.040.020.040.0600.040.020.020
species
0.010.020.010.020.0300.020.010.010
2003504

711470
no rank
0.010.020.010.020.0300.020.010.010
00000100010000001100000

1921553
0.0400.0700.010.03
genus
00000.3333330000000000.33333300000.3333330.666667
0.1200.2100.030.09

0.0400.0700.010.03
1921554
species
0.0800.1400.020.06
00000.50000000000.500000.51

000010000000001000012
0.0400.0700.010.03
1610828
no rank

0.030.030.030.030.030.03
0000000000000000.3333330000
genus
0.010.010.010.010.010.01
1982220

0000000000000000.50000
0.020.020.020.020.020.02
1982221
0.010.010.010.010.010.01
species

00000000000000010000
0.010.010.010.010.010.01
no rank
1636257

genus
1982554
0.020.040.010.010.010.010.01
0.060.120.030.030.030.030.03
000000000.333333000.66666700.33333300.33333300.33333300.33333300.333333

species
0.020.040.010.010.010.010.01
1982555
0.040.080.020.020.020.020.02
000000000.500100.500.500.500.500.5

1383056
no rank
0.020.040.010.010.010.010.01
0000000010020101010101

2.613132.31791.769112.078842.638215.255031.891722.247713.50342.080321.745491.779987.6199815.67462.7657321.7194252.50533.37763.760335.928654.109359.17412.63236
17.0617.7119.360000000000117.630000000000119.6819.370000000000217.640000000000119.660000000000122.0721.040000000000117.1717.6831.1322.160000000000225.9127.120000000000146.6825.980000000000115.7318.080000000000115.7315.9115.4499999999999
no rank
8.468.949.78.839.839.678.959.9811.1110.48.518.815.4811.0312.8613.5423.313.057.859.157.98.047.86
196894

0001001101000011000202
0.020.020.020.030.070.010.020.02
1161901
species

species
1168549
02000000000021
0.040.080.01

species
1572712
7223697194887255198271829121312
0.130.040.060.070.140.080.170.040.210.120.150.160.280.180.350.170.070.340.20.360.160.140.16

0000000001000001000002
0.0100.010.010.030.0100.0200.03
species
1262523

0.020.020.010.010.030.010.010.010.010.010.010.010.010.010.01
00000200001001001000010
species
1486425

10000010000000000000000
0.020.020.010.01000.010.010.01
39103
species

00.010
0000000000000000010000
1647434
species

1887648
species
00010100000000000000102
0.020.010.010.03

0.01
00000000000000001
species
1414739

00100461000100000000001
0.030.030.150.020.020.01
1647438
species

11000322100005020021625
0.020.020.030.050.050.020.040.020.020.010.080.020.07
1403390
species

00010101042000005000000
0.020.020.010.020.010.130.040.0100.040.01
species
1815969

species
43685
83122721221106215123991055
0.160.070.040.060.070.060.060.050.050.060.040.040.010.050.170.140.10.040.10.110.140.060.07

1000131000010100001171
0.020.020.030.020.010.030.010.010.010.010.010.090.01
species
1327959

0.050.020.010.010.020.010.01
00002200000002012100
1868599
species

0.020.030.020.070.030.060.160.210.10.020.150.06
10100203010302331120135
1610872
species

0.020.020.060.050.050.030.040.040.03
11220000212200000000002
species
1109713

1636255
species
00011100000000000000000
0.020.020.010.01000.01

1347760
species
0.020.010.050.010.010.120.010.020.010.0100.0100
10002000000030021000000

species
1506716
01200111110101032100001
0.010.020.060.010.020.020.020.050.020.020.010.030.020.010.010.010.02

0.010
00000000000
1383057
species

10000100000000000230000
0.020.010.010.010.010.010.010.0200.020.030.01
1587907
species

0000000000100000
0.020
1204526
species

000000
0
1527580
species

10704
species
0.020.050.04
000000102000000003

1873895
species
00000000002
0.04

01000000001001032600
0.030.020000.010.030.010.010.030.020.070.01
1873777
species

1873887
species
0.050.010.010.020.020.020.0100.01
00100000101002010000001

1527463
species
0000000000000000000
00.010.0100.0100

0.01
0000000000000001
1538804
species

0.010.010.0100.01
0000010000000000000
1262518
species

species
1852563
0.010.020.020.010.02
000001100001000012

0.020.0200.020.020.0200.070.0100.010.0200.01
11000000001000110000100
species
1582151

1887647
species
002002020021000201
0.060.020.050.040.020.020.01

0.020.050.030.020.020.010.03
10002300000100020010002
877240
species

0.020.020.020.10.020.10.010.010.030.01
1000021400010011102001
745088
species

species
173824
0.020.030.020.010.010.01
0001000001010100010001

0000000000000003
0.0100.010.03
1821562
species

700091844012050130777962370012
0.140.010.240.160.110.110.010.040.042.030.930.510.70.810.290.090.010.010.020.04
species
1505027

1463812
species
0000010010000000000001
0.010.0100.040.01000.01

0.050.110.070.030.060.070.020.060.050.010.020.020.010.020.010.020.010.040.04
25031630122003021101143
species
1838080

species
1165702
0.010.020.010.070.0100.0700.010.010.01
0000112000000010000011

species
1168280
0.020.010.020.01
00000100000101

0.030.060.040.010.030.02
0000030000320000000122
1887651
species

32111300110001411893344010
0.060.040.030.020.020.030.020.040.0500.010.10.10.160.080.410.050.060.010.020
species
1871681

00.010.020.010.010.010
00000100000001001000000
1357713
species

species
40521
0.020.0200.010.020.030.020.020.010.0200.020.01
00011000001103012010

00000300000000000001001
0.010.010.030.010.010.010.0100.01
species
1673876

0.040.020.020.020.020.090.080.040.010.090.070.030.030.010.020.0400.020.01
210112033100013133123021
species
1327772

1679530
species
000.01
0000000000000000001

species
1536604
0.020.050.010.010.020.120.0100.020.020.01
010021001400000020101

1622190
species
0.010.010.01
000000000000000000000

1674927
species
0.02000.020.030.020.070.02
01000001010002102

1536603
species
0.0200.020.010.010.010.030.0100.02
10010000001001000000001

1084722
species
0.030.050.010.050.040
0102010000220000

0.060.030.050.050.040.020.040.040.040.030.140.050.070.030.010.080.030.01
30122500102214206631631
species
1305708

species
1340713
00000.010
0000000000000000000000

0.030.010.0300.030.020.0300.0100.03
00100001000000023000002
species
1527536

01001535232212054342101
0.010.030.020.050.090.120.050.090.040.040.040.020.050.040.040.050.040.020.01
354260
species

0.020.010.010.010.010.050.050.010.010.01
0000010201000000000
species
1327771

00000000001000000000
0.0100.010.010.020.010.010.01
1340708
species

species
1838083
14021321106200000012201
0.020.080.050.020.030.060.040.040.020.130.050.010.020.030.01

0.030.04
00100000002
1647285
species

772054011203230062310461
0.140.160.060.120.030.010.020.020.060.060.080.020.050.020.030.130.060.070.02
species
1124578

1312000002003501513010
0.020.060.040.050.010.010.020.060.010.010.120.040.010.050.020.030.010.01
1056833
species

1536596
species
000000000000000000000
00

species
1673877
00000200103101020100001
0.020.020.060.020.010.020.010.0200.01

10100000020000000000101
0.020.030.060.010.01
1458711
species

0.10.020.070.040.020.050.060.110.040.070.020.030.010.020.010.030.020.01
003135012262010024121221
species
537874

species
1391179
000000
00

species
1871692
0.060.030.020.010.010.010.030.040.360.020.020.150.010.050.010.030.010.01
300111000120930217141210

1873452
species
01000000010000000001001
0.020.030.010.01

0.010.050.01000
00000000000006000000000
species
345922

251001715436102171018721
0.050.120.060.010.010.180.370.110.110.120.020.020.10.060.0100.010.110.10.030.02
673839
species

120002000000012203
0.020.040.020.010.140.020.04
species
756275

00000100001000005
0.010.030.010.010.020.020.0100.04
species
1340818

species
1327974
0000010000000002000002
0.010.020.02

0013000000000110003
0.030.070.010.070.03
151539
species

species
754049
0.040.020.020.040.020.020.060.010.050.01
21001501100301000041

1176422
species
0.021.070.01
000100000000000012800001

1147043
species
0.060.030.050.190.150.020.020.150.080.120.450.10.260.10.070.150.170.030.20.05
0312817110546111402912613142184

000000000000000000
0.020.010.040.040.0100.05
species
0.010.010.020.020.0100.03
1391433

000000000000000000
0.0100.010.0100.01
1391430
no rank

00.010.0100.01
000000000000000000
no rank
1391429

0000010000000000000001
0.010.0100.02000.01
1463811
species

00000000000000
0.0100
species
1327774

species
1821727
001000010000000100000
0.0300.020.010.010.0100.01000

151538
species
0.010.010.04
00000100000001000000003

species
1567471
000001000010000000031
0.010.010.010.020.040.01

00000000000000000000110
000.010.0200.010.010.01
species
1636263

1541824
species
00.0100.020.010.01000.02
00000000000000000001

1675548
species
00100000000000
0.030.010

1168613
species
0.020.020.030.020.020.0400.020.010.010.03
10001001001105000100112

0.020.060.01
01000000020
1838063
species

00.010.010.01000
00000000000000000
373411
species

0.020.030.050.070.010.10.070.020.060.080.020.020.030.010.050.010.010.020.05
10123143124103004141124
species
59241

species
77048
10000000101000000001
0.020.020.020.01

0.210.150.020.030.01
000091700110001
741942
species

0.020.010.170.01
00010100000000000000131
1296646
species

species
423476
0.020.020.010.07
01000000000100018

species
1871687
53027141321023103221321976461014
0.090.070.020.060.170.130.320.050.040.020.040.070.420.230.151.21.650.080.050.080.140.020.06

1340826
species
12205810321203013105204
0.020.040.060.010.120.070.020.080.080.030.040.010.020.010.030.010.010.070.030.05

0000000000000000000
0.010
1647433
species

0.010.020.010.010.040.02
00000100002000000002
species
1109710

species
1458670
0.020.050.030.020.030.060.010.01
010024010130000000101

species
496874
0.020.010.020.020.020.070.020.020.010.030.07
100001111000010002021205

0.060.050.050.040.020.010.010.030.050.010.05
03000022011001004400004
1089117
species

0010000000000001010000
0.030.020.010.0200.010.0100.010.010.010.01
1887652
species

0.0400.010.010.110.060.350.010.010.01000
200000000000274100000
species
1262536

species
348137
0.02
00000000001

species
1647476
0.040.030.020.020.020.040.010.01
02100210000110000000011

species
1821557
00310100101000000000002
0.10.020.010.020.020.03

754056
species
010101000010
0.030.020.010.020.01

0.070.040.030.020.030.050.020.040.020.030.030.010.010.030.10.060.020.050.060.070.020.01
421116001011010117145510
species
12390

00010000010012342235803
0.020.030.490.240.140.210.040.10.04
644007
species

species
1414736
0.020.010.010.0300.010.010.01
00000010000000030000000

110123004301113133047522
0.020.020.020.050.030.090.090.020.040.090.070.030.030.050.090.060.020.03
396359
species

1327776
species
0000000000000001000000
0.010.0100.0100.010

1821564
species
110101000011001
0.020.020.020.010.020.020.07

0.01000.010.02
00000000000000000012
species
1262538

1340709
species
0000010000000000000000
00.010.010.010000.010

0000000000000000000000
00.010.010
1821538
species

31754
species
0.010.020.010.010.020.010.010.0100.020.010.03
01000100000001000002002

species
1340825
0
00000000000000

species
1647304
0.030.050.060.070.020.030.050.030.080.060.010.010.040.020.01
1223130201430001132001

1340831
species
00000000000000000000000
0.010.010.0100.0100.01

100000000000001300000
0.020.010.010000.140.040.0100.01
species
1498171

species
627087
0000020000000011000002
0.0200.070.010.02

1821559
species
0.030.020.0100.040
0010001000000000000300

1204539
species
0.040.060.060.050.080.070.070.10.020.080.080.060.020.070.050.050.10.030.020.040.080.13
232238341243031568323710

0.020.010.010.010
000000001000000000010
1622191
species

000000000000000000000
00.010.010.010.010.01000
1659230
species

947981
species
22010011101110084210102
0.040.040.0200.020.020.020.020.020.040.070.030.020.010.010.03

species
1235649
0.010.010.01
00000000000001010001

1105171
species
000000010000011430001
0.020.020.010.070.040.030.01

1486427
species
0.010.010.030.020.0100.010
0000000000110000001000

111000000000000300100
0.020.020.030.0300.010.01
1458846
species

species
1277672
00000000000000000000
0.010.010000.010.010.01

12417
species
0.050.030.070.050.020.020.020.050.020.010.030.03
00002432101100062010202

0.01
000000000000000000001
species
1383054

species
754042
0.340.130.130.140.210.130.020.10.030.140.090.040.050.070.020.090.010.070.060.050.070.05
18646915140174161211165464

0.070.030.070.020.010.070.040.010.040.040.01
00003300301002080313301
1187894
species

species
1838062
0.050.0200.050.010.060.010.01
0201000200030200001

0.010
00000000000
species
1354514

54756196886210719877731201627236
0.090.080.220.120.140.160.150.210.190.180.040.20.280.140.550.70.060.380.230.20.350.250.08
139872
species

0.020.020.010.020.010.080.03
000111001000010900002
754053
species

species
1591073
0000000000000000000000
0.010.010.010.020.0200000.010

species
1883202
0.020.050.01
0000000120000000001

species
1325953
371441531105420211801651211
0.060.150.450.090.020.040.070.020.230.150.090.040.010.070.010.070.180.060.010.230.01

species
1204525
0000000012120100300111
0.020.060.020.040.010.030.010.010.01

10010001100020000041021
0.020.020.020.020.080.050.010.020.01
species
1815955

00.0100.010.010.030.040.1200.140.01000.02
00000000001130100000001
1195086
species

species
1429912
00000100000000000010013
0.010.010.010.01000.010.020.04

000003000100041604003
0.010.030.050.030.070.060.050.04
99179
species

871584
species
0.060.030.020.050.070.020.120.090.040.060.030.070.020.030.020.080.020.430.130.08
3011063153230412317233126

61340364710214010033233
0.110.030.10.110.030.150.10.180.030.040.040.030.010.040.040.030.030.04
1327981
species

0.020.020.050.010.020.010.010.010.020.010.010.050.010.010.01
11002110000001016000001
species
73422

0.020.010.010.050.040.030.010.0100.0100.020.04
01000111100001000010013
1391431
species

0.020.020.020.040.020.070.020.040.010.01
11010501301201000001
species
1105286

0.070.020.070.170.030.020.140.030.030.050.050.010.010.020.010.010.01
41037310610004056002111
species
139870

00251010001102110218131
0.010.060.120.0200.020.020.020.010.070.010.020.010.110.010.030.01
1445809
species

0.090.040.020.030.080.050.010.030.180.090.030.060.030.030.030.040.040.020.030.060.050.090.05
51013501731304044124384
1042123
species

species
1340821
00.010.0100.01000.01
00000000000000000001

0.010.050.010.010.010.020.020.040.030.0500.0100.010.020.01
00020100011000000000110
species
1838081

species
1673889
000000000001
0.02

100100110002113001001
0.020.020.020.020.040.040.090.010.01
1838064
species

0001000000000000000
0.030
species
1654781

0.010.050.050.02
0000010000000005000401
1262533
species

0.020.020.050.010.020.020.020.070.010.010.04
01020101101000110103
1445726
species

0.020.010.040.060.050.02
0100010000001000005402
446529
species

species
10732
0.060.030.020.030.110.040.040.030.210.04
30000300116200000032163

101570
species
00000000100000002000020
0.020.020.020

0.01
00000000
1675549
species

species
1647469
0.030.070.030.040.020.070.01
00000300312100100000001

species
1527534
0.010.030.010.020.020.0100.03
00000000001000000000002

00000000000000000000000
000.010.0100
species
1821549

0000000000000000000021
00.030.01
1647308
species

0.030.020.0200.020.010.03
0010001000010000021003
species
1175662

000000000010
00.020.01
1654740
species

0.080.030.020.020.010.0100.01
04100000100100010000101
species
1315956

1623132072367151113207121971813132112852
0.30.480.420.470.170.20.150.170.350.330.020.060.810.511.470.880.150.160.150.260.160.930.03
species
475178

00000000000101000010100
00.020.010.010.010
species
1589751

000006000000000000001
0.010.0500.01
1560286
species

0.020.030.040.020.01
00010000012100000001
1647470
species

species
347962
00000000000006000000000
0.010.050.01000

species
1262517
152121622431401052312135504
0.020.120.060.020.050.140.050.050.110.090.020.080.010.050.190.160.150.070.070.05

432200
species
0.01
00000000000000

0.0200.010.02000.010
10000000000000000000010
1194641
species

0.010.010.010.0100.020.0100
0000000000000201000
species
1327952

species
1486473
02020113100000004000020
0.050.060.020.010.040.090.020.0100.04000.020

60002111000202152112022
0.120.010.050.010.020.020.010.050.010.070.050.020.020.020.020.020.03
120574
species

01211531207716441475127331
0.020.060.020.020.040.070.020.050.140.150.650.310.070.040.630.020.020.090.040.030.01
species
1887654

0.010.050.010.010
0000000020000000000000
species
1555233

species
1821561
0.01
00000000000

species
51369
0.060.020.0100.01
002011000000000000001

species
1401669
1261562110121214512442727341210515
0.020.050.190.360.150.180.250.310.280.440.10.030.080.310.140.060.230.040.050.150.140.050.2

1486426
species
120200122103340165101210
0.020.040.020.0600.020.060.060.050.010.060.120.030.150.040.020.010.030.020.01

0.0200.010.020.020.020.070.020.0200.01
0100000000110312200001
1414747
species

0000000001000001000000
0.0100.010.010.030.010.0200
species
1262519

0.020.02000.010
0000000010100000000100
species
432201

658401
species
01100521004001011032001
0.020.030.040.050.020.0800.010.010.010.030.020.01

species
1416012
561326104303091221031241
0.110.130.030.070.050.050.020.090.090.060.070.10.020.020.120.030.010.030.050.02

0.060.040.050.070.020.10.020.060.020.10.020.030.020.010.090.040.01
320028141210014024021741
413066
species

0.02
00000000001
species
1235645

species
1273712
00.010.010.02
000000000000000000001

1133022
species
0.040.020.020.020.010.030.01
21001000100000000100201

00.0100.010.010.01
00000000000001
1195071
species

0.020.030.020.020.010.040.010.030.030.020.010.040.01
01111100000202033213001
species
1552735

000011000000000000110
0.020.010.010.010.01
species
1555235

0.020.020.010.020.060.010.01
0101010010000800011
species
1445729

01202200110201001000007
0.020.060.050.020.020.030.040.010.010.09
1458831
species

1647424
species
0.040.01
00000000000011

010001001111013005311
0.020.010.020.030.020.020.010.210.060.030.010.01
1873985
species

0000000100000003013211
0.040.030.010.030.020.010.01
species
1566990

001000000000101321020
0.040.010.020.040.10.030.020.010.010.020
1536605
species

0.020.020.020.020.050.030.020.040.040.010.010.020.060.04
10010011211211001250003
species
1458842

646413
species
55134445120108030114631917
0.10.110.030.070.10.030.10.140.020.060.010.020.060.0300.010.160.070.050.210.23

species
1262522
000003011000000001011
0.030.020.040.010.010.01

0.210.180.060.070.250.150.120.090.140.060.080.10.010.150.040.080.090.20.150.20.090.18
1182310175362450200497181115813
881953
species

species
1560285
0100000000000101401
0.020.0200.010.020.010.130.02

species
1519999
0.030.010.010.010.010.030.020.0900.060.070.030.010.030.01
10000100000000090462030

0.030.010.01
00100000000000000001001
species
467481

0.010.030.020.02000.01
0000000110000000000001
species
1463809

981323
species
0.020.010.010.020.01
00010100000002020000001

0000010000000000000000
0.010.010.010.0100.0100.010
1458852
species

species
1432658
0.020.010.020.020.010.010.010.020.040.010.03000.010.020.03
10001000000110000000021

0.02
00000000001
1109717
species

species
1340820
0.0200.010.010.020.0100.01
000000001000000000

000000010000000001001
0.020.010.01
species
1647309

species
1589748
000100013041010010101
0.020.020.070.080.020.010.010.010.01

10200000010001005142031
0.020.060.0200.030.010.010.040.020.050.0300.040.01
species
35345

species
1815608
0.010.020.02
00000000001

0.020.050.030.550.010.050.090.020.010.01
100024000000008167211
species
1147137

0
000000000
1662284
species

0.020.020.020.020.020.050.0100.010.01
11000000102000000000
species
1357669

1327993
species
51101200001201020025202
0.090.020.030.020.020.020.040.010.020.020.060.030.03

species
1887655
0.020.150.050.020.060.040.040.020.130.030.010.010.010.010.030.01
17020221106101000111201

species
984175
0.020.010.010.01
00000000000000001

species
1887649
0.10.090.020.0200.010.0100.010.010.010.01
0033020000000001010000

01230101100083022271432
0.010.020.060.070.010.020.040.320.020.020.020.020.090.010.050.030.03
1445858
species

species
1327764
00000000000000
0.0100

species
1852629
0.040.01
0000000000001001

species
1399941
00001000000000000603003
0.020.070.040.04

0.010.020.060.050.020.050.060.020.10.060.020.020.050.020.040.020.060.040.030.020.060.05
01121520421113042432153
species
541865

753084
species
2001014211221501093102
0.040.020.010.10.050.020.030.040.040.040.040.010.120.030.010.02

02010124100000000000000
0.050.040.020.010.050.10.020.0100000
species
1891963

1112111131010201711215111
0.020.030.030.050.020.010.020.020.070.030.010.020.010.020.030.160.10.020.010.060.010.010.01
species
173707

0.020.060.010.030.010.010.020.0200.010.020.020.010.01
10201101100000000001011
species
1429759

species
12348
000001010000000011
0.010.020.09

species
1500757
0001054
0.020.040.1

1147042
species
0.070.070.060.070.10.030.10.10.050.040.10.050.030.070.190.030.120.020.030.10.040.08
4323444421520412141022746

species
1821537
0.020.030.01
00000000010000000000000

0.010.010.010.030.01
00000000000000000000
species
1327939

10000000000000000010201
0.030.010.010.010.030.01
species
1262516

species
181007
00003100100001000000001
0.070.010.020.010.010.010.01

00000000000000000
000.020000
species
432198

0.280.080.080.060.130.150.10.080.240.080.150.180.020.070.050.120.090.060.020.020.030.050.04
14322518431027801001311411142
species
1476886

1647472
species
0.04000.030.02
00100000000000003000001

1606763
species
0.0100.0200.0300.010
00000000001000000200000

100012000000031000511
0.020.020.020.020.070.060.010.01
424716
species

00101100001001
0.030.020.010.020.01
1089121
species

0.020.050.020.010.01
010000021000
1868596
species

00000000000001
00.010.010.0100.0100.01
1445728
species

1567475
species
0.010.020.020.010
00000000001000000000000

279303
species
11000300010000000101
0.020.020.030.030.010.01

1674926
species
00000000020000000000000
0.0200.070.030

species
1245814
0.030.010.010.010.020.010.030.010.020.140.0100.0200
010001010001000150100100

0.020.010.020000
100000100000000000000
species
387905

0.020.030.020.020.01
0111000000100000001
686439
species

668618
species
0.0100.020.040.0400.1000.010.02
000000010011000110000101

1555201
species
0.010.020.010.0100.01
0001000000000000000

0.030.030.50.350.150.10.030.030.030.420.320.150.450.320.150.110.060.080.030.09
1100214164011110452493812105536
species
1871690

000001000000010010001
0.010.010.010.01
1647391
species

151534
species
0.040.060.030.050.020.060.040.060.010.040.010.010.01
23121700002302005001011

species
1029988
3323251322241252512801047105
0.070.070.090.080.050.040.030.080.050.080.040.090.050.010.380.231.080.120.060.090.110.08

species
1887656
0.010.030.020.030.020.010.010.010.020.010.0200.01
01010100001000000000101

52979
species
41110044931000000582143
0.070.020.030.020.10.10.210.090.020.060.090.020.010.040.04

00000000101000000001
0.020.020.01
1150990
species

00000000001101000000001
00.030.020.01000.01
species
1498220

species
673376
0.010.020.060.080.090.070.040.040.130.080.030.020.050.030.050.010.040.030.030.020.070.06
01133711521113051322164

21010000001
0.040.020.020.02
species
45764

species
1327036
00000001002021000000011
00.020.040.080.010.010.01

0.050.040.090.010.030.020.040.010.060.020.030.010.010.040.01000.030.050.020.010.06
22201210201005000024114
species
1340836

11002100000000030112141
0.030.030.010.050.010.010.010.0100.020.0200.0300.010.020.020.020.050.02
1327947
species

0.020.050.020.030.040.010.020.010.04
01020200010202021003
1561065
species

0.020.120.10.020.150.040.010.020.020.020.010.01
100051216000201002022011
764562
species

species
1838070
0.010.010.0200.01000.03
0000000000000000000003

0.010.010.140.080.030.020.030.020.03000.010.010.01
00430300111100000000011
1100814
species

1667322
species
00000100001000020025002
0.010.020.020.020.060.03

species
578234
0.020.020.020.020.020.010.010.050.010.030.010.03
10001001001102000141212

330398
species
0.060.010.020.080.140.060.010.130.010.010.01
3000011000000112710111011

0.010.0200.020.020.020.020
00010000101002000020000
species
1838072

00000000111105010032
0.020.030.020.020.040.010.030.02
species
1229751

10233541110211231265572345
0.020.060.070.070.040.10.270.020.040.020.490.220.140.050.460.090.020.040.050.05
157924
species

22010410101200000001
0.040.040.020.030.020.020.020.040.01
370623
species

1195068
species
0.020.010.020.020.010.010.060.0100.020.010.040.010.010.01
00001200010000000113110

0.010.01
000000000
1327937
species

0.0200.010
00000000100000010
species
1837218

1821541
species
0.020.010.010.01
00000000000000010000000

000000000000
0.01
1340827
species

00.020.0300.040.030.010.070.010.040.010.020.010.02
00010300000014017031111
species
320843

0.050.040.060.090.170.090.010.180.090.150.210.070.030.760.040.020.010.010.01
22247110073810090390302110
species
1589274

0.010.080.020.010.050.030.010.010.010.010.01
00200200011000000110011
species
1647306

20001800000301024012101
0.040.020.070.060.010.020.030.010.020.010.010.01
species
1147044

1327777
species
0000000001
0.0100.03

151599
species
0.020.040.110.020.020.060.050.080.090.020.010.010.020.070.01
12311702330101012600001

species
330397
11213400201060001230061
0.020.020.060.020.070.030.050.020.240.010.020.030.010.070.01

0.040.010.010.030.010.0100.020.080.0300.05000.040.020.01
10010000000124006000310
1385385
species

1555208
species
01330111002100011001103
0.020.10.070.010.020.020.040.020.010.010.010.010.04

51014501741304045034474
0.090.040.020.030.10.050.010.030.180.120.030.060.030.030.040.050.010.040.060.060.090.05
1042122
species

30030001361700000000103
0.060.070.020.840.510.010.04
species
691966

species
573591
0.020.03
0000000011

1465613
species
00000100000000000001100
0.010.010.010.010.010.010.010.0100.0100.010.020.010.01

species
1458669
0100011220000001
0.020.010.020.050.050.01

species
37105
0.020.020.050.10.060.070.050.060.040.050.090.480.040.080.010.090.090.15
1102473202220137410017714

1527511
species
0.020.0200.01
0000100010000000001

0.040.010.03
000000000000100100002
1690817
species

0000000000000201000
0.010.010.010.0100.020.0100
species
649688

species
1527515
000002010000000000031
0.020.020.040.01

species
1446489
10001212100209012112161
0.030.010.010.010.030.020.030.060.040.010.060.070.020.020.020.020.030.010.070.02

0.10.020.030.020.060.010.010.01
0000012001113000010011
species
1109712

0.040.01
00000000000010001
1636271
species

0.010.010.010.01
00000100000000001001001
1282995
species

1567485
species
0000000010000100001
0.020.010.01

28358
species
001001000000165811
0.030.010.040.040.350.070.010.01

764561
species
0.070.020.020.010.120.01
00003000000003021101

species
1445727
0000000000100000000000
0.010.010.020.01

species
947980
0.020.030.10.050.10.040.080.040.040.060.070.050.660.080.020.01
011046413002060784053621

240505023435170177221433
0.040.080.020.120.050.050.080.120.060.110.040.050.160.060.030.020.010.050.040.05
species
1887641

species
1177632
6510040000000601000102
0.110.10.030.030.040.010.010.02

0.010.01
00000000
species
1223259

1476888
species
0.050.070.070.010.180.070.120.120.080.120.020.070.060.180.150.160.271.140.020.030.040.040.04
232078543413124217329312232

691963
species
0.020.020.060.020.010.020.020.060.10.020.03
11210101103500000000022

1126885
species
0.040.030.090.050.020.020.130.060.020.020.080.040.030.020.010.010.040.010.03
02142210521125002113012

1654890
species
0.020.03
0000001001

00.040.0200.010.020.020.010.020.010.040.0200.040
02001000001000021301030
species
1118063

00000000000100001000000
0.020.0100.030.020.020.0100000
591487
species

1458726
species
0.010.010.010.020.020.010.140.0100.020
00000101000000015010010

000001010000000000412004
0.010.020.020.010.0200.010.050.1500.06
1340819
species

0.040.070.070.020.040.060.090.040.080.10.060.070.070.030.180.10.090.140.150.06
23211703104508173158710144
species
1206557

00000000000001000000
00.010.010.010
1551711
species

species
384848
00241801302323031256202
0.060.090.020.070.020.070.040.060.080.020.030.010.020.060.070.030.03

30023211211211033123212
0.060.050.070.020.020.020.050.030.020.040.040.010.030.030.010.020.040.030.010.03
936152
species

1524882
species
0.020.030.050.020.040.020.040.090.080.010.150.030.020.010.040.040.020.05
110215011304010174213323

00100200100000000000001
0.010.030.020.010.020.01000.01
species
1383059

0000000001300000001101
0.030.060.010.010.01
species
437329

0.010.020.010.010.01
000001000001010011
species
1567453

species
1887643
00000000100000000000000
0.020.010.020.030.010000.010000

0.020.020.010.040.020.030.020.0600.010
11000410011300000000100
1647473
species

0.020.010.010.020.020.0200.010
00000000000000000010010
1458714
species

species
445516
000.010.010.01
00000000000001011

00.020.0200.010.030.020.020000
00000000000100000000000
species
546184

1340830
species
0200020030000000010001
0.040.020.0100.080.0200.0100.01

species
1647442
0.02
0001

0.040.020.02
0000.500.500000000000000.5
species
354259
0.020.010.01

no rank
382685
00010100000000000001
0.020.010.01

species
1873891
00000000000000000
0.010

00000000000000000100000
0.020.0100.020
1868170
species

114416
species
0.020.010.050.050.030.020.020.030.020.020.01
00010122010103003100021

species
1437839
0.020.060.020.09
00000200000300000000107

874267
species
00.020.010.010.040.01
000000000010000000020

species
1527493
0.020.040.050.020.030.020.020.030.060.060.040.030.010.010.01
12021401113314000000101

15000500220103000010200
0.020.110.040.010.050.060.020.020.0300.0100.030
1327770
species

0.030.020.020.020.010.020.060.180.070.030.020.010.030.01
10000210001302510410121
species
151537

species
1412875
0.220.090.10.080.120.170.160.050.080.050.070.060.050.090.070.070.030.030.160.260.040.270.07
11433520623133112183214213245

0.020.010.020.01
01000101000001
species
1414737

1556290
species
3422315162435040512281199
0.060.080.060.050.070.130.020.150.050.120.060.10.030.050.10.020.090.010.010.10.12

129009
species
0001000000000001000000
0.010.0200.0300.010.010.010.010

0.040.010.0100.01
20000000000000000000000
1262520
species

010001000000031
0.020.010.020.07
species
1821551

species
1445714
0.050.060.010.010.020.010.010.010.010.020.010.010.090.01
00100600000001002007000

species
1505026
20102100001112001400111
0.050.010.030.010.050.010.010.010.020.020.060.010.010.050.010.010.010.02

species
1168612
000012500000013290024207
0.010.020.020.140.090.140.080.030.050.030.09

0010000000100000100000
0.030.010.010.030.0100.010
species
1567473

1476887
species
0.040.050.050.060.040.030.040.010.040.050.010.020.060.070.050.090.050.040.010.020.040.050.04
2212131011001100105301252

species
1458841
11020211010000031000004
0.020.030.050.020.020.020.030.010.030.01000.05

species
1567468
00000
0.01

species
1654889
0.01
00000000000001

0.010.010.130.020.020.030.040.030.03
004000011102000040002
species
1647305

0.020.020.020.020.030.020.020.010.090.020.060.01
10011200011102000702501
species
547228

species
146269
0.030.050.140.010.070.050.020.060.020.040.180.010.020.030.330.010.010.03
000.51311.752120.51013.600.5201.52700.51
0.060.10.280.010.080.050.020.060.040.060.250.020.020.060.330.010.020.04

001260000011010010030011
0.030.050.140.010.020.020.070.010.030.010.01
279281
no rank

0.0200.020.020.02000.01
01000001000000000000000
species
1543203

0.020.02
00000000001
1527512
species

1647471
species
0.020.020.010.050.010.01
01010100200000001000001

0.010.020.020.020.040.010.010.030.010.030.010.03
00000001100001013100212
151536
species

50011173011412161401620525510
0.10.010.040.260.060.090.260.120.020.040.650.10.150.020.060.020.060.060.13
1327970
species

species
565140
0.0100.010.010.01
00000000000000000000001

0.040.010.0100.010.01
20000000000000000010000
1161930
species

0111010000000504
0.020.030.020.010.040.04
species
1429904

0.030.020.020.020.030.020.0200.04
00100001000103032200003
species
1129145

species
1821563
0.030.020.010.020.020.0400.0400.010.020.020.020
00100000010200000001110

1120011012111000004
0.020.020.060.010.020.020.060.020.020.040.05
species
1664246

00000200000100000000001
0.0100.010.0200.010.010.03000.01
species
1821558

0.130.130.080.050.150.090.210.110.120.060.060.090.090.080.070.170.110.080.170.250.10.10.18
662261184523421111812614207913
1560313
species

species
1815972
10221304111009000012106
0.020.090.050.020.030.010.110.020.030.020.06000.010.020.010.08

0.010.020.010.01
0001000000000000000
1527520
species

1069378
species
0.010.02
000001001

species
1567005
00110000000000000011
00.030.0200.010.020.010.010.010.02

00000000000001000000000
0.02000.010.010.030
species
1690425

species
1204524
0010020000110101001
0.030.020.020.020.010.010.01

0.010.020.010.020.010.01
0000000010100200000000
1486469
species

190478
species
0.040.030.020.040.040.010.030.030.050.070.030.030.08
20101000000211034406236

0.020.030.020.010.020.010.020.01
10101100000100000000121
1838075
species

species
387909
00000000000000000010
0.010.020.010.010.01000.020

0000000000000000000001
0.01
1147140
species

00010000000014000011103
0.010.020.040.0300.010.010.010.04
330396
species

1458697
species
0.020.020.020.020.020.030.050.01
10000010001003023400001

0.020.030.140.040.010.010.010.020.0100.020.01
01106400000001000010
species
330399

species
1622193
01121200000000000000001
0.020.030.050.020.020.01

10001010100000000000000
0.020.020.020.030.020.030.010000.01000
species
1821553

species
438780
221161622105526035133601
0.040.040.030.020.140.140.050.050.020.10.10.080.040.030.050.010.030.040.080.010.01

0.010.020.020.050.0200.10.010.010.01
00001012001000101000101
403905
species

species
1430411
1
0.02

species
1262537
0.070.030.040.040.040.010.020.010.010.010.04
00000031002210012001013

1340833
species
000000
00

1458717
species
0.110.020.010.050.020.010.020.040.010.040.110.020.020.05
003011210020115002100004

0.030.0200.010.020
00100000000000001002000
1076630
species

0.020.010.030.050.030.050.030.01000.010.010.03
101200120100000000002
species
1542912

0.130.040.050.030.050.050.050.060.010.050.010.070.010.01
72002422220002050106101
species
765765

00000100001001
0.010.020.010.010.020.01
species
319522

1887644
species
00000000001
0.03

00.010.01000
00000000000000000
species
561998

0.050.110.070.030.060.070.020.060.050.010.020.020.010.020.010.020.010.040.04
25031630122003021101143
1838082
species

species
1838066
00000001000001101010001
0.020.010.070.010.010.01

0.020.010.010.010.01
0100000000000101000001
347324
species

10130425111214011031348
0.020.010.030.070.030.050.130.030.030.020.040.070.030.010.010.030.020.040.040.11
species
1464127

10000000000101001004110
0.0200.020.010.030.010.0100.060.010.010.01
1032362
species

1070990
species
0.0100.010.010.020.010.01
0000000000100000000

00000000000000000000001
0.010.01
1548918
species

0.040.050.10.090.10.050.050.010.030.070.120.040.030.030.020.070.050.01
22334502013605003031540
species
1647400

species
1821552
0.020.04
00010000000000005

000012120020211155035365
0.020.020.020.050.040.080.010.070.140.040.030.060.040.070.07
864178
species

1465618
species
0.010.0100.010
000000000000000000

00000051000001124011102
0.120.020.010.070.020.030.010.010.010.03
294631
species

00.010.010.010.010000
000000000000000000000
1308897
species

0.130.040.190.050.070.10.10.120.090.030.150.10.040.050.030.050.060.080.020.080.120.11
7262312454185170365726118
320836
species

100000211000000011391155001
0.020.050.020.029.551.410.060.01
1327037
species

1204540
species
0.020.020.020.010.050.010.080.010.070.010.010.050.080.020.080.030.11
11010120000401111472628

0.020.060.020.010.020.040.040.060.020.040.020.050.04
01210201100303000030143
1327035
species

species
1887650
00020000000001010000001
0.050.010.010.01

372457
species
01000101000000000330000
0.030.010.010.010.040.010.0200.040.030.01

00100000021100000100000
0.010.030.010.070.030.0200.010
1647300
species

1450169
species
0.010.010.05
000000000000000000004

10002100100000000101010
0.020.050.010.020.030.02000.020.020.010
species
1582150

00000000001010000000104
0.020.040.010.05
560178
species

0.020.070.020.020.280.022.50.03
0100000300110040220400002
294382
species

0.020.030.040.010.030.090.010.690.020.150.170.0100.02
1100100000102110218130101
species
151535

species
1223260
0.01
000

0.010.020.020.010.030.01
00010001001001
1821723
species

5977519736511610705751399176
0.090.190.220.180.120.160.170.070.140.150.210.120.40.050.050.060.060.150.110.120.190.08
species
1308863

0.020.010.01
00000000000000000000
1340822
species

species
1262535
0.020.0200.020.0200.01
000110000001000200001

1821565
species
00000000000000000000000
0.020.010.0100.010.010.010.01000

0.010.020.070.040.0200.01
0001351000000000000001
1383055
species

species
40522
0.020.020.020.010.020.040.0100.040.01
0000001110000500000031

1481224
species
00000000000000000000000
0000.010

species
1327760
111325001000240100114000
0.020.030.030.070.050.0400.030.0100.970.070.010.010.060.01

1109711
species
0.040.030.020.150.020.01
0211000005100001

species
1458724
01000000000004
0.020.03

1458856
species
0.030.020.010.020.040.020.020.010.020.010.01
10010100000112002001101

700512
species
0.030.010.020.01
0010010000100000000001

00000201122001000000001
0.020.020.020.060.040.010.01
1821560
species

species
416419
1000000000100100000001
0.020.020.010.01

species
1887646
00100110011000013000001
0.030.010.020.010.030.020.010.030.01

0.020.040.01
0000100000000601
310539
species

0.150.220.050.010.080.150.060.020.30.270.080.050.060.070.080.150.040.090.160.020.040.320.05
81010317201294219116471412293
1476889
species

0.020.010.010.010.010.02
00000000000001000000001
1857099
species

000000200000000000001
0.0100.070.0100.02
1399949
species

01000200000000000000000
0.020.020.020.020.0100.010.01
species
1327759

00000200001
0.020.02
species
1873889

0.010.020
0000000000010000000
species
1647427

0.020.020.020.040.06
01000001000118
species
1573458

300304003314521110721241311
0.060.070.030.070.090.020.080.20.150.070.970.180.290.010.040.010.01
1675317
species

00000000000000000000000
0000.010
1223261
species

1510440
species
300121001011112215254103
0.070.020.050.010.020.020.020.040.010.140.020.010.060.020.070.050.110.04

species
1591072
0.020.02000.010.020.0100.01
00000000100000000120001

00000000000000010000112
00.0100.020.010.03
1262515
species

1051631
species
0.010.070.020.020.020.020.950.520.210.320.330.270.060.010.03
00000131101123733353922512

1674930
species
0000020000000000212001
00.020.010.010.0100.020.010.0300.01

species
1838077
0.010.010.0200.0100
0000000000000000

1541819
species
0.010.050.020.030.090.240.050.020.010.030.020.13
020103392010000000200210

00
000000
species
1647311

1536598
species
00.010.0100.010.020.020.01
000000000010

001010000000000002
0.030.020.02
species
1414742

0433012621300111012000002
0.080.10.070.010.050.150.050.030.060.010.070.090.10.03
1837862
species

0.040.070.050.010.020.050.020.040.010.030.01
02032101201200001000201
753085
species

1610509
species
0.010.020.020.030.010.01
00000200000101000000001

03000200101102001001011
0.010.070.010.020.010.020.030.030.030.040.0200.010.0100.020.010.010.02
species
1887645

species
641487
0.060.10.10.090.050.110.020.050.070.030.130.080.310.090.060.050.060.150.030.040.1
35342131231740430974512247

1647421
species
00000000000
0

1675555
species
00000000
0.01

1527513
species
0.020.020.010.010.010.050.050.010.0100.010.01
000001020100000000001

species
1211640
1113913626311411131621412610171110811
0.210.290.320.310.150.220.090.280.090.340.260.320.080.10.070.020.060.130.20.150.130.090.15

0.020.020.040.010.030.010.010.020.010.010.03
00001011010001000121012
species
1647313

0.020.0200.0200.020.020.02
000000.500000000000000000.5
701769
0.010.0100.0100.010.010.01
no rank

0.010.0100.0100.010.010.01
00000100000000000000001
1034149
species

species
35241
1210094041120301243426143
0.020.060.030.020.080.10.090.030.030.050.020.020.110.030.040.050.030.080.160.04

0.0100.010.010.030.0100.0200
0000000001000001000000
1262521
species

0.010.020.020.010.010.010.010.010.020.0100.01
00000000001000000000000
1821556
species

242861
species
00000110000000001000001
0.010.020.010.01

1567466
species
0.020.01
0100

species
767806
0.010.020.030.020.050.080.020.0400.040.0300.020.010.02
01101500000023040320101

210163001000001300120201
0.040.020.020.140.260.020.090.010.020.020.01
species
1239381

561999
species
01000402000100000
00.030.010.030.0600.0200

1308896
species
000000000000000000100
0.010.020.010.010.01000.010.0200.01

0.080.060.040.020.020.020.020.480.060.420.050.160.030.020.0100.020
000.50.500.5000000321.50.84.50.3333330.5000.50
species
0.040.030.020.010.010.010.010.240.030.210.030.080.020.010.0100.010
861046

0011010000006432911001
0.040.030.020.010.010.010.010.240.030.210.020.080.010.010.01
no rank
1429791

981330
species
0.020.020.020.030.020.040.010.020.010.01
01001200011202021000001

species
71366
0.10.01
000000000000000111

1327948
species
0.030.030.010.050.010.010.010.0100.020.0200.0300.010.020.020.020.050.02
11002100000000030112141

663241
species
0.040.020.050.030.050.030.020.040.080.010.010.010.030.050.06
2100242100122200113035

00010000001101
0.020.010.020.020.020.01
species
504346

0.010.010.01
000000000000
1883200
species

species
1486428
0.050.160.060.060.020.160.370.120.110.10.020.010.10.010.170.040.090.120.060.06
2712026155351011011437954

0000010000000000000
0.010.010.010.010.01
species
1168595

1541823
species
0.0100.020.01
00000100000101

species
1071177
0.020.020.020.10.050.020.010.05
1100024200100000014

1086749
species
00000000000000000000000
0.010.0100.01

31000202110440021110101
0.060.030.020.010.060.020.030.010.080.160.020.010.010.020.020.03
species
1821566

0.030.030.040.010.030.020.030.010.040.020.010.020.0100.020.030.010.010.02
11101110100002000121011
1873696
species

00000000002
0.04
255431
species

0.010.030.010.020.010.010.03
00100100001000013
754055
species

0.040.210.080.070.010.010.170.030.07000.050.020.040.01
02630800051300000041300
species
1262532

1406793
species
101100100141100000301
0.020.030.020.020.030.080.020.040.030.01

1871678
species
2155773224211034341268981486
0.040.030.190.120.180.070.090.070.060.130.050.030.430.250.220.380.220.10.110.110.180.090.08

0.020.060.020.020.01
10210000000100000001
species
172667

species
1354510
0.010.020.020.010.010.020.030.01
000000000010

308911
0.010.010
genus
0.020.020
0000000000000000.50000.500

0.010.010
0000000000000001000100
40631
species

00000000000000000
00.030.030.03000
genus
00.010.010.01000
1623294

1982366
00.010.010.01000
species
00.020.020.02000
00000000000000000

no rank
373410
00.010.010.01000
00000000000000000

genus
308912
0.060.10.070.020.040.020.020.060.020.060.030.010.020.010.010.010.010.01
00.8333331.51.50.52.50.500.510.51.50200.500.40.500.50.50.5
0.120.20.140.040.080.040.040.120.040.120.060.020.050.020.020.020.020.02

010000000000000100
0.020.010.01
species
225588

species
10719
0.040.10.070.020.040.020.020.060.020.060.0300.020.010.010.010.010.01
02331510121304000110111

no rank
79205
0.610.661.20.60.870.950.670.660.720.860.770.850.990.450.710.880.731.50.930.760.650.740.74
1.241.32.381.181.731.911.321.281.431.731.571.71.970.911.421.771.4831.851.481.281.461.48
1.467742.969233.319333.669491.988449.015714.151522.06251.636361.959544.057327.194123.720813.483520.5633813.84759.1621622.495.637846.513515.421874.143844

1000090101010111505011
0.020.080.020.030.020.010.070.010.040.060.010.01
382262
species

573174
species
220139652201011030045548
0.040.040.010.020.070.080.150.120.050.060.020.010.070.260.050.060.060.040.11

00020201101401000050510
0.010.020.020.060.020.020.020.020.030.020.030.080.010.030.0600.070.010.01
species
756277

species
573172
6211130200301113248175221021
0.120.050.030.020.020.030.050.010.020.070.450.10.140.440.140.640.020.010.030.01

00100000010001
0.030.030.01
754043
species

754067
species
000003001011
0.030.020.020.02

75590
species
00000000110003011040002
0.020.030.020.010.010.050.03

species
889951
0.020.010.020.040.010.01
000101001000000400100

320831
species
0.010.020.01
0000010000000000002001

0001000110000111000121
0.020.020.020.010.070.010.010.030.01
889954
species

1131316
species
020000002100010711
0.040.050.030.010.060.010.01

1449437
species
000000001100000002
0.010.020.030.020.02

0.020.030.040.05
10100500000000000000004
species
754044

10100000010001000001
0.020.030.030.010.01
756276
species

0.090.260.470.310.120.330.330.20.190.280.320.480.110.080.10.070.050.090.30.320.280.180.23
412141353913889162421117572626211617
536474
species

01111000030000020000111
0.020.030.020.020.090.020.010.010.01
species
754066

754058
species
0.040.020.10.020.020.020.040.020.010.010.010.030.050.010.050.03
21301201002102010134152

1432848
species
00000200000101
0.020.020.020.01

323071431303279015184444122
0.060.040.10.170.120.070.020.070.060.040.280.070.140.150.050.050.050.060.140.04
1567014
species

1536592
species
0.010.020.07
000001010000001

889953
species
0.070.01
0000300000000001

00.0200.010.01
00000000000000000000001
648017
species

573171
species
1111001020210301001102
0.020.020.030.020.020.050.040.020.020.010.010.010.02

0.030.020.010.030.040.070.010.010.010.020.010.01
00110100010200111012101
1325731
species

species
1432847
1010000010000200000001
0.020.030.0100.040.010.010.0100.01

1435411
species
0000000020000100000111
0.0100.060.010.010.0100.010.010.01

10310110002000000061155
0.020.10.020.010.020.040.070.010.010.050.07
species
347327

1567487
species
0.020.01
000010000000000001

749413
species
24202000020203003012022
0.040.080.060.050.060.040.020.030.010.020.020.03

0.07
000000000000001
754074
species

754045
species
0.010.020.020.010.01
000100001000000000

754051
species
0.020.020.110.040.050.010.030.090.030.010.010.020.020.010.070.03
10004402014100001121162

species
1567011
200047010200100305255101
0.040.10.060.020.060.040.030.640.060.060.010.01

species
573173
0.050.010.020.010.020.070.080.0100.010.020.040.010.050.01
20000010003021001023141

1526550
species
0.010.030.010.010.010.01
001001000000020000

1176423
species
0.020.010.01
0100000000000101

0.030.010.070.030.020.050.010.010.030.070.020.020.05
00103312000001013062104
382273
species

423358
0.040.030.020.050.030.150.020.030.010.030.010.020.020.040.040.020.07
family
0.120.10.060.140.10.440.060.10.040.090.030.060.060.10.120.060.21
0.333333000.3333330.2857140.80.95454500.33333300000.500.5555560.3333330.3333330.6666670.50.5833330.3333330.857143

0.020.030.020.050.030.150.020.030.010.010.020.020.010.040.010.07
1123963
no rank
0.040.070.040.090.070.290.040.070.030.020.040.040.020.080.020.14
0.5000.50.4444441.142861.4482800.500000.66666700.500.510.50.8750.51.28571

285013
species
10001230100001000120103
0.020.020.020.020.070.020.020.010.010.020.010.04

00011230000001010101212
0.020.020.020.020.070.020.010.010.010.010.030.010.03
1123964
species

0.040.040.020.040.02
0.50000000000000010.500100.5
0.020.020.010.020.01
680194
genus

0.020.020.010.020.01
1000000000000002100201
315953
species

family
0.030.010.01
292638
000.33333300000000000.3333330000.333333
0.090.030.03

0.060.020.02
000.500000000000.50000.5
genus
564645
0.030.010.01

species
292639
00100000000001
0.030.01

270161
species
0.01
000000000000000001

family
196937
0.090.040.140.050.110.150.050.060.020.040.040.070.970.060.10.020.060.090.020.050.17
0.270.120.420.150.330.450.150.180.060.120.120.212.910.180.30.060.180.270.060.150.51
1.666670.666667020.6666674.3333320.66666700.6666670.3333330.6666670.3333333.333334.66667240.6666671.666672.333330.3333331.333334

249585
0.090.040.140.050.110.150.050.060.020.040.040.070.970.060.10.020.060.090.020.050.17
genus
0.180.080.280.10.220.30.10.120.040.080.080.141.940.120.20.040.120.180.040.10.34
2.510316.531010.510.5573612.53.50.526

5206213620212110146122571412
0.090.040.140.050.110.150.050.060.020.040.040.070.970.060.10.020.060.090.020.050.17
species
342409

10.8372090.3333330.6764710.8051952.684782.666670.71.177220.5652170.840581.4137903.846940.53.051021.890911.652.072461.169011.166672.593751.65625
0.460.430.090.340.770.920.960.50.790.690.690.580.980.840.980.550.60.690.710.420.960.96
1511852
0.130.130.030.090.240.280.320.170.260.210.210.180.280.280.280.170.20.230.220.140.320.29
family

1110703
0.060.060.030.020.190.20.270.120.070.090.10.020.060.210.110.10.150.150.140.080.130.19
no rank
1.51.50.50.51.368424.94.666671.086961.50.8333330.90.501.923080.83333334.13.12.92.148151.18753.076922.94737
0.120.120.060.040.380.40.540.230.140.180.20.040.130.420.220.20.30.30.270.160.260.38

1654582
species
00003911022004181313123
0.070.080.020.020.060.040.030.070.070.010.040.010.040.010.020.04

0.060.060.030.020.050.10.250.070.070.030.020.020.030.140.030.080.10.080.070.040.090.11
33112121033111042310876388
1546257
species

1529056
species
0.070.020.020.040.010.010.010.010.060.020.030.020.04
00003201002001011152223

1511854
0.070.060.070.050.060.020.020.060.040.040.150.140.030.020.050.020.07
no rank
1.333331010.6666672.333330.33333300.3333330.6666670.6666670.6666670705.3333310.66666701.3333300.6666671.66667
0.210.180.210.150.180.060.060.180.120.120.450.420.090.060.150.060.21

29250
0.070.060.070.050.060.020.020.060.040.040.150.140.030.020.050.020.07
species
21.501.513.50.500.5111010.5081.5102012.5
0.140.120.140.10.120.040.040.120.080.080.30.280.060.040.10.040.14

1128424
no rank
4303271012220210163204025
0.070.060.070.050.060.020.020.060.040.040.150.140.030.020.050.020.07

0.030.020.050.160.060.080.120.060.070.030.040.020.080.040.060.160.04
1511853
no rank
000001.50.512.1212112303.083330.51.51.444440.53.50.7142862.55.156251.5
0.060.040.10.330.120.160.240.120.140.060.090.040.160.070.120.320.08

000003122046021321725123
0.030.020.050.050.080.120.010.070.030.020.010.080.020.060.130.04
432587
species

species
92521
0000000052000700310103
0.120.060.050.030.010.010.03

family
573053
0.01
0.03
00000000000000.333333

0.02
00000000000000.5
573054
0.01
genus

437444
species
00000000000001
0.01

352476
0.070.03
genus
0.140.06
000000000000000.5000001

0.070.03
000000000000001000002
128708
species

10240
3.411.681.81.524.662.753.912.73.31.91.551.734.212.0410.582.441.592.814.894.897.435.224.05
family
1.985610.9625360.6548430.7814992.6972.917162.44791.82671.724190.727161.020991.146217.363644.2868719.87354.767873.034233.211546.359425.9789230.683811.25034.26137
14.596.947.329999999999996.2699999999999919.6711.7116.711.3113.858.099999999999996.669999999999997.2517.278.6799999999999943.4910.216.7211.4421.3420.431.9122.3317.37

2.121581.342860.7711170.9161493.988383.119053.25632.493951.736460.6715541.494191.4221611.69653.1815531.7322.612434.348683.457533.939533.571671.995862.939852.25709
6.583.853.673.2210.335.466.749999999999994.959999999999995.543.413.443.3410.713.3626.983.383.045.186.449999999999995.859999999999994.835.325.64
21.231.191.033.31.712.141.611.781.051.081.053.481.068.851.040.971.672.051.911.51.691.73
10241
subfamily

0.60.120.180.360.030.150.060.210.360.240.120.120.150.090.150.270.120.30.210.39
3.333330.6666670.66666701.666670.3333330.6666670.33333311.333331.333330.66666701.6666701.666671.333331.333332.3333312.3333323.33333
573055
0.20.040.060.120.010.050.020.070.120.080.040.040.050.030.050.090.040.10.070.13
genus

0.40.080.120.240.020.10.040.140.240.160.080.080.10.060.10.180.080.20.140.26
51102.50.510.51.522102.502.5223.51.53.535
no rank
573057
0.20.040.060.120.010.050.020.070.120.080.040.040.050.030.050.090.040.10.070.13

1022051213442050544737610
0.20.040.060.120.010.050.020.070.120.080.040.040.050.030.050.090.040.10.070.13
species
305674

0.50.5115.052636234.152170120.51.7500.510.751.50.510.50.5
0.080.040.120.10.570.240.20.30.460.080.160.080.080.040.040.080.060.040.060.020.02
genus
0.040.020.060.050.290.120.10.150.230.040.080.040.040.020.020.040.030.020.030.010.01
10270

0.020.060.050.260.110.10.150.210.040.080.010.010.010.010.01
1022111346902402010001001
species
10271

0.020.020.020.010.020.040.030.010.020.030.030.010.030.01
1100110010001401223121
species
10273

2.981980.9459461.318180.81253.63756.796610.7735852.51.706910.7096772.291670.7857144.920.93.9215712.01255.666675.607592.634151.764716.235293.57647
1.110.370.440.321.61.180.530.240.580.430.310.480.560.50.70.510.81.260.790.820.681.360.85
0.520.180.220.160.810.590.250.120.30.210.140.240.280.250.350.270.40.620.410.410.340.670.43
10242
genus

0.040.030.070.090.060.010.040.030.140.040.050.170.080.060.140
20103102000001424513064130
species
10243

species
28871
10000200001021162501302
0.030.010.010.020.010.020.080.010.070.050.020.0600.020.040.010.03

10256
species
310231010112200040151422
0.060.020.050.080.090.020.020.040.040.0400.0400.010.060.020.050.020.03

0.070.030.060.030.0400.010.0100.040.040.020.04
00003320100010000102313
species
10245

0.120.040.020.020.060.060.040.0400.0200.020.020.040.02
1.50.50000.50.500.500.50.500000000.500.50
species
10244
0.060.020.010.010.030.030.020.0200.0100.010.010.020.01

0.060.020.010.010.030.030.020.0200.0100.010.010.020.01
31000110101100000001010
no rank
619591

10255
species
00010100000000000200010
0.010.010.020.020.020.010.03000.010.010

120133200016280124350912
0.020.040.020.090.030.050.020.120.080.060.010.020.050.040.070.10.17
28874
species

160796
species
0.040.020.030.020.210.090.020.120.050.060.040.020.040.0500.020.010.180.070.080.140.1
21118111522211600211556127

12643
species
1234210210033010132113318961192
0.220.070.130.050.240.180.010.070.090.010.020.10.140.10.280.230.10.070.020.210.03

20000100000000000401100
0.050.010.020.020.010.0500.020.0200
species
28873

genus
1285599
0.210.160.140.020.150.070.020.060.190.140.040.020.030.030.040.020.020.020.01
0.420.320.280.040.30.140.040.120.380.280.080.040.060.060.080.040.040.040.02
052.530131.50.5153.50.51.501.51.51.511010.5

species
1285600
0105602631210713033322021
0.210.160.140.020.150.070.020.060.190.140.040.020.030.030.040.020.020.020.01

0.51.050.250.3750.72.585370.7741940.280.560.90.3333330.33333301.750.50.8333331.503.650.50.7272730.51
0.040.20.20.160.20.410.310.250.250.30.060.060.080.140.060.060.20.020.110.060.06
10257
0.020.10.10.070.10.190.120.10.120.150.040.020.040.070.030.030.10.010.050.030.03
genus

0101070012000201001011
0.030.020.040.020.070.010.020.040.0600.010.010.010.010.01
10258
species

13100320011004020081212
0.020.060.050.010.010.030.060.020.020.030.020.030.020.090.010.030.010.03
129726
species

0.010.020.030.060.040.0100.020.01
334521
no rank
0000.50210.50000.5000000000.5
0.020.040.060.120.080.0200.040.02

000104210001000000001
0.010.020.030.060.040.0100.020.01
1579460
species

0000260122000010300001
0.010.010.070.060.030.050.0600.070.030.01
species
129727

2.927081.026320.9487181.029412.813083.306453.254551.0252.628210.6363642.911761.333330.5833333.020411.703032.883722.648658.275.711865.834713.161294.311113.74038
0.960.380.390.341.070.621.10.40.780.330.340.360.480.491.650.430.3711.181.210.930.91.04
genus
10260
0.450.190.190.160.520.260.520.20.350.150.170.160.20.220.620.210.170.50.520.580.40.420.48

10264
species
0.030.040.040.170.080.030.010.030.030.070.050.020.290.080.070.10.050.05
121079100100041522376854

43024312002418283315476
0.080.070.010.050.10.030.030.050.040.080.040.060.140.080.030.040.020.060.050.080.08
10261
species

0.190.040.10.070.170.050.150.070.20.060.040.060.020.080.130.220.220.050.160.15
10233766382001802911191841511
species
44088

0.140.020.020.040.040.180.120.020.160.060.080.160.080.820.020.060.020.280.120.280.140.16
2000.50.55101.50.50112.530.520.562.55.533
no rank
0.070.010.010.020.020.090.060.010.080.030.040.080.040.410.010.030.010.140.060.140.070.08
336487

0.070.010.010.020.020.090.060.010.080.030.040.080.040.410.010.030.010.140.060.140.070.08
40011102031022561411251166
648998
species

0.070.020.030.020.070.020.250.050.070.030.130.040.040.040.050.010.020.060.160.050.050.12
4111321023172150612513459
species
336486

00.666667000000.333333000.333333000.33333300000.333333000.3333330.333333
0.120.060.060.030.030.030.03
genus
10278
0.040.020.020.010.010.010.01

0.080.040.040.020.020.020.02
01000000.5000.5000.500000.5000.50.5
species
10279
0.040.020.020.010.010.010.01

10280
no rank
0.040.020.020.010.010.010.01
02000001001001000010011

no rank
0.240.040.160.050.90.120.620.570.30.030.080.142.310.197.680.190.150.110.470.220.230.150.19
40070
0.480.080.320.11.80.241.241.140.60.060.160.284.620.3815.360.380.30.220.950.440.460.290.38
6.50.51.71875114.63334.2512.57.631585.633330.513.526.582310.894755.5105.93.6818211.05265.477275.913044.103455.21053

species
1873698
0.020.030.120.030.620.450.020.040.080.020.030.010.30.050.050.030.03
01105425181020230031264432

species
67082
0.240.020.130.050.780.090.120.280.030.040.142.230.177.680.190.120.10.180.170.180.110.16
13142331005121275524111201481513141012

0.080.340.060.10.20.240.140.10.180.060.160.120.240.20.140.080.080.10.180.620.220.080.22
140.51271.5120.521.51.570.522.52412.5424
0.040.170.030.050.10.120.070.050.090.030.080.060.120.10.070.040.040.050.090.310.110.040.11
2005509
genus

1076255
species
28124143241433141454825848
0.040.170.030.050.10.120.070.050.090.030.080.060.120.10.070.040.040.050.090.310.110.040.11

000.52.50.51.503.50.523.52.50100.5001.520.500.5
0.060.240.040.060.340.040.240.260.20.020.020.060.10.020.02
0.030.120.020.030.170.020.120.130.10.010.010.030.050.010.01
686612
no rank

species
240426
00151307147502010034101
0.030.120.020.030.170.020.120.130.10.010.010.030.050.010.01

genus
10265
0.240.10.10.120.310.190.070.070.090.060.040.040.280.060.140.060.10.070.040.050.120.2
0.630.290.240.310.790.540.190.220.240.160.090.110.690.160.420.140.260.190.090.120.340.59
1.603170.5172410.50.5161291.734182.685190.2105260.2272730.541667000.2727271.130430.87502.119051.285710.9230770.6842110.3333330.4166671.294121.9661

59509
0.070.060.020.030.050.080.020.020.020.020.010.010.130.020.060.010.030.020.010.010.040.08
species
2100.50.54.50000001.51030.510.50.50.51.52.5
0.140.120.040.060.10.160.040.040.040.040.020.020.260.040.120.020.060.040.020.020.080.16

no rank
376849
42011900000032061211135
0.070.060.020.030.050.080.020.020.020.020.010.010.130.020.060.010.030.020.010.010.040.08

186805
0.090.030.020.050.120.080.020.050.030.020.010.010.010.020.070.010.030.030.010.010.050.1
species
2.50.5012400.50.50000103.50.511.50.50.523.5
0.180.060.040.10.240.160.040.10.060.040.020.020.020.040.140.020.060.060.020.020.10.2

51024801100002071231147
0.090.030.020.050.120.080.020.050.030.020.010.010.010.020.070.010.030.030.010.010.050.1
376852
no rank

40216310200133024311132
0.070.010.060.030.140.030.040.010.050.020.010.030.130.020.020.040.040.020.010.030.040.03
10266
species

genus
10275
0.160.070.030.090.020.020.10.070.060.020.120.070.010.010.020.10.090.030.10.04
41.50.520.51201.510.501.500.50.50.514.53.514.51.5
0.320.140.060.180.040.040.20.140.120.040.240.140.020.020.040.20.180.060.20.08

10276
species
0.160.070.030.090.020.020.10.070.060.020.120.070.010.010.020.10.090.030.10.04
83141240321030111297293

genus
0.10.060.030.020.120.040.10.050.090.060.060.040.080.070.060.040.130.10.110.150.040.08
10282
1.035711.50.3333330.3333331.666671.181820.810.8333330.66666710.6666670.6666672.501.823531.181822.593751.61.851852.540540.81.2381
0.280.120.090.060.360.110.250.10.240.180.180.120.240.20.170.110.320.250.270.370.10.21

0.140.060.040.240.060.10.10.120.120.080.160.120.10.040.10.10.10.120.040.1
1.500.50.52.5210111.5114031222212
0.070.030.020.120.030.050.050.060.060.040.080.060.050.020.050.050.050.060.020.05
99000
species

0.070.030.020.120.030.050.050.060.060.040.080.060.050.020.050.050.050.060.020.05
30115420223228062444424
132475
no rank

0.040.060.010.050.050.050.010.010.030.090.050.060.10.020.03
23000122200002013745722
38804
species

subfamily
1.160.420.510.471.330.981.650.921.450.750.40.560.570.91.731.190.560.762.692.765.853.362.19
10284
3.891.321.611.494.623.315.73.214.822.521.441.861.953.045.933.921.922.629.559.0319.4211.367.37
3.29821.121211.130431.29532.560615.063443.242112.392522.933611.321430.93751.698920.8102568.434211.377749.576533.572923.8969511.345510.936949.906320.62328.24152

0.340.160.260.140.280.280.440.10.320.360.160.280.320.480.6210.240.540.460.720.560.780.9
4.5221.5384.513.5323.5217227.57111014.510.51817
no rank
39747
0.170.080.130.070.140.140.220.050.160.180.080.140.160.240.310.50.120.270.230.360.280.390.45

944361692764743445514222029213634
0.170.080.130.070.140.140.220.050.160.180.080.140.160.240.310.50.120.270.230.360.280.390.45
83191
species

2.51112.59.52.525.52.5100.522.57.56.5210.522.51097
0.180.080.120.10.240.320.240.20.520.30.080.080.060.70.280.220.10.481.120.520.40.38
0.090.040.060.050.120.160.120.10.260.150.040.040.030.350.140.110.050.240.560.260.20.19
62098
genus

62099
species
0.090.040.060.050.120.160.120.10.260.150.040.040.030.350.140.110.050.240.560.260.20.19
52225195411520145151342145201814

2.210.660.720.782.771.733.371.992.531.110.81.020.981.62.881.450.91.225.924.4312.496.823.9
4.90951.636361.638892.076923.75096.635844.71813.608044.015811.351351.18752.137250.91836710.851.798615.475864.166673.3360716.67414.248376.709431.765410.9692
genus
10286
0.90.290.320.351.070.681.30.771.030.420.290.420.360.631.070.560.330.442.221.835.312.771.55

0.240.030.080.010.270.120.210.070.20.060.090.080.180.050.520.140.090.180.470.30.70.780.38
1993631
species
6.50.5105.5741.54122233.585720.5122735.514
0.480.060.160.020.540.240.420.140.40.120.180.160.360.11.040.280.180.360.940.61.41.560.76

1312011148382444671610144124547128
0.240.030.080.010.270.120.210.070.20.060.090.080.180.050.520.140.090.180.470.30.70.780.38
no rank
1293539

0.330.110.180.150.270.210.310.250.250.120.070.160.060.280.210.160.080.070.60.463.31.260.54
175561124121010438139318105533725411540
species
10288

31.5015.5896.532.51.51.50.5131.53.50.542614.5391310.5
0.220.120.10.520.280.880.640.280.30.120.120.080.380.420.120.020.21.20.722.020.580.58
species
1179664
0.110.060.050.260.140.440.320.140.150.060.060.040.190.210.060.010.10.60.361.010.290.29

630211161813653312637185229782621
0.110.060.050.260.140.440.320.140.150.060.060.040.190.210.060.010.10.60.361.010.290.29
1293572
no rank

842571292141141426341247102115
0.160.080.060.120.170.10.220.050.330.030.020.080.040.030.140.050.030.050.140.580.140.230.21
species
28321

species
1993630
0.060.020.10.110.120.070.120.060.060.040.040.090.140.130.050.410.120.160.210.13
0.120.040.20.220.240.140.240.120.120.080.080.180.280.260.10.820.240.320.420.26
1.5000.5262.51.52.511.510.5607.57.5218569.55

0.060.020.10.110.120.070.120.060.060.040.040.090.140.130.050.410.120.160.210.13
30014125352321120151543610121910
no rank
1293540

3.070420.3333330.60.50.3333331.947371.235291.386360.63157911.333331.250.83.6666704.914891.8235310.38554.333333.677421.304353.023261.77419
0.710.090.250.040.060.190.340.440.190.270.240.320.40.240.470.170.830.450.620.230.430.31
0.250.030.10.020.020.060.120.170.070.090.080.120.160.080.20.060.390.150.230.080.160.12
40069
no rank

0.060.030.020.010.020.070.020.040.080.130.010.320.090.030.050.05
3011011310022001412607244
species
1680908

0.40.060.120.040.120.20.20.10.180.160.160.160.160.140.10.120.30.30.120.220.14
50.5100.532211.52215.503.532.56.55.5252.5
species
39444
0.20.030.060.020.060.10.10.050.090.080.080.080.080.070.050.060.150.150.060.110.07

0.20.030.060.020.060.10.10.050.090.080.080.080.080.070.050.060.150.150.060.110.07
1012016442344211076513114105
930275
no rank

0.330.080.240.080.080.250.270.080.520.240.420.160.30.160.20.40.510.40.240.190.48
0.2424240.250.50.250.250.60.2222220.250.48076900.4166670.2142860.250.56666700.250.350.451.039220.650.8750.3157890.354167
0.070.020.060.020.020.060.070.020.120.060.10.040.060.040.050.10.130.10.060.050.12
1511857
order

157897
0.040.040.020.020.090.060.060.040.040.030.030.060.060.050.050.020.08
family
0.140.130.060.060.340.180.20.120.150.090.090.180.170.150.150.060.25
0.285714000010.3333330.3333330.67647100.5555560.250.3333330.86666700.3333330.5555560.3333330.6470590.8666671.333330.6666670.56

0.040.040.020.020.070.060.030.040.030.020.020.040.050.040.020.05
187213
genus
0.400001.444440.50.5100.8333330.250.51.3333300.510.51.111111.5010.727273
0.10.090.040.040.210.120.080.080.090.040.040.080.090.080.040.11

0.020.030.020.040.010.040.010.020.020.010.040.020.01
10000310002010012113021
157898
species

0.020.070.020.030.01
351054
no rank
0.500000001.5000.502000000000.5
0.040.140.040.060.02

10000000300104000000001
0.020.070.020.030.01
species
537009

00000201001000010130002
0.020.020.020.0100.020.030.03
157899
species

0.040.060.020.020.020.040.020.020.10.06
000000000.5000.500.500.50.50.50.50.5201
no rank
0.020.030.010.010.010.020.010.010.050.03
1895332

1895333
species
0.020.030.010.010.010.020.010.010.050.03
00000000100101011111402

0.3333330.3333330.6666670.3333330.3333330.3333330.28571400.333333000.33333300.44444400.3333330.333333120.8666670.3333330.250.272727
0.120.060.180.060.060.060.140.060.120.090.030.060.120.210.150.030.080.11
family
10477
0.040.020.060.020.020.020.050.020.040.030.010.020.040.070.050.010.030.04

0.020.010.010.010.07
564646
genus
00.50000.50000000000.50.503
0.040.020.020.020.14

300186
species
0.020.010.010.010.07
0100010000000001106

0.080.120.040.040.020.090.040.080.060.020.080.10.020.050.07
0.5010.50.50.50.44444400.5000.500.666667000.51.501.30.50.40.428571
0.040.060.020.020.010.050.020.040.030.010.040.050.010.030.04
341940
genus

species
346883
0.020.020.020.010.01
10000000100101000000001

0.060.020.020.010.01
0021001000000200000001
512792
species

0.020.01
0000000000010000000001
species
346881

00001100000001000003101
0.020.010.010.040.010.01
species
176106

0.020.020.010.0100.01
10000010000000001001001
species
346884

346882
species
0.040
0000000000000000030000

12.498.199.0611.2513.969.411.987.3311.57.597.248.634.5911.057.476.958.7211.4611.1315.4711.1612.5114.67
22.716615.81210.343317.586719.817332.981915.04519.416116.63228.7944712.020714.28164.0283256.67333.124524.312237.263833.763528.295634.715624.58630.962433.2999
549779
3.582.422.63.223.992.63.422.13.332.212.052.471.343.212.1122.53.323.074.263.063.424.14
family

985780
1.30.741.121.451.541.231.220.751.110.810.920.880.491.240.730.691.071.331.471.961.521.581.61
no rank
3.92.23.364.334.623.693.622.213.312.372.722.641.433.682.192.033.23.934.385.844.554.724.82
2311.136411.666720.150121.66674815.78459.4570115.15417.9620314.911814.66673.4895154.84783.3333323.13340.715633.361341.445251.561637.925347.271238.9378

species
1.30.721.121.431.541.231.190.721.090.750.880.880.451.20.730.651.051.271.441.931.51.561.59
985782
2.61.442.242.863.082.462.381.442.181.51.761.760.92.41.461.32.12.542.883.8633.123.18
34.51717.530.532.5722414.52312.523225.5845366251.5637857.571.559

no rank
1269028
1.30.721.121.431.541.231.190.721.090.750.880.880.451.20.730.651.051.271.441.931.51.561.59
6934356165144482946254644111681072124103126156115143118

1586712
species
0.020.020.020.020.020.060.040.040.040.040.030.060.030.020.030.020.03
01010011122016043532222

1.290.630.480.481.351.111.620.91.320.630.631.020.360.721.230.90.480.691.382.311.472.071.89
7.666673.333331.666672.333336.3333314.33337.3333346.333332.333333.666675.66667111.33332116613.333320.666712.66672115.6667
genus
1513234
0.430.210.160.160.450.370.540.30.440.210.210.340.120.240.410.30.160.230.460.770.490.690.63

species
0.430.210.160.160.450.370.540.30.440.210.210.340.120.240.410.30.160.230.460.770.490.690.63
1513235
11.552.53.59.521.51169.53.55.58.51.517316.59920311931.523.5
0.860.420.320.320.90.741.080.60.880.420.420.680.240.480.820.60.320.460.921.540.981.381.26

0.430.210.160.160.450.370.540.30.440.210.210.340.120.240.410.30.160.230.460.770.490.690.63
23105719432212197111733463318184062386347
no rank
693272

1.861.471.311.61211.661.061.771.190.921.250.731.720.971.011.271.761.151.531.041.151.91
315393
genus
3.722.942.623.22423.322.123.542.381.842.51.463.441.942.022.543.522.33.062.082.33.82
49.53520.5344258.533.52137.519.52431.59121755.575.571.55061.54052.571

species
212035
9970416884117674275394863182421411115114310012380105142
1.861.471.311.61211.661.061.771.190.921.250.731.720.971.011.271.761.151.531.041.151.91

2.648650.7333330.6486490.51.752692.148150.9310340.6666672.076920.51.743591.380950.3333333.227270.51.484851.179492.357143.56142.694441.319151.638891.90805
0.740.30.370.280.930.270.580.31.040.360.390.420.120.660.840.330.390.840.570.720.470.360.87
10482
0.240.10.130.090.310.090.20.10.350.120.130.140.040.220.280.110.130.280.190.240.160.120.29
family

genus
10483
0.040.080.060.050.120.010.120.050.120.090.040.10.140.090.070.210.020.050.050.050.12
0.51.2510.4444441.750.51.7511.086960.8333330103.350.52.611111.785714.2439120.81.71.58333
0.080.160.120.090.240.020.240.10.230.180.080.20.280.180.140.410.040.10.10.10.24

265522
species
0.020.060.020.020.020.030.020.010.010.030.040.04
13010010100004021100243

10204000320001112504112
0.020.060.10.070.060.010.070.010.020.060.050.010.010.03
species
419435

10484
species
0.020.020.020.010.10.050.020.030.040.060.070.060.040.130.020.010.05
010111421102091751120104

genus
0.210.020.060.050.190.080.070.050.230.030.130.10.040.120.140.020.060.070.170.190.10.070.17
10485
0.420.040.120.10.380.160.140.10.460.060.260.20.080.240.280.040.120.150.340.380.210.140.34
4.571430.5113.184213.56250.85714314.152170.52.615382.50.56.0833310.51.751.65.852944.578952.5714333.76471

53988
species
0.020.020.010.020.020.030.110.10.020.010.030.040.020.050.030.05
10001110116503013324204

39640
species
101227822901011421431311669
0.190.020.060.050.170.070.050.050.210.020.040.10.140.010.030.040.150.140.080.070.12

0.2987010.456140.31250.379630.2380950.480.8860760.41250.4727270.250.3076920.8333330.2857140.7945210.6666670.8095240.6716420.6170211.031750.303030.4827590.4897960.206897
0.770.570.961.080.420.50.790.80.550.60.520.240.280.730.420.630.670.470.630.330.580.490.29
0.20.150.260.30.120.120.250.250.160.150.130.080.080.20.140.170.160.140.170.10.160.130.09
10508
family

genus
10509
0.140.080.160.160.10.070.170.170.160.060.070.020.080.10.140.10.140.080.110.070.10.10.07
0.3023260.6666670.4565220.6904760.3333330.6363641.828570.750.6666670.40.260870.50.40.733333110.95555611.657140.4705880.7142860.750.285714
0.430.240.460.420.240.220.350.360.390.150.230.040.20.30.280.30.450.220.350.170.280.280.14

species
1986018
0.020.020.010.01
0.500000000.5000000000000.5
0.040.040.020.02

100000001000000000001
0.020.020.010.01
no rank
1715778

0.060.020.020.02
000.500000000000.5000.5000000.5
0.030.010.010.01
413684
no rank

00100000000000000000001
0.030.01
species
1923976

1324794
species
0.010.01
00000000000001001

species
2015374
0.040.020.010.020.010.01
0.080.040.020.040.020.02
10000.50.50.5000000100000

no rank
1788434
0.040.020.010.020.010.01
2000111000000200000

0.020.02
0000000000000100.5
1986019
0.010.01
species

0.010.01
0000000000000201
no rank
909210

114424
species
0.020.010.020.050.03
0001010121

0.080.040.040.060.10.040.020.02
0100.50.500000000202.51000.500.5
0.040.020.020.030.050.020.010.01
696859
species

0201100000000405200101
0.040.020.020.030.050.020.010.01
573199
no rank

0.020.020.01
129951
species
0.040.040.02
00000000000000000000.5

28285
no rank
00000000000000000001
0.010.010.01

00000000
0.01
no rank
10515

no rank
10533
0.01
0

species
0.04
1986020
0.08
0000000000000.5

no rank
585059
0.04
0000000000001

species
0.01
536079
0.02
0000000000000000.5

0.01
0000000000000001
310540
no rank

00.04
00000000000000001
00.02
1986023
species

1641642
no rank
00000000000000002
00.02

0.060.040.020.040.10.060.040.020.080.06
000.50.500.500.510.50.5000.50000001.51.5
no rank
0.030.020.010.020.050.030.020.010.040.03
346173

0.01
00000000000001
species
10535

0.030.020.010.020.050.030.020.040.03
0011010121100000000033
740971
species

0.040.020.14
00000010000000002.21429
species
108098
0.020.020.07

no rank
565303
0.03
00000000000000001.42857
0.07

0.01
00000000000000001
10522
no rank

no rank
198503
0.03
00000000000000003

no rank
0.010
565302
00000000000000000
0.020

no rank
10519
00000000000000000
0.010

0.010.010.01
242710
species
000000.50000000000000000.50.5
0.020.020.02

00000100000000000000011
0.010.010.01
47680
no rank

129875
species
0.140.010.01
00000000000000200000101

0.070.040.120.070.010.020.01
0.4285710000.5053000001000002001
0.040.020.120.070.010.020.01
130310
species

0.030.02
10001
651580
no rank

species
130308
000000000000000001
0.01

species
129955
0.02
000000000001

0.030.020.010.010.080.010.030.03
1986024
species
000.500000000.5000.5000.503.50.511.5
0.060.040.020.020.160.020.060.06

943083
no rank
0010000000100100107123
0.030.020.010.010.080.010.030.03

0.02
000000001
129953
species

2015375
0.010.010.01
species
0.020.020.02
00000000000000.5000.500

1788435
no rank
0000000000000100100
0.010.010.01

0000000110001
0.020.020.04
species
129954

species
0.010.010.01
1986017
0000000000000000.50000.500.5
0.020.020.02

0000000000000001000101
0.010.010.01
38432
no rank

0.020.040.010.02
000000.500000.500000000.5
species
10537
0.010.020.010.01

0
00000000000000000
10512
no rank

10514
no rank
0000010000100000001
0.010.020.01

0.01
1986021
species
0.02
0000000000000000.5

0000000000000001
0.01
1560346
no rank

0.020.01
00001000000001
1146877
species

0.01
1986022
species
0.02
0000000000000000000.5

no rank
994672
0.01
0000000000000000001

0.02
0001
129956
species

0111.5000000000000
0.080.120.140
species
1146874
0.040.060.070

696069
no rank
0.040.060.070
0223000000000000

0.0200.0200.02
000000000000000000
0.0100.0100.01
1962299
species

000000000000000000
0.0100.0100.01
no rank
995022

0.010.01
00000000000000000000000
species
2015370
0.010.01

1788436
no rank
00000000000000000000000
00

00
00000000000000000000000
no rank
1788433

00000000000000000000000
00
1788432
no rank

0.01
000000000000000001
species
130371

0.030.010.010.05
130369
species
000001.50000000000.50.52
0.060.020.020.1

0.030.010.010.05
000003000000000114
no rank
35265

0000.3333330.3333330.3333330000.33333300000000.5000.58333300.5
0.060.060.030.180.020.120.02
0.020.020.010.060.010.040.01
129876
genus

0.03
691961
species
0000000000.5
0.06

0.03
0000000001
1520002
no rank

00000000000000000100001
0.010.01
298333
species

0000.5000000.500000000001
0.040.060.06
0.020.030.03
298331
species

114102
no rank
0.020.030.03
000100000100000000002

00000.50.5000000000000000.5
0.040.020.02
0.020.010.01
886980
species

000011000000000000001
0.020.010.01
1520004
no rank

10552
0.020.060.060.070.030.050.020.040.070.060.020.020.020.01
genus
0.060.180.180.170.090.110.060.120.170.160.060.050.050.02
0.3333330.5555560.3333330.35294100.5555560.3636360.333333000.666667001.8823501.31250.3333330.40.4000.5

0.080.02
010000000000000000.5
0.040.01
1548417
species

020000000000000001
0.040.01
764030
no rank

0.04
00000000000005
species
190063

species
190064
0000000000000000001
0.01

species
0.020.020.01
1962301
00.500000.5000000000.5
0.040.040.02

1408257
no rank
0.020.020.01
0100001000000001

species
1146875
0.020.030.010.020.040.030.01
0.500.5000.500.5001002000.5
0.040.060.020.040.080.060.02

0.020.030.010.020.040.030.01
10100101002004001
878329
no rank

0.04
000001
species
0.02
1534553

0.02
000002
no rank
1520006

0.020.020.040.010.01
0001001000000004010001
190061
species

0000.5
0.04
no rank
0.02
280050

1907767
species
0001
0.02

0.020.010.01
1193420
species
00000000000000010.500.5
0.040.020.02

1193422
no rank
0.020.010.01
0000000000000002101

0.060.04
000.50.5
0.030.02
190062
species

0011
0.030.02
no rank
172861

genus
100953
0.040.030.050.020.020.050.030.020.060.030.020.030.020.010.020.01
100.50.307692010.250.30769200.250.51.500.66666700010.8333330.3333330.50.3333330.25
0.080.060.130.040.080.130.120.040.120.060.040.060.060.020.060.04

0000.333333000.3333330.33333300.3333330000000000.33333300.3333330.333333
0.060.060.060.090.030.030.03
no rank
327833
0.020.020.020.030.010.010.01

species
130500
0.020.020.020.030.010.010.01
0.040.040.040.060.020.020.02
0000.5000.50.500.50000000000.500.50.5

00010011010000000001011
0.020.020.020.030.010.010.01
no rank
111167

0.030.020.020.060.010.010.01
0011020000030100001001
species
130328

0.040.020.020.010.02
2000000100100200002
130327
species

0.010.020.010.01
000000000000010002011
130499
species

family
0.150.060.060.140.10.240.070.220.140.060.060.10.080.110.070.240.070.140.060.110.280.210.12
43682
0.7843140.4545450.51.148150.6388895.698920.750.7948720.9019610.2857140.4545451.133330.2857141.842110.254.109891.0344820.523811.323532.233012.948051.27907
0.510.220.240.540.360.930.280.780.510.210.220.30.280.380.280.910.290.550.210.341.030.770.43

0.50000101100.520.540100.5031.50.52
0.040.040.10.10.040.160.080.120.040.020.140.080.020.1
genus
0.020.020.050.050.020.080.040.060.020.010.070.040.010.05
1921707

species
158683
10000202201418020106314
0.020.020.050.050.020.080.040.060.020.010.070.040.010.05

genus
0.130.060.060.140.10.220.070.170.090.060.040.020.040.050.070.220.070.130.060.040.240.20.07
43680
1.18750.6250.6666671.550.8846158.0923111.130431.333330.40.6666670.50.3333331.466670.3333335.873021.363642.794870.7333330.3333333.253734.185191.66667
0.320.160.180.40.260.650.210.460.270.150.120.040.120.150.210.630.220.390.150.090.670.540.21

113366
0.020.020.010.010.020.010.010.010.03
species
0.50000000.500000100.510.50.50.51
0.040.040.020.020.040.020.020.020.06

260797
no rank
0.020.020.010.010.020.010.010.010.03
100000010000020121112

species
113370
4101110201010003101144
0.070.020.020.020.010.050.030.020.030.010.010.010.050.04

no rank
328613
0.040.040.060.120.070.210.070.10.090.030.040.040.040.070.180.050.120.030.010.160.150.07
0.080.080.120.240.140.420.140.20.180.060.080.080.080.140.360.10.240.060.020.320.30.14
1112.51.512.51.5220.5100.52.50.5102.54.51.50.5672.5

222532534412015120593112145
0.040.040.060.120.070.210.070.10.090.030.040.040.040.070.180.050.120.030.010.160.150.07
328615
species

0.060.180.090.150.420.270.150.180.180.120.040.090.150.160.10.550.040.03
0.33333310.3333330.6666670.5952381.1111100.66666700.666667010.3333330.25011.066670.81250.51.8181800.250.333333
10656
0.020.060.030.050.120.080.050.060.060.040.010.030.050.050.030.150.010.01
family

0.040.120.060.10.30.190.10.120.120.080.030.060.10.110.070.40.030.02
0.51.50.510.8333331.57895010101.50.50.33333301.51.61.181820.7142862.500.3333330.5
10657
0.020.060.030.050.120.080.050.060.060.040.010.030.050.050.030.150.010.01
genus

03121502020300002324
0.060.030.050.020.040.050.060.060.020.040.020.05
10658
species

no rank
341937
0.070.030.010.010.010.1
0.140.060.020.020.020.2
00001.51.500000000.50000.50.54

307240
species
0.1
00000000000000000008

663237
species
0.070.030.010.010.01
0000330000000100011

species
513550
10001100000010034000001
0.020.020.010.040.030.030.01

species
341938
0.01
0000000000000000000000.5
0.02

0.01
0000000000000000000001
no rank
236750

0.060.120.030.060.060.030.03
000000.666667000000.33333300.33333300.6666670.66666700.333333000.333333
0.020.040.010.020.020.010.01
1511858
family

000001000000.500.501100.5000.5
0.040.080.020.040.040.020.02
1511859
0.020.040.010.020.020.010.01
genus

0000000000010100201001
0.020.010.020.010.01
269145
species

754004
species
0.020.020.02
0000020000010002

548681
1.791.821.491.161.281.630.91.411.190.971.731.041.213.041.272.120.71.20.81.071.082.070.97
order
5.941773.493860.4580470.4312620.7812021.737890.3630140.6617860.4683760.5331950.8625431.124280.99673722.06450.5923574.389990.8108881.735550.6485150.6119690.81226113.41680.687371
7.899999999999998.139999999999997.275.309999999999996.489999999999998.049999999999994.386.829999999999995.854.828.729999999999985.236.1315.046.2810.593.495.709999999999994.045.179999999999995.229.139999999999984.82999999999999

15.98929.874521.138891.252870.4516133.908051.185191.516480.7058821.368421.551721.644440.33333316.42770.3333335.511361.357141.89610.4285710.975611.3384631.56431.83333
2.792.630.720.870.310.870.540.910.510.570.870.450.241.730.210.880.280.770.140.410.653.810.72
family
0.930.880.240.290.110.290.180.310.170.190.290.150.080.570.070.290.090.260.050.140.221.270.24
548682

0.020.060.02
00000000000000.501.500.5
genus
0.010.030.01
172653

10401
species
0.010.030.01
000000000000010301

00.500.50.50.83333300.51.50.50.5001.166670.5101000.50.51.77778
0.040.040.040.060.040.140.060.080.060.140.040.040.020.020.18
0.020.020.020.030.020.070.030.040.030.070.020.020.010.010.09
692605
genus

00001201301001120200013
0.020.020.020.070.020.010.070.020.020.010.04
species
389214

85655
species
01010100011003000000104
0.020.020.010.030.020.020.010.05

1.861.710.480.540.160.520.360.560.20.320.50.30.161.080.490.190.450.090.270.412.520.3
23.983915.17541.708331.981480.756.442311.777782.428570.752.343752.622.466670.526.240709.6326523.133330.6666671.481482.0975647.71833.33333
0.930.860.240.270.080.260.180.280.10.160.250.150.080.540.240.090.230.050.140.211.260.15
692606
genus

0000000100011200010221
0.010.020.020.040.010.010.020.030.01
species
150286

180230
species
4935431164025400401241256
0.910.730.130.070.020.140.10.050.150.080.030.010.020.050.010.020.060.07

140321134102006035324538
0.020.080.010.070.050.090.080.110.030.010.040.010.050.030.050.040.020.050.060.040.11
species
317878

0.040.10.130.030.150.020.130.120.040.450.210.030.120.010.040.051.140.04
023504061076162022391341053
317858
species

0.220.120.140.040.020.040.040.020.060.020.040.020.02
31.501.50.50.500.50.50000101.50.51000.500.5
0.110.060.070.020.010.020.020.010.030.010.020.010.01
860343
no rank

63031101100002031200101
0.110.060.070.020.010.020.020.010.030.010.020.010.01
1241371
species

family
548685
0.060.030.020.020.020.010.030.040.030.020.080.010.01
100.3333330.33333300.66666700.33333300000000.3333331110.66666720.3333330.333333
0.180.090.060.060.060.030.090.120.090.060.240.030.03

1.500.50.50100.500000000.51.51.51.5130.50.5
0.120.060.040.040.040.020.060.080.060.040.160.020.02
genus
0.060.030.020.020.020.010.030.040.030.020.080.010.01
548686

30110201000000013332611
0.060.030.020.020.020.010.030.040.030.020.080.010.01
261939
species

0.690.891.220.771.151.310.731.0510.781.450.91.132.451.21.80.580.870.720.90.770.790.71
10292
family
2.923.574.973.084.859999999999995.469999999999992.944.409999999999994.113.286.129999999999993.744.6810.254.87.499999999999992.43.583.013.643.233.233.09
0.5102740.6414570.4989940.3149351.010291.926870.3231290.7029480.5742090.5457321.008161.374331.2884629.6010.7604175.538670.9791672.315640.8205980.750.8916410.730650.640777

subfamily
0.350.540.80.330.720.670.320.540.540.420.640.520.730.160.970.740.220.250.280.350.250.380.25
10357
1.161.662.470.982.432.21.051.871.691.382.071.622.310.512.912.380.720.760.931.080.851.230.9
0.870691.078310.7732790.51.584363.359090.4285711.229950.8934910.739132.120772.765432.333330.6470591.182137.596640.7222220.8552630.9784951.027781.364711.317070.888889

0.210.270.320.190.230.220.140.070.190.090.320.320.530.060.140.130.070.110.050.070.080.110.08
10358
genus
0.420.540.630.370.460.460.320.140.370.180.650.641.060.110.280.310.140.240.090.140.170.220.18
1.642862.462961.3015911.456524.043480.968750.8571431.351350.8333335.184626.254.745280.9090910.51.354841.071431.1250.6666670.7142861.0588210.722222

no rank
342596
0.010.0500.0400.020.010.010.01
0.020.100.0800.040.020.020.02
000000.5100000000200.500.500.50

1919083
species
00000120000000040101010
0.010.0500.0400.020.010.010.01

57213120240131411310140021
0.090.150.060.020.070.10.010.050.090.250.280.450.020.070.010.050.030.01
species
50290

0.020.030.020.010.020.010.02
0111010000100001002
1535247
species

22111200210211123201232
0.040.040.030.020.040.020.050.030.040.040.010.070.020.030.020.010.030.030.03
species
188763

10359
species
03424830122010021012231
0.060.130.050.10.070.070.020.060.040.040.020.010.010.020.030.030.01

20001201101002042112012
0.040.020.020.020.020.020.010.040.020.010.010.020.010.03
50292
species

47929
species
20230000000001010000010
0.040.060.070.010.0100.010.010.01

no rank
0.040.040.130.070.140.220.040.150.160.060.10.060.410.430.030.010.070.030.04
686603
0.080.080.250.130.280.440.070.290.320.120.20.120.820.870.060.020.140.060.08
110.960.4615382.214298.090910.2857141.13793212.51.500319.56320.50.501.5714300.50.875

10397
species
222152003525300643110101
0.040.040.060.020.120.170.010.070.120.060.10.060.410.390.010.010.010.01

species
33706
0.060.020.030.020.020.020.050.010.010.010.03
00210411100000051001012

species
1608255
00011202100000001004011
0.020.020.020.050.020.010.050.010.01

0.3333330.9090910.77419400.3636361.083330.2857142.666670.3636360.66666710.5555560.3333330.50.51.21.187501.545450.400.40.666667
0.060.220.310.110.120.140.60.110.180.180.180.120.080.140.150.160.220.050.050.09
genus
548689
0.020.080.130.050.040.050.20.050.060.060.060.040.030.070.060.060.080.020.020.03

0.040.120.120.040.080.090.40.040.120.120.120.080.040.060.070.120.020.020.06
0.51.5100.51.6250.44444440.511.50.8333330.50.7501.51.4285702.50.500.51
0.020.060.060.020.040.050.20.020.060.060.060.040.020.030.030.060.010.010.03
548912
no rank

0.060.060.020.030.020.20.020.060.060.040.040.010.030.030.060.01
0320141812321103305001
548914
species

0.020.010.020.020.010.010.010.03
10000110000102001001002
species
768738

146015
species
0.020.060.020.020.010.070.030.030.020.010.01
0120100010000113302101

0.040.160.260.090.10.110.090.090.040.240.180.160.070.560.080.020.070.130.190.130.24
0.50.7520.44444411.090910.4444440.4444440.520.7777781.2500.7142861.251.250.50.7142860.9230771.842111.384623.20833
genus
0.020.080.130.050.050.050.050.050.020.120.090.080.040.280.040.010.040.070.10.060.12
10365

species
28304
0.020.020.020.020.020.020.020.020.010.070.010.020.060.030.01
1101021100110110102521

1261657
species
0100020000100201012102
0.020.020.020.010.010.010.020.010.02

0.040.130.020.050.020.020.020.020.120.050.060.010.210.030.020.020.020.040.09
0241221114230233022238
species
10366

genus
0.070.060.10.020.260.140.050.070.120.090.080.160.040.070.090.060.090.090.070.10.10.11
40272
0.9523810.50.4090910.3333333.184214.30.363636110.6250.90909100.81.555560.51.565221.166671.777781.8571422.666672.071431.8
0.210.120.220.060.760.40.110.210.310.240.220.40.090.140.230.120.180.210.210.30.280.3

0.120.060.040.480.240.040.140.140.120.120.160.020.080.020.020.060.140.20.160.16
1.500.50.5570.51.51.511.5011020.50.51.5343.53
no rank
431037
0.060.030.020.240.120.020.070.070.060.060.080.010.040.010.010.030.070.10.080.08

3011101413323022041136876
0.060.030.020.240.120.020.070.070.060.060.080.010.040.010.010.030.070.10.080.08
1940555
species

species
32604
0.020.030.0200.01
011000000010000001

11101210210024044550012
0.020.020.030.020.020.020.050.030.080.030.040.030.060.060.010.03
10372
species

0100000000000012100001
0.020.070.020.010.01
species
32603

0.470.240.370.340.410.550.60.270.820.180.530.360.364.890.272.160.431.050.460.380.60.40.68
0.3404260.3333330.4864860.2941180.4390241.018180.4666670.2962960.6951220.6666670.4905660.4444440.55555657.53990.2592599.851852.325586.209520.8695650.5789470.8166670.6250.735294
10374
0.150.080.130.120.140.180.210.10.270.060.170.120.121.620.090.710.140.360.160.140.20.140.21
subfamily

genus
0.060.020.060.050.020.010.020.020.020.040.080.020.020.010.010.020.020.090.040.04
548687
0.120.040.120.090.040.020.040.040.040.080.160.050.040.020.020.040.040.180.080.08
0.50.510.4444440.50.50.50.50.5000.510.6010.50.50.511.510.875

0.020.060.020.010.01
1021000000000100001
species
10398

0.020.020.020.020.080.010.020.010.020.050.01
00010010100121020102401
species
35252

138184
species
0.020.020.020.010.010.010.020.03
10000001000101001000122

1100110000000000001022
0.020.020.020.010.010.030.02
1504288
species

no rank
35249
0.030.040.020.050.020.010.020.010.020.010.010.01
0.060.080.040.10.040.020.040.020.040.020.020.02
000.5002.50.50100.5000.50100.510.50.50.5

0.030.040.020.050.020.010.020.010.020.010.010.01
0010051020100102012111
1452540
species

0.50.500.4444440.500.8333330.50.823529010098.3063015.98394.5511.510.50.50.51.28571
0.080.040.090.0400.120.040.170.082.840.040.620.20.560.040.020.040.020.14
548688
0.040.020.050.0200.060.020.080.041.420.020.310.10.280.020.010.020.010.07
genus

10371
species
1101000110000198033102320103
0.020.020.0200.010.020.041.410.020.30.090.280.020.020.04

12657
species
10011020202001011001012
0.020.020.020.050.050.040.010.010.010.010.010.03

0.060.040.030.020.10.120.10.050.120.060.10.060.040.160.070.340.030.040.090.070.050.040.08
10379
genus
0.120.080.060.040.190.240.190.090.240.120.20.120.080.330.140.690.070.070.180.140.10.080.16
0.50.50.50.50.7368421.416670.7368420.4444441.2916710.70.8333330.56.454550.516.30431.142860.7142861.666671.071431.30.751.1875

species
37296
0.020.030.01
10100000000000000001

species
10385
0.020.010.020.070.020.020.01
01000100001000102000021

species
1605972
0.020.020.020.020.020.040.010.01
1000021000111100001

species
154334
00000100000001000101
0.010.010.010.01

0000140130020170350230102
0.020.030.020.070.040.120.320.020.030.010.03
85618
species

00001210202003010003313
0.020.020.020.050.040.020.010.040.040.010.04
species
10381

species
33708
1101242102100102204101
0.020.020.020.050.030.050.020.060.020.010.020.020.050.010.01

000000.33333300000.50.500.33333300.66666700.500.511.50.666667
0.030.040.040.030.060.020.020.060.060.09
genus
0.010.020.020.010.020.010.010.030.030.03
10375

species
106331
0.030.03
0000000000000000000023

0.020.020.010.01
00000000001100000101
45455
species

0.020.020.040.06
000000.500000000.5010000001
species
10376
0.010.010.020.03

0.010.010.020.03
00000100000001020000002
no rank
12509

0.5333330.5384620.4285710.3838381.011491.836880.3928570.5901640.4666670.6914890.7355770.5813950.59.045830.3333331.879311.238811.244441.288891.093751.217820.6049380.85
0.60.780.910.990.871.410.561.220.60.942.080.860.882.40.421.160.670.90.91.281.010.810.8
subfamily
10293
0.190.260.290.330.290.450.20.410.190.30.640.260.280.670.140.340.220.270.280.420.320.270.24

0.040.20.060.250.140.190.150.040.060.610.080.20.220.060.220.10.110.190.080.130.14
0.510.50.6402.428570.7368420.5333330.50.51.180330.50.42.8636400.52.409090.40.8181822.315790.8751.076920.928571
10294
0.020.10.030.120.070.10.070.020.030.290.040.080.110.030.110.050.060.10.040.060.07
genus

10353
species
0.020.020.020.050.020.040.040.010.07
11010020102010000016

species
10310
0.030.040.010.01
001000000020000100001

species
340907
0000001000200400000021
0.010.020.040.0300.030.01

00000000000.50000000.5
0.040.02
342410
0.020.01
no rank

0.020.01
000000000010000001
species
1343901

species
10317
0.010.020.020.020.040.010.010.010.03
00010211000000011000002

species
10298
00000000002108001100001
0.040.020.060.010.010.01

481315
species
0.020.010.010.020.01
00000000001001000121

species
10325
03020601014100003021032
0.060.050.050.040.030.090.020.030.020.010.040.03

species
2003507
0.020.040.01
0.040.080.02
0000.5000000000.5000000000.5

0001000000001000000001
0.020.040.01
332937
no rank

0.010.010.060.01
000000000000020171
species
137443

10000100.4444440010.50.51.5010.5100.50.51.25
0.080.040.090.080.040.080.040.040.020.040.040.020.08
genus
180255
0.040.020.050.040.020.040.020.020.010.020.020.010.04

0.040.020.040.020.010.03
2000000100210000000103
species
50294

10386
species
0000020100001302120111
0.020.020.040.020.020.010.020.010.010.01

0.060.090.040.060.060.060.060.060.190.150.120.060.21
0.33333300.3333330.50.3333330.66666700.33333300.50.33333300000001.263161.3333310.6666671.66667
1232637
0.020.030.020.020.020.020.030.020.070.050.040.020.07
genus

0.500.500.5100.5000.50000000221.512.5
0.040.060.040.040.040.040.10.10.080.040.14
1561225
0.020.030.020.020.020.020.050.050.040.020.07
no rank

10101201001000000044325
0.020.030.020.020.020.020.050.050.040.020.07
species
1561226

0001000001000000002
0.020.030.02
species
702736

genus
180252
0.040.030.10.020.120.050.020.010.070.030.10.10.080.480.240.070.170.020.110.060.040.05
0.50.2857140.6818180.3333331.750.8333330.500.6315790.3333331.133331.21429115.537303.327871.62.266670.21.636361.615380.751.09091
0.080.070.220.060.240.120.040.030.190.090.30.280.161.340.610.150.450.050.220.130.080.11

0.020.020.020.020.020.080.060.10.040.010.060.010.03
100012101000280115105012
species
104388

1100000000100001000001
0.020.020.020.010.01
species
35250

1098912
0.010.030.020.010.010.050.030.090.080.380.120.020.110.010.010.01
no rank
000.50.500.50010.51.777782026.503.6666713.6818200.50.5
0.020.060.040.020.020.10.060.180.160.760.240.040.220.020.020.02

0.010.010.030.010.380.080.020.10.01
00000000010005309280
species
1510155

species
1890678
001101002044000401011
0.030.020.010.050.080.080.040.010.010.01

species
37108
0.010.040.020.050.01
00000100000005020401

10390
species
0.060.10.020.020.010.020.050.020.03
00204200000100000012422

1.066670.80.720.5806451.571433.450.4615381.208330.7777781.302330.9487180.511.538460.50.7272730.8333330.53.037041.461542.411760.7894740.8
0.150.250.250.310.280.60.130.480.180.430.390.20.160.130.280.110.060.040.270.260.340.190.1
0.070.130.130.160.140.30.070.250.090.210.190.10.080.060.140.050.030.020.140.140.170.10.05
10319
genus

0.040.010.0200.01
20001000000000000001
species
10326

0.010.060.050.030.01
0000010201100001
species
10320

species
35246
0011121100012000003021
0.030.020.020.020.020.020.020.080.030.030.01

21000611011100110001321
0.040.020.050.020.020.030.020.020.070.010.010.040.020.01
species
10335

0.060.020.10.10.050.060.030.070.010.010.090.060.090.020.03
002141200203004111085722
species
170325

35244
species
0.040.030.010.020.050.020.010.010.01
02000400002100000111

55744
species
0100000000000000000001
0.020.0100.01

species
10334
0.010.010.020.01
00000100000001000002001

species
10331
00020001101001000001
0.050.020.020.020.010.01

0.020.030.020.010.020.010.01
0001030000010001210001
80341
species

species
10345
0.040.030.020.050.020.10.020.120.020.020.020.020.010.02
0211061414110302000012

family
0.390.270.130.070.330.170.170.150.140.210.190.20.160.180.10.050.070.30.190.210.20.48
944644
2.444441.049380.3846150.3157891.02971.705880.750.51.095240.7142861.103451.133330.51.8518501.38710.750.7142862.067421.963641.380953.33.01399
1.170.810.390.191.010.510.520.440.420.630.580.60.480.540.310.160.210.890.550.630.61.43

2100.5111.5000.50.510.51020020.51.502
0.140.10.040.040.120.040.180.020.060.040.080.080.040.080.10.040.080.020.1
genus
0.070.050.020.020.060.020.090.010.030.020.040.040.020.040.050.020.040.010.05
1513458

42012230011212040041304
0.070.050.020.020.060.020.090.010.030.020.040.040.020.040.050.020.040.010.05
694581
species

2.615381.704550.750.51.833332.277780.7058820.7058821.642861.321.87512.7727302.363641.142861.166673.272733.642862.291676.066674.96721
0.260.440.160.040.480.180.170.170.280.30.270.320.160.220.110.070.120.330.280.240.30.61
no rank
0.130.220.080.020.240.090.090.090.140.150.130.160.080.110.050.030.060.170.140.120.150.31
1513461

0.020.090.020.040.010.010.010.020.040.020.050.010.030.020.030.010.03
14001100001203050131202
1560514
species

1955558
species
0.060.060.020.080.020.020.050.090.090.020.020.050.020.040.030.010.010.140.09
032132124311070023311137

1826170
species
630056212255250121996013
0.110.070.120.060.050.020.050.060.10.10.080.040.010.020.010.10.110.080.18

500.50.50.53.50100.5100.53.500.510.53.51224.5
0.380.060.040.080.120.10.060.080.080.10.020.040.020.160.040.10.080.24
no rank
1513460
0.190.030.020.040.060.050.030.040.040.050.010.020.010.080.020.050.040.12

species
999883
0.190.030.020.040.060.050.030.040.040.050.010.020.010.080.020.050.040.12
100111702012017012172449

0.710.060.210.170.890.390.610.220.280.180.20.140.241.713.142.242.660.160.110.290.290.28
0.5352110.3333330.2857140.2352941.404490.5641030.475410.2727270.2857140.3333330.30.2857140.33333313.1228019.850311.589312.95860.250.4545450.3448280.5517240.535714
0.210.020.060.050.240.10.170.070.090.060.060.040.080.430.790.560.670.050.040.080.090.09
151341
family

0.333333000.3333332.3333310.3333330.333333000.33333300.50.5002.37500.3333330.3333330.58333301
0.120.060.510.140.060.060.060.080.060.240.030.030.120.06
genus
0.040.020.170.050.020.020.020.040.020.080.010.010.040.03
1891713

0.5
0.04
1891734
0.02
species

no rank
928211
1
0.02

1891727
0.01
species
0.02
00000000000000000000.5

862909
no rank
00000000000000000001
0.01

0.01
1891742
species
0.02
00000000000000.5

no rank
1219896
0.01
00000000000001

species
1891728
0.03
000000000000000000001
0.06

no rank
1497391
000000000000000000002
0.03

species
1891717
0.020.170.030.020.020.010.07
0.50003.520.50000.5001004
0.040.340.060.040.040.020.14

1623684
no rank
10007410001002008
0.020.170.030.020.020.010.07

species
1891741
0.01
0.02
00000000000000000.5

no rank
1604875
0.01
00000000000000001

species
1236398
00000000000010000000002
0.040.03

0.02
1891740
species
0.04
0000.5

1286213
no rank
0.02
0001

0000000000000000000.5
0.02
0.01
1891726
species

0000000000000000001
0.01
493803
no rank

000001
0.01
1606502
species

1891721
0.01
species
000000.5
0.02

0.01
000001
no rank
1623687

species
1891720
0.020.01
00000000.50000000000000.5
0.040.02

000000010000000000001
0.020.01
no rank
1286215

0.090.030.060.060.020.04
000.333333000.3333330.33333300.333333000000000.50001
genus
1891715
0.030.010.020.020.010.02

0.060.020.040.04
000.5000.50.500.5
species
1891745
0.030.010.020.02

no rank
1277649
0.030.010.020.02
001001101

0.010.02
0000000000000000010002
species
746831

0.210.040.060.060.040.060.040.090.130.120.080.080.020.040.060.020.060.040.1
1.047620.50.50.3333330.50.8333330.50.4444440.4615380.50.500.50.5011.500.500.3333330.50.8
no rank
1723728
0.090.020.030.020.020.030.020.050.070.060.040.040.010.020.030.010.030.020.05

1970065
species
0.060.020.010.03
30000000001000000000012

00000000000001
0.01
species
1236402

1662286
species
0.030.020.03
0010000011

1273090
species
00000001000000000000001
0.020.01

0.01
000000000000000000001
305677
species

species
1572341
000000001
0.02

0.03
00000000000000003
species
1276187

species
1399914
000000001
0.02

1236401
species
000001
0.01

1561705
species
0.03
0000000001

1904410
species
0000000000000000000001
0.01

00000010000010000000001
0.020.040.01
888441
species

species
1365206
00000001
0.02

1000000000000002001
0.020.020.01
1623686
species

0.040.040.02
0.5000.500000000000000000.5
0.020.020.01
1582477
species

100100000000000000001
0.020.020.01
no rank
928212

species
1965344
01001200001
0.020.020.020.02

genus
1891716
0.020.010.40.750.460.650.020.010.030.01
0.060.031.22.251.381.950.060.030.090.03
00000.3333330.333333000000018.6667027.666718.333317.66670.33333300.33333310.333333

00000.50.50000000000000.500.500.5
0.040.020.020.020.02
species
1891749
0.020.010.010.010.01

0.020.010.010.010.01
00001100000000000010101
no rank
1394033

species
0.40.750.460.650.010.03
1891750
0.81.50.921.30.020.06
000000000000028041.527.526.50.5001.5

no rank
1590650
00000000000005608355531003
0.40.750.460.650.010.03

0.170.040.030.280.10.060.020.040.03
0.7058820000.50.3333330.82142900000.40000.66666700.50100.333333
1891714
0.070.020.010.10.040.020.010.020.01
genus

0.04
0000000.5
0.02
1891757
species

0000001
0.02
no rank
1623685

1891760
0.070.01
species
0.140.02
0000001.5000000000000000.5

0000003000000000000001
0.070.01
1623688
no rank

0.5
0.04
species
0.02
2006158

1762023
no rank
1
0.02

species
1236392
0.02
00000000000000000002

000000000000000001
0.01
1608323
species

0.04
000000000000.5
1891774
0.02
species

no rank
452475
000000000001
0.02

1606501
species
0.020.020.02
100010000001

1606503
species
2
0.04

1891765
0.01
species
000000.5
0.02

no rank
440266
0.01
000001

0.02
1891771
species
0000000000000001
0.04

0.02
0000000000000002
563775
no rank

137992
0.020.020.020.020.020.060.030.010.01
family
0.3333330.3333330000.6666670.33333300.33333300000002.3333301000.3333330.333333
0.060.060.060.060.060.180.090.030.03

genus
0.020.020.020.020.020.060.030.010.01
39743
0.50.500010.500.500000003.501.5000.50.5
0.040.040.040.040.040.120.060.020.02

11000210100000007030011
0.020.020.020.020.020.060.030.010.01
10497
species

family
2.351.841.391.562.832.142.21.662.631.981.631.823.341.594.952.394.322.462.071.992.032.59
10501
8.358513.071191.69672.024257.944510.41733.848192.014735.928812.903853.691613.203951.4480830.69931.7918357.447618.094830.68919.769238.043548.804697.550457.02989
7.816.044.555.369.197.077.185.438.856.765.486.087.0310.745.1415.647.713.678.326.896.46.748.7

2.971.951.431.23.672.252.521.42.621.981.881.71.453.382.365.092.275.242.732.352.632.252.5
20.70037.794874.342664.4333318.277926.68449.503975.3428617.11077.96979.159577.564712.9103464.23373.66525112.5636.65261.417925.139221.204320.205319.706719.108
no rank
1.480.980.720.61.841.121.260.71.310.990.940.850.731.691.182.551.132.621.371.171.321.131.25
455363

0.030.050.040.030.020.030.010.050.0300.050.01000.030.010.030.020.030.050.08
10111210100200003131246
696472
species

0.040.10.020.020.050.10.0100.010.040.010.030.01
25010220005001001003131
1586714
species

0100640002100000032002
0.020.140.030.060.0200.040.020.02
species
1586715

1474867
species
54211210517028943222620769101532186855455444
1.010.440.380.231.210.60.690.221.010.660.490.40.30.50.690.140.270.220.770.680.590.590.59

455364
species
19167111245101476141561604251941583731453340
0.350.330.220.260.30.390.250.350.160.180.270.30.241.140.282.270.791.940.420.390.580.360.53

0.060.040.030.050.150.040.220.120.070.030.060.060.170.040.040.040.040.060.050.060.050.04
32126595313345045354553
species
642926

0.040.030.030.020.020.030.040.210.090.370.060.040.02
021003101102003100305031
1586713
species

0.040.080.060.040.040.20.140.090.120.040.120.080.020.070.020.020.090.020.15
0.510.50.50.5601.50.44444410.50.8333330.50.5001.428570.50.50.8888890.51.66667
genus
0.020.040.030.020.020.10.070.050.060.020.060.040.010.030.010.010.050.010.08
181085

0010000310011000300214
0.030.070.020.020.040.030.020.010.04
species
39420

0.020.040.020.020.10.020.060.020.040.010.010.010.010.020.03
12011120012120100111203
37665
species

genus
0.310.270.250.370.220.270.40.230.380.360.320.390.180.140.140.250.20.260.390.260.210.360.5
181084
1.020.830.851.320.670.861.210.741.231.171.051.250.620.450.490.90.710.971.320.880.691.161.54
0.9901961.216870.6117651.674241.014932.569771.37190.5945950.9105690.8376071.447621.5440.5483871.155560.2857142.844442.323942.824743.060611.761361.246381.775863

1.422541.803570.8666672.326321.511113.745762.049380.8627451.317651.209882.082192.244190.7727271.677420.43.938463.235293.859154.344092.51.791672.5754.44231
0.710.560.60.950.450.590.810.510.850.810.730.860.440.310.350.650.510.710.930.620.480.81.04
no rank
0.310.270.250.370.220.270.40.230.380.360.320.390.180.140.140.250.20.260.390.260.210.360.5
880158

1663209
species
0.050.030.030.110.010.070.0300.010.030.070.020.030.010.020.030.040.12
21000340210004124001249

933845
0.090.020.10.210.020.040.020.050.090.090.110.080.080.050.070.140.110.180.140.110.050.080.05
species
2.50.51.54.50.52.50.5121.53213.50.57.56.5764.523.52
0.180.040.20.420.040.080.040.10.180.180.220.160.160.10.140.280.220.360.280.220.10.160.1

no rank
880159
0.090.020.10.210.020.040.020.050.090.090.110.080.080.050.070.140.110.180.140.110.050.080.05
513915124364271151314129474

0.080.050.010.050.030.110.160.060.050.080.020.070.010.050.020.010.050.020.080.040.19
4202112622213000520416414
species
880162

0.020.030.010.020.020.020.010.010.010.010.010.020.080.05
10100200100102011012074
1663210
species

0.050.130.080.060.030.010.030.060.10.040.060.050.090.010.020.020.0500.030.050.08
26221012413221002140246
1663208
species

880161
species
12125832246802041413626
0.020.040.030.050.120.070.070.050.050.120.110.160.010.040.010.050.150.080.030.07

0.780.730.440.750.850.790.90.791.341.420.720.752.683.020.564.141.942.730.781.030.820.661.33
1.230771.219181.2272731.294124.341771.777781.291141.910451.915491.902784.786672.0522436.10260.62577.188427.407234.43592.141032.456312.365853.909094.07519
181083
0.30.310.220.360.330.340.390.350.560.550.30.340.971.310.211.960.911.270.320.40.340.280.57
genus

0.150.10.020.120.070.060.050.20.290.090.070.510.380.140.190.090.120.070.140.110.070.15
346674
no rank
0.290.20.040.240.150.120.10.40.580.180.141.010.760.280.370.180.240.140.280.220.140.3
1.931031.200.51.833332.533330.50.42.13.396551.777781.53.3267318.460518.756763.333332.51.52.52.454551.571432.76667

1917232
species
4301461158434432198635645
0.070.070.020.110.060.040.040.120.240.080.070.160.310.140.170.070.080.030.070.080.050.07

410001103100810012335216
0.070.030.010.020.020.010.080.050.010.340.070.010.020.040.040.070.030.020.08
380598
species

11210412521014045130416
0.020.030.060.020.040.020.060.130.080.020.060.030.040.050.020.040.010.050.010.08
species
46021

0.020.050.10.070.050.050.060.060.140.030.050.010.060.070.020.080.010.050.120.030.030.09
12332522612009129049237
10506
species

0.070.10.060.220.110.150.170.140.070.140.130.260.160.821.690.661.070.090.040.120.160.21
452941775347134115018778878391416
species
322019

10.500.511.51.50.50.500.503101.522.533.510.51.5
0.080.040.040.120.060.140.080.040.040.040.480.040.060.060.120.140.180.060.040.08
species
381899
0.040.020.020.060.030.070.040.020.020.020.240.020.030.030.060.070.090.030.020.04

21012331101062034567213
0.040.020.020.060.030.070.040.020.020.020.240.020.030.030.060.070.090.030.020.04
no rank
399781

genus
0.170.150.060.070.290.230.070.120.30.030.040.140.040.130.180.10.090.250.090.050.110.2
181086
0.510.450.180.210.870.690.210.360.90.090.120.420.120.390.540.30.270.750.270.150.330.6
32.333330.66666714911.666674.333330.3333330.6666672.333330.333333606.3333342.333337.333332.333331.333333.333335

4.53.511.5613.51.52.56.50.513.50.5909.563.5113.5257.5
0.340.30.120.140.580.460.140.240.60.060.080.280.080.260.360.20.180.50.180.10.220.4
0.170.150.060.070.290.230.070.120.30.030.040.140.040.130.180.10.090.250.090.050.110.2
358403
no rank

97231227351312711801912722741015
0.170.150.060.070.290.230.070.120.30.030.040.140.040.130.180.10.090.250.090.050.110.2
species
251749

genus
346673
0.070.080.10.140.130.070.070.170.020.020.020.040.070.070.010.010.060.130.10.050.080.07
0.140.160.20.280.260.140.140.340.040.040.040.080.140.140.020.020.120.260.20.10.160.14
221.532.541.53.50.500.50.50.550.50.50.52.55.5423.52.5

4436583710111101115118475
0.070.080.10.140.130.070.070.170.020.020.020.040.070.070.010.010.060.130.10.050.080.07
181082
species

0.060.190.020.050.10.020.030.030.080.170.050.020.050.130.160.050.080.23
151340
family
0.240.650.080.160.450.080.120.10.320.670.230.080.180.430.610.20.270.83
0.250.33846200.250.250.3555560.25000.250.20.37501.119400.3478260.250.3333330.3023260.9180330.40.4814810.819277

genus
334211
0.010.01
0000000000000000000.333333000.333333
0.030.03

0.010.01
334212
species
0.020.02
0000000000000000000.5000.5

0000000000000000001001
0.010.01
no rank
392505

genus
0.020.01
325456
00.5000000000000000000000.5
0.040.02

01000000000000000000001
0.020.01
species
10567

00000000000000.333333
0.03
936058
0.01
genus

0.02
00000000000000.5
1176150
0.01
species

936059
no rank
0.01
00000000000001

000000000000000000000.33333302
0.030.24
1513240
0.010.08
genus

species
1513249
0.010.08
0.020.16
000000000000000000000.503

00000000000000000000106
0.010.08
no rank
1081054

0.01
333921
genus
0.02
0000000000000000000.5

0000000000000000001
0.01
species
10561

00000000000000000000.333333
0.03
genus
0.01
333918

species
333919
0.01
00000000000000000000.5
0.02

00000000000000000001
0.01
333920
no rank

325455
0.020.020.050.020.030.010.020.090.010.010.030.060.040.07
genus
0.3333330.333333000.3636360.3333330000.33333300.33333302.2800.33333300.3333330.3333331.1666700.9090910.736842
0.060.060.110.060.090.030.060.250.030.030.090.180.110.19

species
1175851
0.04
00000000000000000000001.5
0.08

no rank
931210
00000000000000000000003
0.04

0.02
0000000000000000000.5
species
1961678
0.01

1420545
no rank
0.01
0000000000000000001

0000000000000000000.5
0.02
species
0.01
1513260

0000000000000000001
0.01
no rank
1070408

0.040.06
00000.500000.5
species
1175848
0.020.03

0000100001
0.020.03
no rank
338323

0000000000000000000001
0.01
1513265
species

0.02
00000000000000000000000.5
species
1513261
0.01

00000000000000000000001
0.01
1070410
no rank

000000000000000000.5
0.02
species
1513264
0.01

no rank
1315259
0.01
000000000000000001

species
0.010.060.010.03
1513256
000000.500000004.500.5000001.5
0.020.120.020.06

0000010000000901000003
0.010.060.010.03
1055684
no rank

0000000000000000000.5
0.02
species
0.01
1513262

1070417
no rank
0.01
0000000000000000001

0.040.040.1
00.50000000000.500000002
0.020.020.05
333928
species

0.020.020.05
01000000000100000004
no rank
40538

00000000000001
0.02
species
0.01
1513259

0.01
00000000000002
no rank
909330

0.02
00001
species
1513263

species
0.010.01
1175849
00000000000000000000.5
0.020.02

0.010.01
00000000000000000001
no rank
915426

species
1513257
0.020.010.010.01
0.500000.500000000.5000000000.5
0.040.020.020.02

10000100000001000000001
0.020.010.010.01
746832
no rank

0.060.030.060.03
00.3333330000.3333330.3333330000000000000000.333333
0.020.010.020.01
334202
genus

00.50000.50.50000000000000000.5
0.040.020.040.02
334204
0.020.010.020.01
species

0.020.010.020.01
01000110000000000000001
no rank
28311

0.03
0000000000000000.333333
0.01
1513239
genus

0.02
0000000000000000.5
species
0.01
1513248

0.01
0000000000000001
no rank
759701

genus
0.02
325454
0.06
0000.333333

no rank
470213
0.02
0.04
0000.5

species
996651
0.02
0001

genus
936057
0.01
0.03
00000000000000000.333333

00000000000000000.5
0.02
species
0.01
1176146

no rank
291589
0.01
00000000000000001

0.01
935641
genus
0.03
0000000000000000.333333

0.02
0000000000000000.5
0.01
1177767
species

445217
no rank
0.01
0000000000000001

genus
0.04
934800
000000000000.666667
0.12

0.08
000000000001
1177759
0.04
species

464980
no rank
000000000002
0.04

000000.3333330000000000.333333
0.030.03
genus
0.010.01
325462

0000000000000000.5
0.02
no rank
1104916
0.01

0000000000000001
0.01
species
1472717

0.02
000000.5
species
0.01
1513272

0.01
000001
1104917
no rank

00.8333330000.375000000.33333300.500000.714286000.50.333333
0.120.080.060.040.070.020.09
333750
0.060.030.020.010.030.010.04
genus

0000000000000000000.50000.5
0.020.02
species
337048
0.010.01

333763
no rank
00000000000000000010001
0.010.01

species
333766
02000000000000000020011
0.040.020.010.01

species
333754
0.010.01
0.020.02
000000.500000000000000000.5

10600
no rank
00000100000000000000001
0.010.01

0.02
00000000000000.5
species
337039
0.01

333759
no rank
00000000000001
0.01

10570
species
010001
0.020.01

species
337043
00000000000001
0.01

0.02
333757
species
0.04
000000000000.5

0.02
000000000001
no rank
10620

000000.5
0.02
0.01
337049
species

333764
no rank
000001
0.01

genus
0.01
934799
0000000000000000000.333333
0.03

1961786
0.01
species
0.02
0000000000000000000.5

1336600
no rank
0000000000000000001
0.01

genus
934802
0.02
0.06
00.333333

1177757
0.02
species
00.5
0.04

01
0.02
338903
no rank

000000000000000000000.666667
0.09
0.03
935650
genus

000000000000000000001
0.06
0.03
1177762
species

no rank
369584
000000000000000000002
0.03

00.333333
0.06
936056
0.02
genus

1177764
0.02
species
0.04
00.5

no rank
446138
0.02
01

genus
333922
0.020.020.01
00000000000001000000.6666670.333333
0.060.060.03

species
0.02
337051
0.04
00000000000001.5

no rank
333923
0.02
00000000000003

0.040.02
000000000000000000010.5
species
334209
0.020.01

000000000000000000021
0.020.01
247269
no rank

000000.666667
0.06
0.02
325459
genus

0.04
000001
1777073
0.02
no rank

0.02
000002
1434070
species

0.3333330.3333330000.55555600000000.7500.6666670.3333330.50.3333331.6666700.3333330.333333
0.120.060.090.120.060.030.10.060.180.030.03
no rank
333774
0.040.020.030.040.020.010.040.020.060.010.01

333938
0.020.020.020.040.020.010.010.010.01
no rank
0.040.040.040.080.040.020.020.020.02
0.50.5000100000001.125010.50.50.50000.5

1
0.02
706526
species

species
1887214
0000020000000000001
0.020.01

01000000000003021100001
0.020.020.020.010.010.01
species
1335477

00000000000002
0.01
species
706525

species
1887215
0.01
00000000000001

173087
0.020.010.020.010.060.01
no rank
0.500000.50000000000010.52.500.5
0.040.020.040.020.120.02

10566
species
0.020.01
1000000000000000001

1070409
species
0.010.06
00000100000000000005

1070413
species
000000000000000002
0.02

1195796
species
0.01
0000000000000000000001

00000000000.5
0.04
genus
0.02
325457

0.02
00000000001
species
122291

10442
1.341.021.4611.430.81.180.961.30.720.971.011.231.81.313.441.322.821.351.871.11.281.16
family
4.409999999999993.474.743.234.599999999999992.743.913.194.242.533.173.334.016.649999999999994.1613.224.9499999999999910.574.469999999999996.089999999999993.629999999999994.019999999999993.91999999999999
0.9569160.4956770.6097050.5944270.810871.010950.8363170.5454550.6886790.3043480.7129340.4864862.5411518.89920.75721243.85713.957625.45411.102912.656811.074381.711440.808673

no rank
35255
0.020.020.020.020.020.080.010.040.010.060.350.140.070.04
0.040.040.040.040.040.160.020.080.020.120.70.280.140.08
0.5000.50.51000000.5110200.52.5145.531.5

0.020.020.020.020.020.080.010.040.010.060.350.140.070.04
1001120000012204015281163
1675865
species

0.170.120.110.040.060.140.140.150.040.10.020.140.060.090.160.250.060.050.17
0.7058820.33333300.3636360.50.3333330.85714300.2857140.40.50.400.50.501.50.8888891.3751.2810.41.05882
558018
0.070.050.050.020.020.070.050.060.020.040.010.070.030.050.070.110.030.020.07
genus

0.040.040.040.020.080.060.040.020.060.020.06
0.50.500.500.5000.50.500.50000000100.51
1233523
0.020.020.020.010.040.030.020.010.030.010.03
no rank

204507
species
0.020.020.020.010.040.030.020.010.030.010.03
11010100110100000002012

0.020.020.010.050.030.010.020.050.050.030.01
1100012001000100024421
species
111874

species
249151
0.040.010.020.020.010.020.010.020.020.070.030.020.020.030.04
20011010001100103212003

2.11.572.791.611.821.351.691.842.061.591.531.442.381.281.462.380.831.521.872.441.561.71.51
1.604760.7006370.9390681.006210.8516481.525931.426040.7989130.9708740.4465411.16340.6254.180672.039061.438363.865551.108431.960531.641712.0327913.088241.03974
558016
0.910.661.230.720.780.560.740.790.880.630.690.621.070.540.690.940.330.640.81.030.660.810.64
genus

1010110200000101221
0.020.030.030.010.050.010.010.020.020.02
74660
species

207830
species
11301802111001110102013
0.020.020.10.020.070.050.020.030.020.010.070.010.010.030.010.04

species
78219
20000000000000000002010
0.050.010.020.010.020.020.010.030.020.01

204440
species
20000000010100000032010
0.050.010.020.010.050.020.020.040.030.020.01

0.020.010.050.020.040.050.070.020.020.010.020.040.010.01
01000121000207122102311
59376
species

101850
species
00220400101003002284221
0.060.050.030.020.020.0200.020.020.090.050.030.020.01

species
51313
001002010000010410101
0.030.020.020.010.040.010.010.01

1580580
species
0.020.020.010.020.020.020.020.010.020.010.010.01
01001110100100020102111

species
320432
00000011113100033321203
0.020.020.020.030.060.020.030.030.040.020.020.030.04

00242101102315507000001
0.060.090.050.010.020.020.040.060.610.040.060.01
10455
species

12210001001001
0.020.040.060.020.020.020.01
262177
species

00101000003012001001203
0.030.020.060.040.010.010.010.030.04
1307957
species

000302300010040140022111
0.070.020.070.020.030.1300.020.020.010.010.01
70600
species

species
224399
11100201301102041101001
0.020.020.050.020.010.020.070.020.020.010.040.010.010.010.02

species
332054
0.190.040.160.020.070.020.020.020.070.030.10.040.070.030.060.070.010.17
102513211315200130055116

208973
species
00011200000101002100102
0.020.020.020.020.010.020.010.010.03

species
1307954
00100161400004002021420
0.030.010.150.020.090.030.020.020.010.050.020.01

species
307467
41112321101103001115271
0.070.020.030.020.050.030.050.020.020.020.020.020.020.010.010.010.060.030.080.01

0.020.020.020.030.020.010.010.020.040.010.050.040.030.04
11010400000101012314333
490711
species

2000100010012202224011
0.040.020.020.020.080.010.020.020.020.050.010.01
species
28288

3312021011110000001301
0.060.060.030.050.020.020.020.030.020.020.010.040.01
10454
species

1307956
species
1011120201120302040208
0.020.030.020.020.020.050.030.020.040.020.020.050.020.09

0.50.5001000100000.5010.52.5411.510.5
0.040.040.10.10.020.040.020.120.20.040.080.040.02
species
0.020.020.050.050.010.020.010.060.10.020.040.020.01
1207438

0.020.020.050.050.010.020.010.060.10.020.040.020.01
11002000200001021582321
no rank
1592335

species
1962501
0.060.020.060.090.050.010.030.020.010.010.020.040.010.020.07
31242100010003011203125

0.030.020.020.010.010.0100.020.03
1000010000000001002
species
268591

01003023104011500034122
0.020.070.050.070.020.080.040.010.350.030.060.010.020.03
species
46242

0100000000000000002011
0.020.020.020.010.01
species
271108

1010010011110102010141
0.030.030.010.040.030.020.020.010.020.010.010.050.01
10449
species

307456
0.040.020.050.060.010.010.030.040.03
species
1000000011000000.50.501.501.501
0.080.040.10.120.020.020.060.080.06

20000000220000011030302
0.040.020.050.060.010.010.030.040.03
46015
no rank

0.030.020.020.030.010.010.01
00000000011104000101001
161494
species

1001010000210010012014
0.020.020.010.040.020.070.010.020.010.04
species
208013

no rank
0.170.260.320.190.230.190.230.320.230.260.110.20.240.170.070.440.120.220.170.380.170.080.19
745176
0.350.510.640.380.460.390.470.630.460.520.230.40.480.340.140.890.240.440.330.760.340.150.39
0.7142861.058821.1250.9210531.217392.025641.914891.206351.521740.5769230.9565220.61.166673.823530.56.9101124.431821.272733.9210510.7333331.20513

0.050.110.020.040.020.020.030.020.020.090.140.020.150.010.010.03
02301401111101301621201102
1850906
species

1642929
species
10001100010130011000004
0.020.020.010.030.020.120.010.010.05

species
31507
0000010000000101002
0.010.020.020.010.010.010.010.020.03

10004051523212000008121
0.020.020.10.140.020.120.060.060.040.040.010.10.010.020.02
542343
species

0.020.060.030.050.020.010.010.010.050.010.03
13000402100000011014012
species
1367203

species
1563660
0.050.020.020.080.020.070.030.010.010.150.02
00020000100123130111202

22310200010100000300302
0.040.040.10.020.020.030.020.040.040.03
species
1346829

02010124010101040111112
0.040.020.010.050.10.030.020.010.040.010.010.010.010.010.03
1367205
species

species
566270
0.040.020.070.020.060.050.050.040.030.020.020.010.180.030.050.020.030.010.01
210317221111010204042211

species
307461
0.030.050.010.030.010.030.040.03
001021000100000100332

001000020001010161001
0.030.010.050.010.010.020.010.010.050.0100.01
1592576
species

121001010001010000202
0.030.040.030.010.020.010.020.010.020.03
species
745177

0110111120100100021007
0.020.030.020.010.020.020.050.020.010.020.010.08
species
74320

genus
0.020.020.020.070.030.090.03
558019
0.50.500000.501.500002005.500001.5
0.040.040.040.140.060.180.06

130556
species
0.020.020.020.070.030.090.03
11000010300004001100003

genus
558017
0.320.290.220.210.590.20.340.170.30.030.270.340.081.210.552.470.882.120.430.380.270.350.41
0.720.720.490.471.270.490.860.390.60.070.630.740.243.471.257.322.566.120.970.830.630.791
0.9583330.7777780.551020.5106381.685041.326530.8372090.6923081.1166700.7301590.8918920.33333335.42360.78477.927626.20743.46081.381441.32531.17461.392411.3

0.020.030.020.020.020.070.020.020.040.010.010.020.020.070.060.040.03
1011101310120101226532
10464
species

0.030.020.020.010.020.020.010.01
001111000011000000101
283675
species

0.120.290.120.140.190.20.390.130.040.160.120.162.10.284.731.583.790.250.160.180.210.34
0.51.034480.50.8571431.210531.551.384620.3076920.501.250.8333330.558.20950.5120.52642.329169.88391.320.68751.51.428571.79412
no rank
342107
0.060.150.060.070.10.10.20.070.020.080.060.081.050.142.360.791.90.130.090.090.110.17

307444
species
0.030.010.070.010.010.010.04
0010000000000111001104

020000000030113112518914730425
0.040.060.040.930.072.270.751.80.030.050.020.07
364745
species

11120110000001010412001
0.020.020.030.050.010.020.010.010.010.050.010.020.01
species
252584

1000000000000000000203
0.020.020.03
1352289
species

species
307454
0.020.020.020.10.020.020.040.030.020.020.010.010.030.04
11010041100204020211203

species
1136024
030003200000010001
0.060.030.050.010.01

00001310000003011110011
0.020.030.020.010.020.010.010.010.010.010.01
1675862
species

00003401001115054041103
0.070.030.020.020.020.040.040.050.030.050.010.010.04
1675863
species

0.020.020.010.010.010.010.03
10000010000001000101012
98383
species

0.020.10.010.020.020.040.010.010.010.030.05
010041101002020110304
species
170617

359919
species
0.020.020.040.020.070.010.110.010.020.050.01
00000010100203110912051

01011000301001021200002
0.020.020.020.070.020.010.020.010.020.03
species
28289

262175
species
0.050.010.01
0000200000000000011

0.020.170.050.020.040.040.010.010.010.010.030.050.010.010.03
00017600102202011134112
species
192584

166056
species
10000000001000001034304
0.020.020.010.030.050.040.05

489830
species
0.070.020.030.0200.010.020.010.010.020.010.030.010.030.06
31101000000001000112034

51677
species
21001001000100100022020
0.050.020.0200.020.010.020.010.0200.070.010.020.030.010.020.01

0.060.060.020.050.010.020.060.060.070.010.010.050.040.010.03
3301210000130811104313
36355
species

0.040.060.020.020.010.050.090.040.020.020.210.050.030.020.050.03
20211120402103354020052
35254
species

000000.333333000.333333000.3333330000000.333333
0.030.060.060.03
0.010.020.020.01
10472
family

0.020.040.040.02
000000.5000.5000.50000000.5
genus
0.010.020.020.01
10473

0.010.020.020.01
0000010010010000001
species
46014

family
0.02
324686
00.333333
0.06

0.04
00.5
0.02
1298634
genus

species
1032474
01
0.02

4.212.051.922.676.133.233.182.785.253.182.361.622.453.092.391.762.741.024.184.545.585.663.75
0.9429930.5317070.43751.280911.829720.635220.575541.377141.264150.7627120.4691360.4408162.233010.234310.8181821.288320.5588241.483252.599121.935483.217311.26933
0.860.460.380.581.280.670.660.621.10.720.510.320.530.640.480.370.560.210.870.951.121.120.76
10486
family

180169
0.050.050.080.030.050.060.150.060.040.040.070.030.050.010.070.020.030.010.01
no rank
0.10.110.160.060.090.120.30.120.080.090.140.060.10.020.140.040.060.020.02
00.800.9090911.3125200.4444440.8333331.71.5102.333330.51.52.10.53110.50.5

0.010.010.010.020.010.0300.010.010.01
000000010100010011
1887316
species

02023401243205135062211
0.040.050.070.030.020.050.120.060.040.040.070.030.040.070.020.030.010.01
1465751
species

1.004590.7361110.5373130.6176471.364382.592780.8727270.8333330.9365850.350.4050630.5789470.5555562.355030.1735541.031581.641380.7968751.528213.698922.561641.975811.61472
2.180.720.670.683.651.941.651.22.050.60.791.141.081.691.210.951.450.641.952.793.652.482.31
subfamily
2017757
0.540.190.160.190.90.490.440.320.520.150.190.260.280.430.280.240.340.160.470.730.90.590.57

0.040.060.10.040.040.020.020.020.040.080.10.04
0.500001.5100.500.5000.500.500.511.521
10491
0.020.030.050.020.020.010.010.010.020.040.050.02
genus

1000032010100101012342
0.020.030.050.020.020.010.010.010.020.040.050.02
species
345201

1.60.530.510.492.751.391.110.881.490.450.560.880.81.240.930.691.110.461.441.982.651.851.74
1.3562510.7058820.8571431.810913.553961.207211.136361.275170.4666670.5357140.750.753.201610.2258061.40582.144141.086962.041675.151523.452832.627032.14368
0.520.190.160.190.90.460.390.320.490.150.170.260.280.420.280.230.340.150.450.690.840.570.57
10487
genus

1.737370.8076921.028570.2857142.531795.093021.233330.9090911.477270.6666670.6285711.066671.166674.175680.3230771.634152.61.61292.914897.530434.953493.669422.52381
0.990.260.350.211.730.860.60.440.880.270.350.60.480.740.650.410.70.310.941.151.721.211.05
0.430.10.160.090.780.390.270.20.370.120.130.240.240.340.280.180.280.150.40.560.750.50.45
327984
no rank

5030462341113704567111134
0.110.10.120.050.060.080.090.030.020.030.120.050.040.040.070.080.140.140.040.06
345585
species

345198
species
31007800101128133125894
0.060.030.010.190.070.010.010.040.020.030.080.060.070.030.030.010.030.060.110.10.07

0.120.060.040.180.10.10.080.220.060.080.080.060.060.060.160.020.060.040.360.120.12
1.500.50.51.5310.520.51101.501.550.510.56.52.52
species
345195
0.060.030.020.090.050.050.040.110.030.040.040.030.030.030.080.010.030.020.180.060.06

0.060.030.020.090.050.050.040.110.030.040.040.030.030.030.080.010.030.020.180.060.06
3011362141220303101211354
no rank
68347

species
345199
0.060.060.020.070.030.020.040.080.030.070.010.070.020.110.010.030.160.08
0.120.120.040.140.060.040.080.160.060.140.020.140.040.220.020.060.320.16
1.51.500.51.51.5000.5012020.50.54150.517.53

72201
no rank
0.060.060.020.070.030.020.040.080.030.070.010.070.020.110.010.030.160.08
3301330010240411821012156

species
1301280
0.090.010.030.020.170.160.10.060.110.060.020.060.040.080.030.040.010.010.090.290.20.060.13
501171942421311104118231559

300063200000013144143764
0.060.010.150.030.060.010.010.090.070.040.040.010.060.040.10.070.06
1301279
species

176652
species
0.090.080.090.120.070.120.120.120.030.040.020.040.080.050.070.050.140.090.070.12
5404585551211110680411769

1.170.770.871.31.040.560.870.871.981.560.940.080.840.670.560.380.630.151.220.760.752.040.66
1.521370.6233770.5517242.230770.9038461.357140.6666670.6436782.631312.115381.382980.250.5714294.044780.50.9736841.492060.3333332.295081.894741.853336.519611.54545
subfamily
2017756
0.320.220.220.340.30.150.220.250.530.420.260.020.240.170.140.10.170.040.330.20.190.520.17

0.040.180.30.140.070.020.150.140.070.060.130.160.030.020.050.020.020.02
0.50.33333310.1428570.2857140.50.6666670.8571430.2857140.50.46153800.50.66666700.50.400.5000.50.5
genus
0.020.080.10.060.040.010.050.070.040.030.070.080.010.010.020.010.010.01
10492

10493
species
01000000001001
0.020.020.01

0.040.20.020.10.02
00.51.500010000000000.5
0.020.10.010.050.01
327987
no rank

0.020.10.010.050.01
01300020000000001
638660
species

84739
species
0000000100001
0.010.010.0200.04

0.010.020.010.010
00000000000
100217
species

0.010.02
00000000001
species
265294

species
262968
0.020.020.020.020.010.050.020.030.020.040.010.010.010.010.010.01
11011102111011011010011

00.5000.510000000000001
0.040.040.040.04
genus
308906
0.020.020.020.02

180170
species
0100120000000000002
0.020.020.020.02

0.810.330.350.820.630.350.50.481.381.080.550.060.440.470.420.260.410.110.830.560.561.50.47
2.172841.212120.5142863.51221.428572.028570.960.9166673.760873.027782.254550.3333330.9090915.72340.6666671.384622.24390.4545453.313252.571432.482148.862.14894
10494
0.30.130.130.280.240.130.170.170.490.390.190.020.160.160.140.090.150.040.30.20.190.510.16
genus

42114501431011025062131
0.070.040.030.020.10.040.010.020.090.090.020.040.010.020.040.070.020.010.030.01
species
36363

no rank
0.220.080.10.260.140.090.160.150.40.30.170.020.120.150.140.070.110.040.230.170.180.480.15
345690
3.3636420.7894745.51.724142.833331.51.448286.037553.588240.51.58.9333312.133333.272730.7142865.065224.117653.8333313.753.33333
0.440.160.190.520.290.180.320.290.80.60.340.040.240.30.280.150.220.070.460.340.360.960.3

1898060
species
44211273314108102046111109333
0.070.080.060.260.050.060.070.070.330.30.150.020.010.040.050.010.130.120.120.360.04

256729
species
8010433330103192472945118
0.150.030.10.030.090.070.070.020.120.140.140.040.060.020.10.050.060.120.11

0
3.382.632.392.713.462.974.543.693.43.022.772.431.792.712.962.021.513.22.463.033.454.143.14
